# Yeast population dynamics in Brazilian bioethanol production

**DOI:** 10.1101/2022.10.31.514616

**Authors:** Artur Rego-Costa, I-Ting Huang, Michael M. Desai, Andreas K. Gombert

## Abstract

The large scale and non-aseptic fermentation of sugarcane feedstocks into fuel ethanol in biorefineries represents a unique ecological niche, in which the yeast *Saccharomyces cerevisiae* is the predominant organism. Several factors, such as sugarcane variety, process design, and operating and weather conditions, make each of the ∼400 industrial units currently operating in Brazil a unique ecosystem. Here, we track yeast population dynamics in two different biorefineries through two production seasons (April to November of 2018 and 2019), using a novel statistical framework on a combination of metagenomic and clonal sequencing data. We find that variation from season to season in one biorefinery is small compared to the differences between the two units. In one biorefinery, all lineages present during the entire production period derive from one of the starter strains, while in the other, invading lineages took over the population and displaced the starter strain. However, despite the presence of invading lineages and the non-aseptic nature of the process, all yeast clones we isolated are phylogenetically related to other previously sequenced bioethanol yeast strains, indicating a common origin from this industrial niche. Despite the substantial changes observed in yeast populations through time in each biorefinery, key process indicators remained quite stable through both production seasons, suggesting that the process is robust to the details of these population dynamics.

## INTRODUCTION

Fuel ethanol is used throughout the world to power light vehicles, either on its own or, more commonly, mixed with gasoline for increased octane rating^1^. Brazil is the second largest ethanol producer in the world, surpassed only by the United States, and accounts for roughly 30% (or 31.66 billion liters predicted for 2022) of the world’s fuel ethanol production^2^. While American ethanol is mostly corn-based and requires enzymatic hydrolysis of starch prior to fermentation by the yeast *S. cerevisiae*, most of Brazil’s ethanol is produced from sucrose, glucose, and fructose-rich sugarcane products which can be directly fermented.

The Brazilian process is also unique in that it maintains a very large population of yeast in non-aseptic conditions throughout the 8-month-long sugarcane harvesting season^3–5^ (Fig. 1A). The yeast cells are recycled at every ∼12 h fed-batch fermentation-holding-centrifugation-treatment cycle, allowing for large inocula and short turnaround times. Acid wash and antimicrobials serve to control the ever-present bacterial contamination, which competes against yeast for carbon, but also affects fermentation in ways that are not completely understood^6,7^. These practices are key to the high efficiency of the sugarcane-ethanol industrial process and drastically lower greenhouse gas emissions in comparison to corn-based ethanol^8,9^. However, inconsistencies in fermentation performance associated with cell recycling remain a costly challenge and point to microbiological routes for process improvement^3,7,10^.

**Figure 1.**
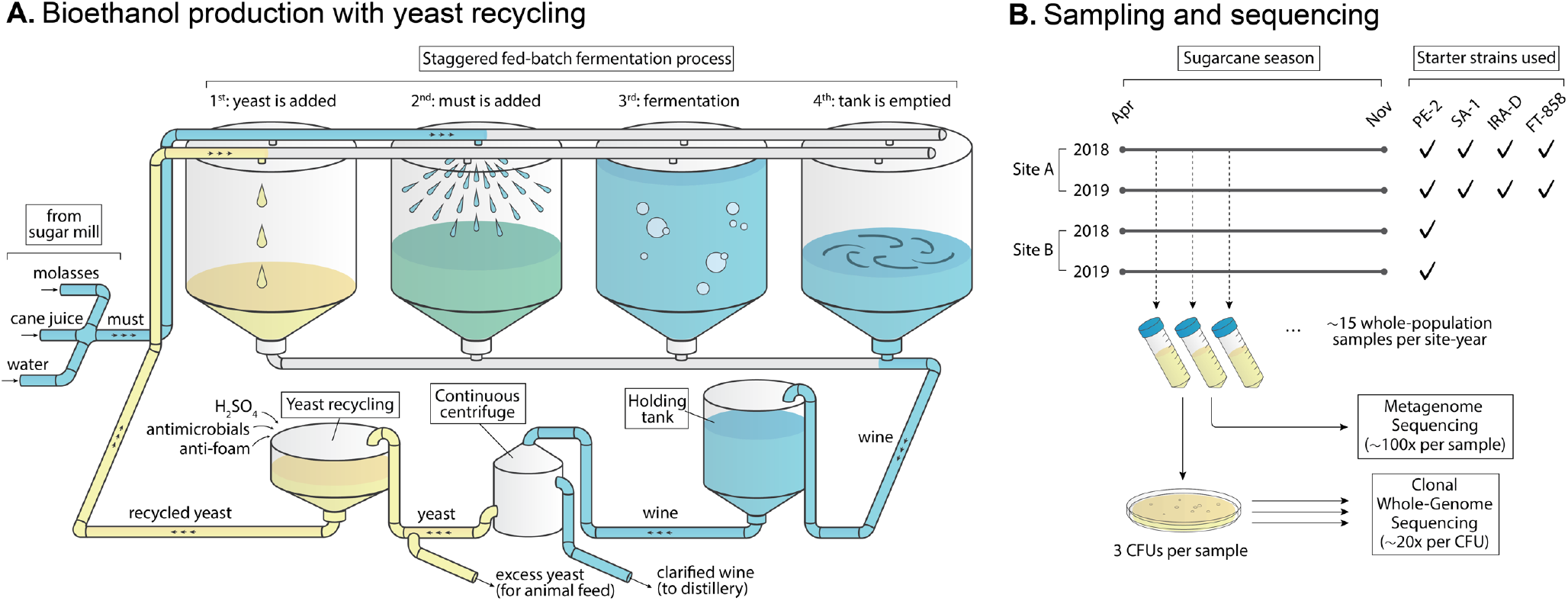
Schematics of the fermentation process and sequencing strategy. **(A)** A large population (∼10^17^ individuals) of the yeast *S. cerevisiae* is maintained over the course of an eight-month-long fermentation season. Yeast ferments must, a mix of molasses, sugarcane juice and water, to produce ethanol in a fed-batch process that takes ∼8h and runs in a staggered parallel fashion across several fermentors (8–16 in any one plant, each with a ∼500,000 ℓ capacity). The fermented broth (wine) from different fermentors is loaded into a single holding tank, which continuously feeds a centrifuge for separation of the yeast from the liquid fraction. Holding tanks are larger than fermentors themselves and allow for mixing between batches. The yeast cells are then treated with chemicals to control for bacterial growth and are later reused in the process. The yeast population grows by ∼10% every 12h, leading to approximately 66 generations over the course of an ∼8 months fermentation season. The season is started with selected industrial strains which are commercialized by yeast suppliers. **(B)** We collected whole-population samples of the yeast used for fermentation through two seasons (2018 and 2019) in two plants (Site A and Site B) located ∼18 km apart in the state of São Paulo, Brazil. The two plants are owned by different companies and use different sets of starter strains in their process. We employed a combination of whole-population metagenome sequencing and clonal whole-genome sequencing to observe the temporal dynamics of genetic diversity in each site-year. See Supp. Table 1–3 for a complete list of collected samples and isolates.

Yeast strains differ in their suitability for industrial-scale fermentation. Traditionally, the readily available baker’s yeast was used to kickstart the fermentation season, but due to its susceptibility to invasion by foreign *S. cerevisiae* lineages, production has largely shifted towards specialized starter strains. A major strain selection program conducted between 1993 and 2005 solidified the potential for these invading strains themselves to serve as a source of new industrially relevant variants^11^. Strains isolated from this program, namely PE-2, CAT-1, SA-1, BG-1, VR-1, and their derivatives, as well as JP-1 (isolated from a similar effort^12^) are the basis for the bulk of today’s ethanol production and have successfully helped maintain the overall high yield of the industry. Still, invasion by foreign strains remains common, as fermentation conditions across the ∼400 bioethanol plants operating around the country span a range of industrial practices, environmental conditions, sugarcane varieties, and other factors, in addition to the yet-little-explored possibility of evolutionary change over the course of a fermentation season.

To identify and track these yeast population dynamics in industry, chromosomal karyotyping became popular in the 1990s and is still commonly used for process monitoring^11–13^. More recently, PCR-based methods have helped in decreasing the cost of strain surveillance^14–17^. However, these methods cannot readily differentiate closely related strains, which may differ by few mutations anywhere along the whole genome. Moreover, these methods estimate lineage frequencies based on fraction of picked isolates from agar plate streaks, which leaves room for biased assessments of strain dominance if strains differ in culturability.

Whole-genome metagenomic shotgun sequencing is a potential culture-independent alternative method for strain differentiation^18^. Temporal metagenomic datasets have been used to assess microbial community dynamics with subspecies resolution, largely in the context of human gut microbiomes^19–28^. However, inference of the underlying strain movements from metagenomic frequency trajectories remains challenging and methods are mostly limited to low-diversity and prokaryotic populations. Non-haploidy complicates this inference even further, as the diploid or polyploid genotype of individual variants (which itself may vary among individuals in a population) must also be accounted for.

Here, we present a novel framework for inferring the population dynamics of highly diverse, non-haploid, asexual microbial populations from a combination of clonal sequences and temporal metagenomic data. We employ this method to investigate the dynamics of yeast genetic diversity across two fermentation seasons, in two independently run bioethanol plants in Brazil. More specifically, we ask whether starter strains tend to persist and dominate through an entire production season, and if not, what strains they are replaced with. We also investigate the differences between seasons and production facilities, the origin of invading strains, and the effects they have on the process. Our focus here is on the yeast dynamics, but our sequencing data also contains information on other microbial species, which remains to be analyzed in future work.

## RESULTS

### Sampling and sequencing strategy

We collected whole-population microbiological samples from two independent industrial units, which we refer to as *Site A* and *Site B*, through two fermentation seasons, *2018* and *2019* (Fig. 1). Sampling started on the first day of the fermentation season for Site A 2018, and ∼14 days into the season for the other site-years (see sampling dates in Supp. Table 1). The two sites are owned by different companies and are located 18 km apart in the region of Piracicaba, São Paulo, Brazil. Site A used a mix of four strains to start both the 2018 and 2019 fermentation periods—namely strains PE-2, SA-1, FT-858, and IRA-D. While the first three are common commercially available industrial strains, IRA-D is an in-house strain isolated from Site A in a previous fermentation season. In contrast, Site B informed us that they have used PE-2 as their sole starter strain in both fermentation seasons, although we would later find evidence suggestive of a second starter strain being used, possibly unknowingly, in 2019 (see results below).

Samples were taken directly from fermentation or holding tanks and were composed of a mix of fermentation liquid and cells. Glycerol was immediately added for cryopreservation. For metagenome sequencing, we simply pelleted cells, extracted their DNA and performed sequencing library preparation using a tagmentation-based approach for short-read whole-genome sequencing, for a total of ∼15 timepoints from each site-year. From each of these sequenced timepoints, we also streaked the original sample on rich medium agar plates, picked up to three colonies from each, and used the same tagmentation-based approach for clonal whole genome sequencing (see Supp. Table 2 and 3 for isolate information). We did the same with samples of the four starter strains (see Methods). All reads from metagenome and clonal isolates were then aligned to the reference genome of *S. cerevisiae* S288c and used to call SNPs (single-nucleotide polymorphisms) for further analyses. Our final dataset is thus composed of *alternate allele counts* and *depth of sequencing* (hereon referred to as simply count and depth) at each called variant site both for individual clonal isolates and whole population timepoints. In both cases, *allele frequency* will refer to the quotient count/depth. See Methods for details.

### High genetic diversity among industrial isolates

We began by investigating genetic diversity in the studied populations. Using our variant calling pipelines (see Methods), we find a total of 145,066 SNPs among all 134 fermentation and 11 starter strain isolates. 14,200 (9.8%) of these mutations are singletons, while 15,749 (10.5%) are seen in all sequenced clones (see Ext. Data Fig. 1 for the full distribution). We also find a similar number of SNPs (150,265) in the whole-population metagenome data across all four site-years, with an overlap of 126,845 between the clonal and the metagenomic datasets. This suggests that the clonal genotyping data covers a substantial fraction of the genetic diversity of these populations, especially given that the metagenomic data (i) samples from the whole population, and (ii) represents a sequencing effort of 6154x over all timepoints, which is larger than that of clonal genotyping (4,341x over all isolates). The 168,486 SNPs uncovered in the whole dataset are widely distributed along the genome, hitting 6,370 out of all 6,579 genes in the annotated S288c genome. 129,697 of these SNPs have been previously observed in the 1011 yeast genomes project^29^, which itself uncovered 1,544,489 SNPs.

*S. cerevisiae* may exist at different ploidies, and so we examined allele frequencies in the clonal isolate data to infer isolate ploidy (see Methods for details). We found that 64 of our isolates are triploid, while the remaining 70 are diploid (Fig. 2A). All isolates of starter strains FT-858 and IRA-D are triploid, while those of PE-2 and SA-1 are diploid (as described in ref.^11,30,31^). An examination of allele frequencies and sequencing depth along the genome revealed that a small number of isolates carry structural variations, such as gain or loss of whole chromosomes or sections of chromosomes (Supp. Fig. 3). Given the small number of affected isolates, and in each case a minor fraction of the genome being affected, we keep these isolates in all further analyses.

**Figure 2.**
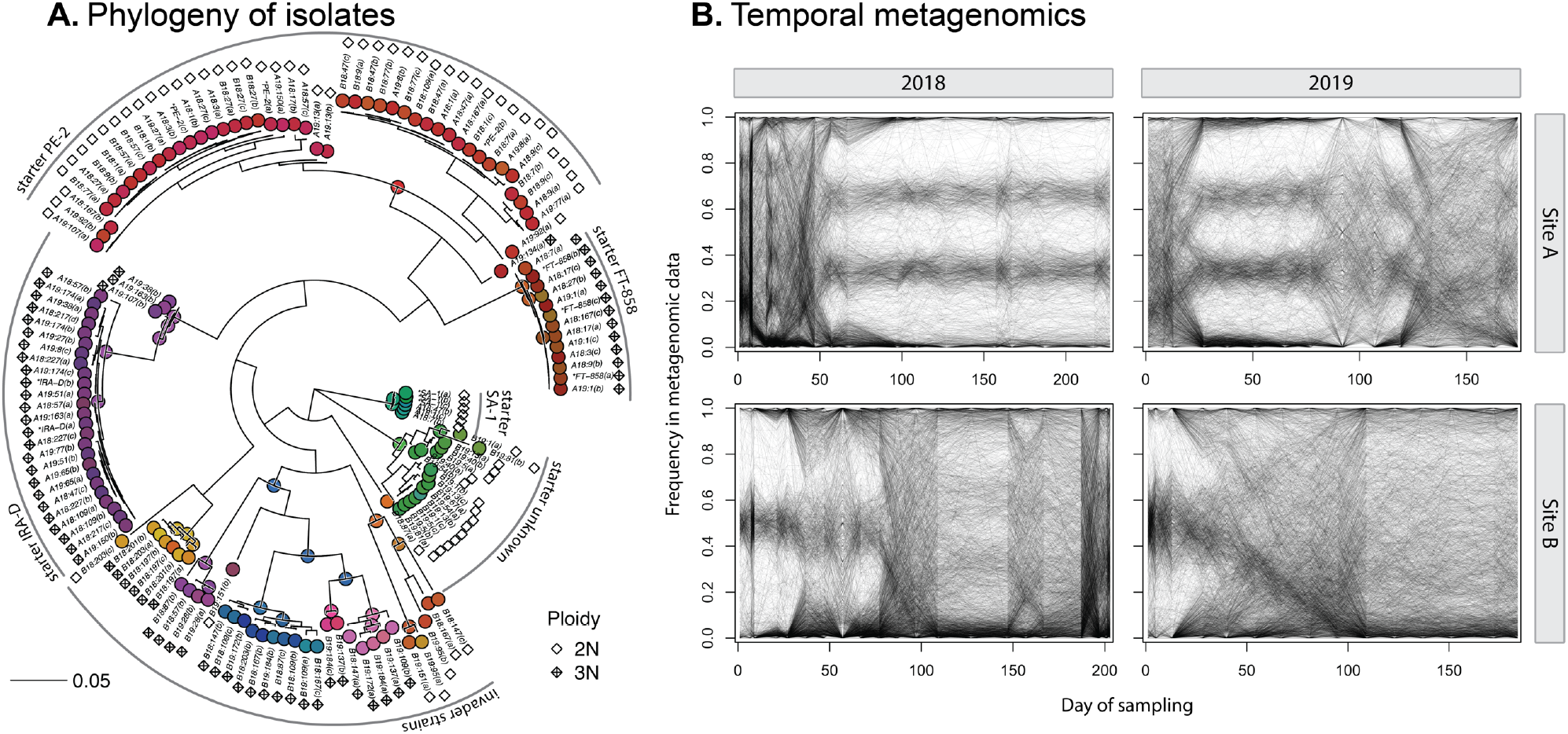
Yeast populations in bioethanol fermentors are genetically diverse and dynamic. **(A)** Phylogenetic tree of isolated clonal strains from all site-years, as well as known starter strains used. Most isolates are closely related the known starter strains, but several are not. The tree was inferred with a maximum likelihood model using the data of 27,229 SNPs. Ploidy of each isolate, assessed as described in the Methods, is indicated by diamonds. Nodes and tips are colored as in Figs. 4 and 5. The tree is rooted in the same place as the independently inferred tree in Fig. 6. Isolates are grouped as in Figs. 4–6. Isolates are named as *<site><year>:<timepoint>(<letter identifier>)*, while starter strain isolates are marked with an asterisk. The associated Newick tree can be found in Supp. Data 1. The allele frequency data used for ploidy assessment can be visualized in Supp. Fig. 3. Selected examples of a diploid and triploid strain can be seen in Ext. Data Fig. 2. **(B)** Frequency of alternate allele (in relation to the reference genome of strain s288c) through time for an arbitrary subset of 2000 mutations (out of ∼100k) per site-year. Overall, mutation trajectories indicate alternation between periods of stasis, when one major strain dominates, and periods of transition, when many mutations change in frequency in a correlated way indicative of strain dynamics. Noise in mutation trajectories comes from random sampling (approximately binomial), as well as sequencing and mapping errors, which is not homogeneous across mutations.

We then used the called SNP data to infer a maximum-likelihood phylogenetic tree between all sequenced isolates (Fig 2A). As expected, we find that several of the isolated clones are closely related to the starter strains used to initiate the industrial process. We note that PE-2 isolates form two major clades, which are both represented in starter and fermentation isolates from both sites and years. We also find several other groups of closely related isolates, mostly triploid, that diverge from the starter strains by thousands of SNPs. These groups are all composed of isolates from Site B, whereas all Site A isolates fall close to the known starter strains.

### Lineage inference

We turned to the whole-population metagenomic data to investigate the yeast population dynamics through the fermentation season (Fig. 2B). We are interested in understanding how starter strains change in frequency through the fermentation, as well as identifying events of selection of novel mutations or invasion by foreign strains. Examining the raw metagenomic allele frequencies through time, we observe periods when large cohorts of mutations move together, indicative of competition between divergent strains, as well as periods of stability when allele frequencies remain mostly constant. Correlation between allele frequency trajectories is indicative of co-segregation and has been used as the signal for inference of population dynamics in previous studies^21,25^.

However, this type of inference is complicated by several factors. First, our populations are highly genetically diverse and mutations are shared between different strains in complex patterns. These patterns are presumably created by earlier, potentially sexual population dynamics that led to the creation of these strains in the unknown other environments in which they evolved. This means that individual metagenomic mutation trajectories can depend on the frequency changes of potentially multiple different strains that carry that mutation. This is complicated by the fact that these different strains may carry a given mutation at different genotypes (i.e. as homozygous or heterozygous diploids, or in one to three copies in triploids). Finally, it is not immediately clear how to polarize mutations for lineage frequency inference (i.e. which one should be considered the references versus alternative allele), which leads to an overall pattern of mirrored mutation trajectories in the raw metagenomic data (Fig. 2B).

Here, we developed and employed a novel framework for jointly inferring the frequencies of nested asexual lineages of descent through time from whole-population metagenomic data (Fig 3; see Methods and Supplementary Information for details). This approach takes advantage of our clonal sequencing data to phase an informative subset of all mutations into cohorts that segregate together in the population, completely ignoring the metagenomic data for this purpose. While we are limited to the genetic diversity that is sampled by picked isolates, by following this approach we overcome the challenges described above, as well as have higher power to identify small lineages, whose metagenomic trajectories may be indistinguishable from sequencing noise in correlation-based grouping methods^21,25^. In doing so, our pipeline automates an approach similar to that of Zhao and colleagues^27^, while handling high genetic diversity and ploidy variation in the population.

**Figure 3.**
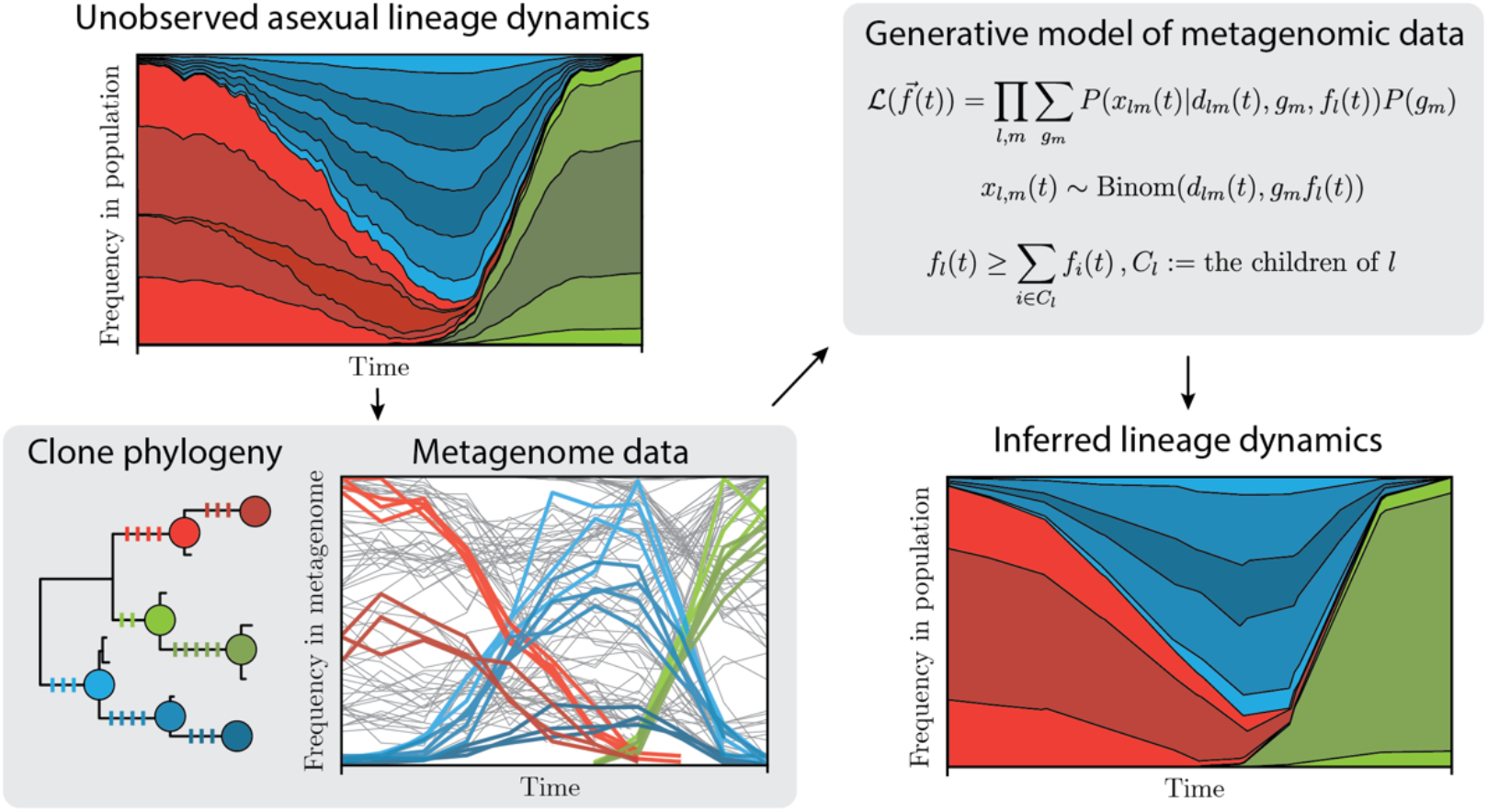
Schematics of lineage inference procedure. We use temporal metagenomics and clonal isolate whole-genome sequencing to infer the unobserved frequencies of asexual lineages in the original population over the course of a fermentation season. (Upper left) Starter, invading, and newly mutated lineages change in frequency through time due to selective and random factors. (Lower left) A phylogeny of clonal isolates is used to select the sets of clade-defining variants (colored bars on tree branches) that we will later search in the metagenomic data and use for lineage inference. (Upper right) At each timepoint t, we jointly infer the frequencies 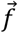 of all asexual lineages by optimizing a likelihood model of 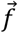 given the metagenomic allele counts *x*_*lm*_ of variant *m*, which is a clade-defining variant for lineage *l*, the read depth *d*_*lm*_, and the variant’s genotype *g*_*m*_ (which takes values 0, 0.5 or 1 for diploid, and 0, 1/3, 2/3 or 1 for triploid lineages). The frequencies of all lineages are jointly inferred and constrained such that the summed frequencies of sister lineages do not exceed that of the respective parent lineage. (Lower right) Undersampling of genetic diversity by isolates will cause whole lineages to be left out, but that should not bias the frequency estimation of included lineages.

Among the four site-years, we infer the frequencies of a total of 197 lineages, spanning a wide range of lineage sizes, with a median maximum lineage frequency of 6.7% (see Ext. Data Fig. 3 for the full distribution). The inferred results pass basic soundness checks: the timepoints at which different isolates were picked largely correspond to times when their associated inferred lineage frequencies are high, and lineage frequency trajectories are smooth, even though timepoints are inferred independently from each other.

### Stable dynamics dominated by in-house strain in Site A

In Site A, we only observe lineages closely related to the known starter strains (Fig. 4). In particular, we find that IRA-D, a triploid strain, dominates the process in both years. Curiously, IRA-D is an in-house strain which was found to invade the process in a previous fermentation season, and since then it has been included in the starter strain mix. While these observations suggest that IRA-D is the best adapted to these fermentation conditions among all four starter strains, we observe that it does not completely displace PE-2 in 2019, which continues at a low frequency in the process even in later timepoints. Coexistence for such a long timescale is suggestive of some ecological process, such as niche partitioning, or negative frequency dependence. However, it is unclear why the same dynamics are not seen in 2018, when PE-2 seems to be completely outcompeted. Either the population itself is genetically different between the years (although isolates from both seasons are closely related) or differences in agricultural and industrial practices, or weather patterns, may have affected fermentation conditions.

**Figure 4.**
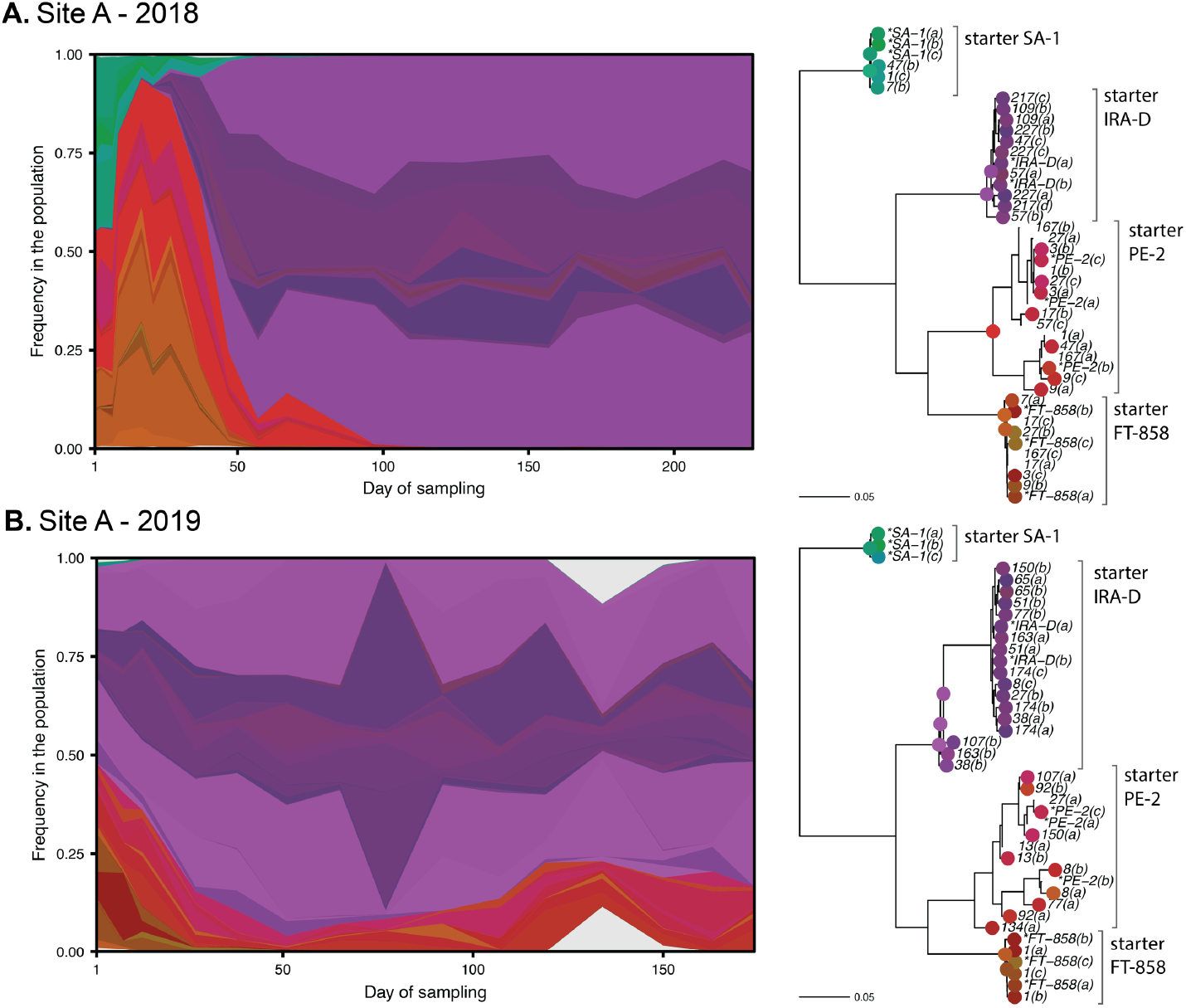
In Site A the in-house starter strain IRA-D consistently dominates over other starter strains. On the left, inferred strain dynamics in Site A over the two fermentation seasons. White space corresponds to non-inferred genetic diversity in the population. On the right, subtrees of the tree in Fig. 2A including only the isolates from each respective site-year. Circles on nodes and tips indicate inferred lineages and their respective colors.

### Foreign lineages systematically invade Site B

In Site B, we observe a very different picture, where several large lineages are distantly related to the starter strain PE-2 (Fig. 5). While PE-2 dominates at the start of 2018, it is a minor fraction at the start of 2019, when the process is instead dominated by a different lineage (labeled “starter unknown” in Fig. 2A and 5), suggesting a different starter strain mix for that year.

**Figure 5.**
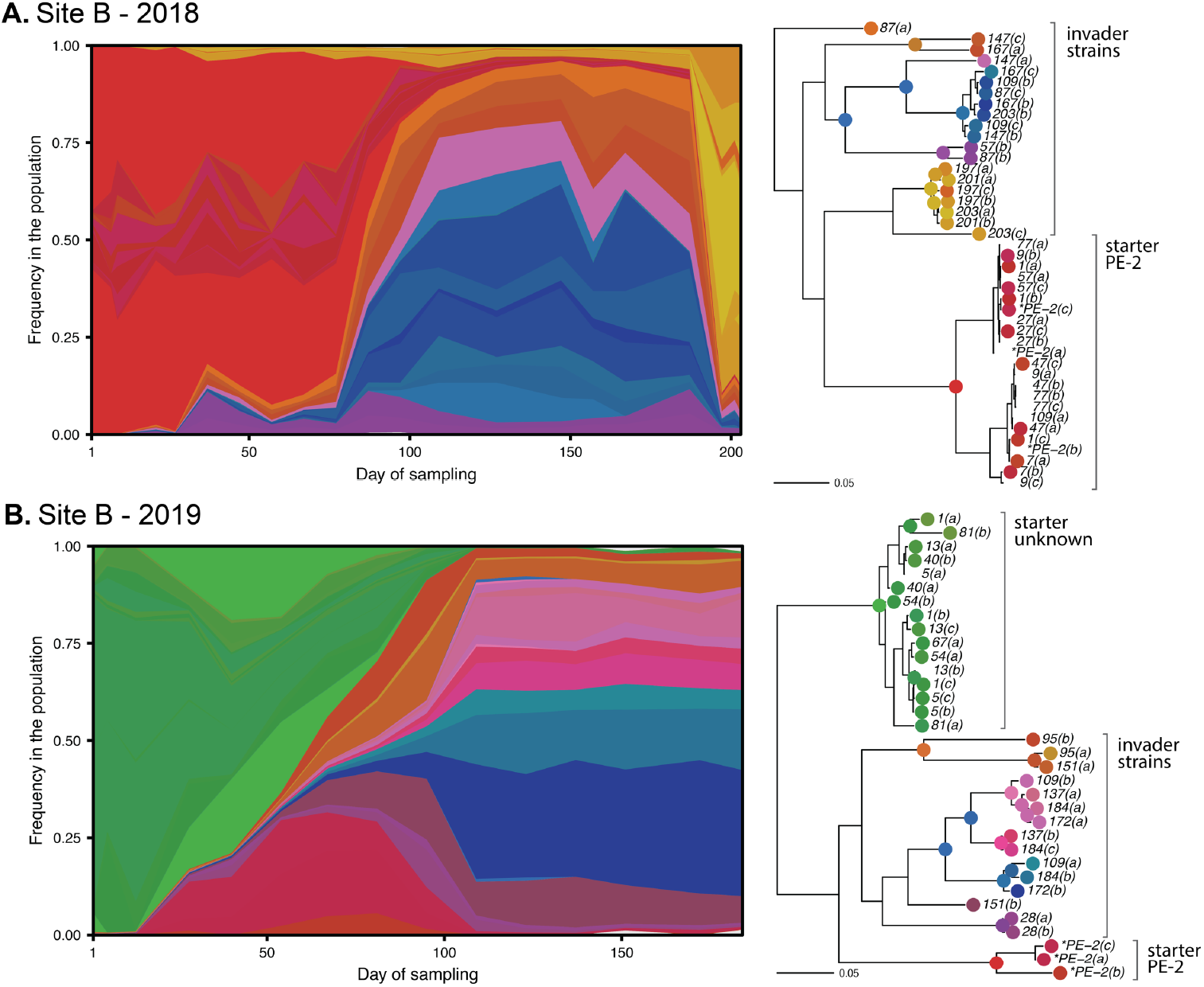
In Site B, a group of diverse invading strains systematically takes over the process. Despite the genetic diversity among invader strains, they seem to coexist, except for the second substitution event in 2018, which involves a different set of invading strains. In the 2019 fermentation season the process starts with a large amount of an unexpected unknown strain. See Fig. 4 for a description of the diagrams.

In both years, the population gets substituted by a cohort of much fitter strains halfway into the season (labeled invader strains in Fig. 2A and 5). Most of these strains are triploid, except for a small group present in both years (Fig. 2A and 5). While their genetic distance to other starter strains and minute presence in early timepoints suggest that they invade the fermentation process, we cannot rule out that they were already present in the starter inoculum or have their origin in the industrial equipment itself, where they might find a reservoir from one production season to the next. The fact that closely related isolates are seen in both 2018 and 2019 is indicative of some systematic source of contamination. Surprisingly, despite the large degree of genetic diversity and the ploidy variation within this cohort, these different invading strains stably coexist in the timescale of the fermentation season. Here again, an ecological explanation is suggested.

Finally, we observe a second substitution event in the final timepoints of Site B’s 2018 season. The inference suggests that this set of strains were already present since early in the season, remaining at low frequency until they suddenly displace all other strains. This event does not seem to be driven by selection for a novel mutation, since the expanding lineage retains significant diversity within itself, and instead may be caused by a sudden change in fermentation conditions.

### Origin of invading yeast strains

We further investigate the origin of Site B’s invader strains. While we cannot assess industrial procedures directly, we can examine the phylogenetic relationship of these strains to other known isolates. For that purpose, the 1011 Yeast Genomes Project (YGP) represents the largest and broadest whole-genome sampling of *S. cerevisiae* genetic diversity^29^. Most importantly, it includes 37 isolates related to the Brazilian bioethanol industry. Here, we compare all our picked isolates to the YGP collection by inferring a combined phylogeny of both studies (Fig 6; see Methods for details). The inferred unrooted tree largely replicates the structure of previous inferred trees of broad yeast diversity^29,32–34^.

**Figure 6.**
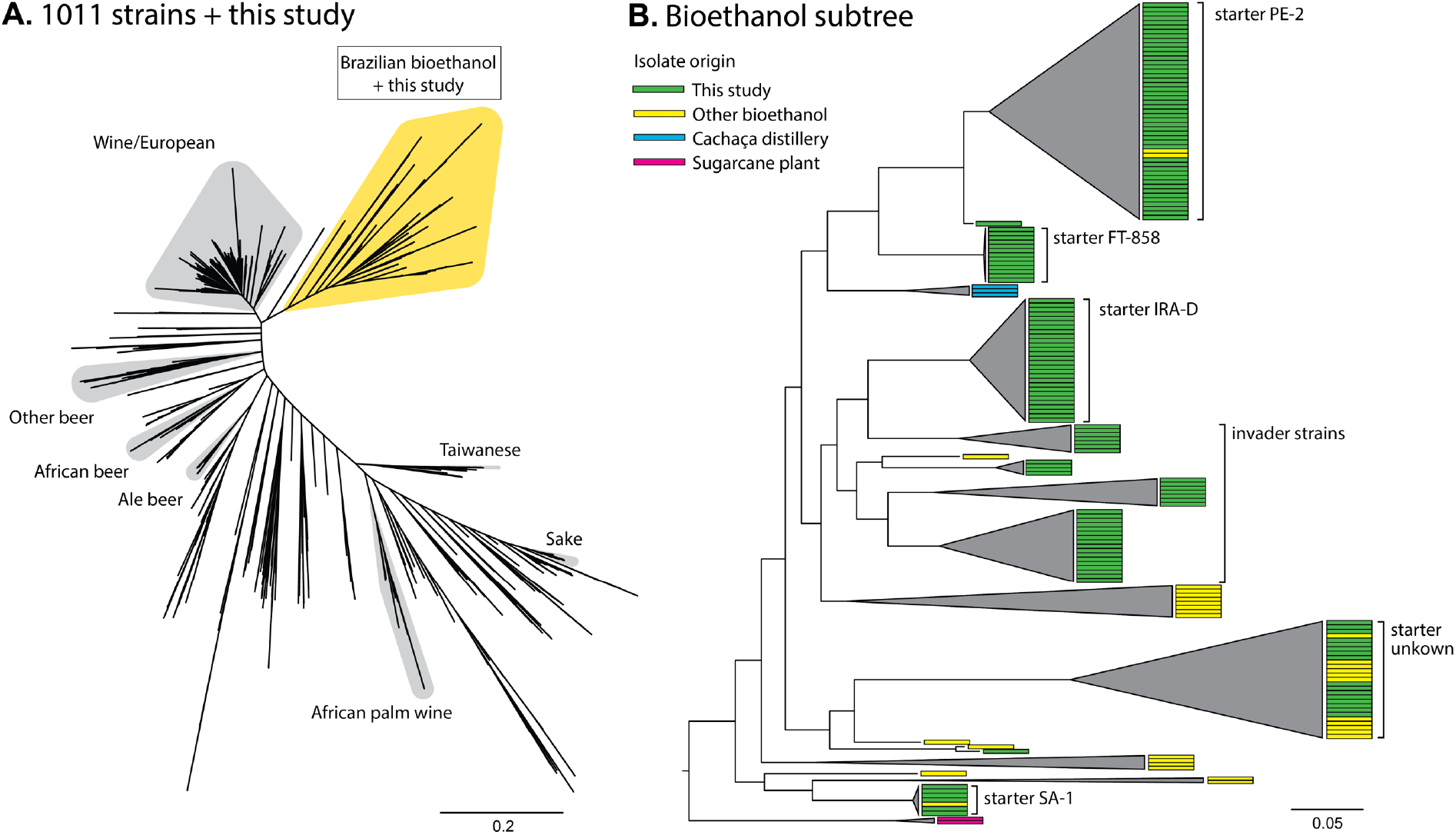
Starter and invader isolates all cluster together within a larger group of Brazilian Bioethanol strains. **(A)** A SNP-based maximum likelihood phylogeny combining isolates from the current study and from the 1011 Yeast Genomes Project^29^. Other groups of domesticated strains are highlighted for reference. This tree was inferred based on 42,012 SNPs. **(B)** Subtree of bioethanol-related isolates. Isolates from the current study are closely associated with isolates from the bioethanol industry and cachaça distilleries (a sugarcane-based spirit). Individual isolate origins are indicated with colored rectangles. Branches are collapsed to aid visualization. A full phylogeny can be seen in Ext. Data Fig. 4, and its associated Newick tree can be found in Supp. Data 2.

First, we find that all Brazilian bioethanol isolates from both studies form a monophyletic group and are closely related to a large group of European wine strains, in agreement with previous studies^29,34^ (Fig. 6A). As shown in Fig. 6B, we note that among the 37 isolates classified in the Brazilian bioethanol group in the 1011 YGP, 3 were isolated from cachaça distilleries (a traditional sugarcane-based spirit), while 2 were from the sugarcane plant or from sugarcane juice (although further detail is missing), while the remainder were isolated from different bioethanol plants. Among these isolates from the bioethanol industry, several are closely related to PE-2, SA-1, and most notably, to the “unknown starter” strain in Site B’s 2019 season. Finally, Site B’s “invader strains” do not seem to be represented in the 1011 YGP, but their close association with other bioethanol isolates points to an industrial origin (e.g. shared equipment, supplies, or sugarcane), as opposed to invasion by wild strains brought to the industrial environment by vectors such as insects or birds from foreign niches.

### Stability of macroscopic fermentation parameters despite strain dynamics

Yeast strains vary in their suitability for the industrial process due to, among other factors, their ability to produce and withstand high ethanol concentrations, their propensity to generate foam or cell aggregates in large industrial settings, or their tendency to be outcompeted by poorer performing strains^11^ (in terms of the final ethanol yield on sugars). Thus, invasion by unknown strains may harm the fermentation process and the profitability of the industry, due to decreased ethanol production and/or to higher costs involved with the use of chemicals, such as sulfuric acid, antimicrobials, antifoaming agents and dispersants. In the case of Site B’s 2018 and 2019 seasons, we have not found a connection between general industrial metrics and inferred events of population substitution (Ext. Data Fig. 5). Nonetheless, it may still be possible that this stability was accomplished by the employment of commonly used but costly corrective measures, such as those outlined above.

## DISCUSSION

In this study, we described the population dynamics of the yeast used for bioethanol production via fermentation in sugarcane-based biorefineries through the course of two fermentation seasons (2018 and 2019) in two independently run industrial plants. The method we developed for this purpose allowed for an unprecedented description of how the starter strains used in the process change in frequency through time and how the fermentation environment may be invaded by foreign strains. We observe that these large populations (estimated to be ∼10^17^ individuals) harbor a vast amount of genetic diversity, recovering ∼8% of alleles previously found in a *S. cerevisiae*-wide survey^29^, plus novel ones. This diversity is not only observed in invading strains, but also within the starter strains themselves, whose same subtypes are sampled across years and sites (most notably the two major groups within PE-2; Fig. 2A). This may be due to how propagation companies, which sell large initial inocula to bioethanol producers, keep and propagate their own stocks: companies may not start from single colonies every year, and new mutations may accumulate during propagation. Similar observations of strain genotypic (and phenotypic) heterogeneity have also been made in the baking, wine and beer industries^35^.

Such large populations must harbor many novel mutations. At an approximate rate of 5 × 10^− 10^ mutations/bp/ generation^36^, and at least 66 generations during one fermentation season, a total of 8 × 10^16^ or more mutations should occur in a diploid population of this size. In fact, at this rate, any given SNP in the yeast genome should independently occur ∼3 × 10^7^times per generation. We cannot know how many of these mutations would be adaptive in the industrial environment, but decades of microbial experimental evolution, including in yeast populations, show that adaptation in large asexual populations is not mutation-limited^37–42^. Yet, we do not find clear signs of selection for novel mutations in our results, which would be observed as either an inferred lineage that increases in frequency much faster than its closely related counterparts, or inferred lineages being deflected by some unobserved rising lineage. A likely explanation is that the timescale of a fermentation season (in number of generations) is too short for selected lineages, carrying novel adaptive mutations of a typical fitness effect, to increase in frequency enough to be sampled by our sparse isolate picking strategy. All in all, what this suggests is that as long as starter inocula are not produced from the previous year’s final population, or that the equipment itself is not contaminated with large amounts of previous populations, evolution on a single-strain background is likely not a consequential factor in the timescale of a fermentation season due strictly to the large population sizes and dynamics of selection.

Ecological dynamics may explain the observed long periods of coexistence between distantly related lineages in both sites, such as in PE-2’s permanence in Site A 2019, or the stable relative frequencies of invader strains in Site B 2019. While it is possible that these observations simply reflect small differences in fitness in the fermentation environment, the large phylogenetic distance between strains argues against this hypothesis. Large genetic differences may lead to diversity in resource usage (niche partitioning), and/or in how strains benefit or not from each other’s presence (frequency dependence). Such ecological dynamics are by no means rare in microbiological communities in the wild^43,44^, and have been unintentionally evolved in laboratory *E. coli* and *S. cerevisiae* populations^39,45^. Strain interactions could open up avenues for designed strain mixes that take advantage of synergistic interactions in terms of fermentation output and management. We also should not discount the potential bacterial contribution to these dynamics, as bacteria have been shown to interact both positively and negatively with yeast during fermentation^7,10^. The analyses carried out for the current study do not include bacterial data, but such microbial consortia compose an interesting avenue for future work.

The fact that results have varied more between industrial plants than between years suggests that systematic differences in industrial practices and/or starter strain mix largely explain differences in population dynamics. Additionally, observed fluctuations in strain frequencies through time (e.g. the strain responsible for the second substitution event in Site B 2018) indicate that fluctuations in fermentation conditions may make certain strains more or less fit to the industrial environment. This is not unexpected, as (i) fermentors are only partially protected from external temperature fluctuations, (ii) incoming sugarcane varieties change through the year and result in different must compositions, (iii) the ratio of sugarcane juice and molasses in the must is adjusted daily depending on current sugar and ethanol prices, (iv) clean-in-place (CIP) practices are carried out on a regular or as-needed basis, and (v) recycling practice may be adjusted depending on levels of bacterial contamination, among other factors. Further collaborations with companies, including access to a detailed record of industrial practices and strain-tracking as done in this study, may shed further light into the causes behind fermentation fluctuations. These records should especially contain information on the usage of chemicals (e.g. sulfuric acid, antimicrobials, antifoaming agent and dispersant, among others), which remediate fermentation output, but add to production cost and greenhouse gas emissions.

Our observation that the in-house strain IRA-D dominates the process throughout the two observed seasons in site A underscores the potential of *in loco* isolation of industrial strains. Invading strains have been documented to cause harm, but they also served as the source for most if not all of the currently used strains in the industry^11,46,47^. Previous studies had shown that these known bioethanol strains are phylogenetically related and harbor genomic signals of domestication, some which are shared with wine strains and others that are specific to bioethanol strains^34^. These strains also cluster very far apart known natural *S. cerevisiae* isolates from other Brazilian biomes, further suggesting a non-natural origin^48,49^. Our results show that currently invading strains in Site B are closely related to these known domesticated bioethanol strains. On top of that, we note that the dominant strains across all sites and years are largely triploid, suggesting a systematic advantage of higher ploidy in this industrial environment (Ext. Data Fig. 6). Taken all together, we hypothesize that the same patterns hold in most strain invasion events in bioethanol plants that follow a process similar to Site A and B (Fig. 1A). The observed large genetic diversity among invading strains should be further explored as a potential resource for future strain isolation. Strain monitoring as carried out in the current study is thus not only a sanity check, but also a productive assistive strategy for the selection of novel and locally adapted industrial strains. For this purpose, industrial plants should have protocols in place for the isolation of invading strains, record-keeping of associated fermentation metrics, and subsequent testing in blocked off portions of the industrial pipeline and scaled-down systems that mimic the industrial process^50^.

Our study used metagenomics and a newly developed framework to extract individual lineages to illuminate the yeast population dynamics in industrial sugarcane-based bioethanol production, with the goal of finding routes towards more consistent fermentation performance. The resolution obtained with this approach surpasses by far previously described and utilized methods, such as chromosomal karyotyping and PCR-based methods. We observed that over two sampled production periods in two independent industrial units, the yeast population dynamics varied more dramatically between units than between years. In one site we observed dominance and persistence of an in-house strain in both years, whereas in the other site, foreign strains invaded the process and displaced the starter strain used to initiate the production period. The several individual clones sequenced, including invading strains, are phylogenetically grouped with other known bioethanol strains, producing strong evidence that the invading strains originate from the sugarcane environment itself, and not from natural niches. The data presented, as well as the statistical framework developed, represent useful material for future investigations on sugarcane biorefineries (as well as other microbial communities of mixed ploidy). This, in turn, might lead us to a deeper understanding of the yeast and other microbial ecology in this peculiar environment, opening the way for process improvements, decreased consumption of costly chemicals, and increased ethanol yields. A potential new paradigm of industrial practice includes the design of synergistic yeast strain mixes, and the inoculation of beneficial (or probiotic) bacteria in the process.

## METHODS

### Sample collection

Whole-population microbiological samples were collected from two bioethanol plants, here named Site A and Site B, through the 2018 and 2019 sugarcane-crushing seasons, which ran from April/May through November/December. See Supp. Table 1 for sampling dates and estimated correspondence with days in season. The two sampling sites are owned by different companies and are located 18 km apart (on a straight line) in the region of Piracicaba, SP, Brazil. Samples (∼10 ml) were collected daily (2018) or weekly (2019), after fermentation was completed, directly from fermentors or holding tanks, into pre-sterilized 15 ml tubes containing 3 ml glycerol. After mixing by vortexing, samples were stored at –20°C for a period of between one and three months before being transferred to a –80°C ultrafreezer. Finally, samples were shipped from Brazil to the US in dry ice, where they were stored at –80°C. Starter strains PE-2, FT-858 and SA-1 were shipped as active dry yeast (ADY), whereas strain IRA-D was shipped as colonies on agar slants, without dry ice. The collection and shipping of samples has been registered at the Sistema Nacional de Gestão do Patrimônio Genético e do Conhecimento Tradicional Associado (SisGen, Brazilian federal government) under numbers R40E57A, RB42674, R193AED and RAD5521 (for the shippings), and AF14971 (for the sampling). A full list of samples with associated collection dates can be found in Supp. Table 1.

### DNA extraction and sequencing

We selected 15 to 20 samples from each site-year for whole-genome metagenomic and clonal sequencing. For metagenomic sequencing, samples were completely thawed and vortexed, after which 1 ml was aliquoted and centrifuged to remove the supernatant. Whole DNA extraction was carried out using an in-house protocol^40^. Sequencing library preparation was done using the transposase-based protocol^51^.

For clonal isolate sequencing, the same 15 to 20 thawed and homogenized samples were used for plating onto Yeast Extract-Peptone-Dextrose(YPD)-agar (Supp. Table 2). Plates were incubated at 30°C for 24 - 48 h. From each plate, 2 or 3 CFUs were picked and grown in 5 ml liquid YPD overnight at 30°C, after which DNA extraction and library preparation proceeded as for metagenomic sequencing. Starter strains were inoculated in liquid YPD, left to grow overnight at 30°C, plated and prepared in the same manner (Supp. Table 3).

Sequencing was carried out in two Illumina NextSeq and one Illumina Miseq runs, following a 300 bp paired-end workflow. Mean coverage after mapping to the reference strain S288c genome and haplotype inference (see *Bioinformatics* section) was 87x for metagenomic samples and 26x for clonal isolates. FASTQ files with all sequencing reads produced for this study were deposited in the NCBI SRA database (see Data and Code Availability).

### Variant calling bioinformatic pipeline

We called variant sites (SNPs only) in relation to the *S. cerevisiae* S288c reference genome (yeastgenome.org, release R64) in all our metagenomic and clonal isolate data. The full pipelines with specific tools and settings used can be found in the GitHub repository (see Data and Code Availability). In summary, all sequencing reads were first trimmed of sequencing adapters using NGmerge^52^, and then aligned to the reference genome using BWA^53^. Variant calling was done with the haplotype inference tools in the Broad Institute’s GATK^54^. In essence, these tools assemble local haplotypes from aligned reads, calculate the posterior probability of each read coming from each of the assembled haplotypes, and finally infer variant sites jointly across a group of samples for added power to call true low-frequency variants: intuitively, an observed variant is less likely to be a sequencing error if it is observed in more than one sample. Given different probabilistic prior models of allele frequency for clonal and non-clonal data, variant calling of isolate clonal data is done with HaplotypeCaller jointly across all isolates, while that of the metagenomic data is done using Mutect2 jointly across all timepoints within each site-year, in line with GATK guidelines^54^. Alternate and reference allele counts (AD field in the VCFs) outputted by the variant calling tools are estimates based on inferred haplotype membership of aligned reads (instead of being simple observations from aligned reads). These are the numbers that we use for all later analyses. For convenience, when referring to a variant site, we will often refer to alternate allele counts as simply *counts*, and the sum of alternate and reference allele counts as simply *depth*. In all further sections, *allele frequency* at a variant site is defined as the number that ranges from 0 to 1 given by counts divided by depth. For the sake of simplifying, we exclude from analyses the small number of variant sites for which we observe more than one alternate allele.

### Isolate ploidy

Isolate ploidy was assessed based on visual examination of the distribution of allele frequencies in clonal isolate data over the whole genome (upper right corner of each panel in Supp. Fig. 3): diploid strains have a multimodal distribution peaked at values 0, 0.5 and 1, while triploid strains, at 0, 1/3, 2/3, and 1. Example allele frequency distributions from a diploid and a triploid strain are shown in Ext. Data Fig. 2.

### Phylogenetic analyses

We infer two phylogenetic trees in this study, both using whole-genome SNP data. *Tree 1* was run with the SNPhylo pipeline^55^ using default parameters. The tree is inferred based on a total of 27,229 SNPs across all clonal isolates from all site-years, including isolates from the four starter strains (Fig. 2A; Newick format tree in Supp. Data 1). *Tree 2* includes the same clonal isolates, plus all isolates from the 1011 Yeast Genomes Project^29^ (Fig. 6; Ext. Data Fig. 4; Newick format tree in Supp. Data 2). For this tree, SNPs were first filtered and aligned using SNPhylo with a missing rate of 0.001, and a maximum likelihood tree was constructed from 42,012 SNP markers using RAxML^56^ with 1000 bootstrap replicates, employing the general time reversible nucleotide substitution model with the GAMMA model of rate heterogeneity. For the purposes of downstream analyses and presentation, Tree 1 was rerooted in a node analogue to that from which the Bioethanol subtree of Tree 2 branches from the remainder of the tree.

### Inference of population dynamics

We assume the reproduction during fermentation is exclusively asexual. Therefore, the population is composed of some large but discrete number of clonal strains of asexually dividing individuals which may have three origins: (1) preexisting diversity in starting inoculum; (2) invading strains during the course of the fermentation season; (3) new strains founded by mutational events during fermentation.

Clonal strains share phylogenetic history, and therefore alleles. Assuming no recombination, and no mutation reversal, we assume that these lineages organize themselves into a hierarchical tree-like structure which defines clades, herein referred to as *lineages*, each with a particular set of synapomorphic alleles: i.e. alleles that are shared by all clonal strains within that lineage, but no strain outside of it. In effect, the inference pipeline should be able to handle some amount of departure from these assumptions due to past history of recombination, mutation reversals, and noise, but we expect this pattern to compose the bulk of the observed data.

Our goal was to use the metagenomic data to infer the frequencies through time of as many lineages as possible in order to characterize the population dynamics over the course of the fermentation season in each site-year. Our inference consists of (i) identifying lineages and their synapomorphic alleles based on a maximum-likelihood phylogeny inferred from our sequenced clones; and (ii) looking for each lineage’s set of synapomorphic alleles among the metagenomic sequencing data to infer lineage frequencies using a maximum-likelihood framework. The rationale for this approach is that the metagenomic data samples genetic diversity among chromosomes in the population in an unbiased way, while the clonal genome sequencing informs us of how to group alleles that segregate together in the same lineages. We do not assume any particular dynamical model of evolution, and instead infer lineage frequencies at each timepoint independently. A crucial feature of this inference is that genetic diversity that is not sampled among sequenced clones does not bias the frequency estimates of other lineages.

A detailed description of the inference pipeline is described in the Supplementary Information. The code developed for this inference is available in the GitHub repository (see Data and Code Availability).

## DATA AND CODE AVAILABILITY

Raw sequencing reads for clonal and metagenomic samples have been deposited in the NCBI BioProject database under accession number PRJNA865262. Code for the variant calling pipeline, lineage inference, and figure generation, as well as parsed called variant data for clonal and metagenomic samples can be found in the GitHub repository (https://github.com/arturrc/bioethanol_inference).

## AUTHOR CONTRIBUTIONS

M.M.D. and A.K.G. designed the project; A.K.G. sequenced samples; A.R.-C and I.H. developed inference methods; A.R.-C., I.H., and A.K.G. analyzed the data. A.R.-C., I.H., M.M.D., and A.K.G. wrote the paper.

## ACKNOWLEDGMENTS

We thank the Bauer Core facility at Harvard for assistance with sequencing. M.M.D. acknowledges support from grant PHY-1914916 from the NSF. A.K.G. acknowledges support from the Harvard Lemann Brazil Research Fund, and from grant 2018/04962-5 from FAPESP. The computations in this paper were run on the FASRC Cannon cluster supported by the FAS Division of Science Research Computing Group at Harvard University.

## EXTENDED DATA FIGURES

**Extended Data Figure 1.**
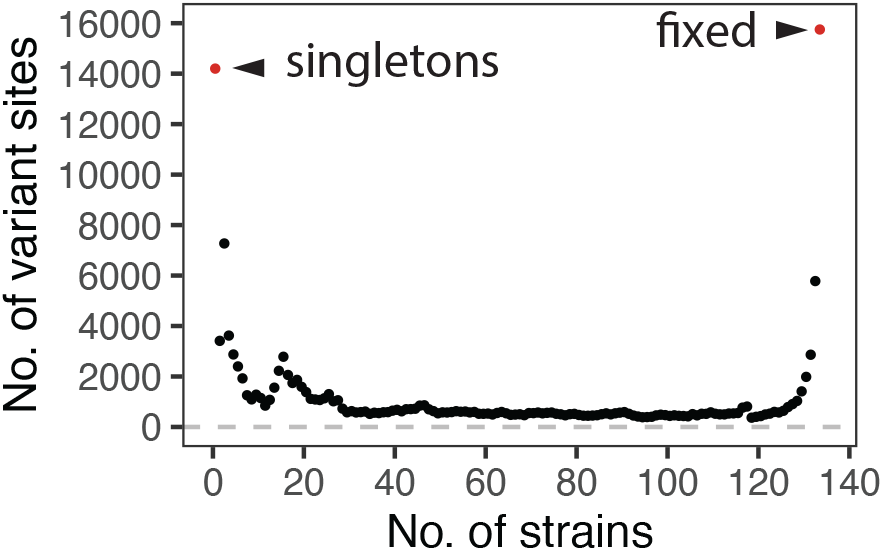
Histogram of number of isolates observed to carry a given alternate allele in the clonal sequencing data. Starter strains were excluded.

**Extended Data Figure 2.**
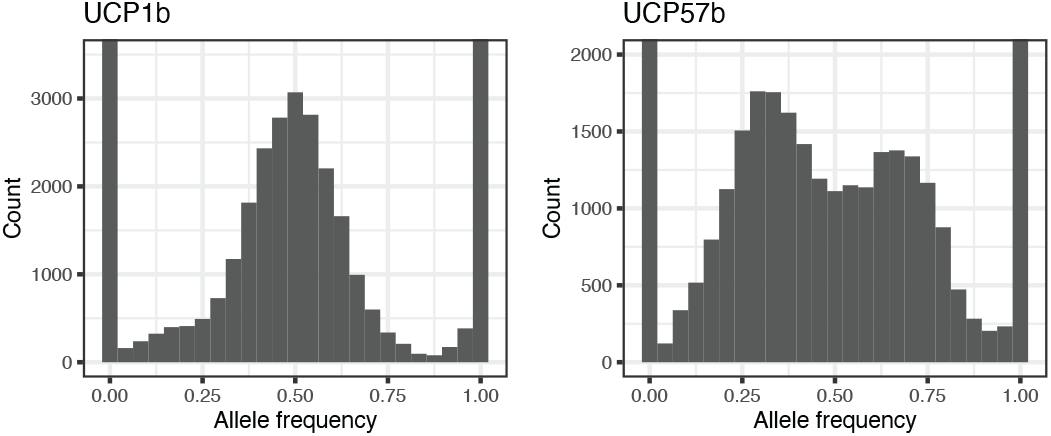
Representative examples of diploid and triploid whole-genome allele frequency distribution in the clonal sequencing data. The y-axes are cropped for better visualization.

**Extended Data Figure 3.**
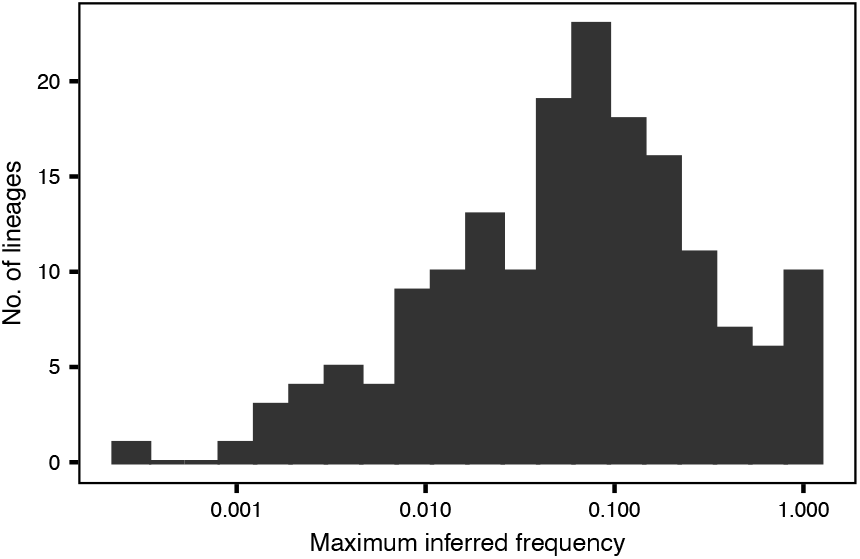
Distribution of maximum inferred frequency (over all timepoints) for all 197 inferred lineages across all site-years.

**Extended Data Figure 4.**
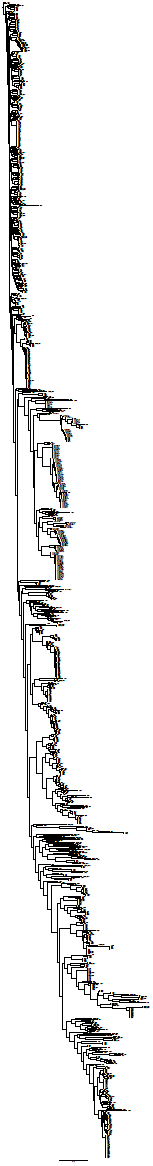
Midrooted labeled version of the tree in Fig. 6A. Clones from this study are labeled as in Supp. Table 2 and 3. Clones from the 1011 YGP are labeled as in Supp. Table 1 of ref.^29^.

**Extended Data Figure 5.**
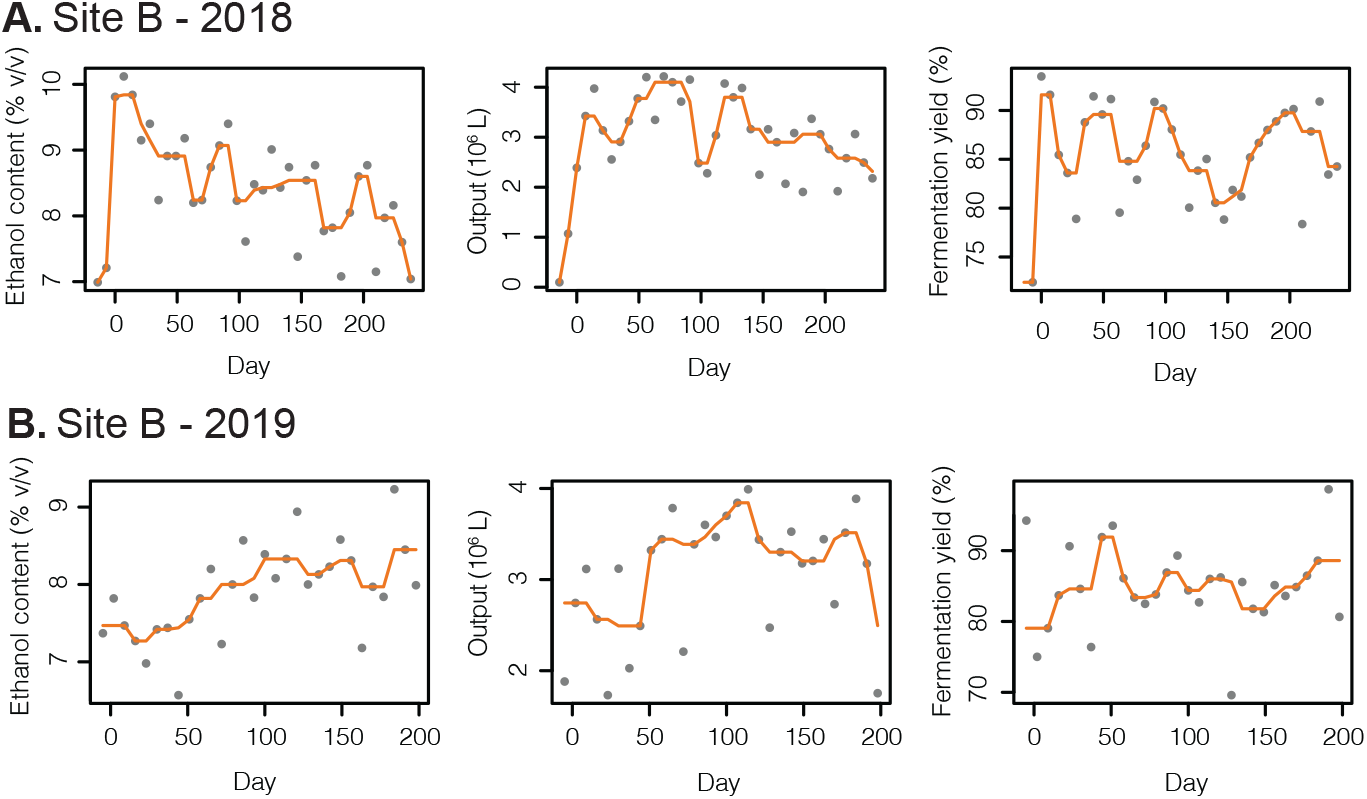
Fermentation metrics in Site B show no clear relationship with invasion by foreign strains. We show weekly data over the 2018 and 2019 fermentation seasons for (left) ethanol content of fermented wine, (middle) total bioethanol output, and (right) fermentation yield, as a measure of amount of ethanol produced out of a theoretical maximum. A running average is shown as an aid (orange line). The raw data can be found in Supp. Table 4.

**Extended Data Figure 6.**
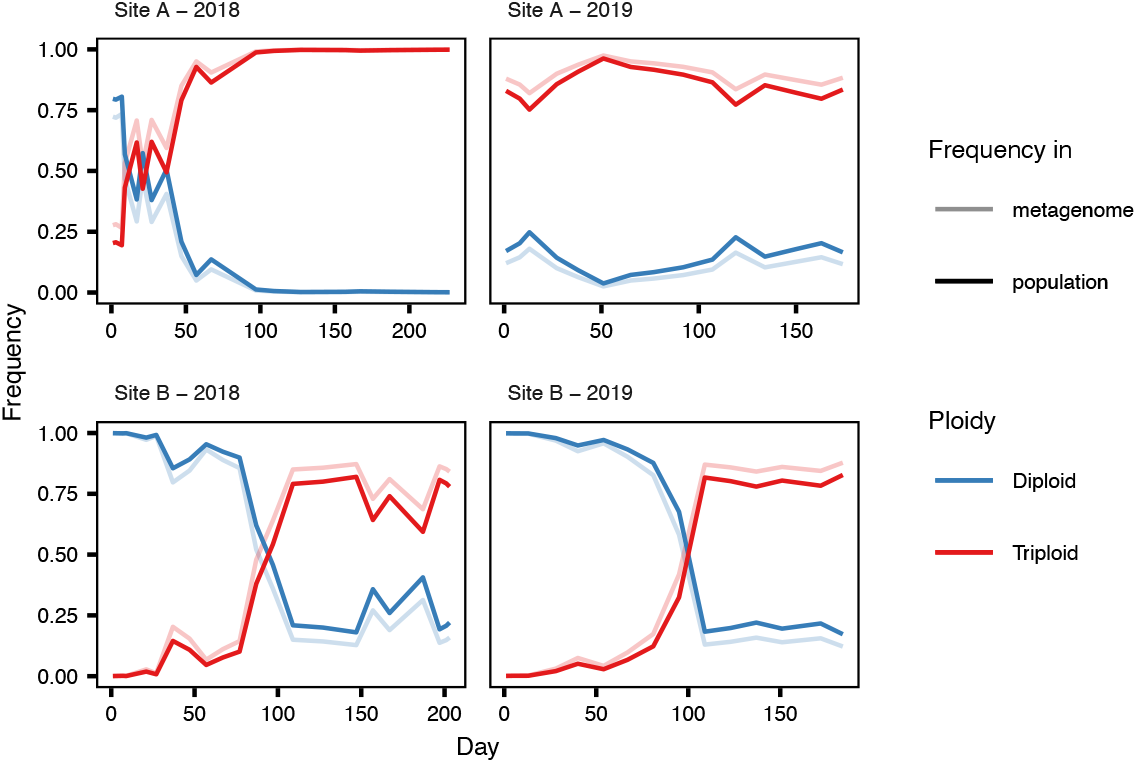
Inferred fraction of diploid and triploid strains along time based on inferred lineages’ frequencies and ploidies. Estimated frequencies in both the metagenome (*i*.*e*. fraction of genetic material of the population that can be assigned to diploid or triploid individuals) and in the population (fraction of individuals) are shown. See Section “Calculation of lineage frequency in the population” of the Supp. Information for details.

## SUPPLEMENTARY MATERIAL

***Supplementary Information***

Details on lineage inference pipeline. Supplementary Figs. 1 and 2.

***Supplementary Figure 3***

Allele frequency and coverage along the genome of each sequenced isolate. Each file corresponds to a sequenced clone and contains four panels. (Top) Allele frequency (alternate allele counts/depth) along the genome, and histogram of allele frequency. (Bottom) Coverage along the genome, and histogram of coverage. Histograms are cropped for visualization. Red bars represent boundaries between each of the 16 chromosomes in the reference strain s288c.

***Supplementary Tables 1–4***

List of collected samples, and sequenced isolates. Site B’s weekly fermentation metrics along 2018 and 2019 production seasons.

***Supplementary Data 1***

Newick format tree of inferred maximum likelihood phylogeny of all sequenced isolates. See Methods for details.

***Supplementary Data 2***

Newick format tree of inferred maximum likelihood phylogeny of all 1011 YPG and this study’s sequenced isolates. See Methods for details.

