## Supplementary Information for "Yeast population dynamics in Brazilian bioethanol production"

### INFERENCE OF POPULATION DYNAMICS

As described in the Methods, we assume that the population at each site-year is composed of a large but finite number of clonal strains which are related by some phylogenetic history in a tree-like manner. Clades in this tree represent lineages of descent from a common ancestor and is what we will be referring to as *lineages* throughout the text.

Our goal here is to (i) use the whole-genome clonal isolate data to identify as many as possible sets of lineage-defining synapomorphic alleles, and (ii) use the metagenomic frequencies of these synapomorphic alleles to infer their respective lineage frequencies in the population through the course of a fermentation season. By doing this, we ignore correlation between mutations in the metagenomic data as signal of coinheritance, something that has been previously done in literature [refs]. The advantage of following this route is higher power to identify low-frequency lineages, whose mutations' metagenomic trajectories would be too overpowered by noise to ever have a significant correlation signal (although our ability to identify these low-frequency lineages is still ultimately limited by the clonal isolate sampling).

In spirit, we follow a strategy similar to that of [Tami's paper], with the important difference that our populations are highly diverse and non-haploid. The consequence is that a large number of mutations will be unsuitable for inference, either because they are not monophyletically shared in the inferred phylogeny, or because their genotype (i.e. number of allele copies within an isolate's genome) varies among isolates that carry it, thus complicating the mathematical relationship between lineage frequency in the population and allele frequency in the metagenomic data.

#### Lineage assignment

We will define lineages as monophyletic clades in the phylogenetic tree inferred for all isolates from our experiment, which is in principle an unrooted tree (Fig. 2A). Since we observe (in a second inferred tree of all our isolates and those from the 1011 genomes project; see Methods for details) that all Brazilian bioethanol isolates cluster together, and within that cluster the SA-1 isolates are the most basal among our isolates (Fig. 6B), we root that first tree of isolates in the analogue node (as shown in Fig. 2A). From this rerooted tree, we define all lineages, (i) which include the very base of the tree with all isolates in the experiment, (ii) all internal nodes and their respective descendant isolates, and (iii) each tip with its associated isolate. Note that since this tree is inferred from isolates, it is most likely undersampling the genetic diversity of the population. Some lineages, especially the smaller ones, will most likely be missed (as illustrated in Fig. 3).

#### Finding lineage-specific alleles

For each one of the lineages defined above, we first would like to find a set of alleles that are unique to it. We cannot assess all individuals in the original population, and so instead we use the observed alternate allele counts and depth of coverage at variant sites in the clonal isolate data as a proxy. Therefore, for each lineage, we first flag all variant sites for which either (i) counts in all lineage members are larger than zero, while counts in non-lineage members are zero, or (ii) counts in all lineage members are less than the depth, while counts in all non-lineage members equal the depth. The second case covers variant sites for which the reference allele is the derived (synapomorphic) one in the phylogeny. For these mutations, in all analyses described below, *counts* will refer to the count of reference allele (instead of alternate allele).

If the lineage under consideration has a single isolate, then all flagged mutations are kept. Otherwise, we must select only those mutations for which we believe all isolates in the lineage to have the same genotype. For a diploid strain, the genotype of the mutation $m$ in isolate $i$ takes values $g_{mi} \in \{0, {}^{1}/_{2}, 1\}$, while for a triploid strain, $g_{mi} \in \{0, {}^{1}/_{3}, {}^{2}/_{3}, 1\}$. For this reason, we exclude from further analyses any lineages composed of a mix of diploid and triploid isolates. For each of the mutations flagged for a lineage we apply a statistical test of genotype heterogeneity, explained in more detail in the section below, where the null hypothesis is that all isolates in the lineage carry that mutation at the same genotype. We then use a procedure similar to Benjamini-Hochberg to select mutations for which we do not reject the null at a False Omission Rate of 0.05 (defined as false negatives/[false negatives + true negatives]).

We apply some filters before arriving at a final list of lineages and mutations for later frequency inference. First, we only keep those mutations that we also observe in the metagenomic dataset. Second, we limit the total number of mutations in a lineage to 500 to keep later steps computationally tractable. When this limit is imposed, mutations are chosen arbitrarily. Third, we filter mutations based on their observed depths in the metagenomic dataset, as they suggest underlying read mapping issues: we remove any mutations that have median depth in the metagenomic data lower than 10, or that has any metagenomic timepoint with depth equal to 0. Finally, we exclude any lineages for which we have selected 3 or less mutations, as we have observed that to result in noisy frequency inference.

#### Genotype heterogeneity test

As described in the section above, we would like to test whether a mutation is carried at the same genotype across all isolates from a lineage. For that we do a chi-squared test of goodness of fit to the model that all isolates have the same genotype.

Let $a_{mi}$ and $b_{mi}$ be the counts and depths of mutation $m$ in isolate $i$. We first would like to define a generative model for the data so that we can compute the likelihood $P(a_{mi}|b_{mi}g_{mi})$. We choose a simple approach that assumes that $a_{mi}$ is largely binomially distributed, except for a small probability of random errors, which can shift the count $a_{mi}$ upwards or downwards. These errors may come from any of the preceding steps in data generation and analysis (e.g. sequencing and mapping errors), and they need to be accounted for the correct genotyping of homozygous sites that show a small (erroneous) count towards the opposite allele. We assume that the observed count $a_{mi}$ is the result of a mixture of two populations of reads observed at site $i$: *true* and *error* reads. The $b_{mi}^{T}$ true reads contribute with an alternate allele count $a_{mi}^{T} \sim \mathrm{Binom}\left(b_{mi}^{T}, g_{mi}\right)$, while the $b_{mi}^{E}$ error reads contribute with an alternate allele count $a_{mi}^{E} \sim \mathrm{Binom}\left(b_{mi}^{E}, 0.5\right)$. We further assume that error reads are independent of each other and occur with equal probability $p_{error}$, such that $b_{mi}^{E} \sim \mathrm{Binom}(b_{mi}, p_{error})$. Since $b_{mi}^{E}$ and $a_{mi}^{E}$ are unobserved quantities, we marginalize over their possible values, and thus

$$P(a_{mi}|b_{mi}g_{mi}) = \sum_{b_{mi}^{E}=0}^{b_{mi}} \sum_{a_{mi}^{E}=0}^{\min(b_{mi}^{E}, a_{mi})} P\left(a_{mi}^{T} = a_{mi} - a_{mi}^{E} \middle| b_{mi}^{T} = b_{mi} - b_{mi}^{E}, g_{mi}\right) P\left(a_{mi}^{E} \middle| b_{mi}^{E}\right) P\left(b_{mi}^{E} \middle| b_{mi}\right),$$

where each probability above is calculated based on the probability mass function of the binomial distribution. Finally, we assume $p_{error} = 0.01$, which accomplishes our goal of a less stringent genotyping criterion at homozygous sites (Supp. Fig. 1).

If the null hypothesis that all isolates have the same genotype is true, then all inference could be done on the summed counts and depths $a_m = \Sigma_i a_{mi}$ and $b_m = \Sigma_i b_{mi}$, in which case the most likely genotype $\hat{g}_m$ for that mutation is

$$\hat{g}_m = \max_{g_m}[P(a_m|b_m, g_m)],$$

where $P(a_m|b_m, g_m)$ is calculated as described above.

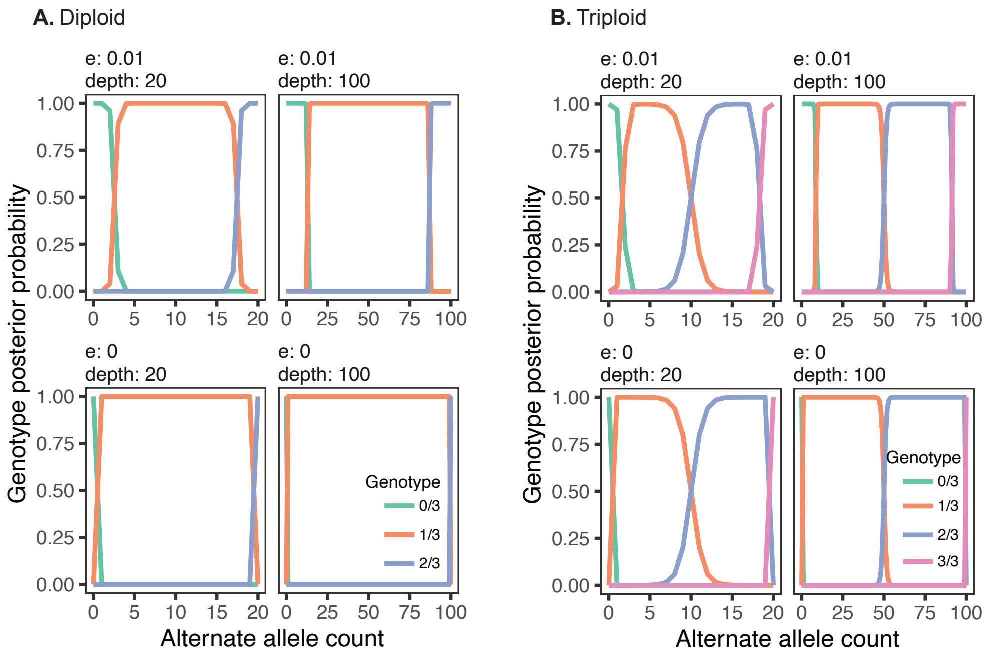

**Supplementary Figure 1. Probability of isolate data given genotype allowing for sequencing error.** We show the computed probability of observing an alternate allele count value based on a given the depth of coverage at that site, the probability of count errors $p_{error}$ (*e* in the figure), and the isolate ploidy.

We calculate the expected counts if the null is true as $\hat{a}_{mi} = \hat{g}_m b_{mi}$, with which we compute the test statistic

$$x^2 = \sum_i \frac{(a_{mi} - \hat{a}_{mi})^2}{\hat{a}_{mi}}.$$

If $\hat{a}_{mi} > 5$ for all $i$, we compute an exact $p$-value taking $x^2 \sim \chi^2_{\text{df=\#of isolates}-1}$ under the null assumption. Otherwise, we calculate an empirical $p$-value from 1,000 permutations of alternate and reference allele observations keeping the isolate depths constant.

#### Genotype posterior probability

In the later lineage frequency inference step, we would like to marginalize the likelihood of a mutation's metagenomic counts and depths by its genotype $g_m$, which effectively serves to downweight mutations for which we have less certainty about their genotype. For that we use an Expectation-Maximization procedure. We compute the posterior probability of the genotype $g_m$ given the summed isolate clonal counts and depths $a_m$ and $b_m$ (see section above) as

$$P(g_m | a_m, b_m) = \frac{P(a_m | g_m, b_m) P(g_m)}{\sum_{g_m^*} P(a_m | g_m^*, b_m) P(g_m^*)},\ a_m \sim \text{Binom}(b_m, g_m).$$

At first, we assume a uniform prior for $P(g_m)$, but having calculated the posteriors, we can update the priors as

$$P(g_m) = \sum_{m^*} P(g_{m^*} = g_m | a_{m^*}, b_{m^*}),$$

where $m^*$ iterates over all mutations selected for a given lineage. We iterate over the two equations above until values converge enough, using a stop criterion on the change per iteration of the total likelihood of the data.

#### Joint inference of lineage frequencies in the metagenome

At this point, we have a list of lineages and their associated synapomorphic mutations. Note that, by definition, there is no overlap between the mutations used to identify any two lineages. We would like to use the metagenomic data for these mutations to infer the frequencies of the lineages during the fermentation season. For now, we will infer the frequency $f_l(t)$ of *chromosomes* of lineage $l$ among all chromosomes in the population. This differs from the frequency $f_l^*(t)$ of *individuals* of lineage $l$ among all individuals in the population because our populations are composed of a mix of diploid and triploid strains. We calculate this latter quantity in the section below.

We will do this inference independently for each timepoint, to avoid having to assume any particular model about how these lineages change in frequency through time. At each timepoint, we infer frequencies for all lineages jointly. If we allowed frequencies to vary freely, this would be equivalent to inferring each lineage's frequency independently. However, our lineages are hierarchically organized according to the inferred phylogenetic tree used to define them (as shown in Fig. 2A): we will use the term parent, child, and sibling lineages to point to the relationship between lineages in this hierarchy. In the most basal part of the tree, we will have one or more lineages that have no parent. Therefore, the frequencies $\vec{f}(t)$ of all lineages at a timepoint $t$ are constrained by the set of inequalities

$\sum_{l \in B} f_l(t) \leq 1$, for the set of sibling basal lineages $B$, and

$\sum_{l \in C_p} f_l(t) \leq f_p(t)$, for the set $C_p$ of children of a given lineage $p$.

We assume that the error in metagenomic counts for different mutations are independent from each other, which is an assumption that only breaks in the case of mutations that are close enough in the genome that they may be covered by a same sequencing read. We therefore calculate the likelihood of a given model of lineage frequencies given the data as (suppressing $t$ for convenience)

$$\mathcal{L}(\vec{f}|\text{data}) = \prod_l \prod_m \sum_{g_m} P(x_m|d_m, g_m, f_l) P(g_m|a_m, b_m),$$

where $x_m$ and $d_m$ are the counts and depths of mutation $m$ in the metagenomic data, and we assume $x_m \sim \text{Binom}(d_m, g_m f_l)$.

We maximize the likelihood model above using a gradient descent method with a log-barrier that bounds solutions to the inequalities above as implemented in the function `constrOptim` in base R [ref]. To make this inference computationally tractable we do not infer the frequencies of all lineages at once, and instead follow an iterative procedure where at each step we infer the frequencies of a parent and all its children jointly starting from the most basal lineages:

(1) jointly fit frequencies of basal lineages $l \in B$, keeping $\sum_{l \in B} f_l(t) \leq 1$;

(2) randomly sort basal lineages; following this order jointly fit the frequency of basal lineage $p$ and children lineages $C_p$, with inequalities

$$f_p \leq 1 - \sum_{p^* \in B | p^* \neq p} f_{p^*}, \text{ and}$$

$$\sum_{l \in C_p} f_l(t) \leq f_p(t);$$

(3) keep this new frequency $f_p$;

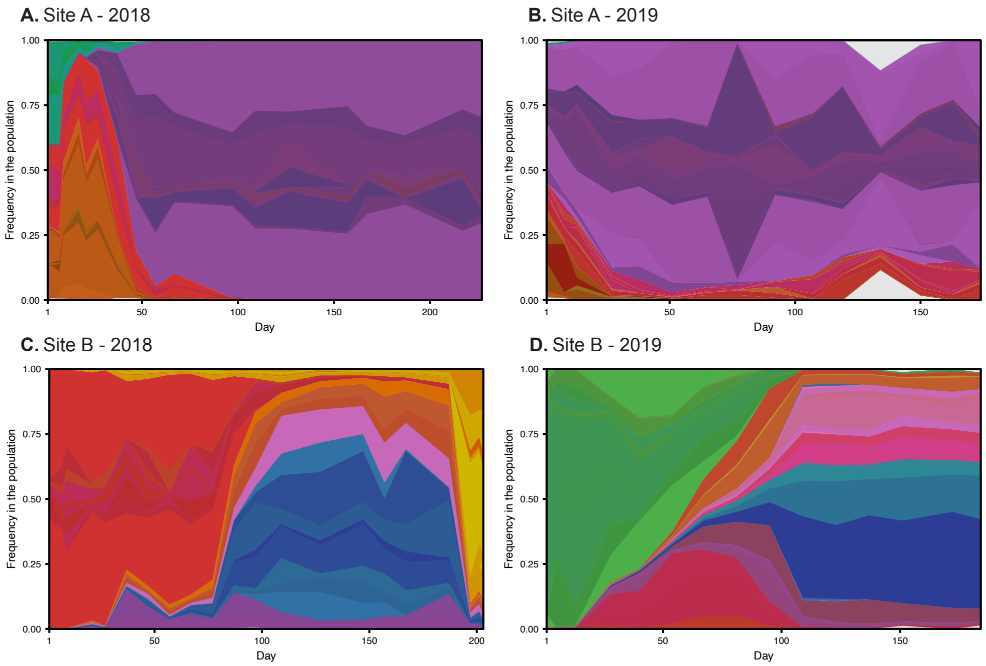

**Supplementary Figure 2. Inferred frequency of lineages in the metagenome.** These frequencies are inferred with the procedure described in the section above and are later used to compute the frequencies of lineages in the population, as shown in Figs. 4 and 5. Lineages are color-labeled as in Fig. 4 and 5.

(4) for each fit grandparent lineage $g$, randomly sort its (also already fit) children $C_g$; following this order, fit jointly the frequencies of lineage $p \in C_g$ and its respective children $l \in C_p$, with inequalities

$$f_p \le f_g - \sum_{p^* \in C_g | p^* \neq p} f_{p^*}, \text{ and}$$

$$\sum_{l \in C_p} f_l(t) \le f_p(t);$$

(5) keep this new frequency $f_p$;

(6) repeat steps (4) and (5) until there are no more lineages to be fit.

We show inferred $\vec{f}(t)$ in Supp. Fig. 2.

### Calculation of lineage frequency in the population

Having inferred the frequencies $\vec{f}(t)$ of all lineages in the metagenome, we proceed to calculating frequencies $\vec{f}^*(t)$ of all lineages in the population. These two quantities are related as (suppressing $t$ for convenience)

$$f_l = \frac{p_l}{\bar{p}} f_l^*$$

where $p_l \in \{2,3\}$ is the ploidy of lineage $l$, and $\bar{p}$ is the mean ploidy in the population. Notice that if the whole population is composed of individuals of the same ploidy, then $f_l = f_l^*$.

We cannot directly assess the ploidy of all individuals in the original population, so instead we use inferred $\vec{f}(t)$ and respective lineage ploidies to estimate the mean ploidy in the population, but with two caveats. First, our isolate sampling may have missed ploidy heterogeneity within lineages. Second, our inference is not bound to infer frequencies that sum to 1 in the population, and thus may leave some portion of the population uninferred and of unknown ploidy. This is not a significant fraction in our study (see Supp. Fig. 2), but it may be in other systems. We therefore make two assumptions: that (*i*) we are not missing ploidy heterogeneity in the inferred portion of the population, and that (*ii*) any non-inferred portion of the population has the same mean ploidy as the inferred portion.

Let $F_2(t)$ and $F_3(t)$ be the total frequency of diploid and triploid strains in the metagenome as computed from inferred $\vec{f}(t)$. The frequencies $F_2^*(t)$ and $F_3^*(t)$ of diploid and triploid strains in the population are, thus, given by (suppressing $t$ for convenience)

$$F_p^* = \frac{\frac{F_p}{p}}{\frac{F_2}{2} + \frac{F_3}{3}},$$

from which we compute the mean ploidy in the population as

$$\bar{p} = 2F_2^* + 3F_3^*.$$

We show computed $F_p(t)$ and $F_p^*(t)$ in Ext. Data Fig. 7, and inferred $\vec{f}^*(t)$ in Figures 4 and 5 of the main text. Effectively, they only slightly deviate from inferred $\vec{f}(t)$ (Supp. Fig. 2).
