## Supplementary figures and images for "Yeast population dynamics in Brazilian bioethanol production"

### A18-9-a.png

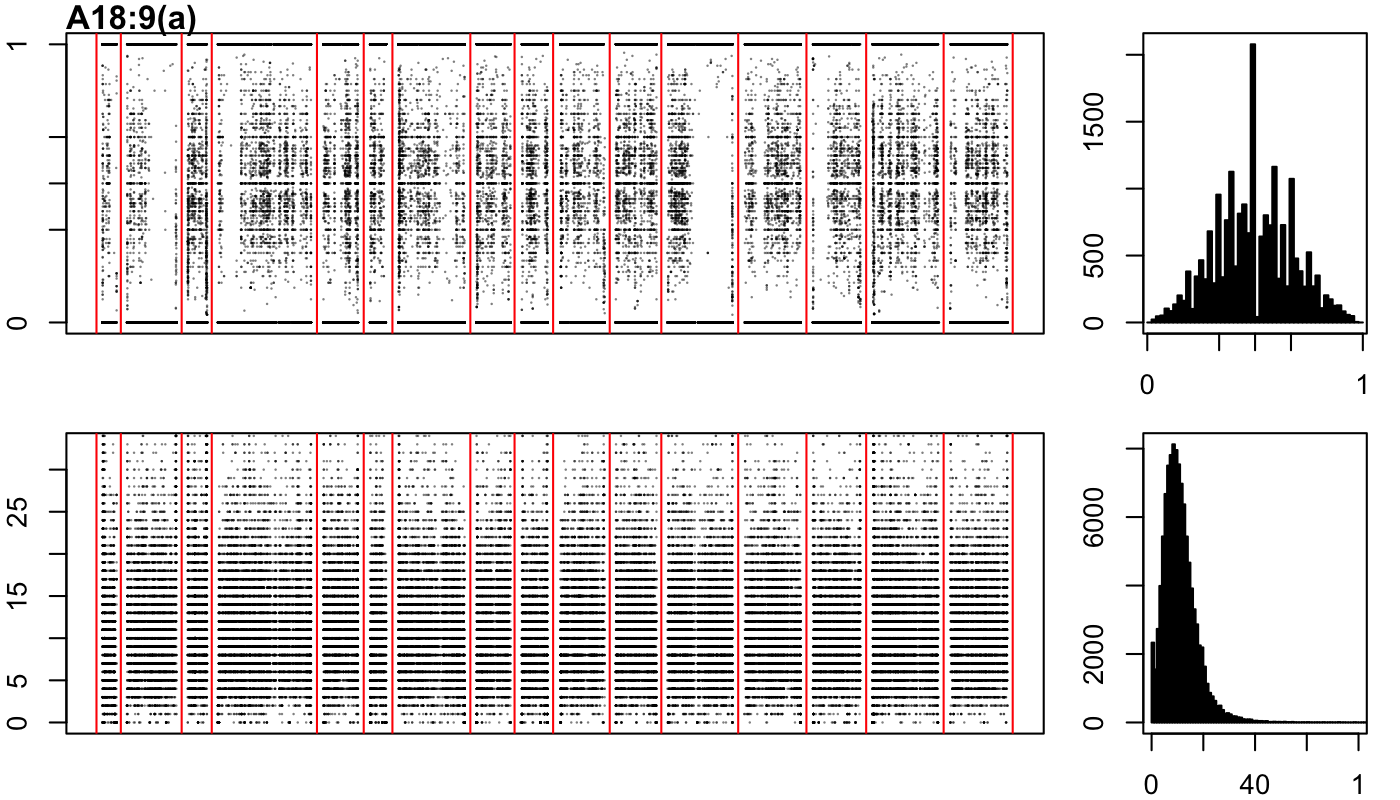

### A18-9-b.png

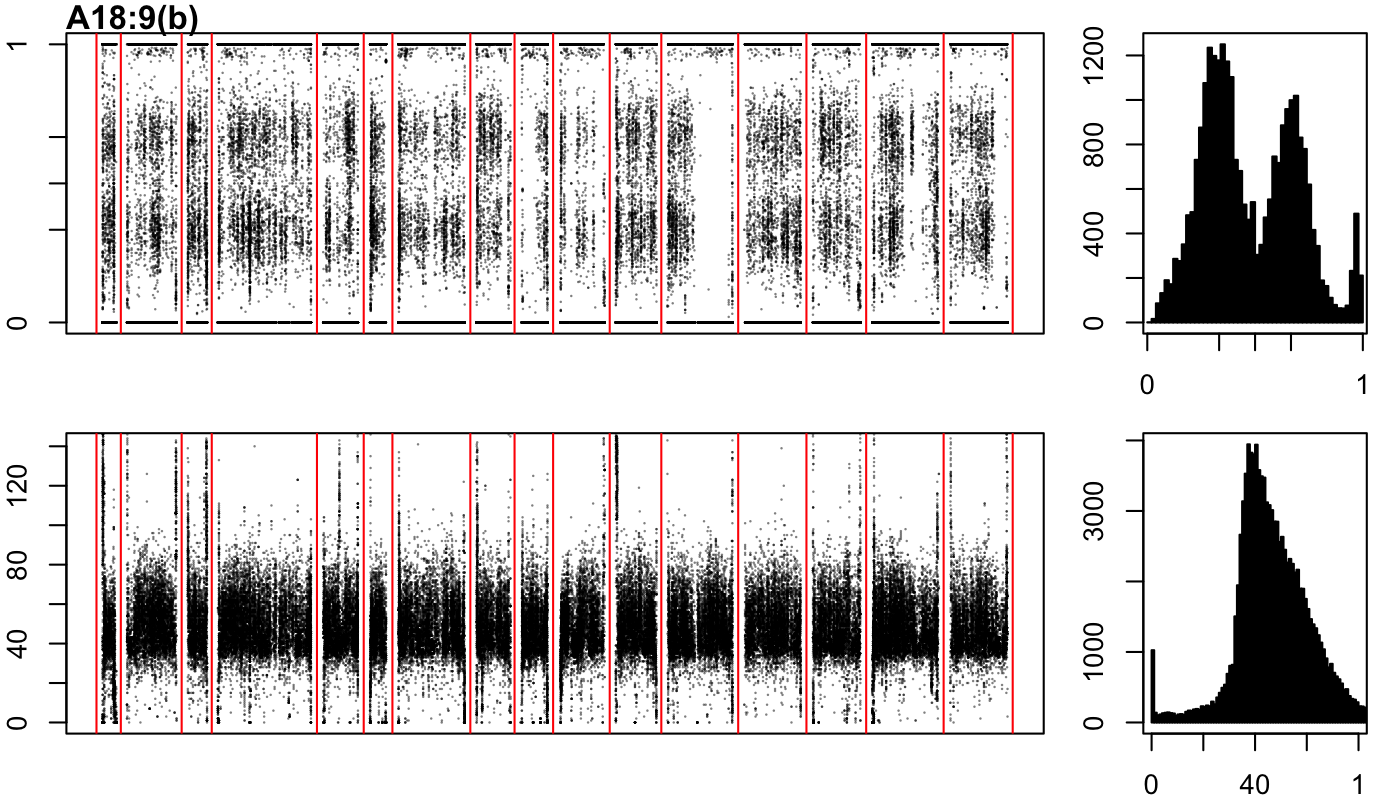

### A18-9-c.png

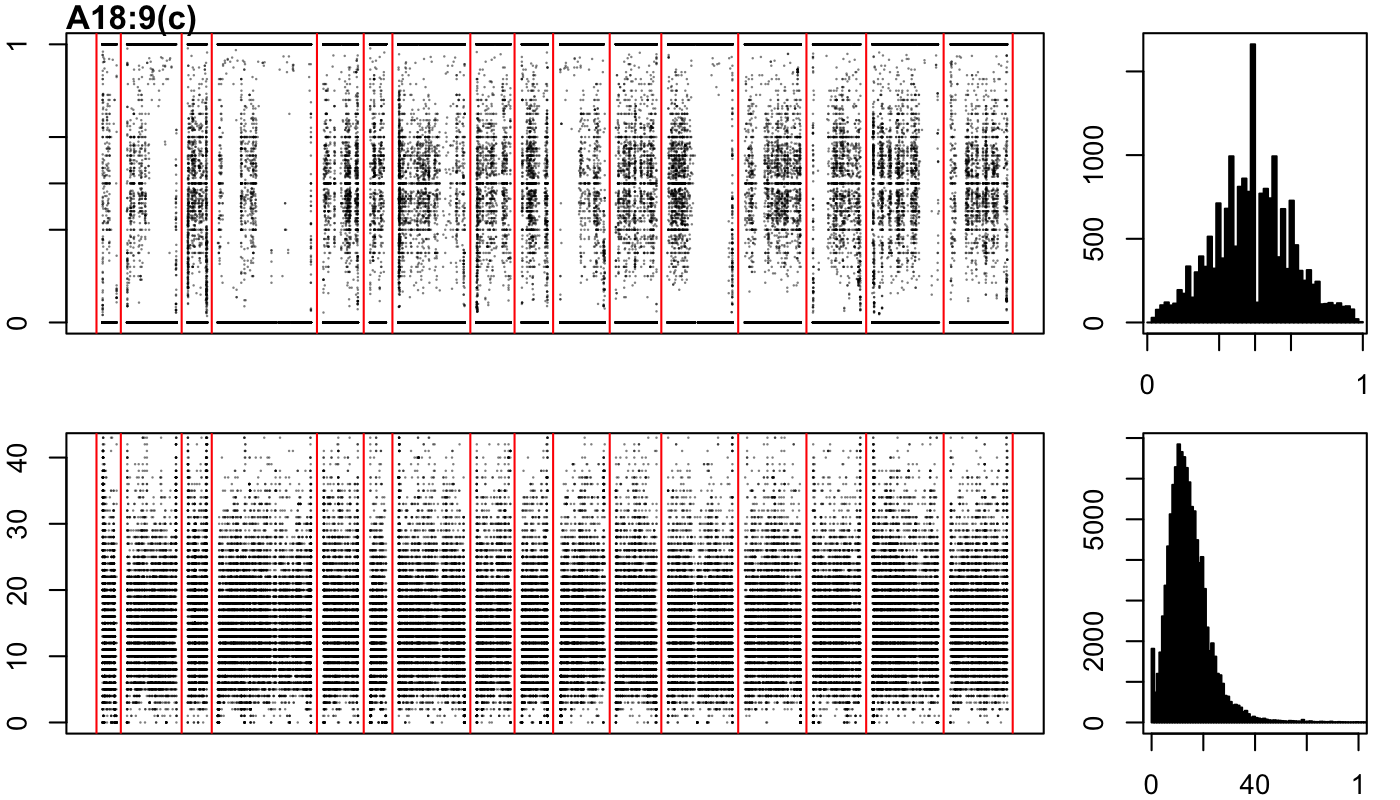

### A18-17-a.png

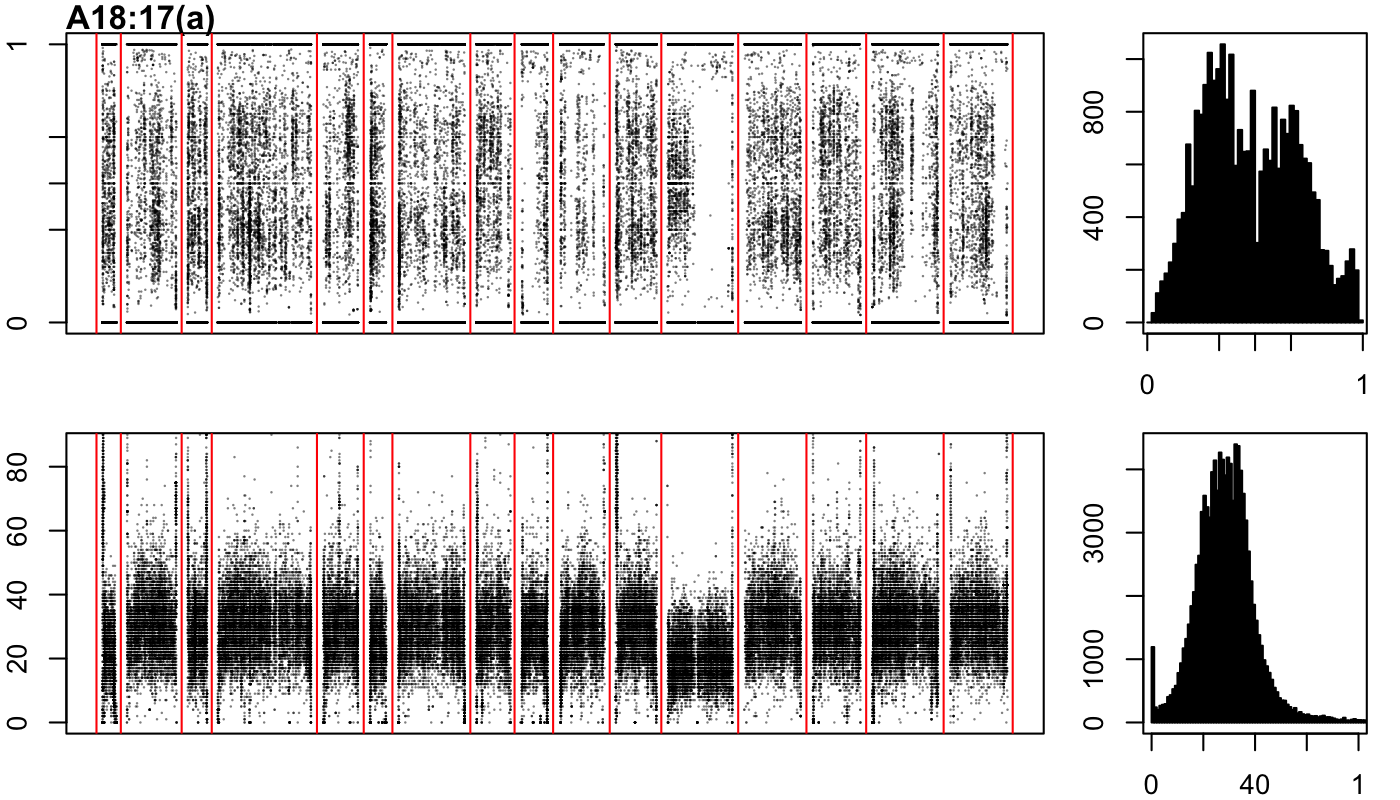

### A18-17-b.png

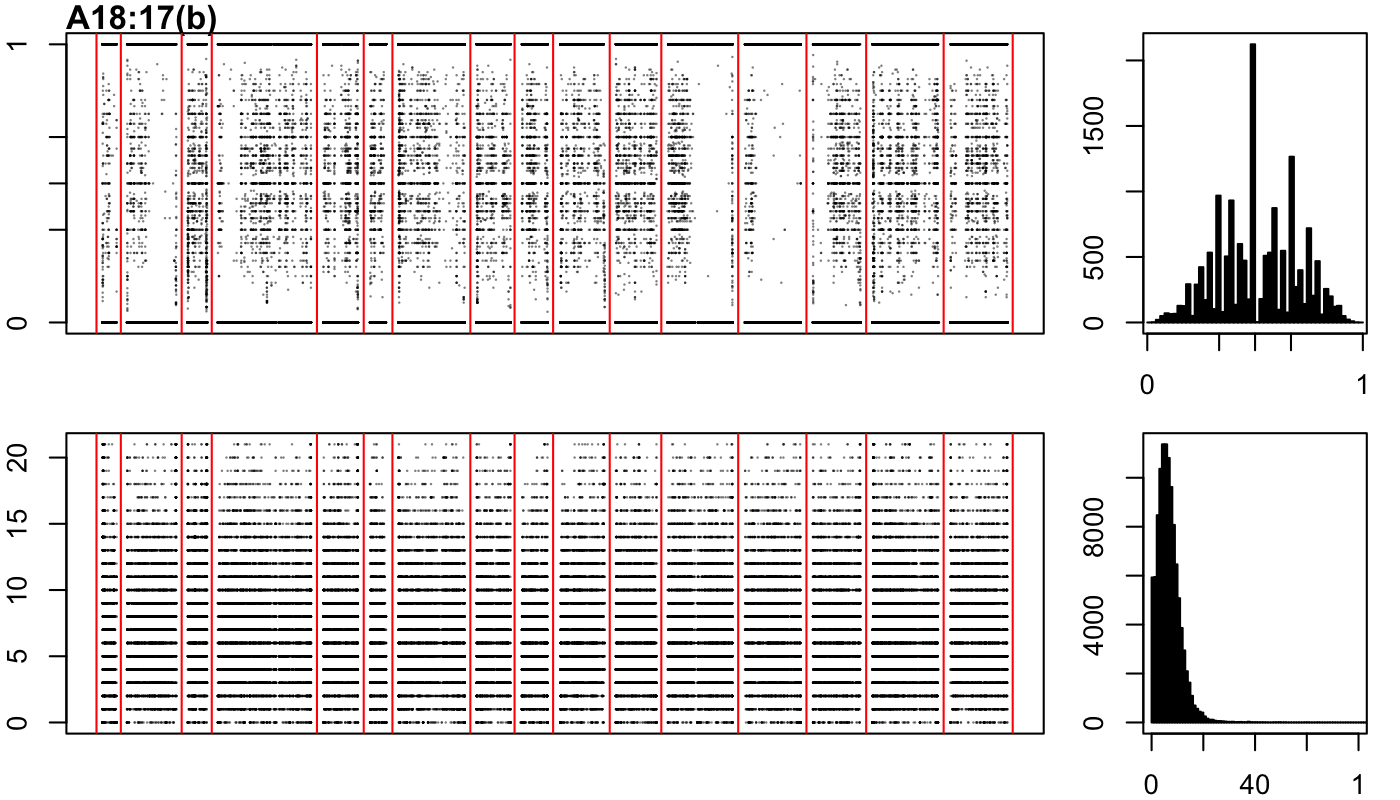

### A18-17-c.png

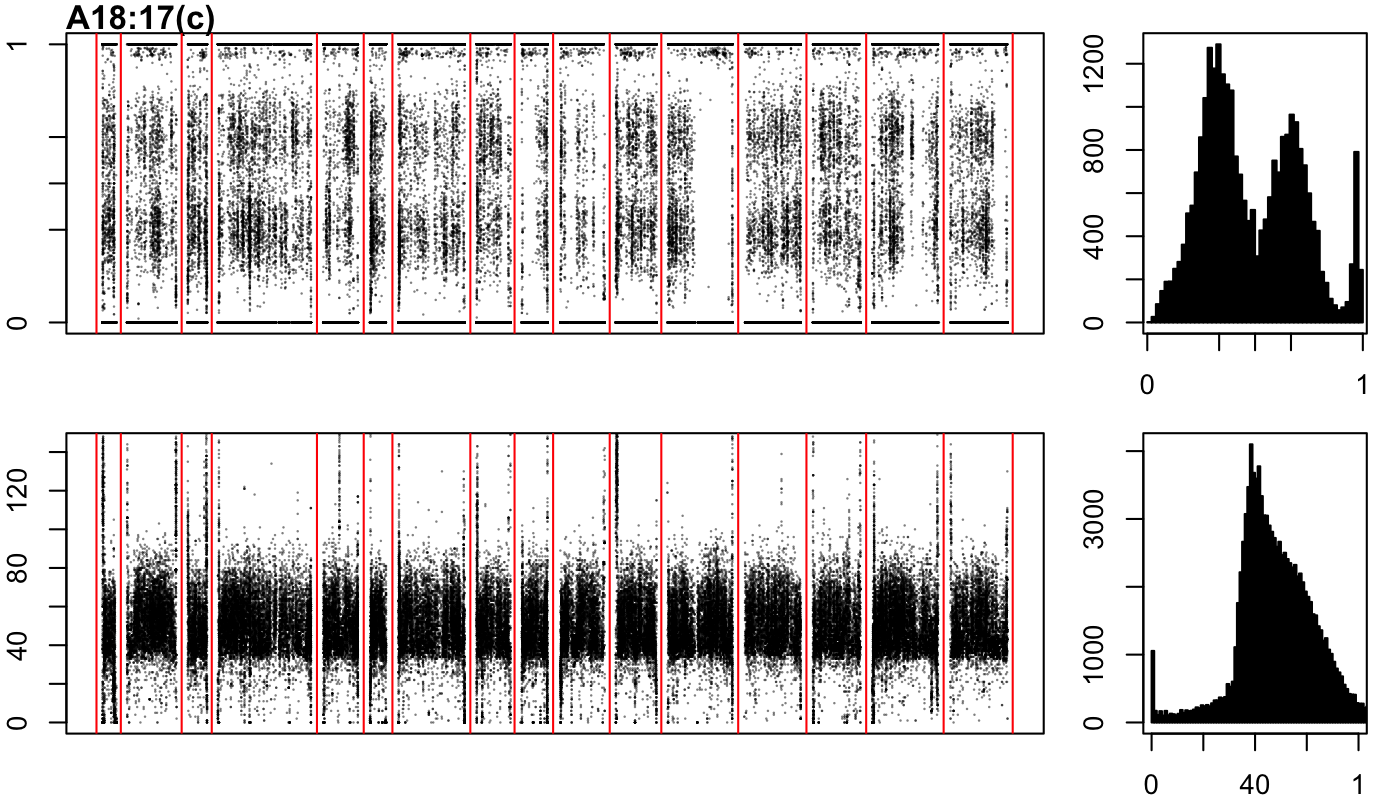

### A18-109-b.png

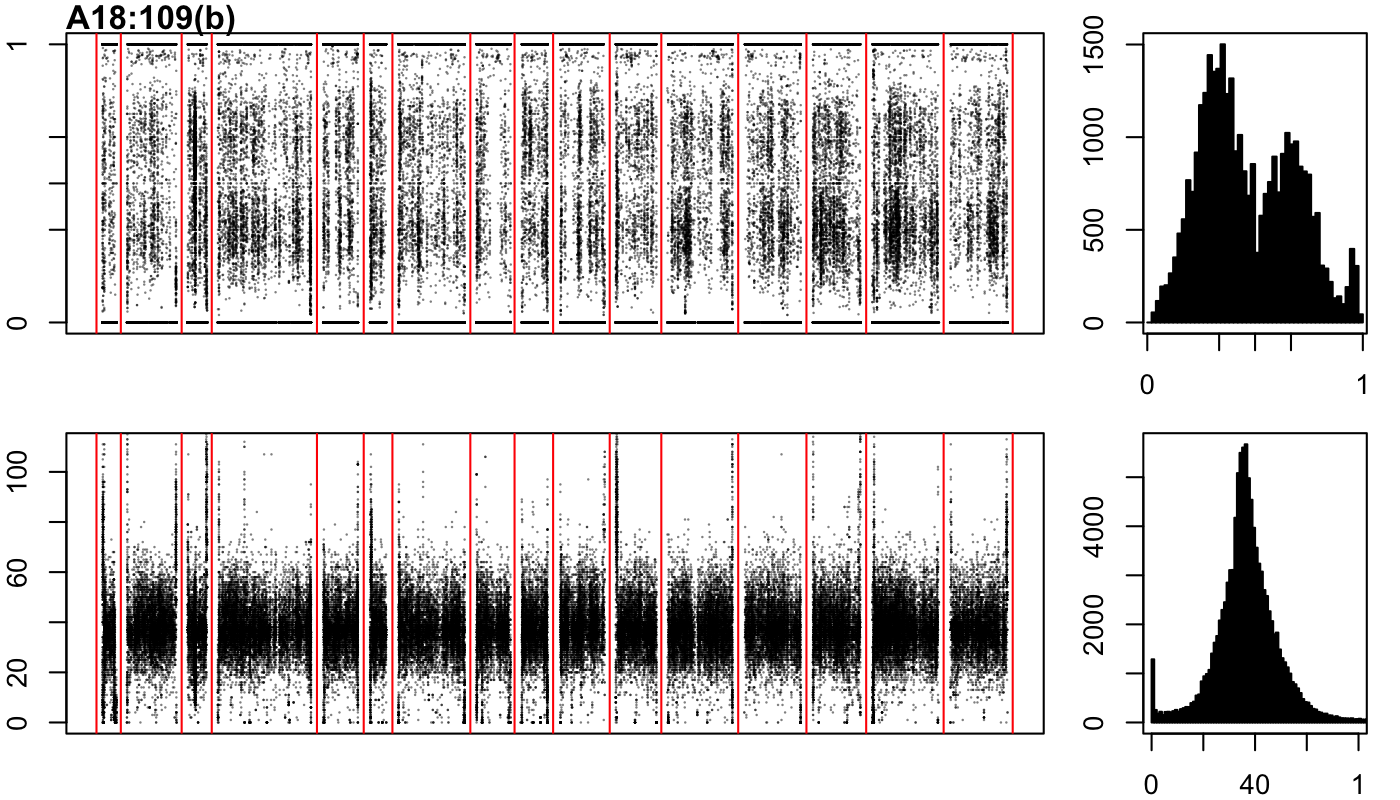

### A18-109-c.png

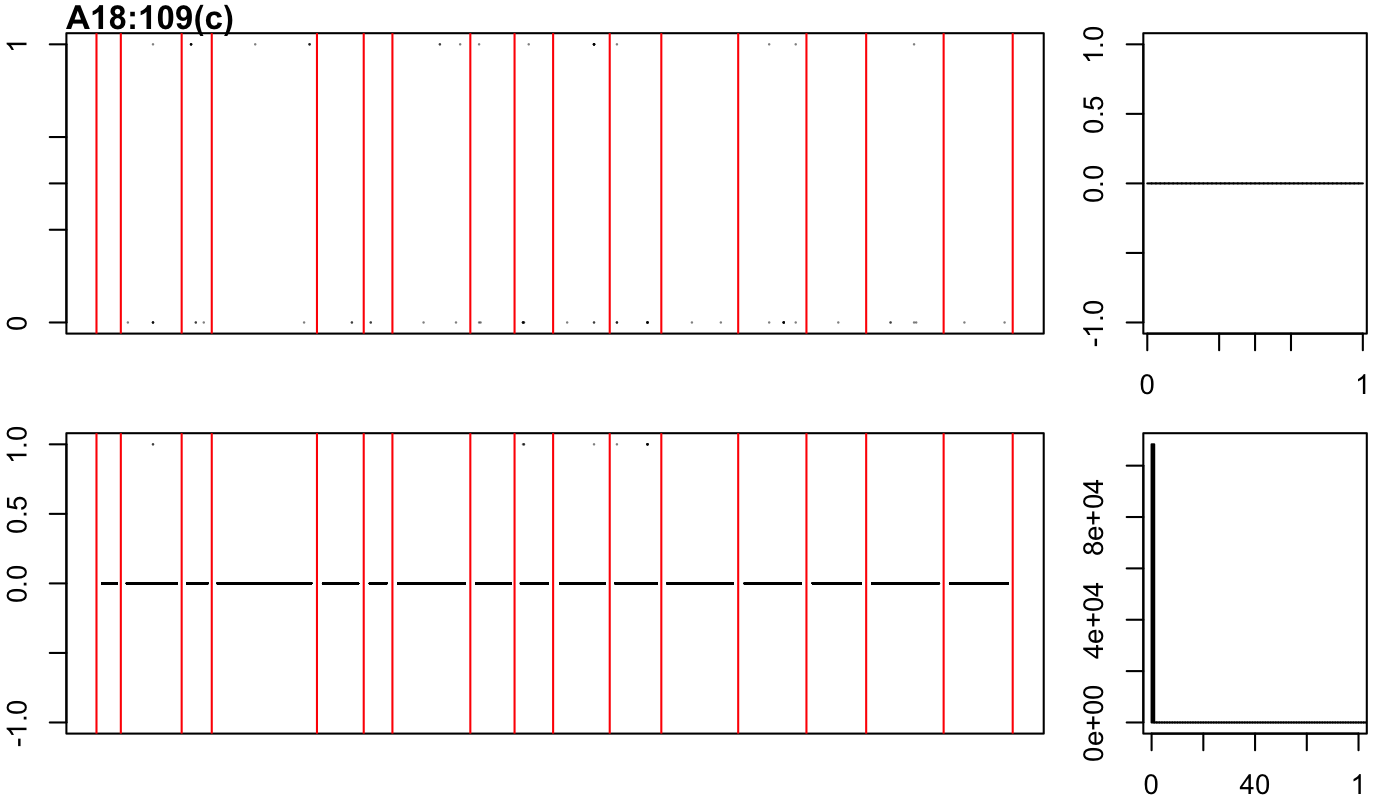

### A19-13-a.png

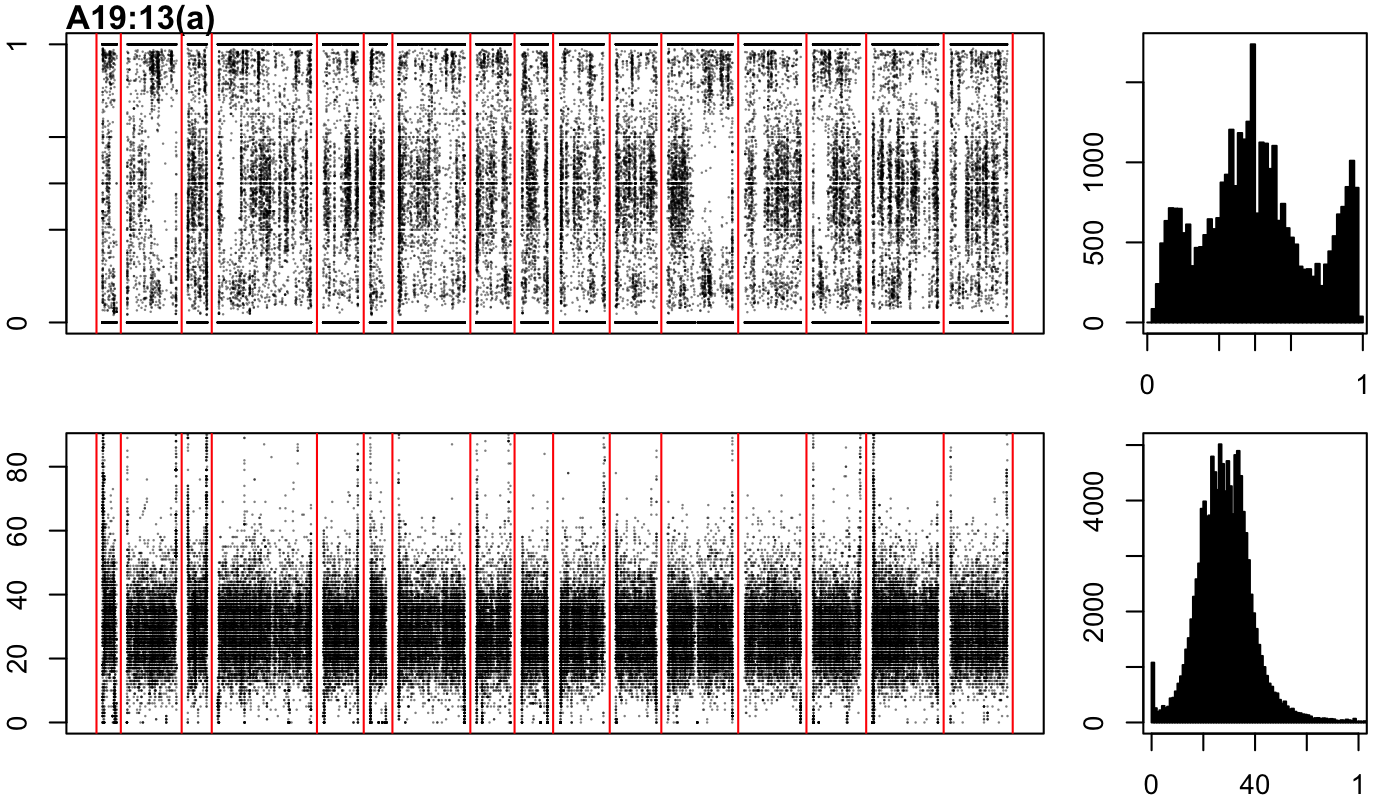

### A19-13-b.png

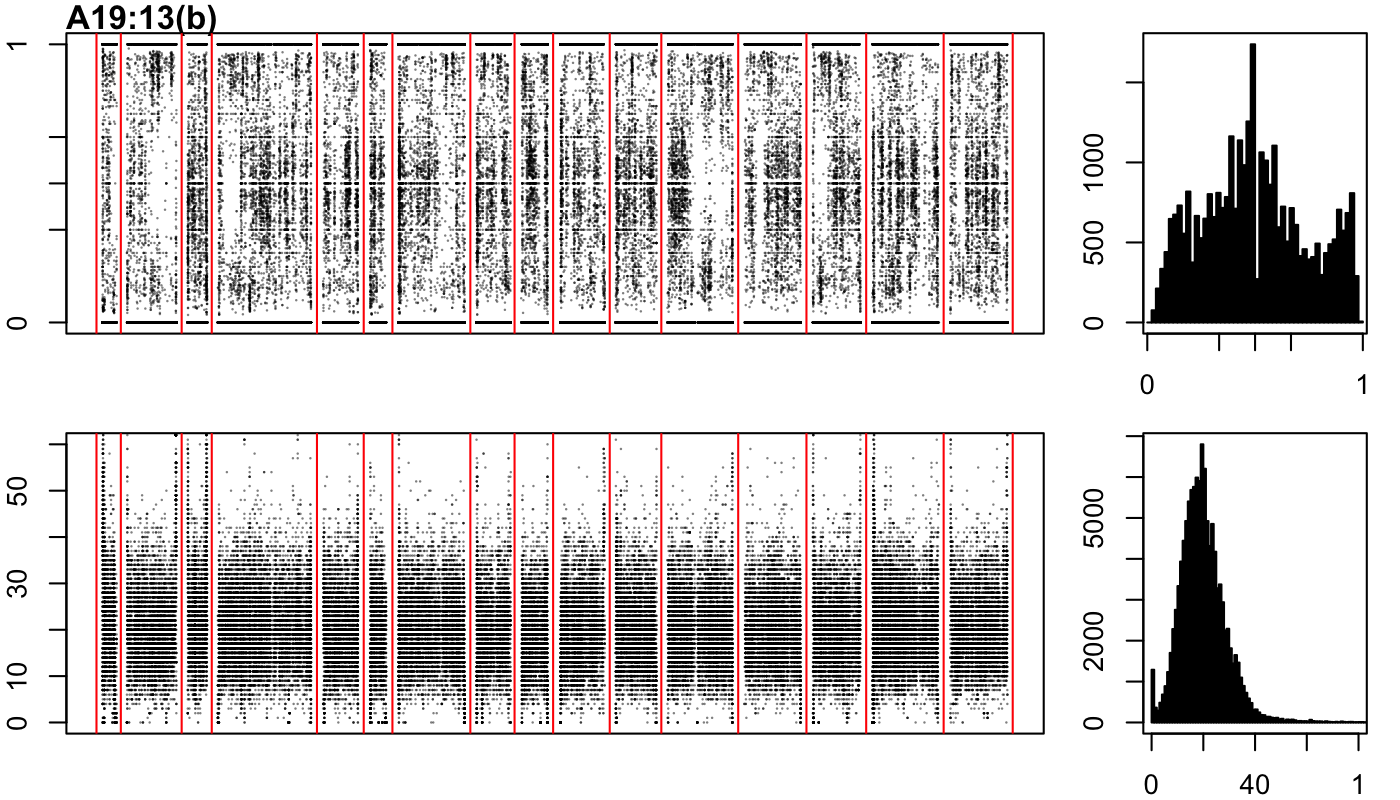

### A19-51-b.png

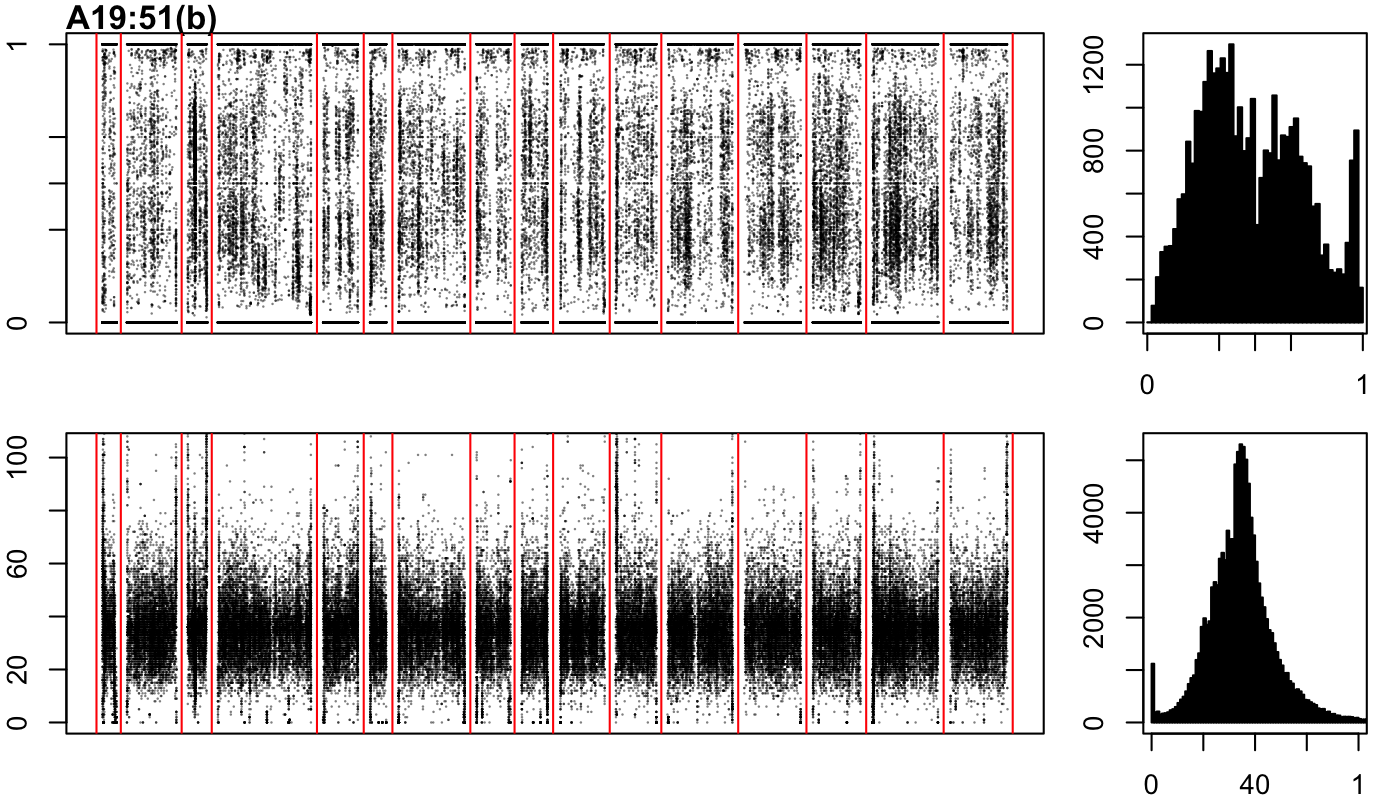

### A19-77-a.png

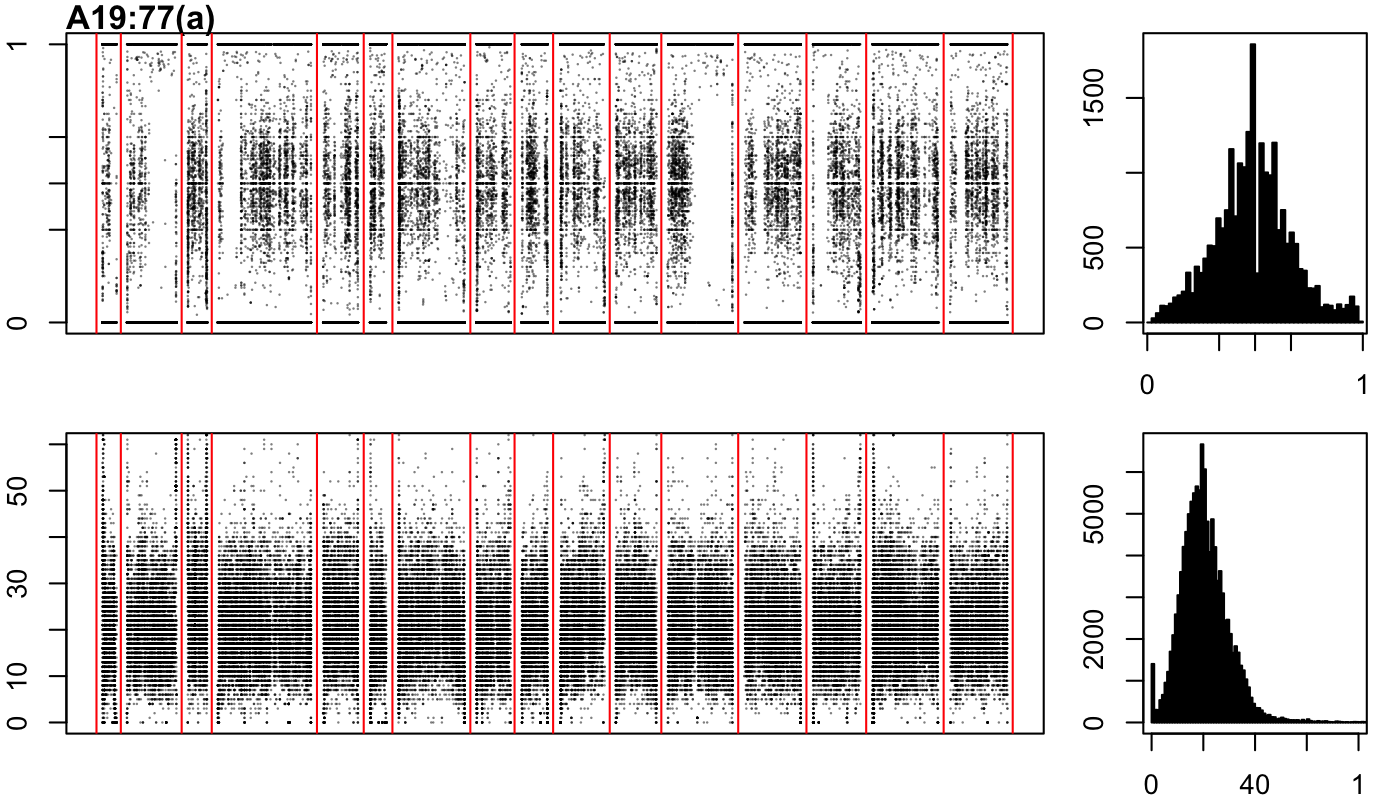

### A19-92-a.png

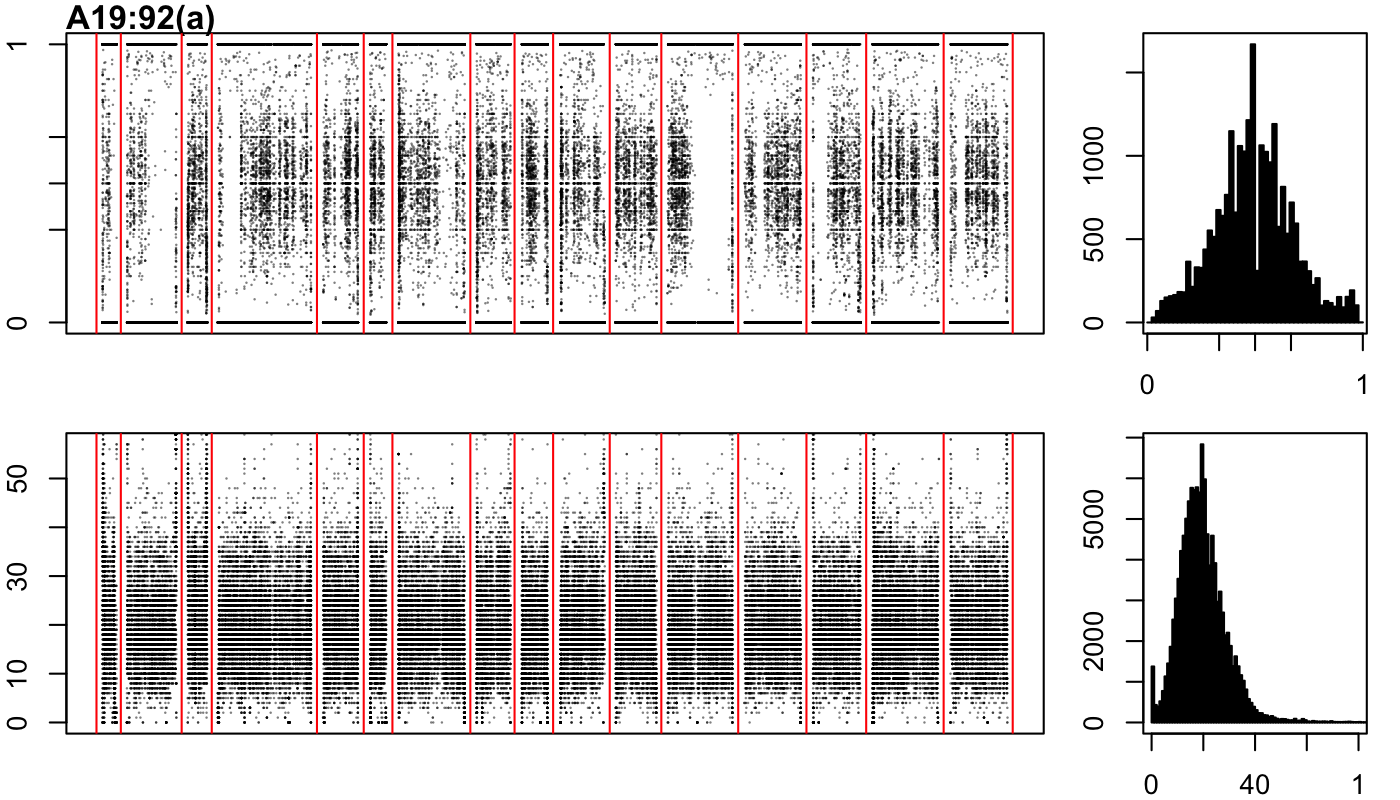

### A19-92-b.png

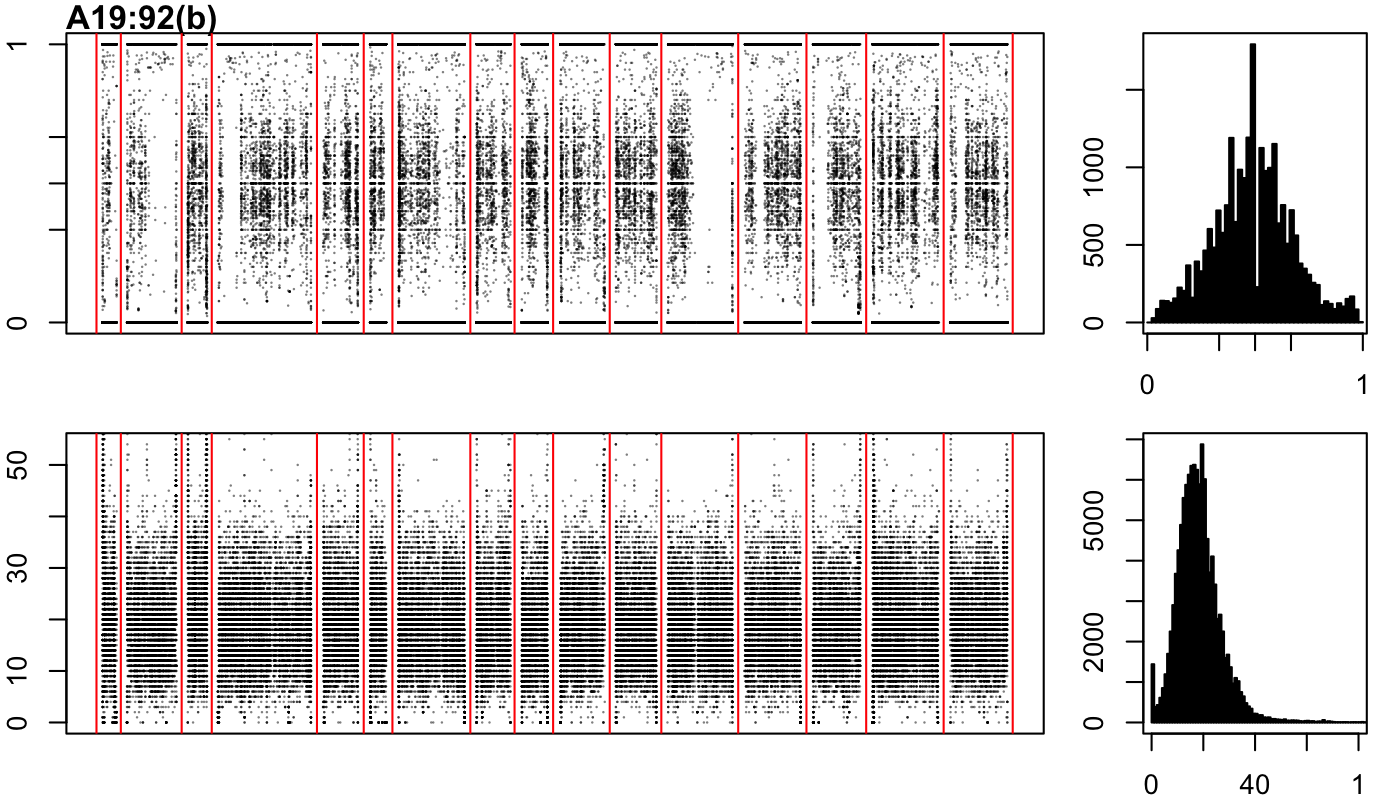

### A19-134-a.png

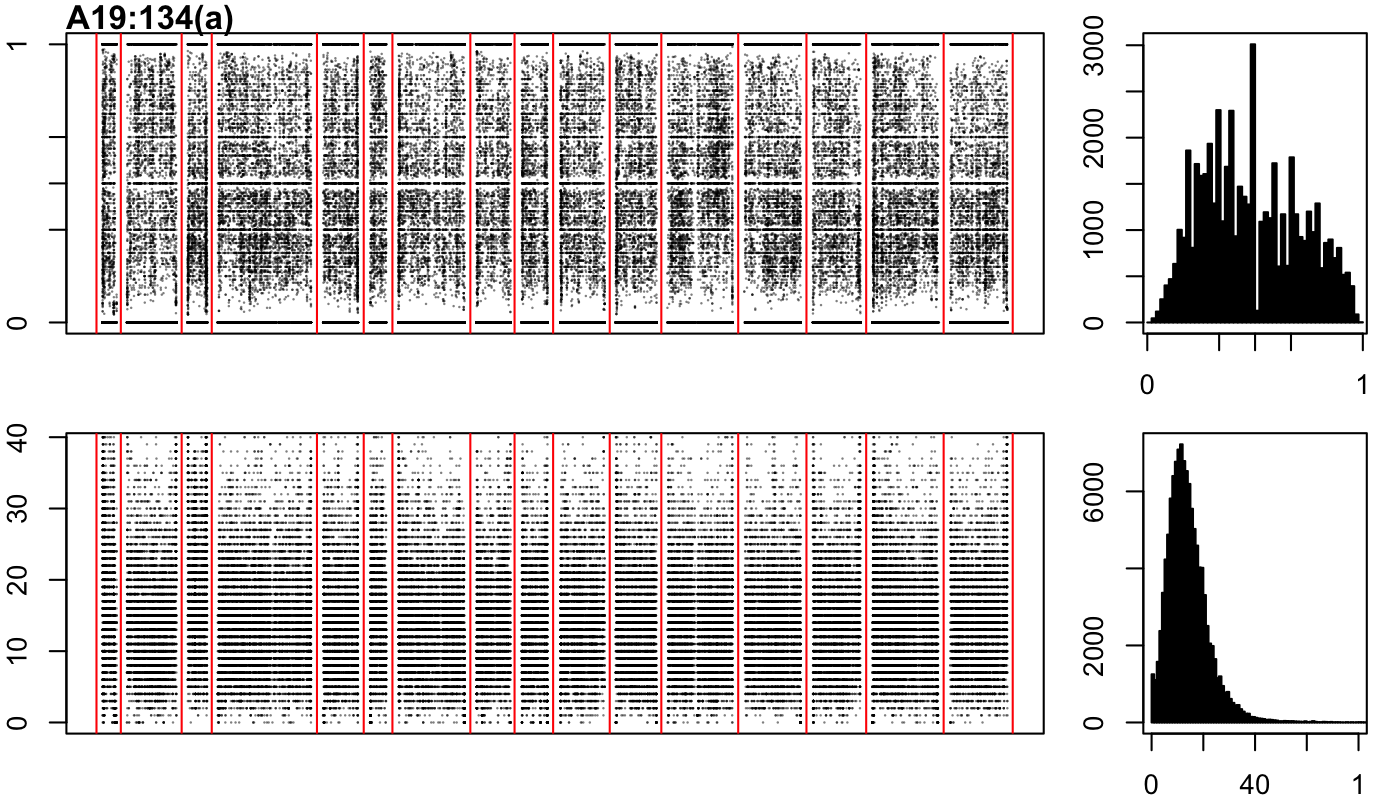

### A19-134-b.png

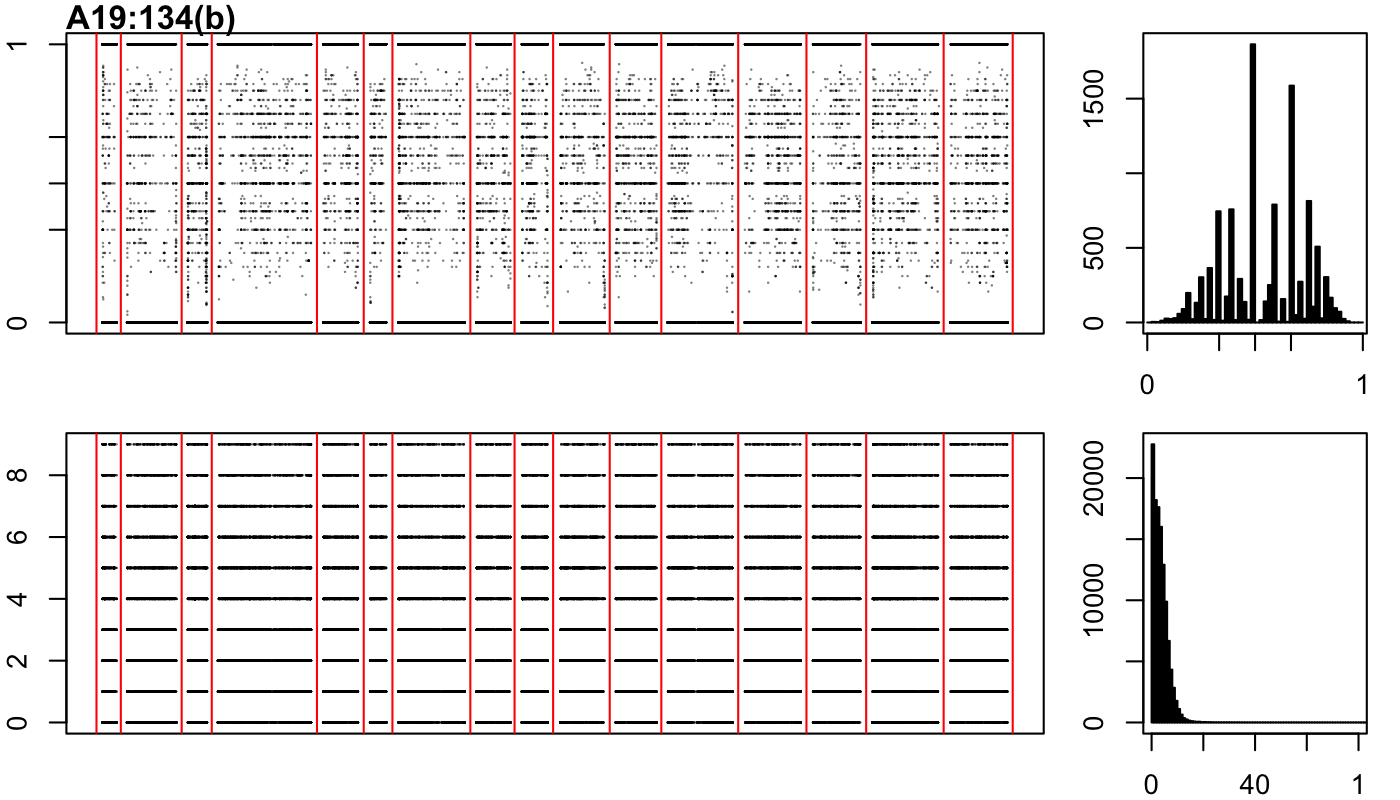

### B18-1-a.png

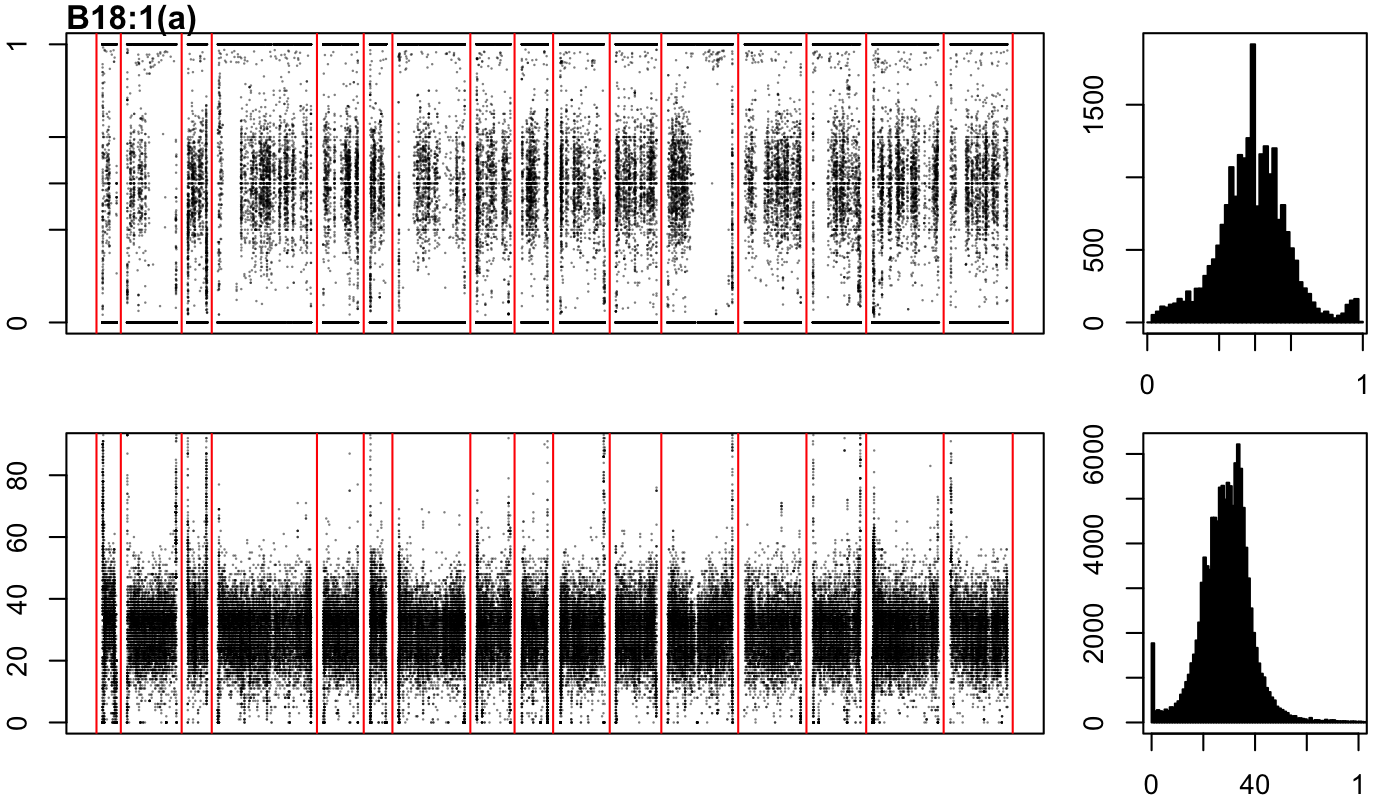

### B18-1-b.png

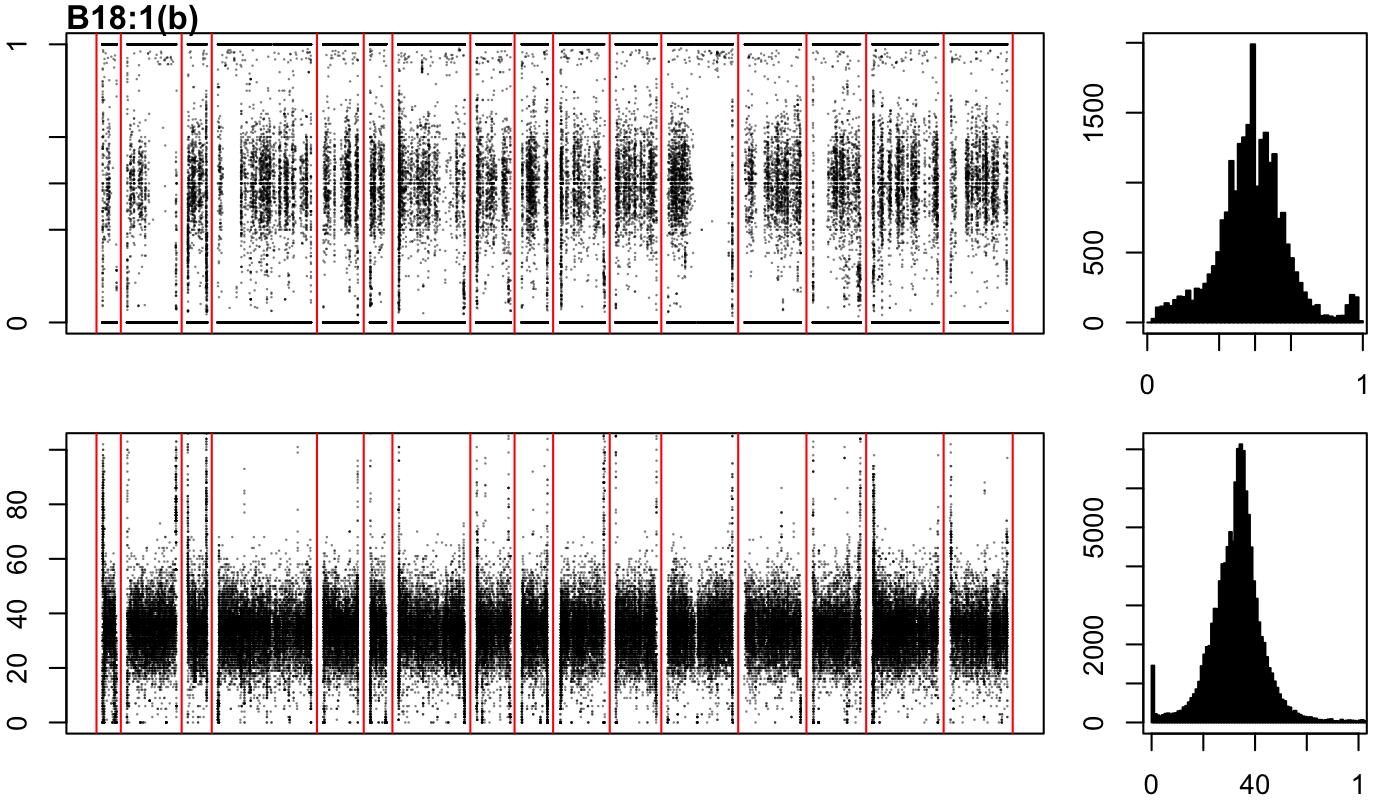

### B18-1-c.png

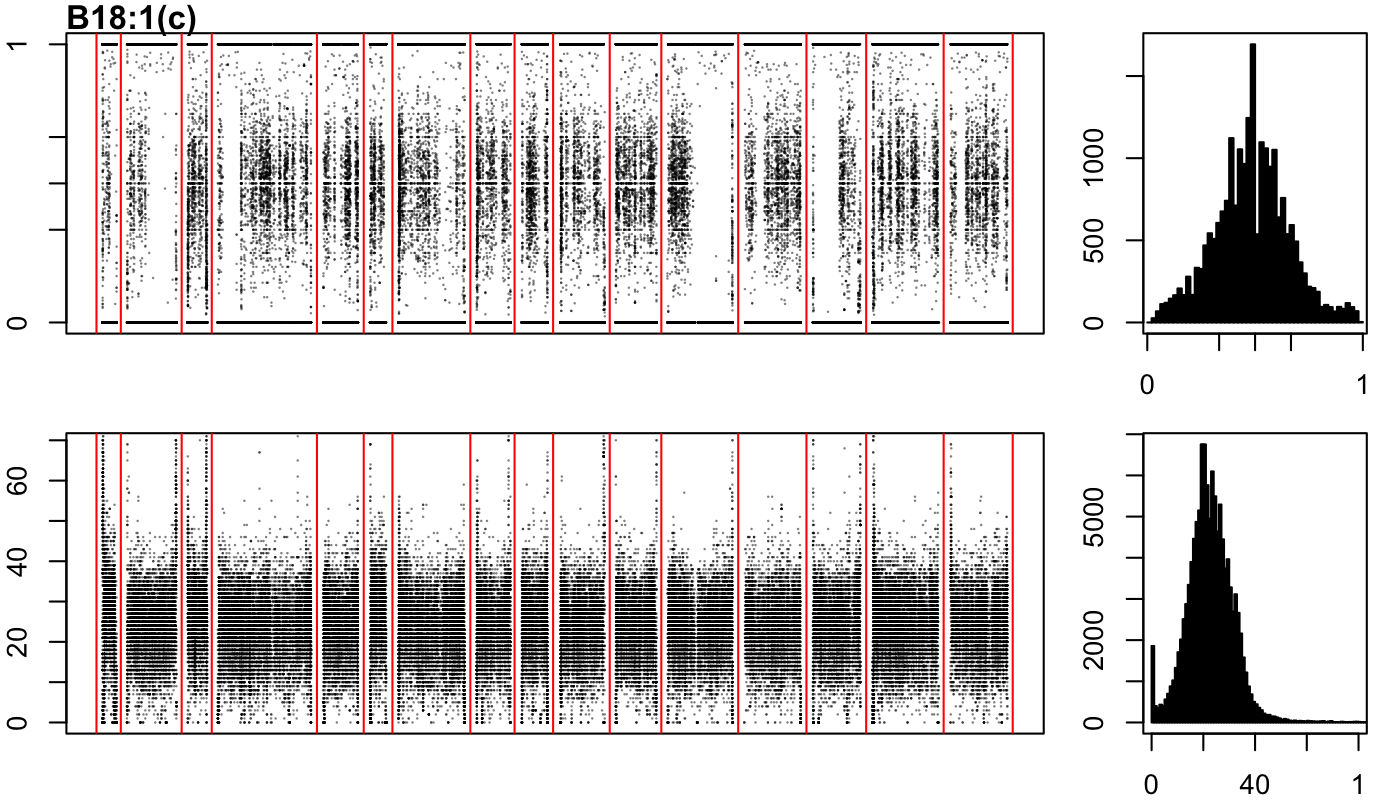

### B18-7-a.png

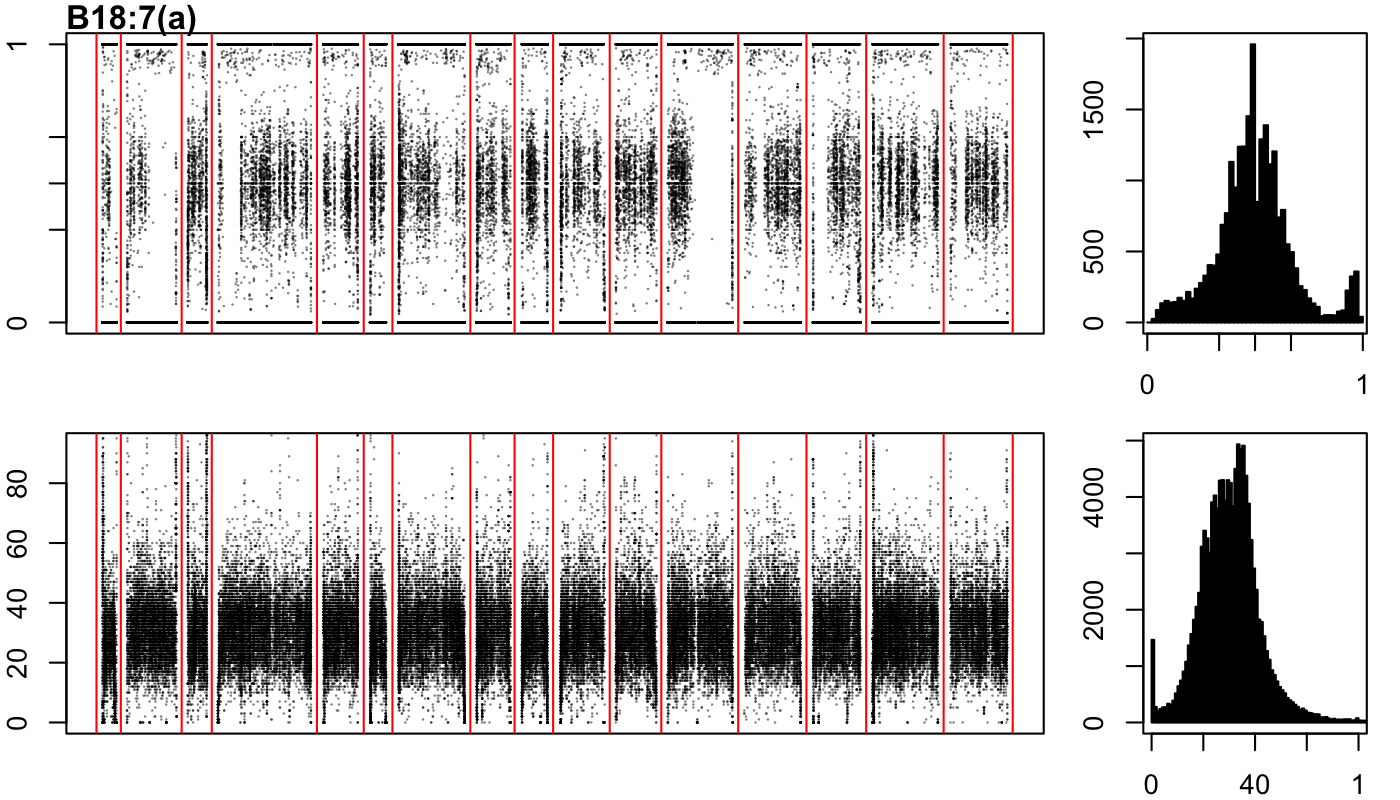

### B18-7-b.png

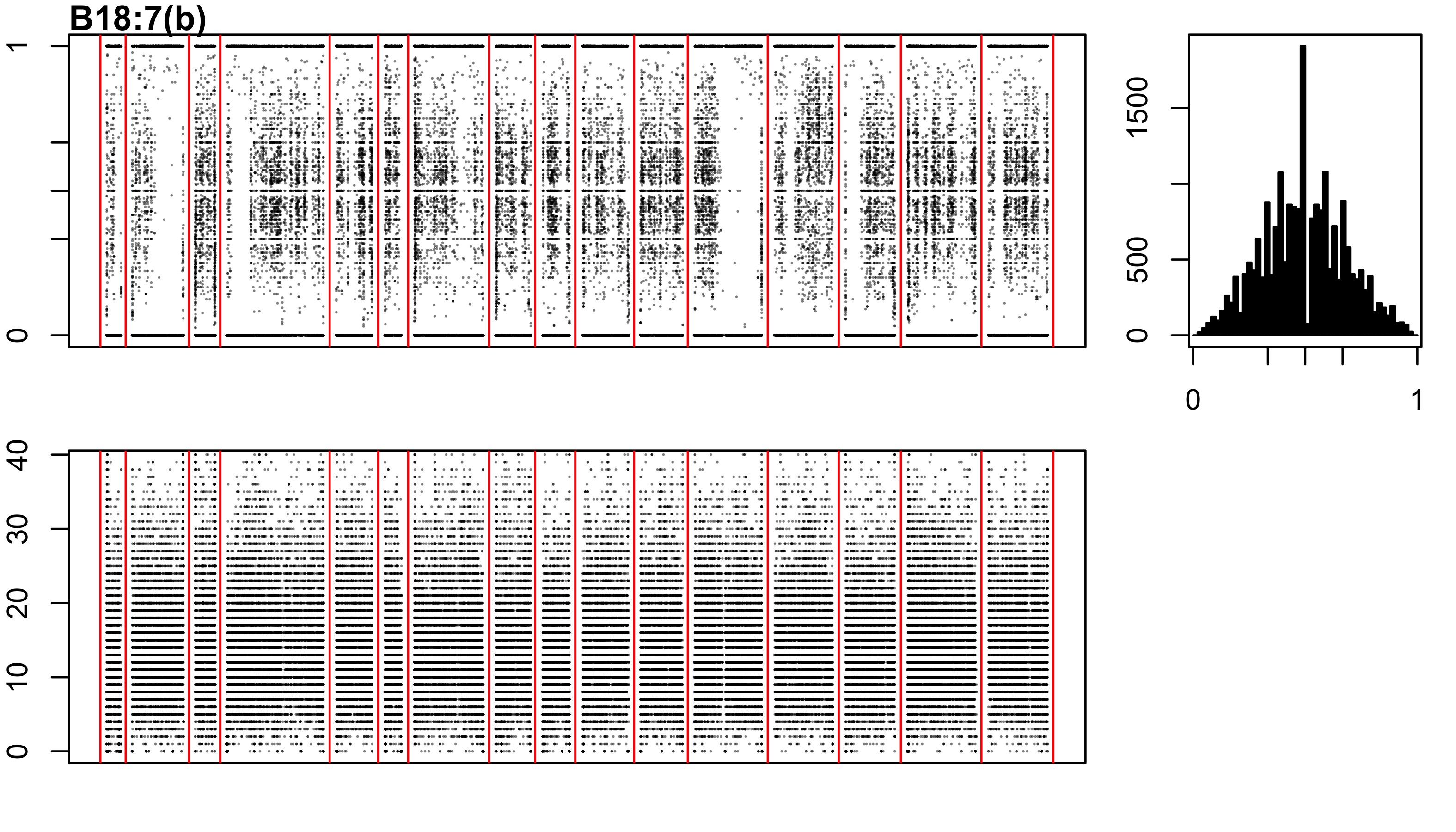

### B18-27-b.png

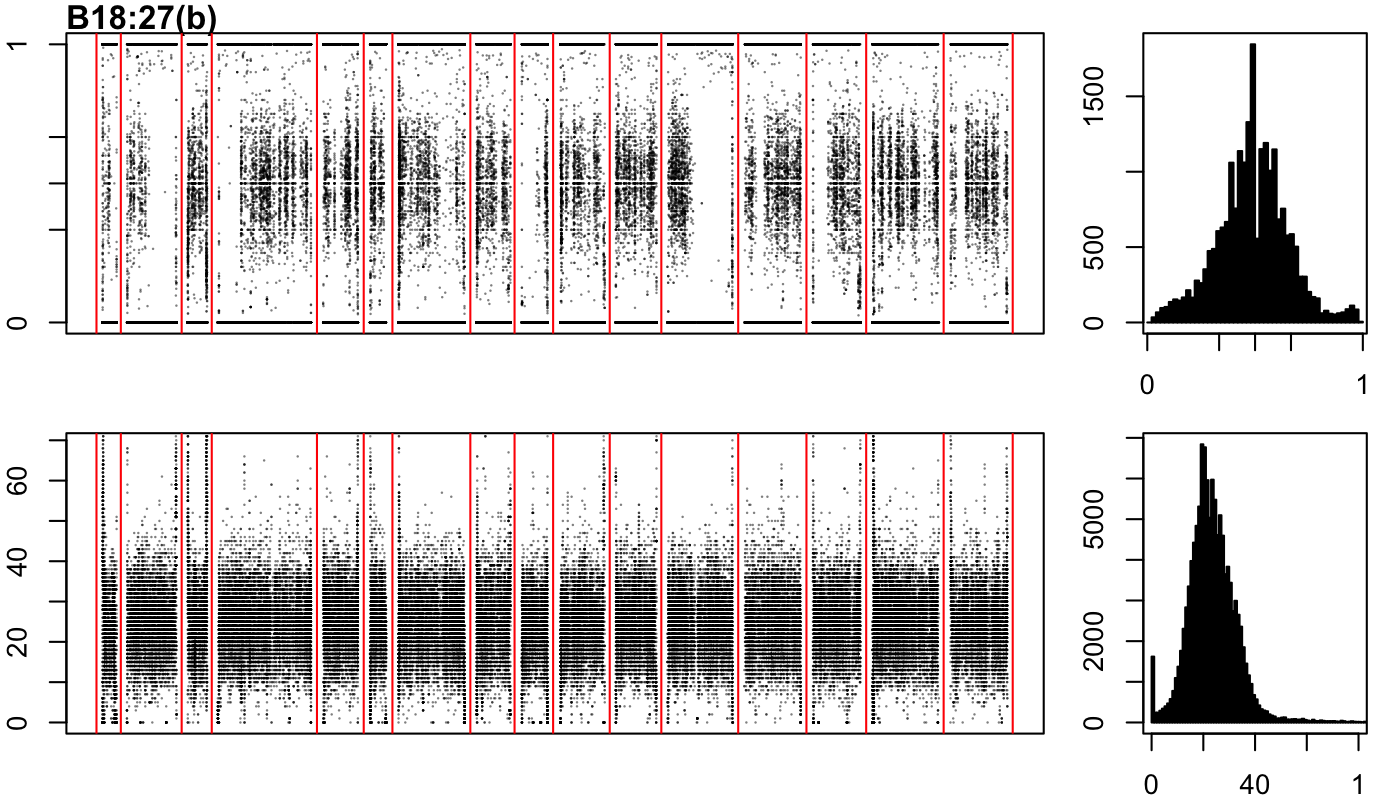

### B18-27-c.png

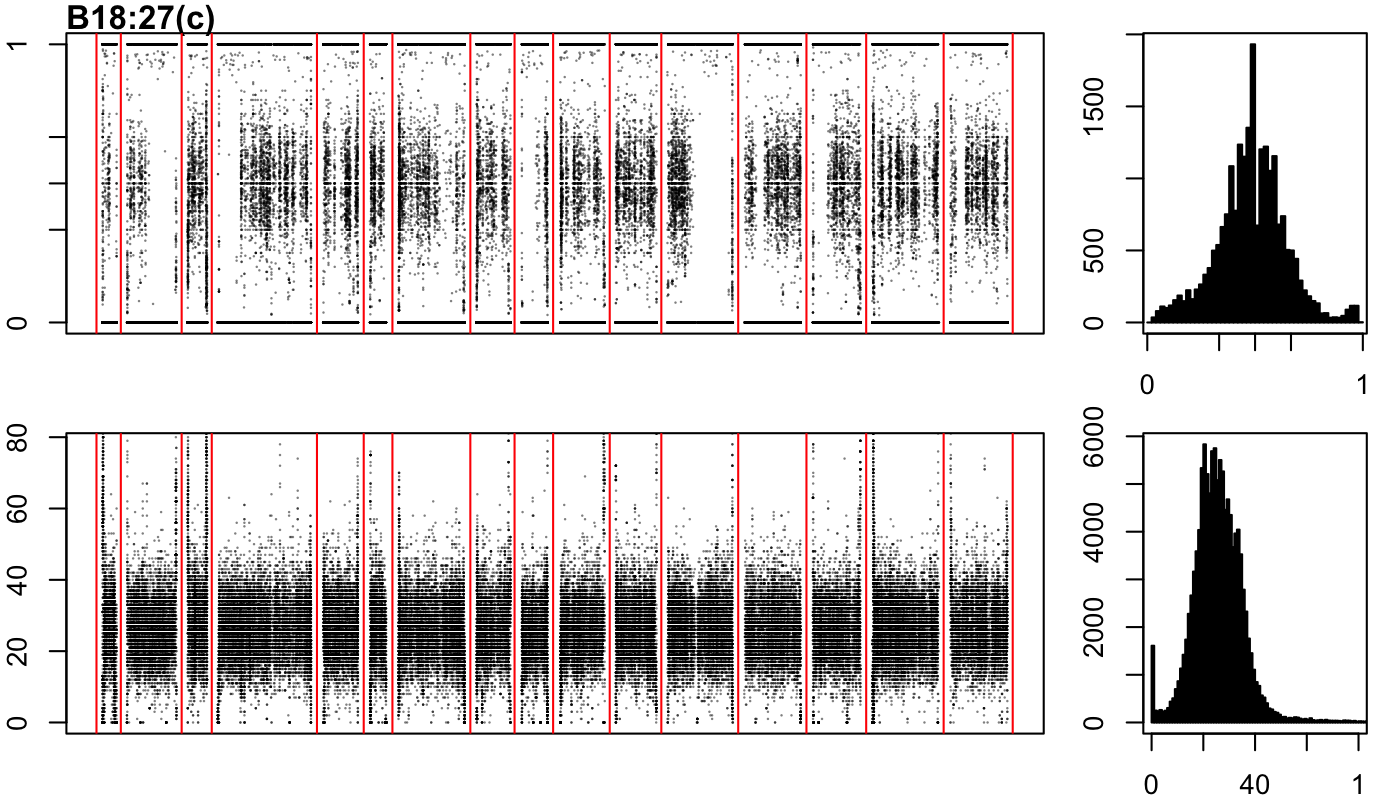

### B18-47-a.png

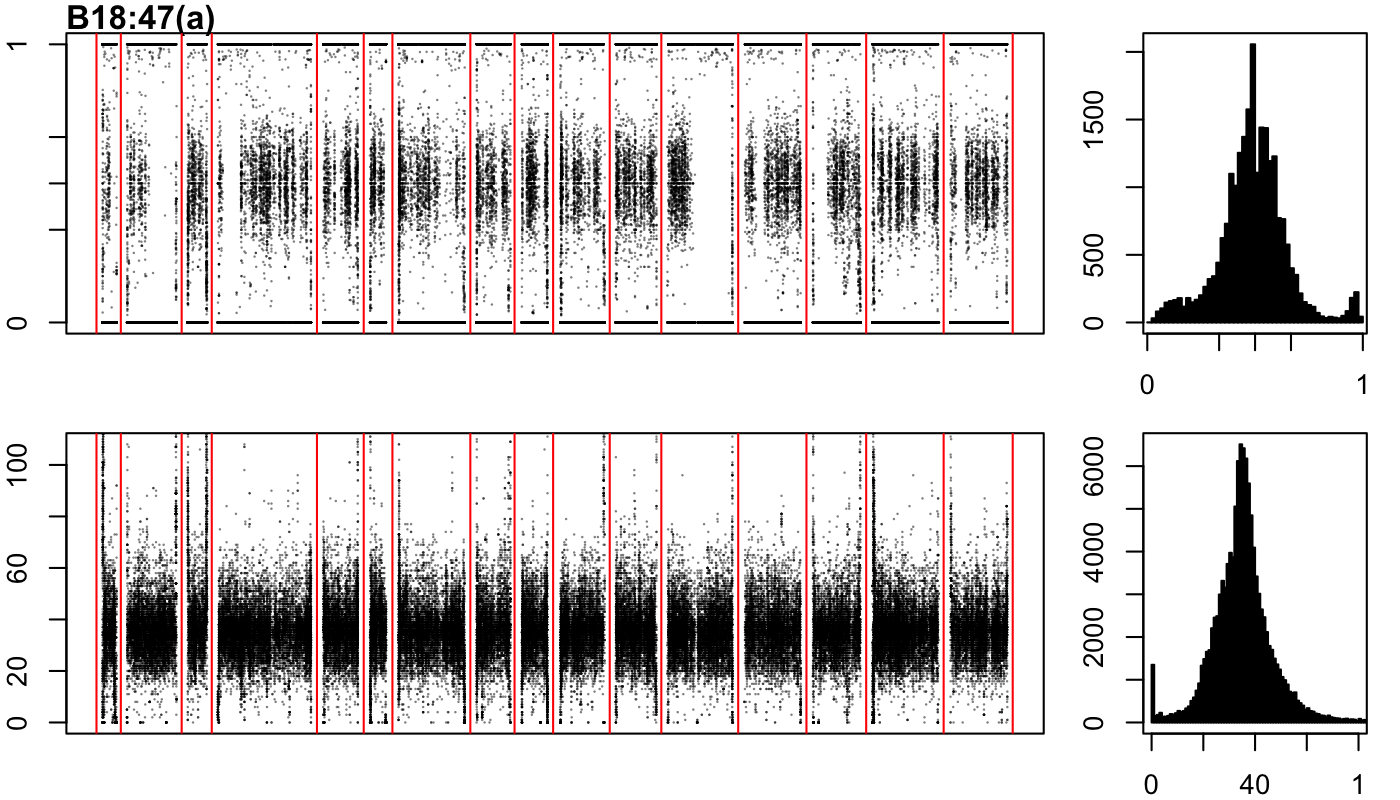

### B18-47-b.png

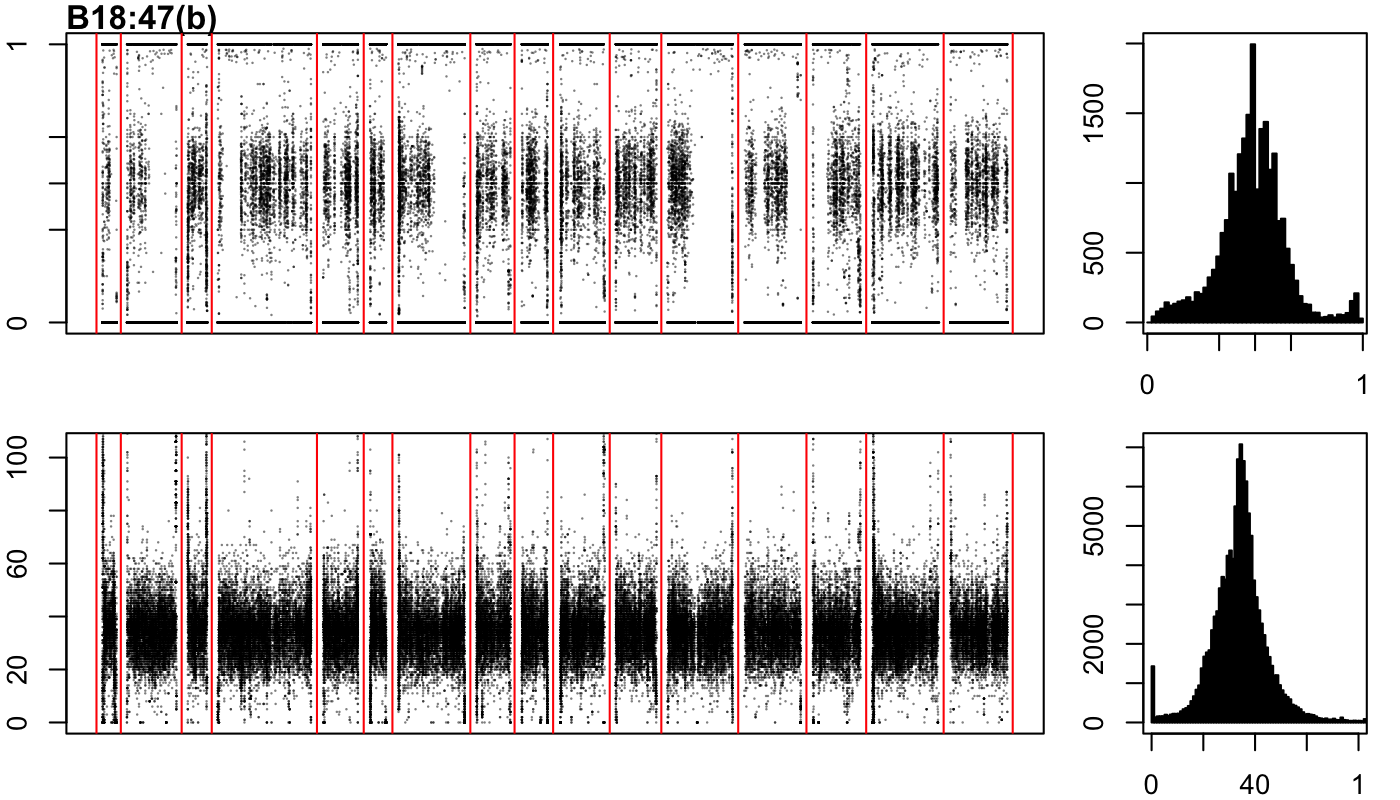

### B18-47-c.png

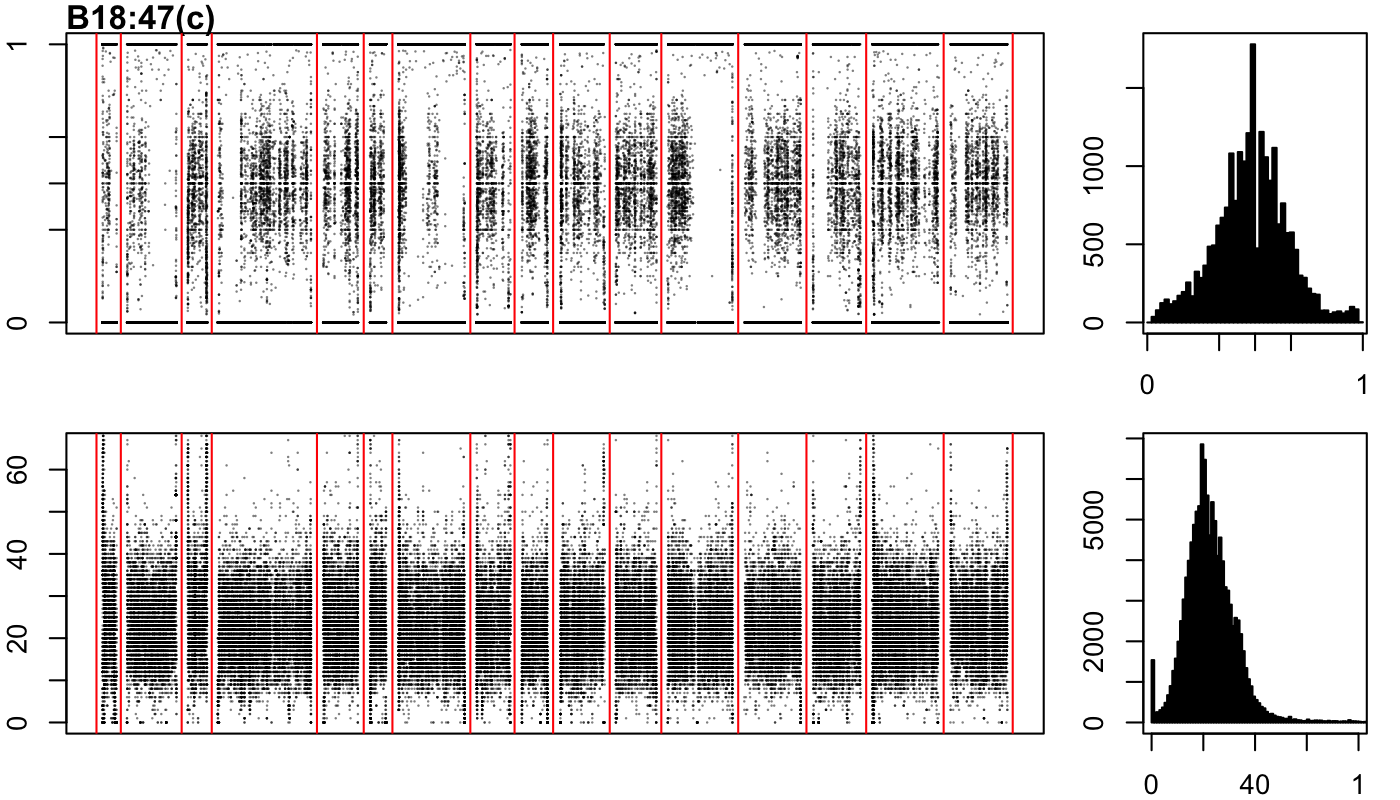

### B18-87-a.png

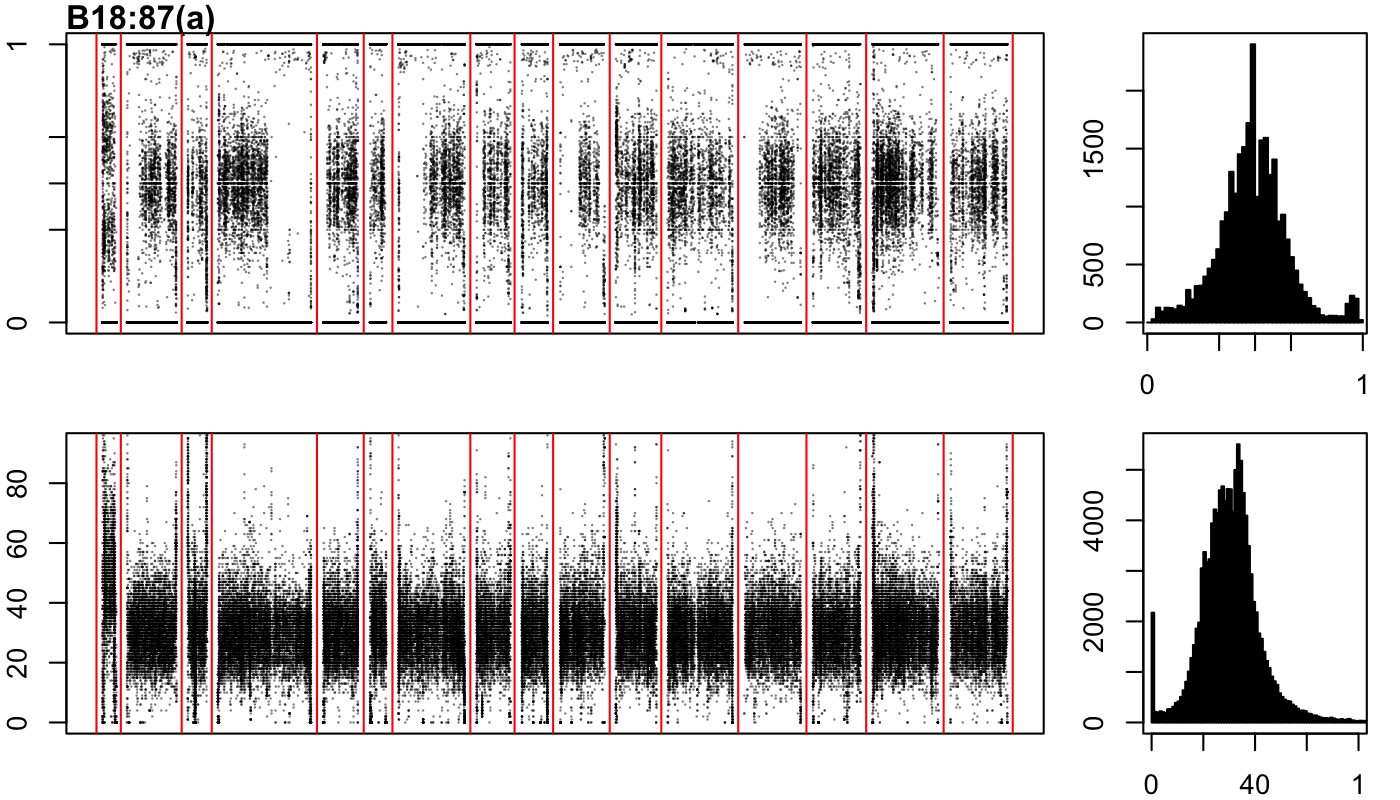

### B18-87-b.png

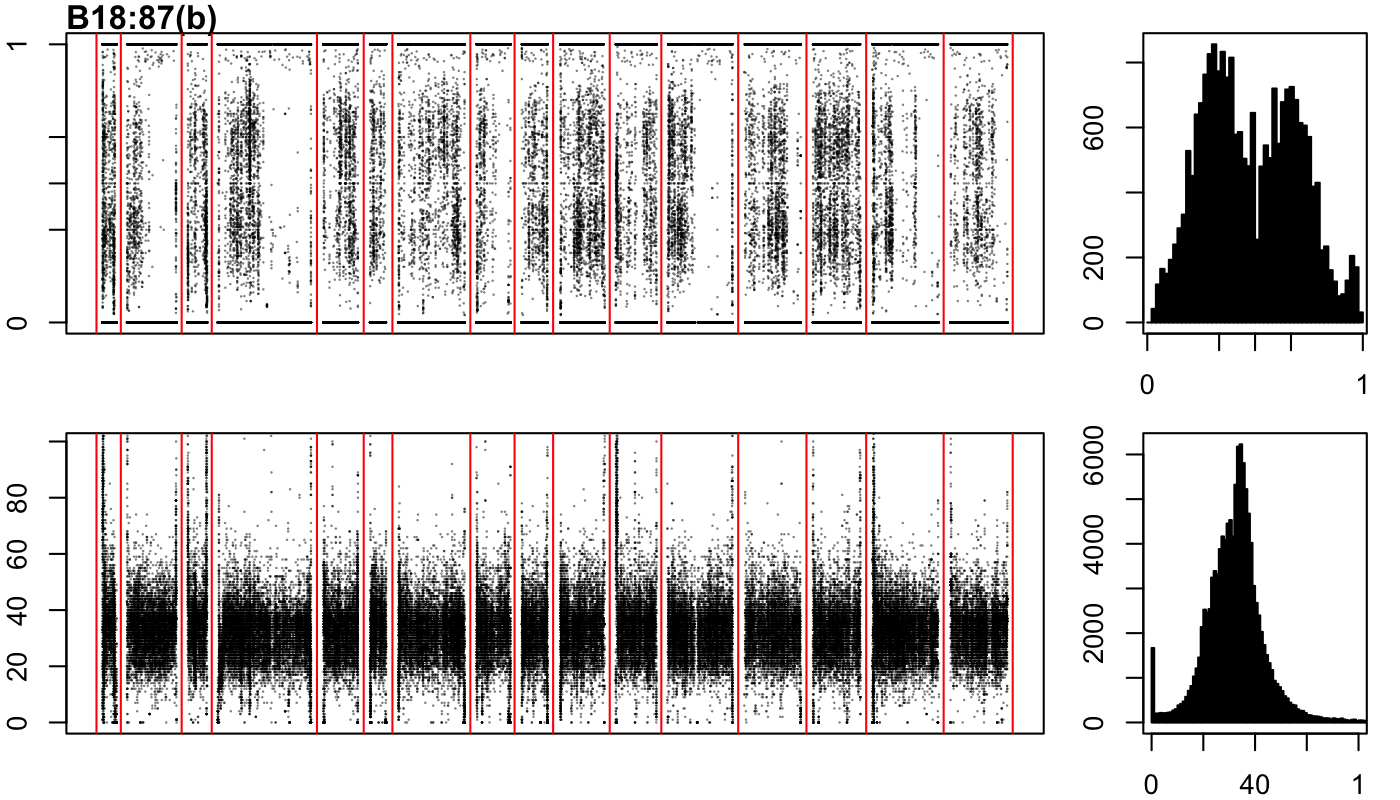

### B18-87-c.png

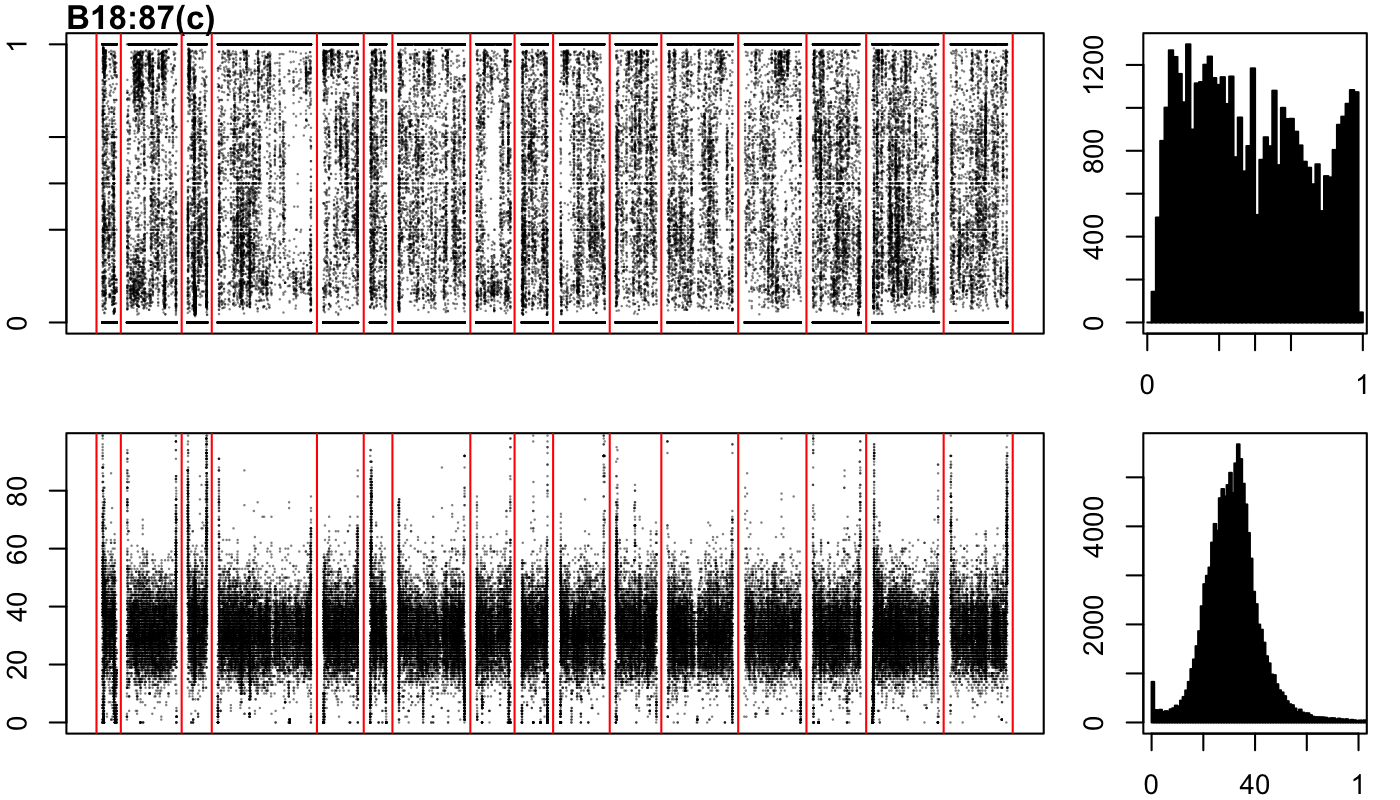

### B18-109-a.png

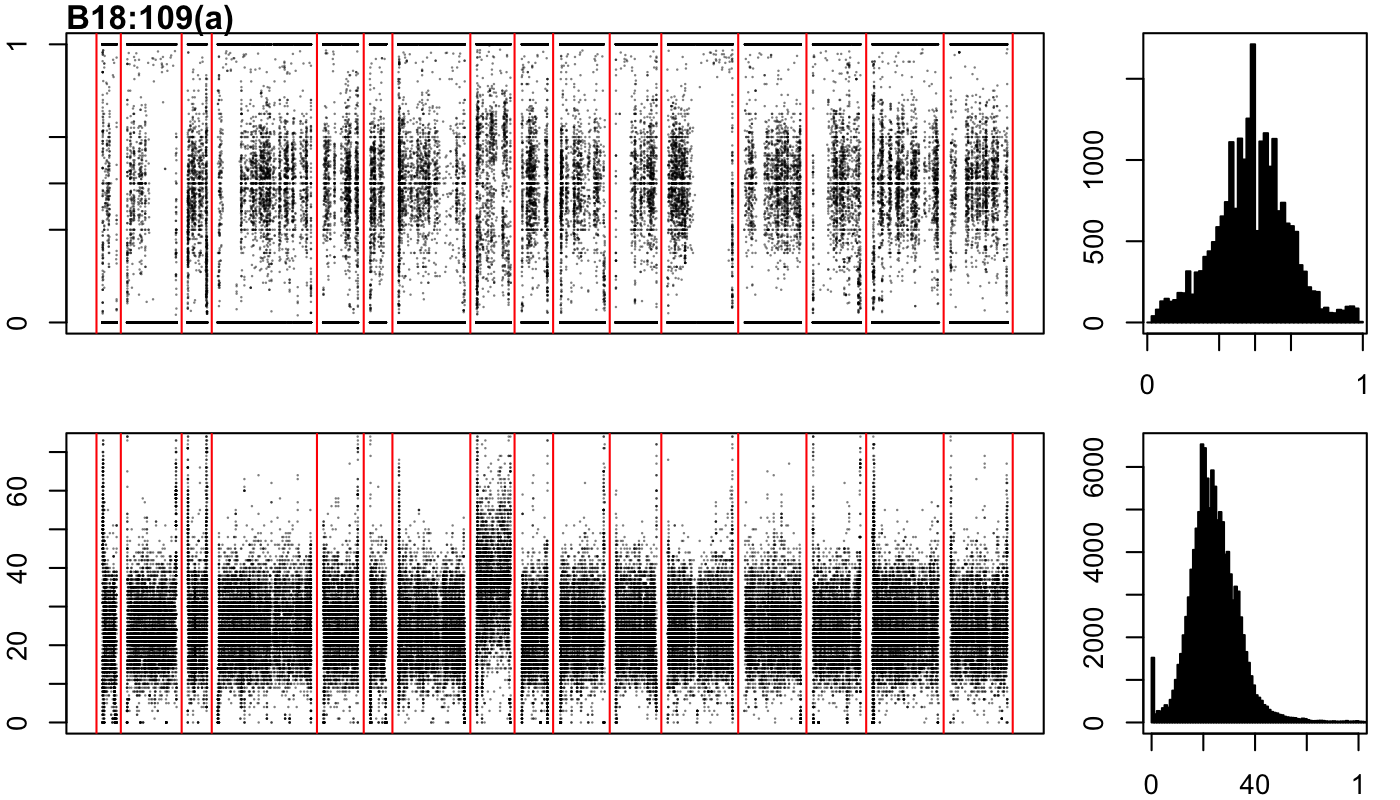

### B18-109-b.png

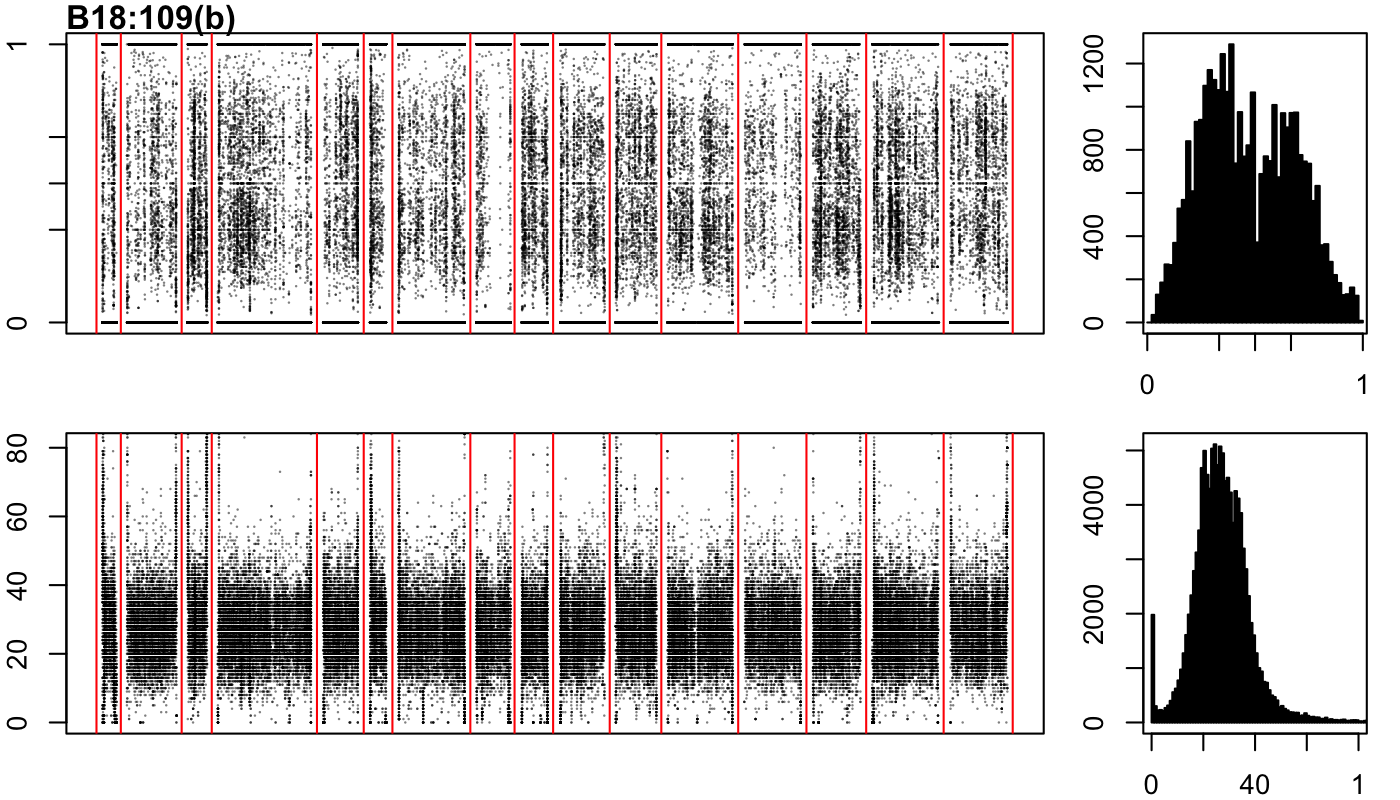

### B18-109-c.png

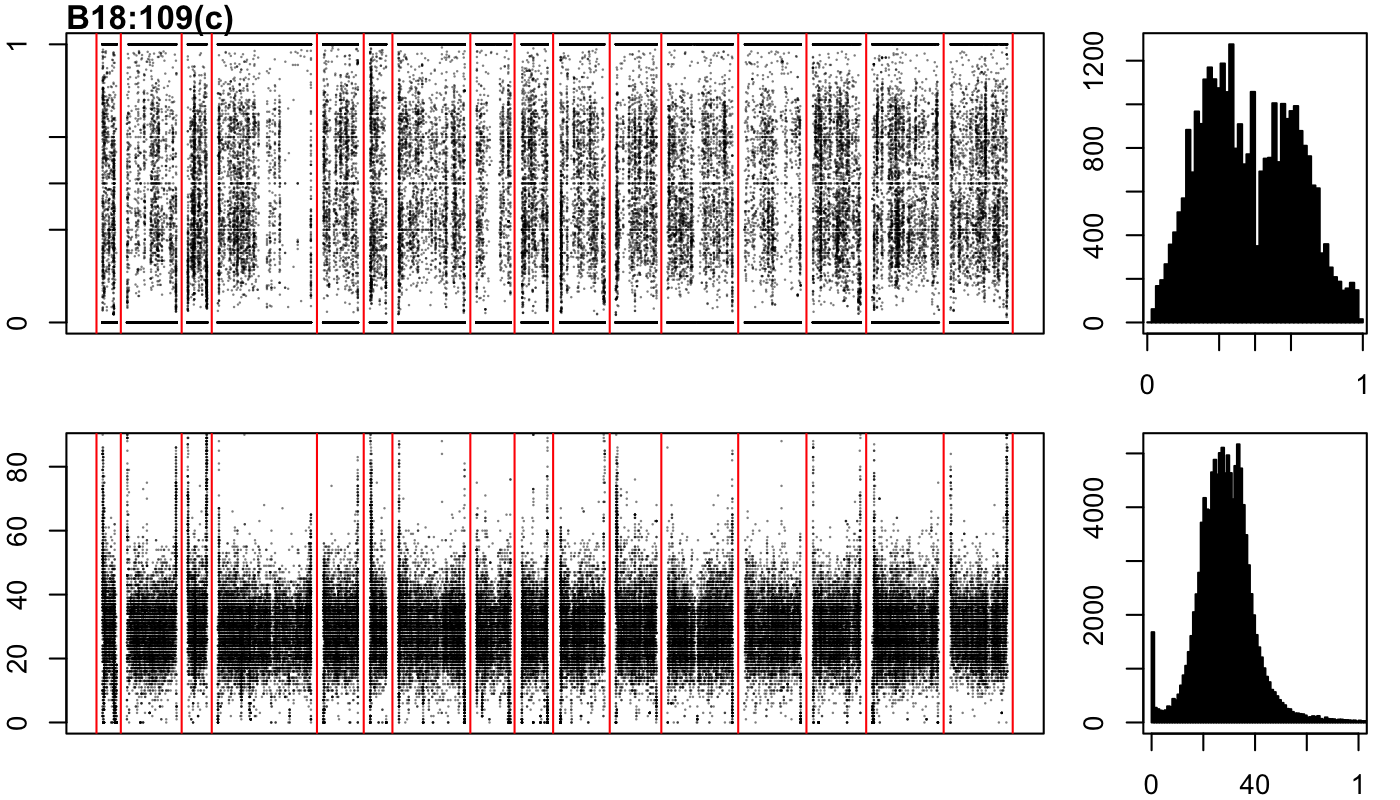

### B18-197-a.png

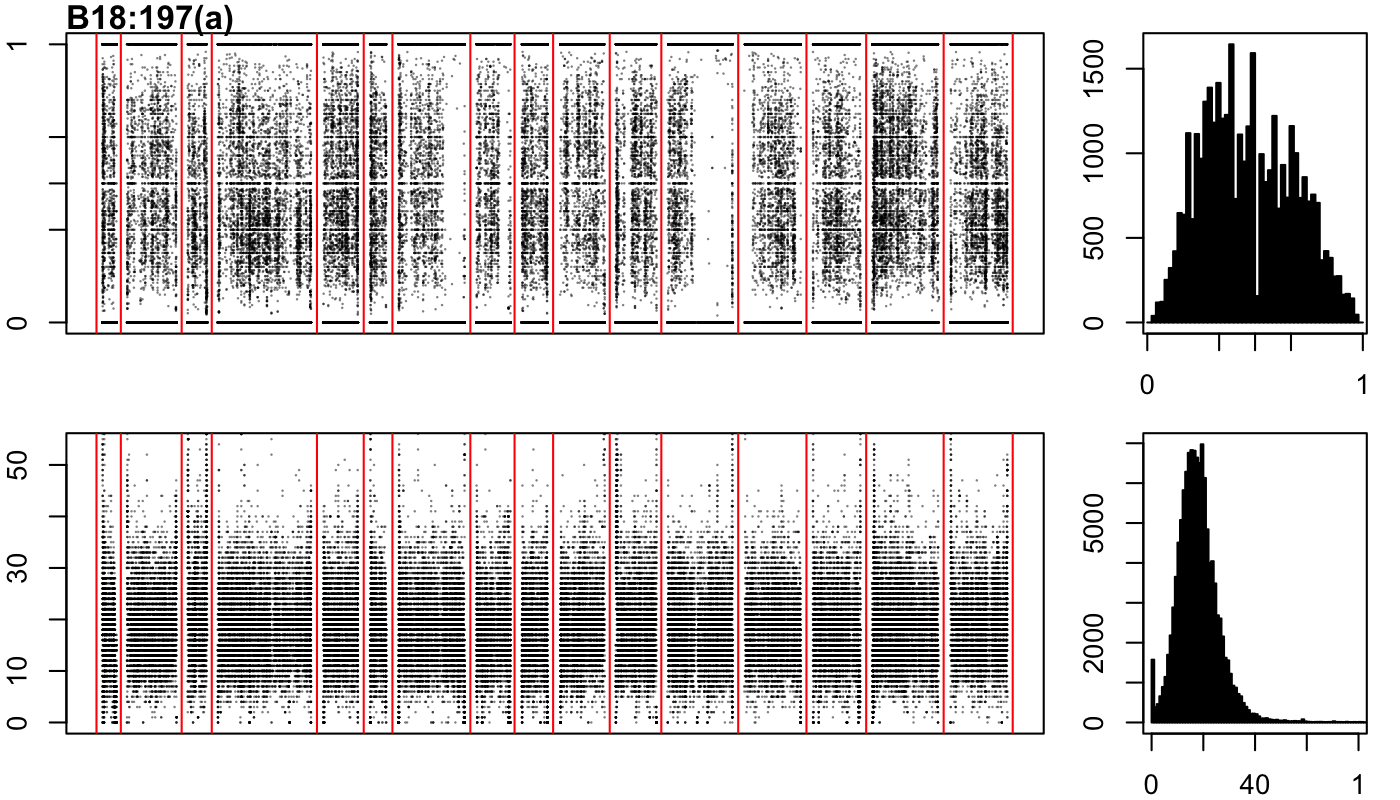

### B18-197-b.png

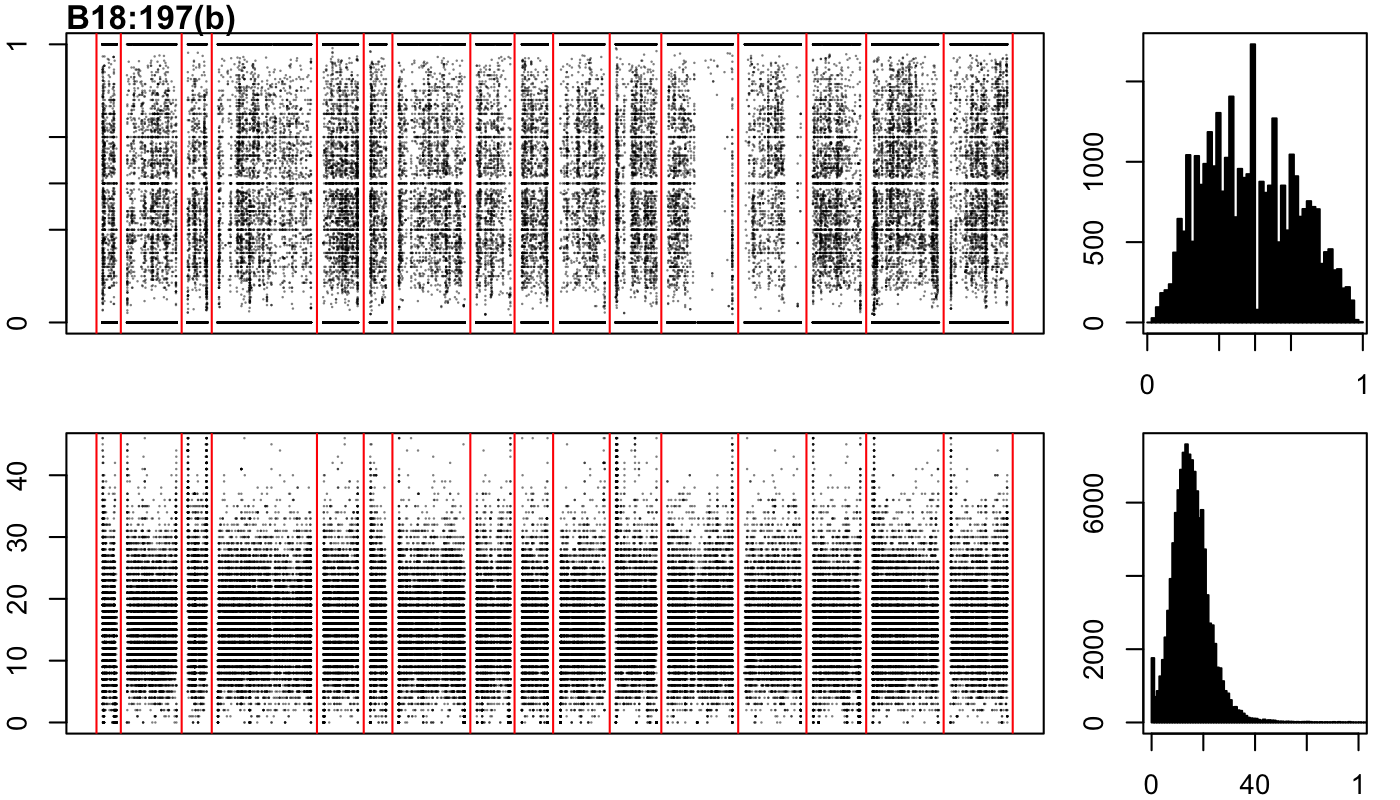

### B18-197-c.png

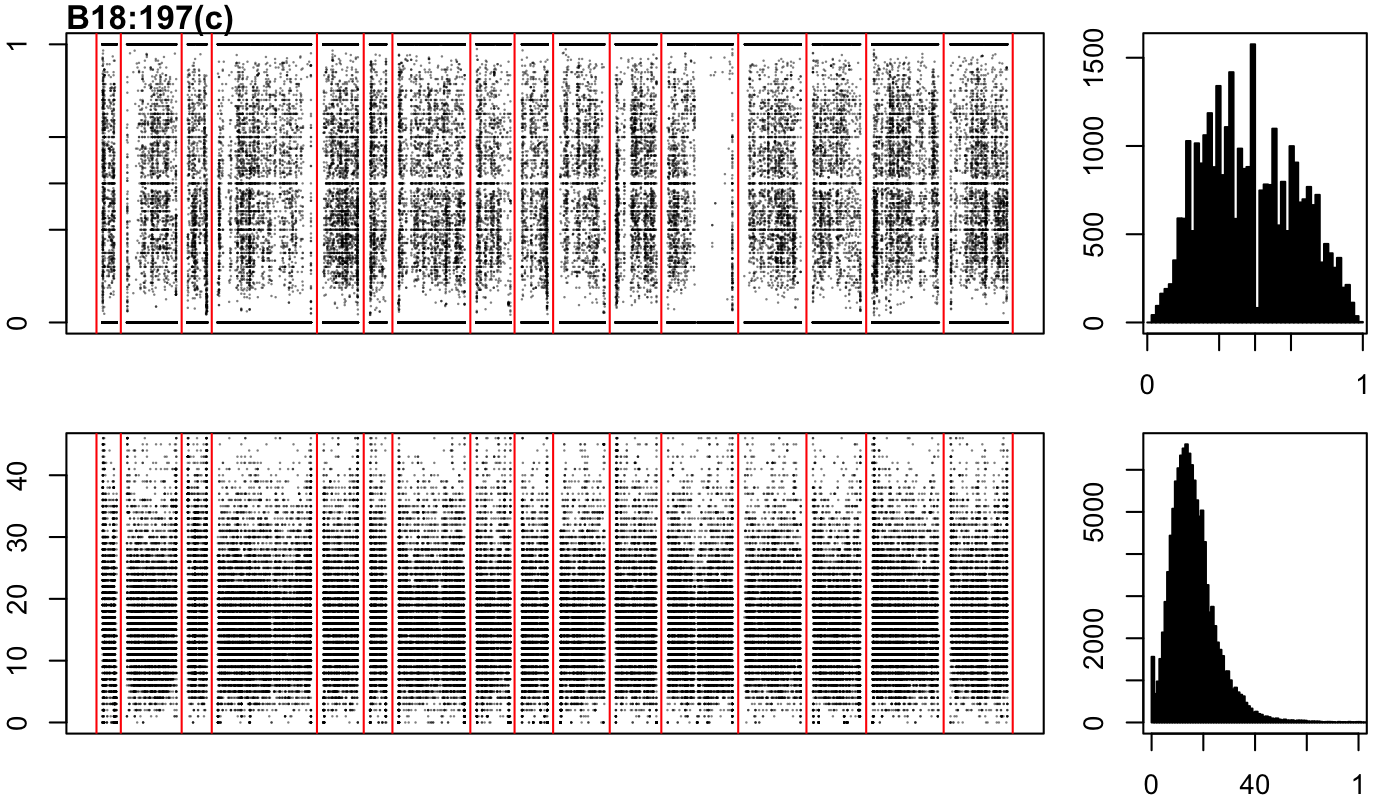

### B19-1-a.png

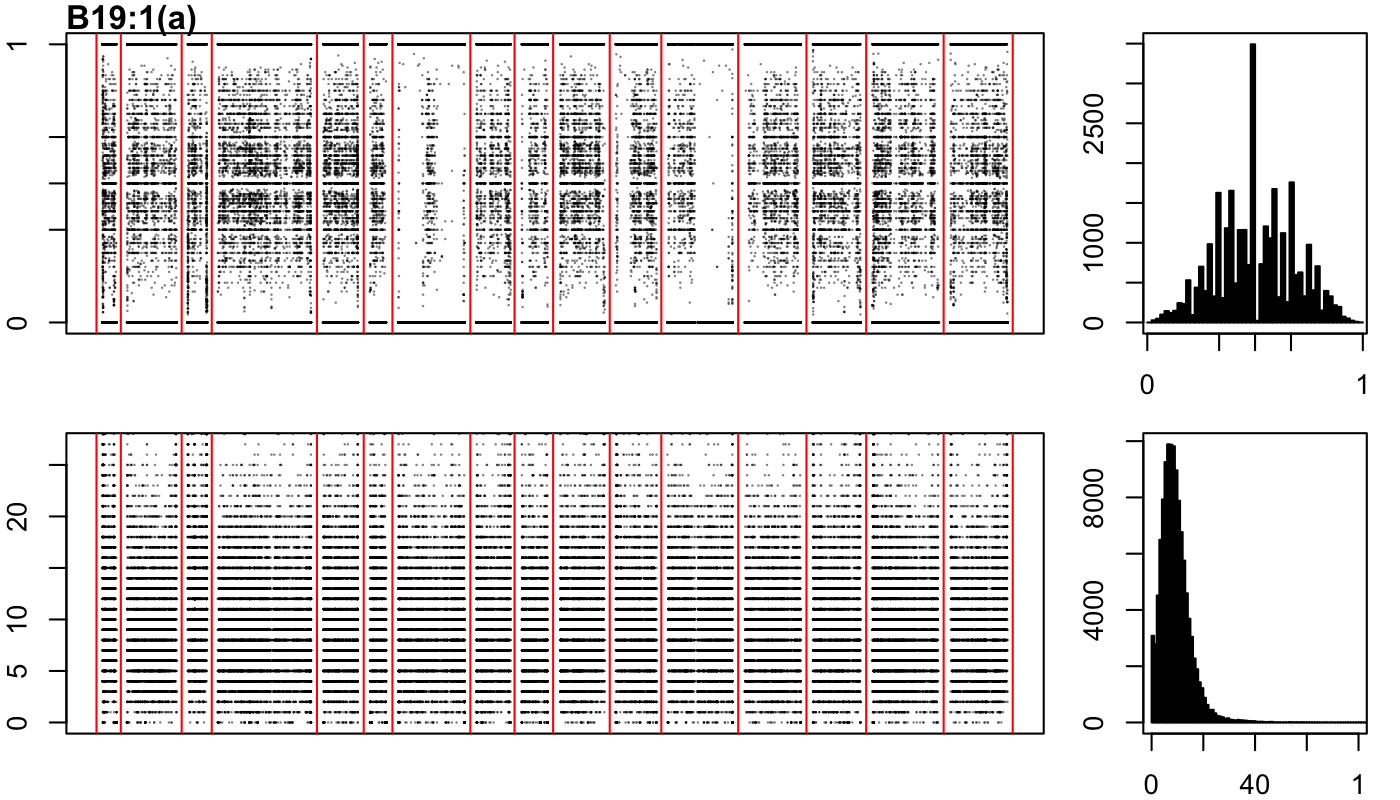

### B19-1-b.png

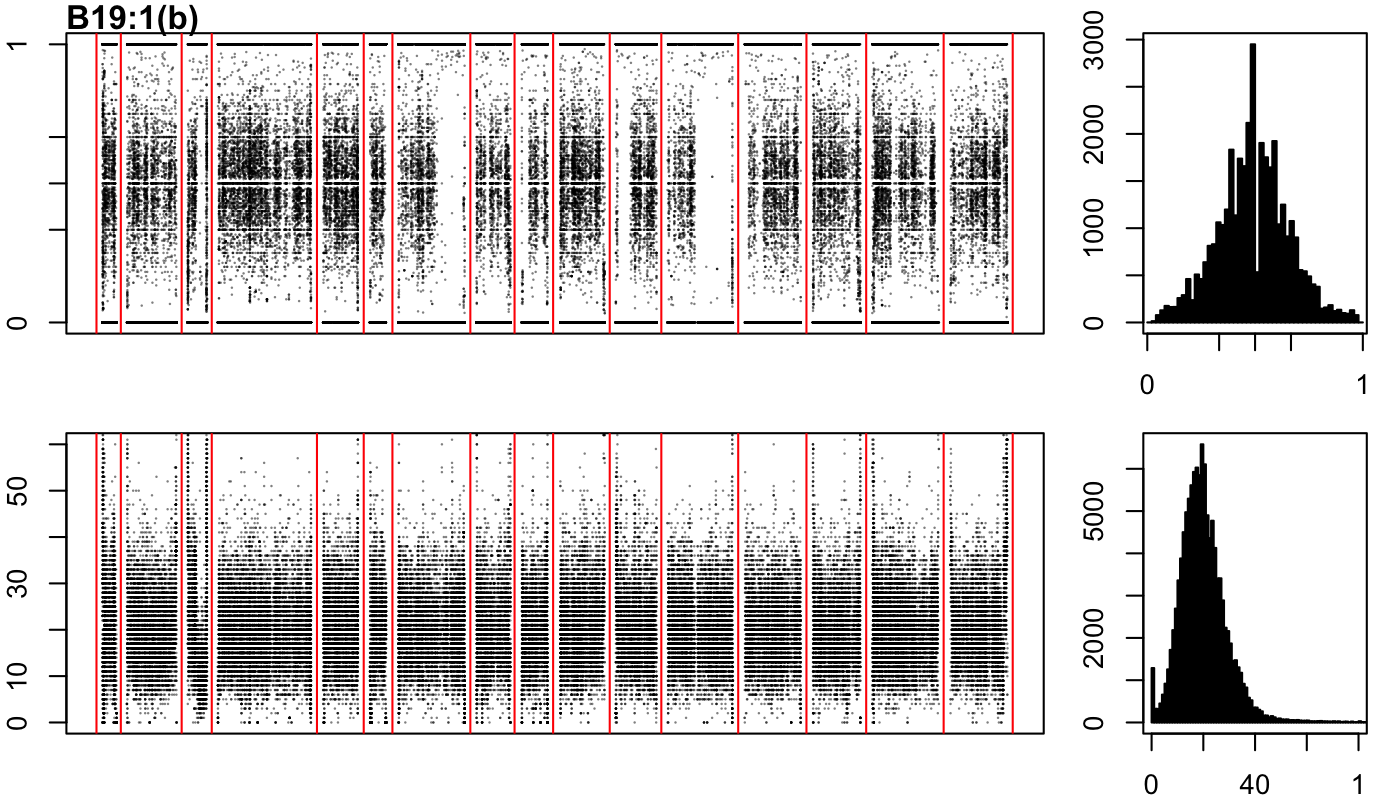

### B19-1-c.png

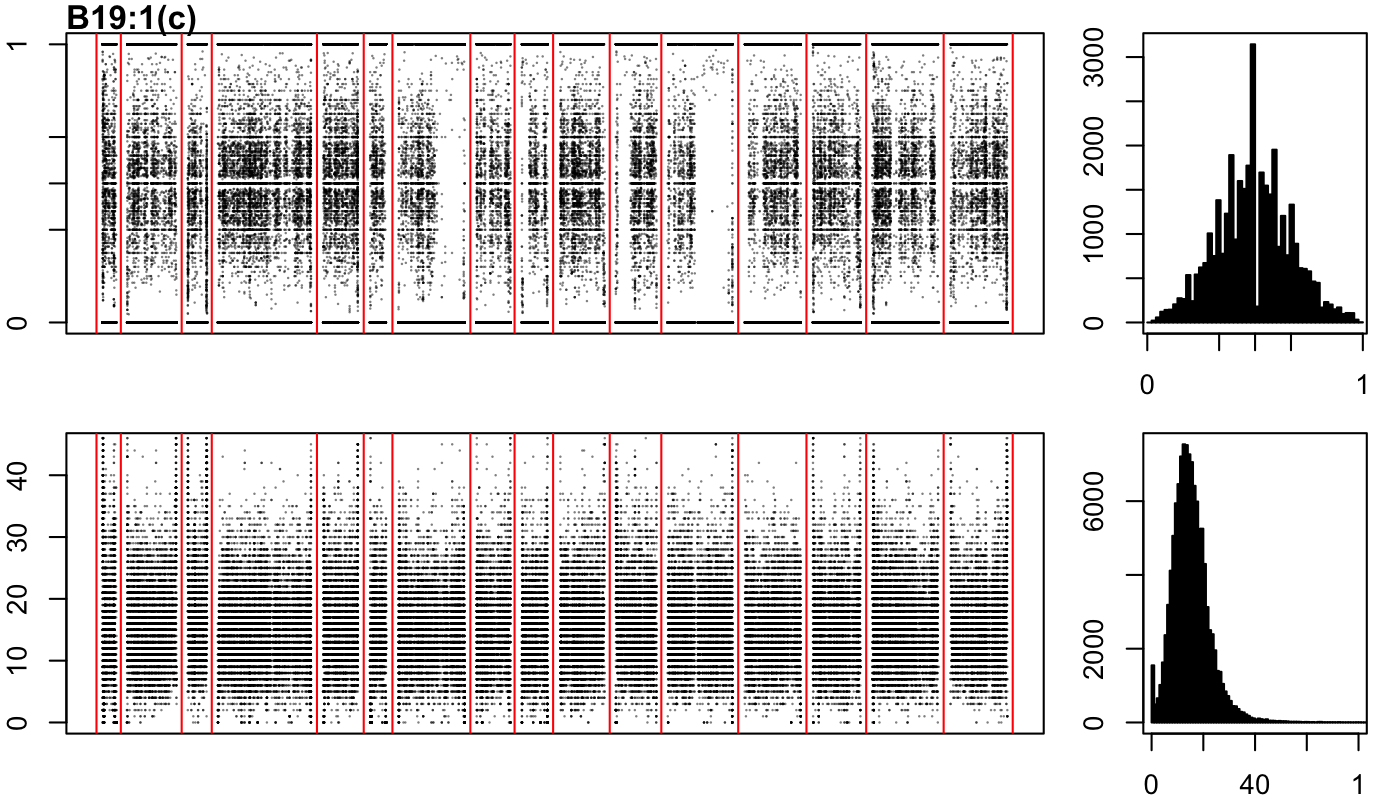

### B19-5-a.png

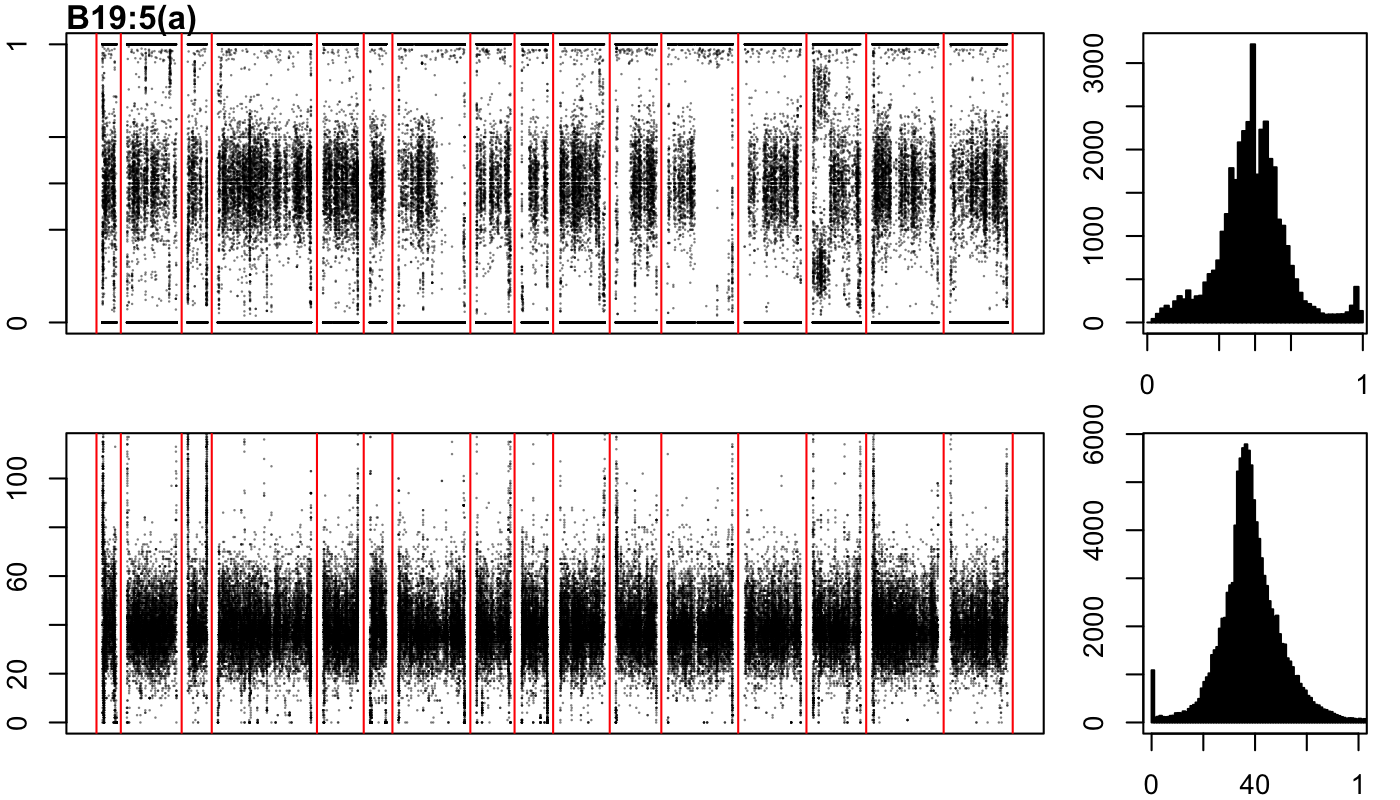

### B19-5-b.png

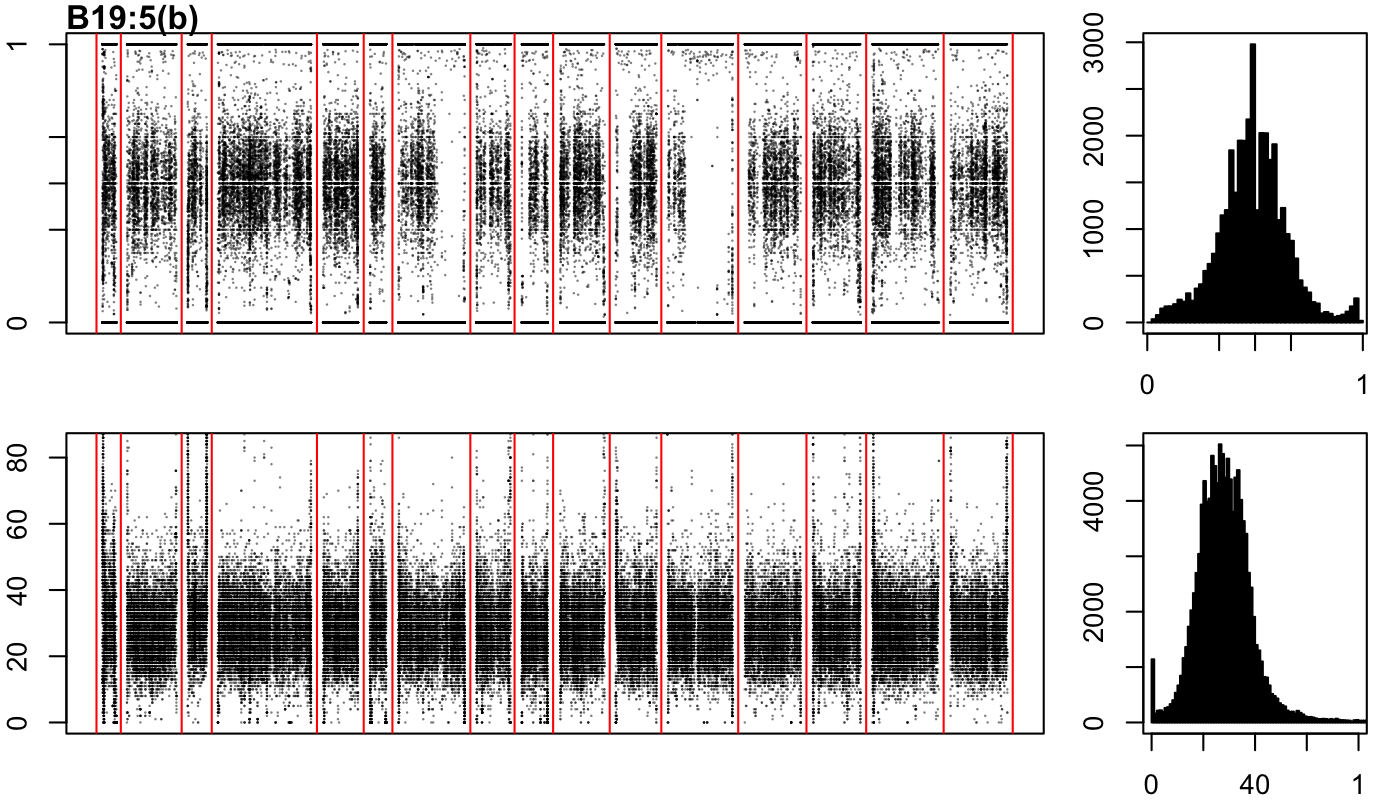

### B19-5-c.png

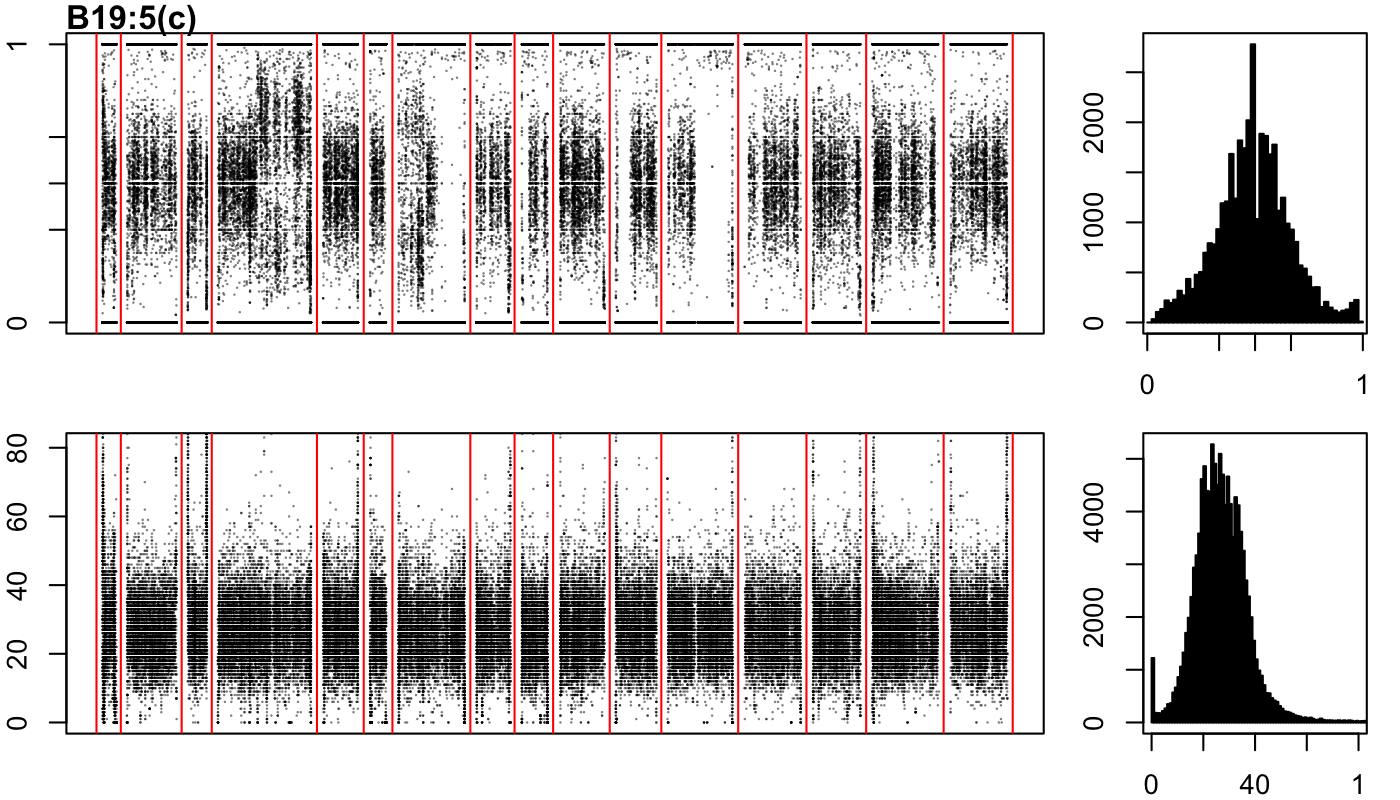

### B19-67-a.png

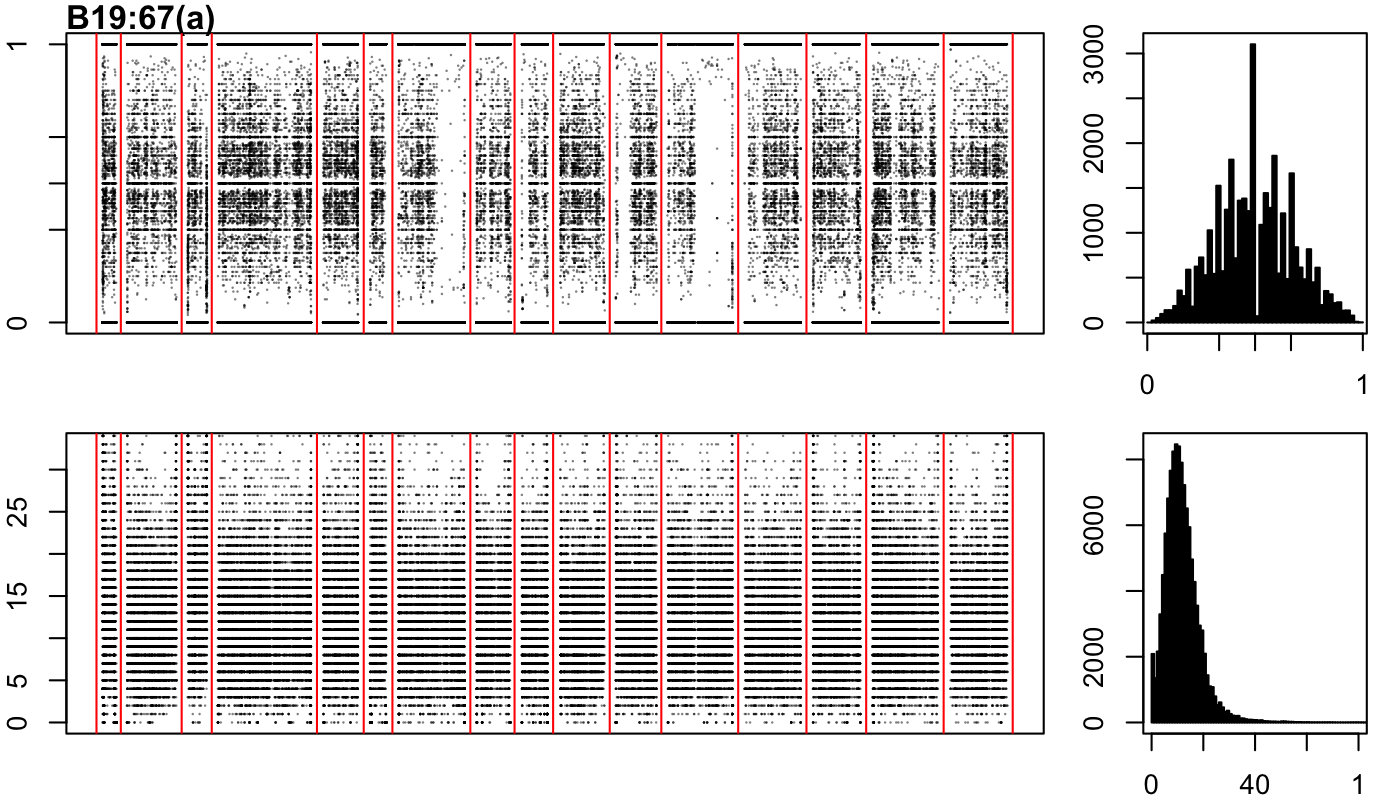

### B19-67-b.png

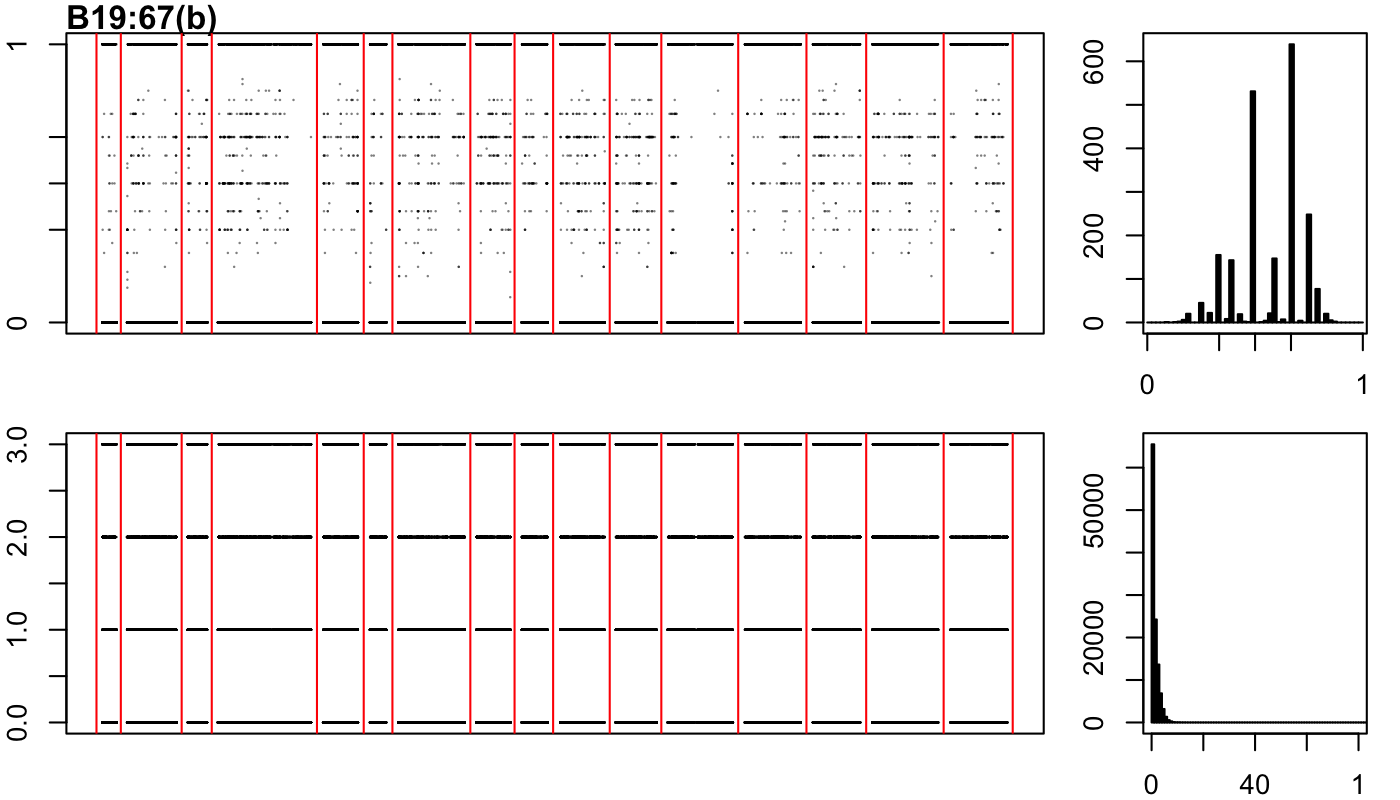

### B19-81-a.png

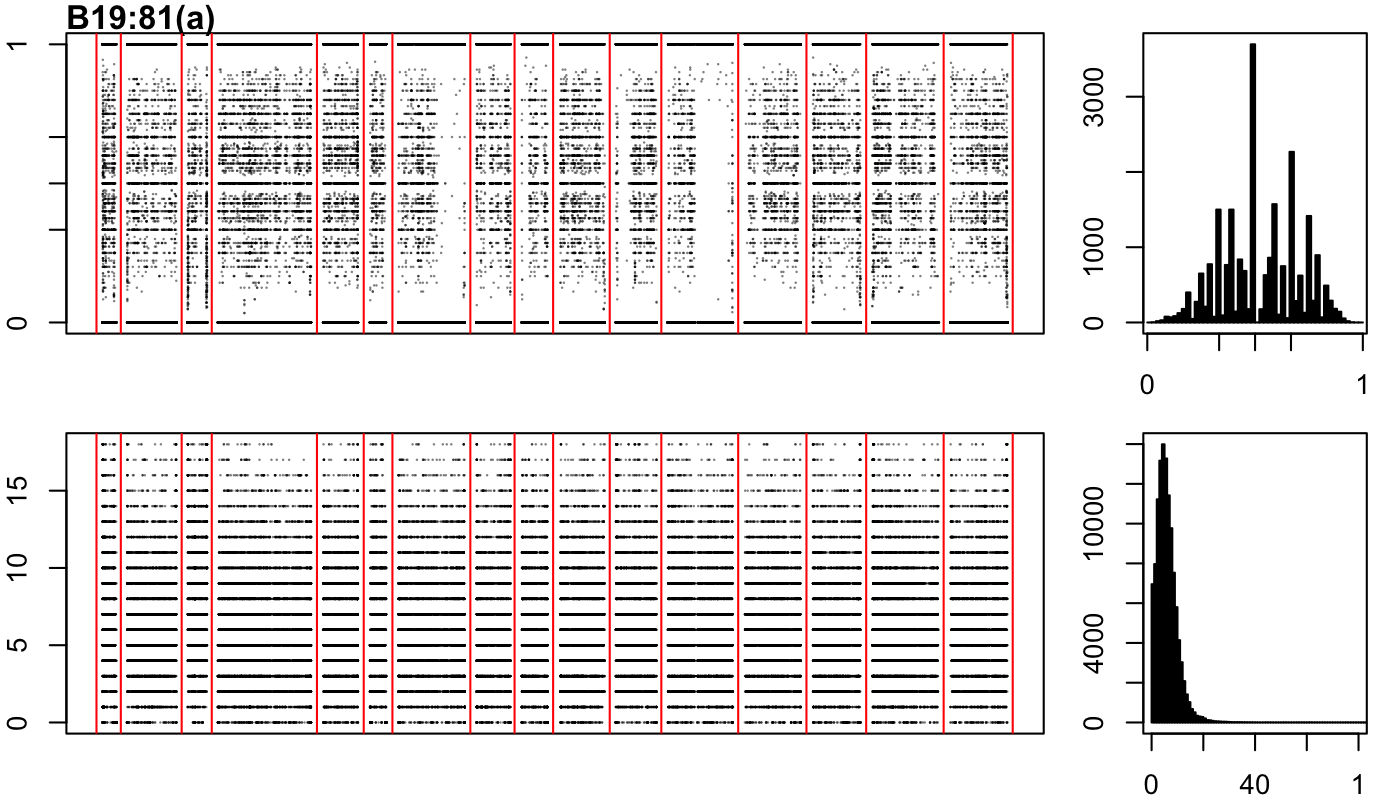

### B19-81-b.png

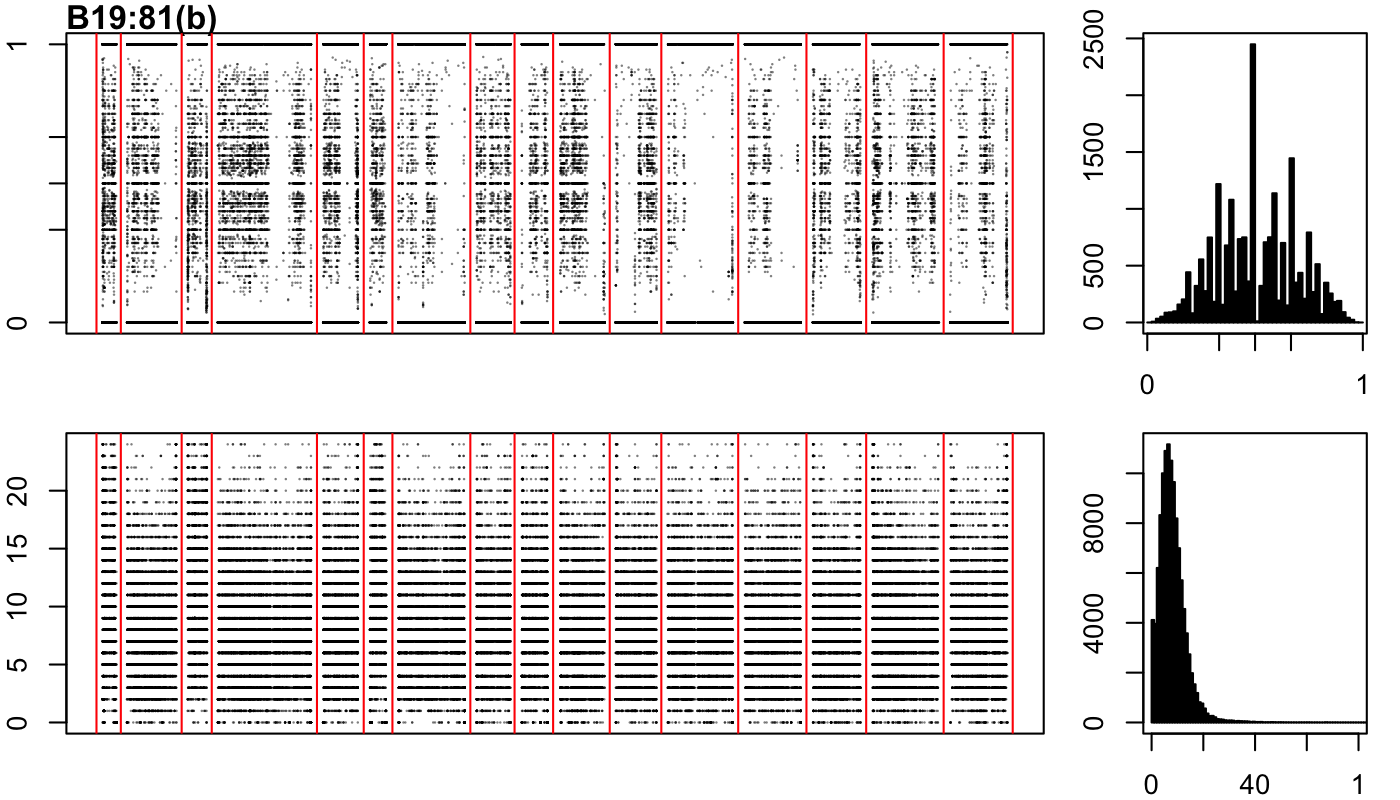

### B19-137-a.png

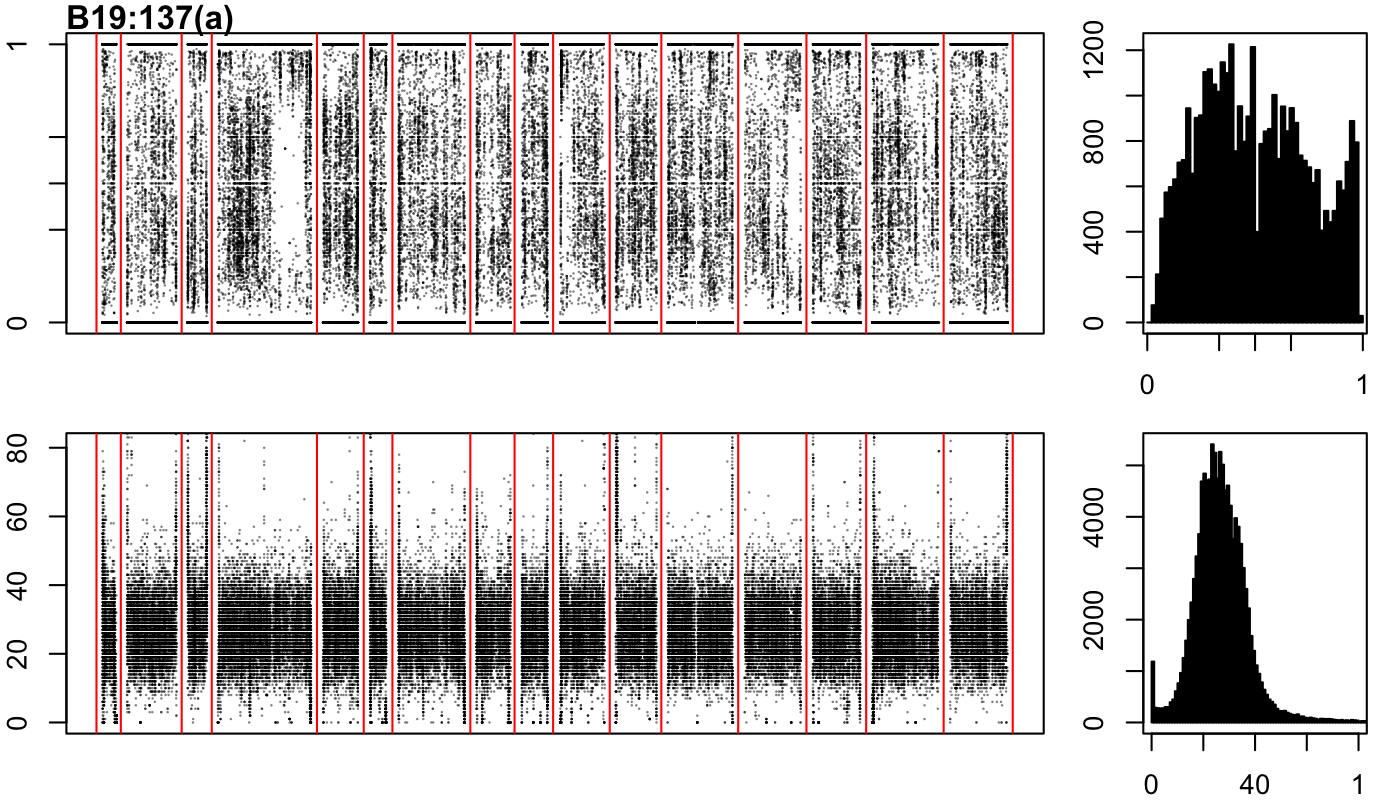

### B19-137-b.png

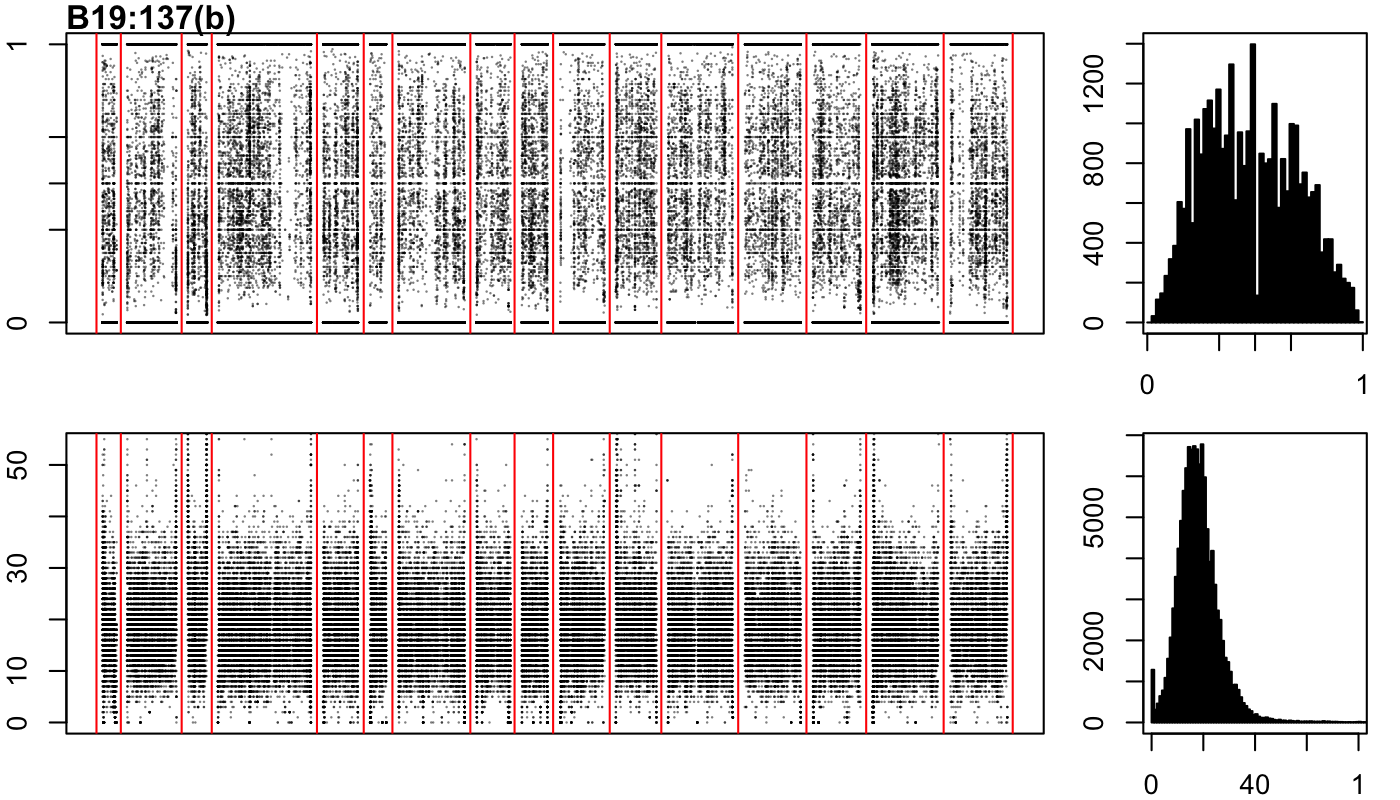

### B19-151-b.png

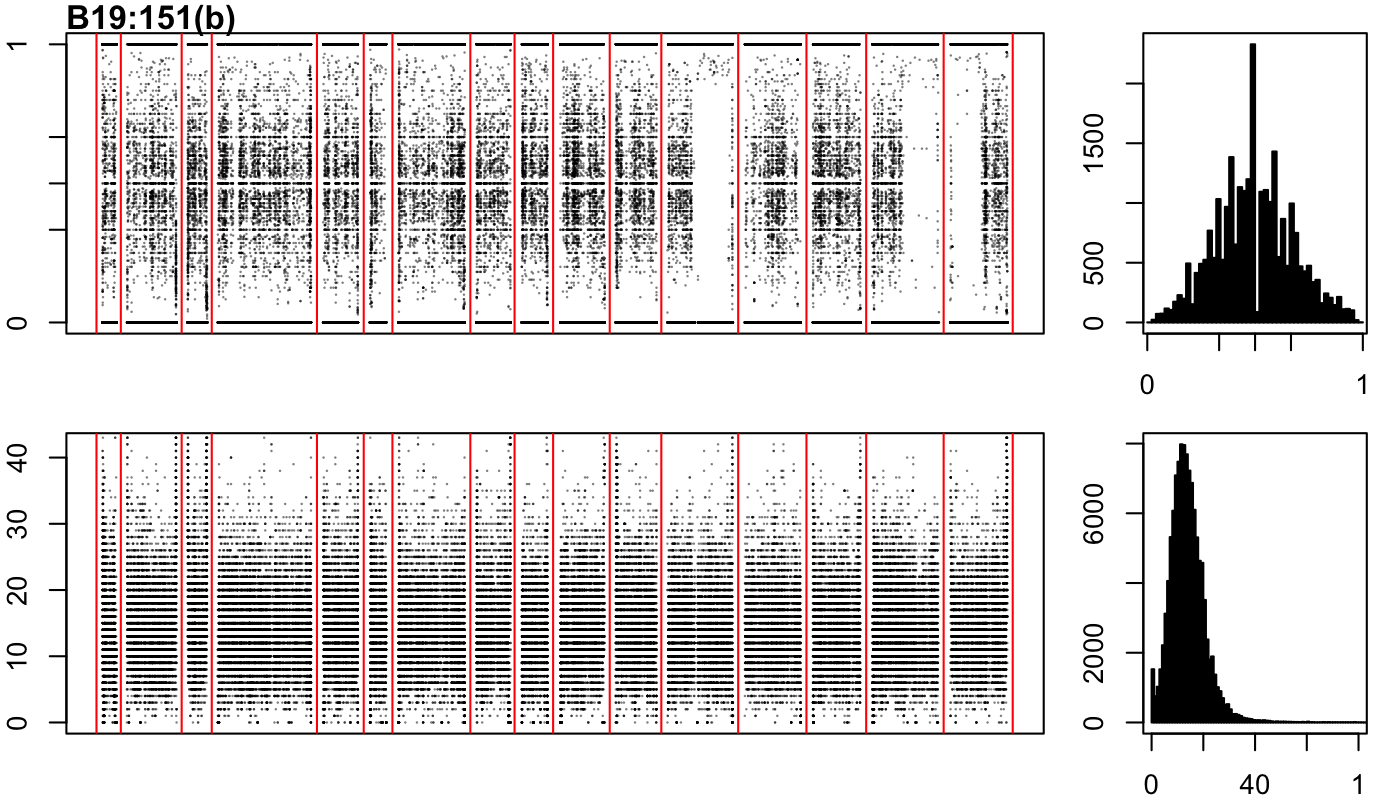

### B19-172-a.png

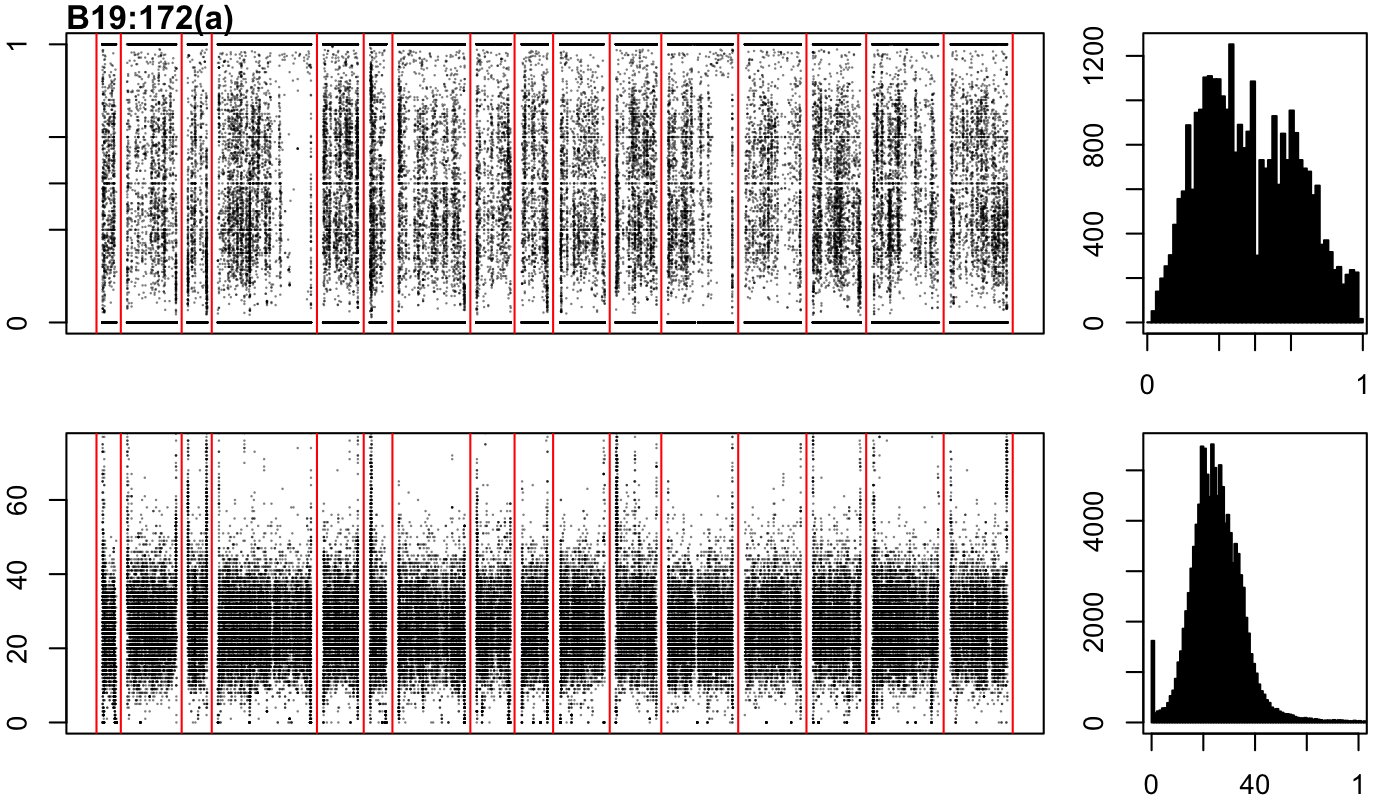

### B19-172-b.png

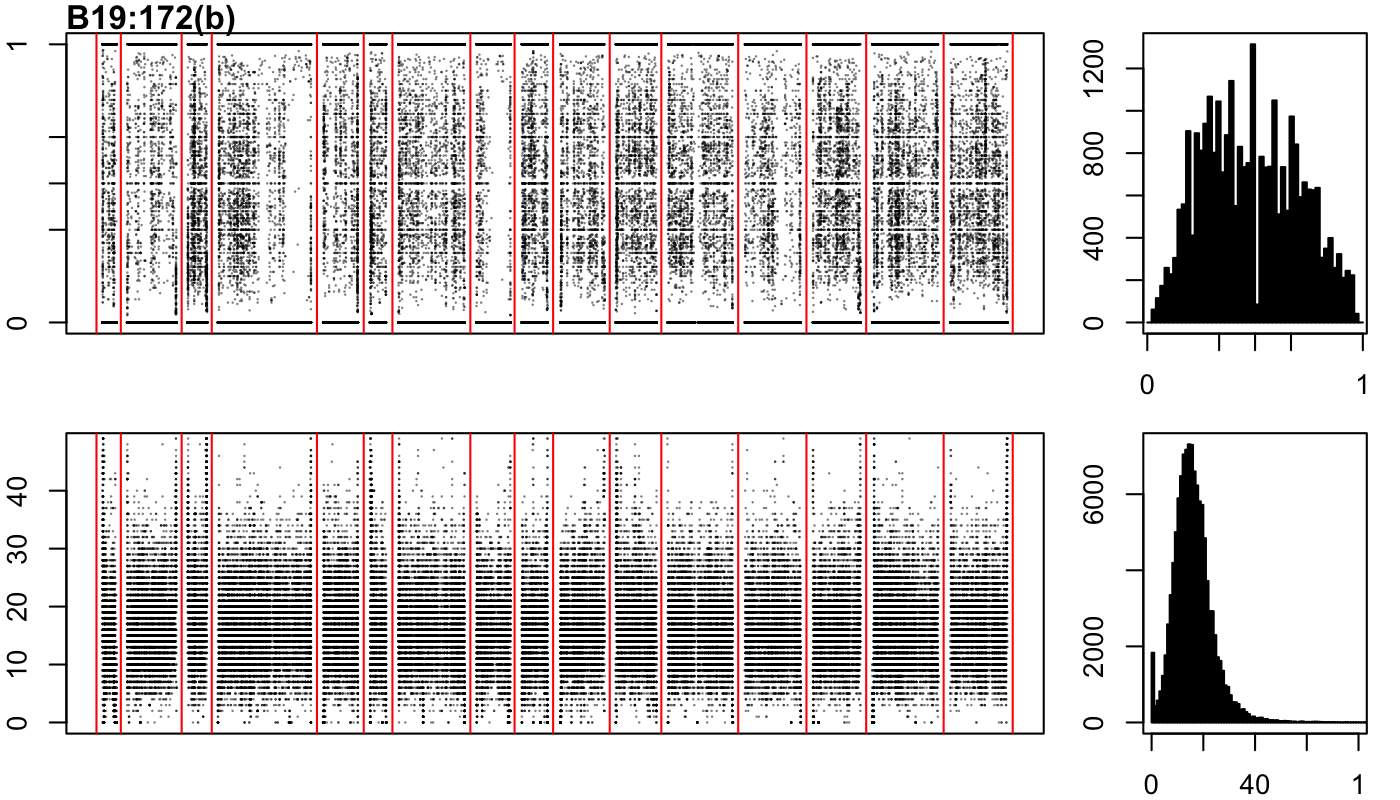

### UCP_2018.png

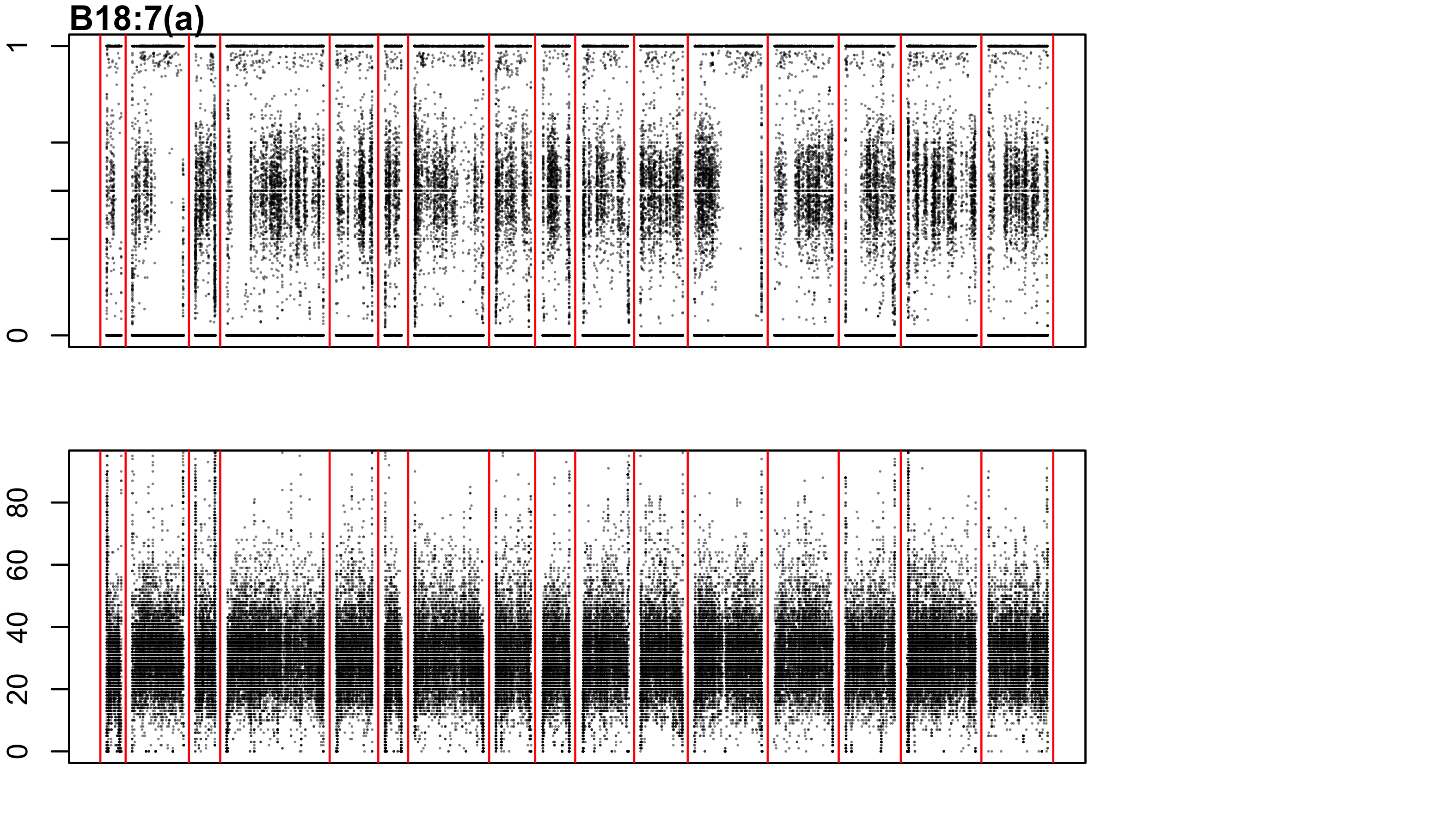
