## Supplementary figures and images for "Yeast population dynamics in Brazilian bioethanol production"

### A18-1-a.png

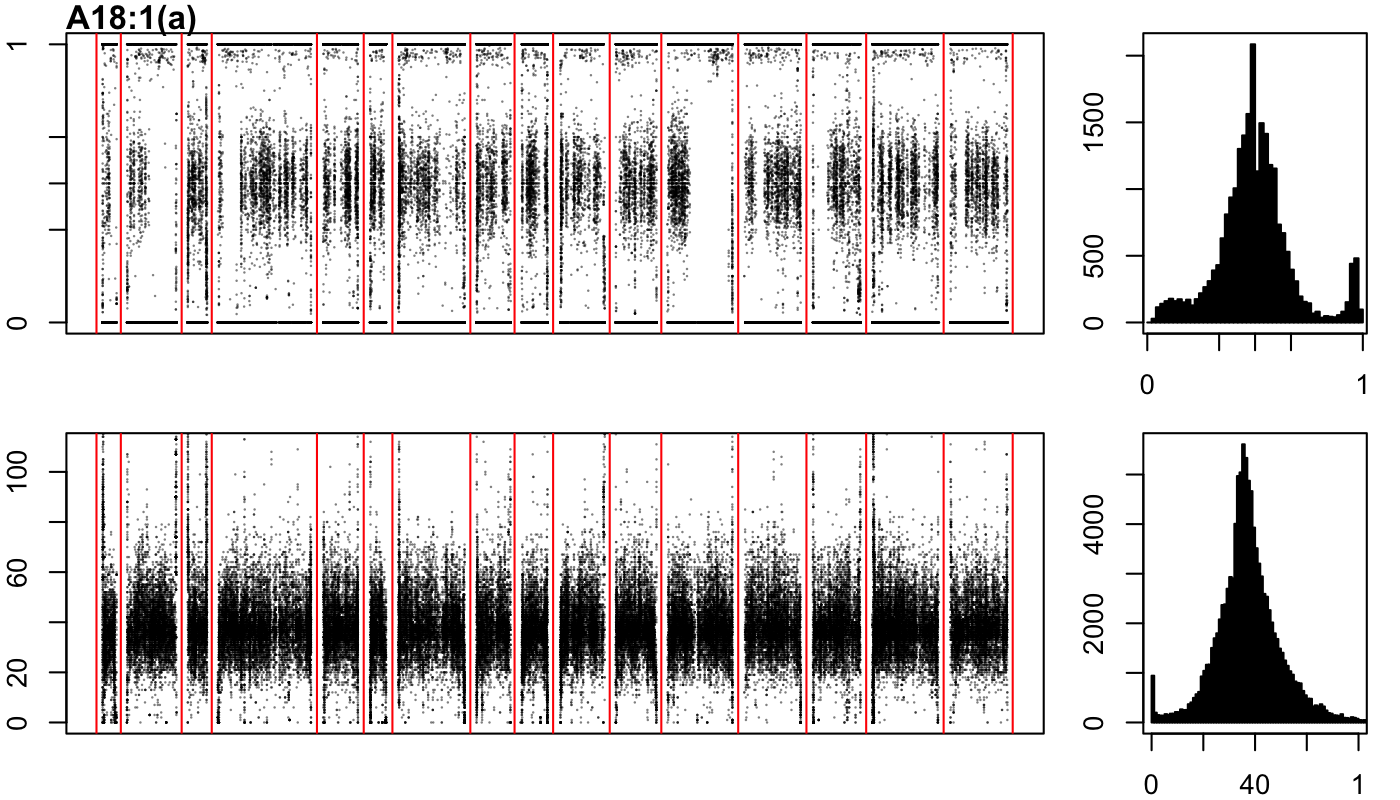

### A18-1-b.png

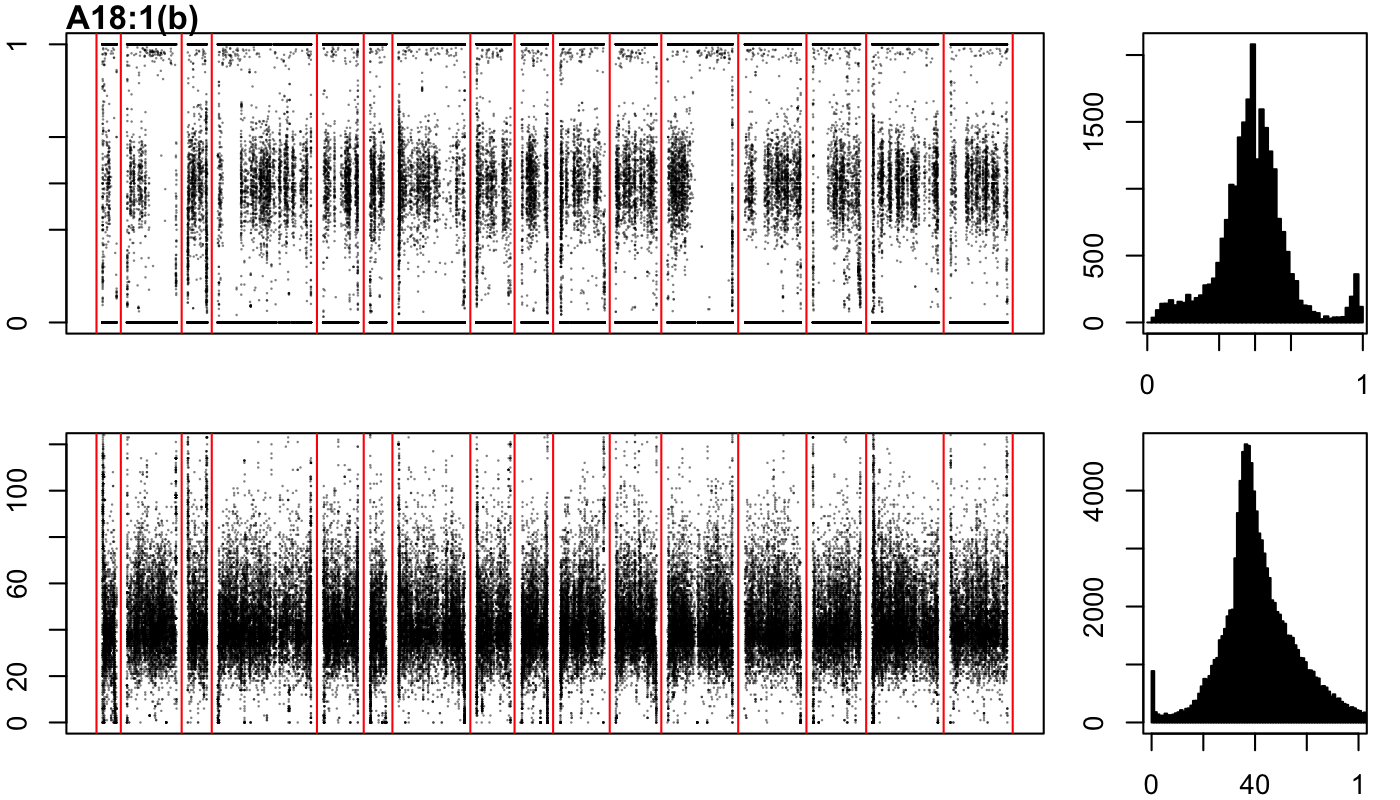

### A18-1-c.png

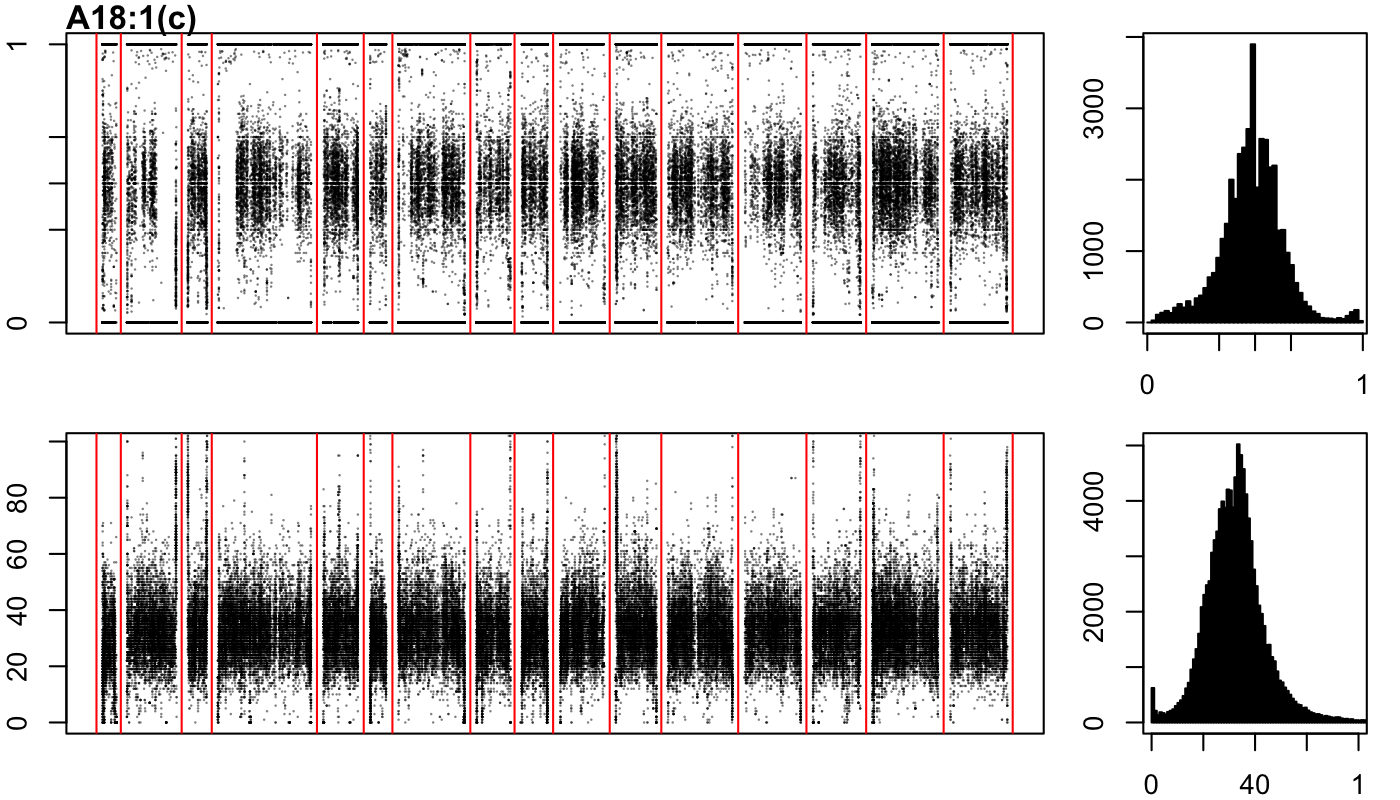

### A18-3-a.png

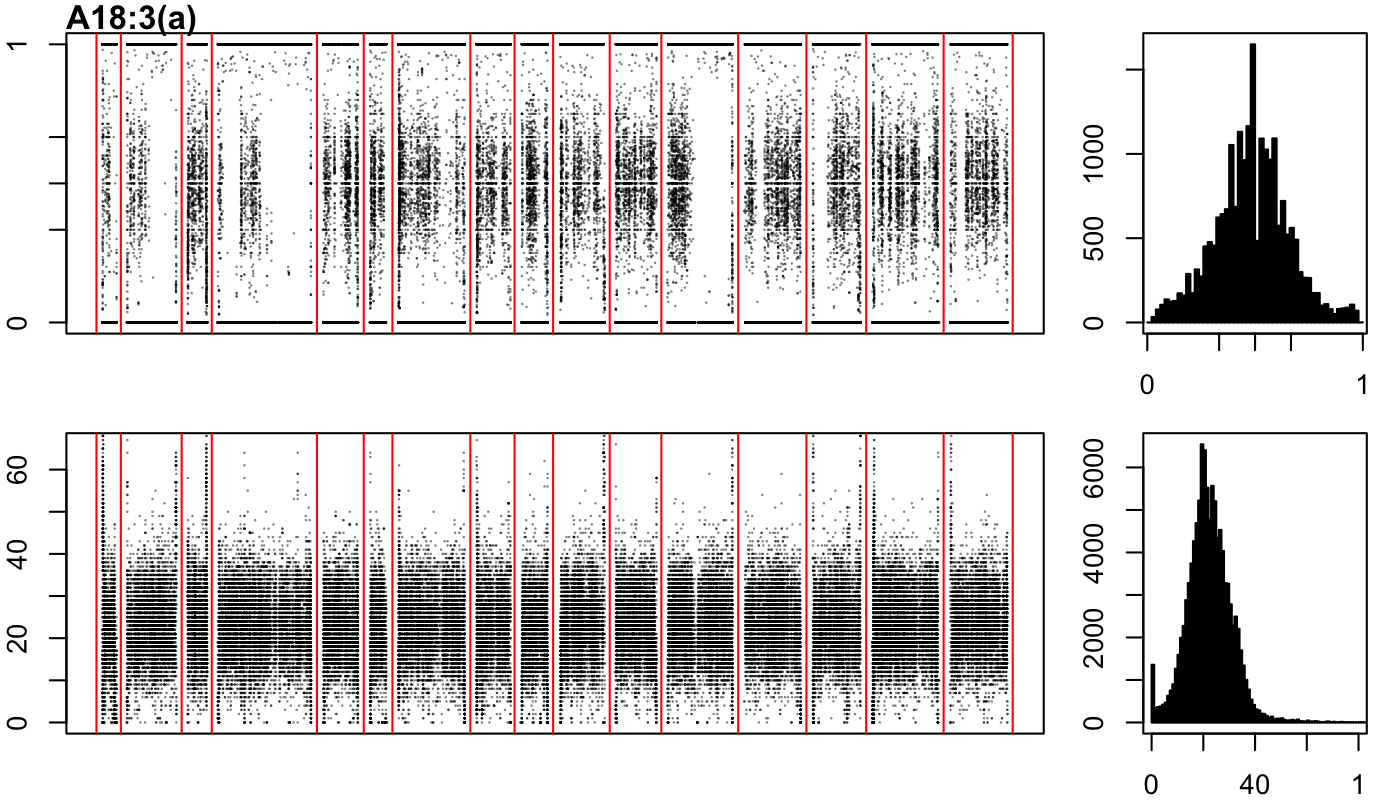

### A18-3-b.png

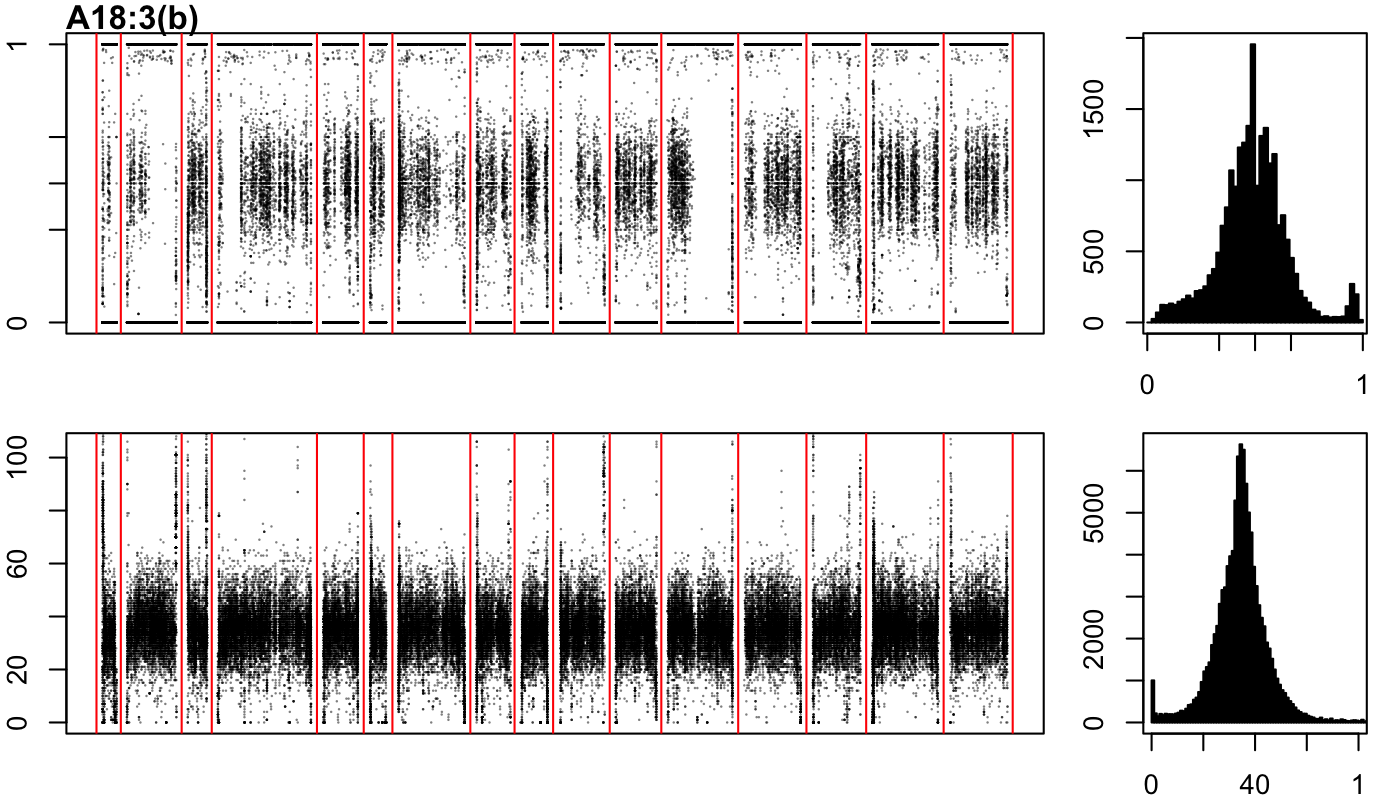

### A18-3-c.png

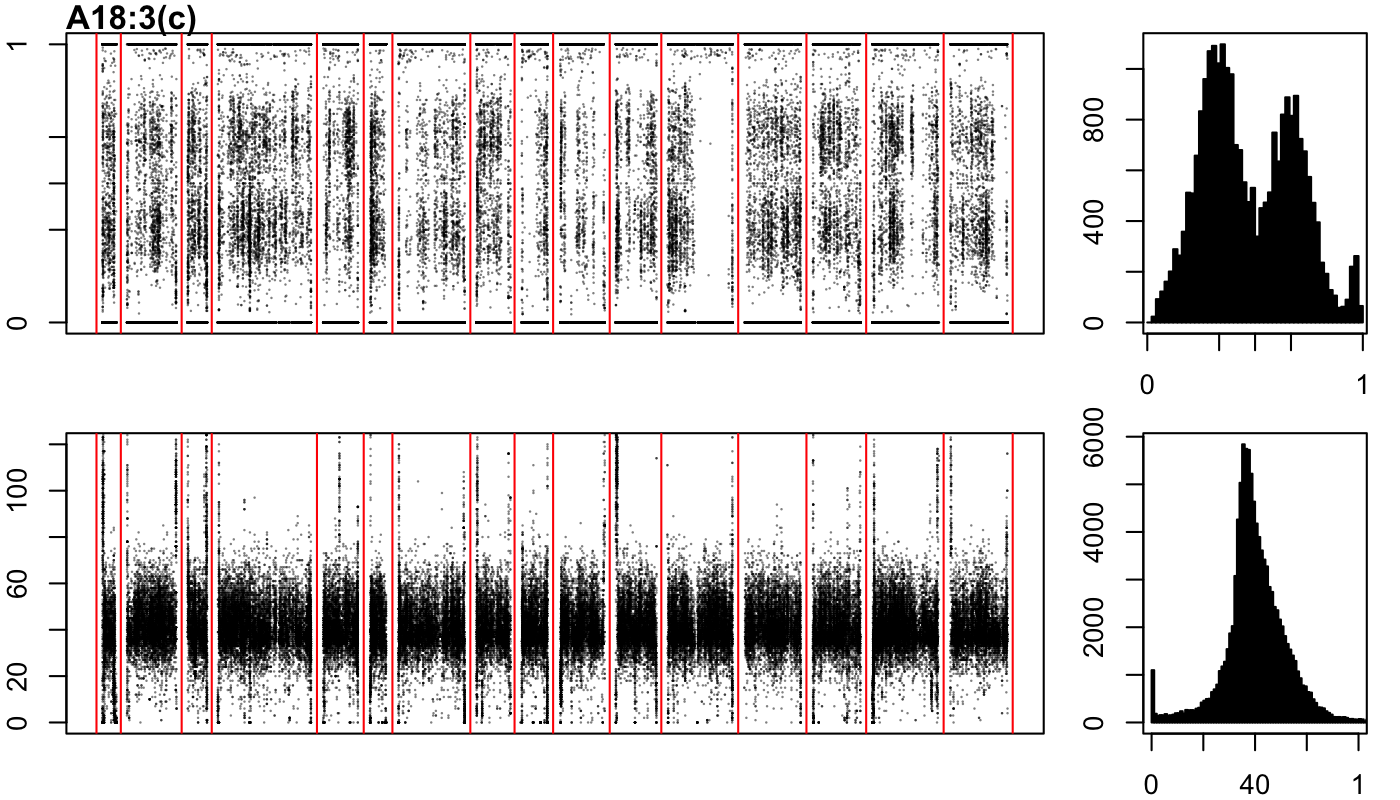

### A18-7-a.png

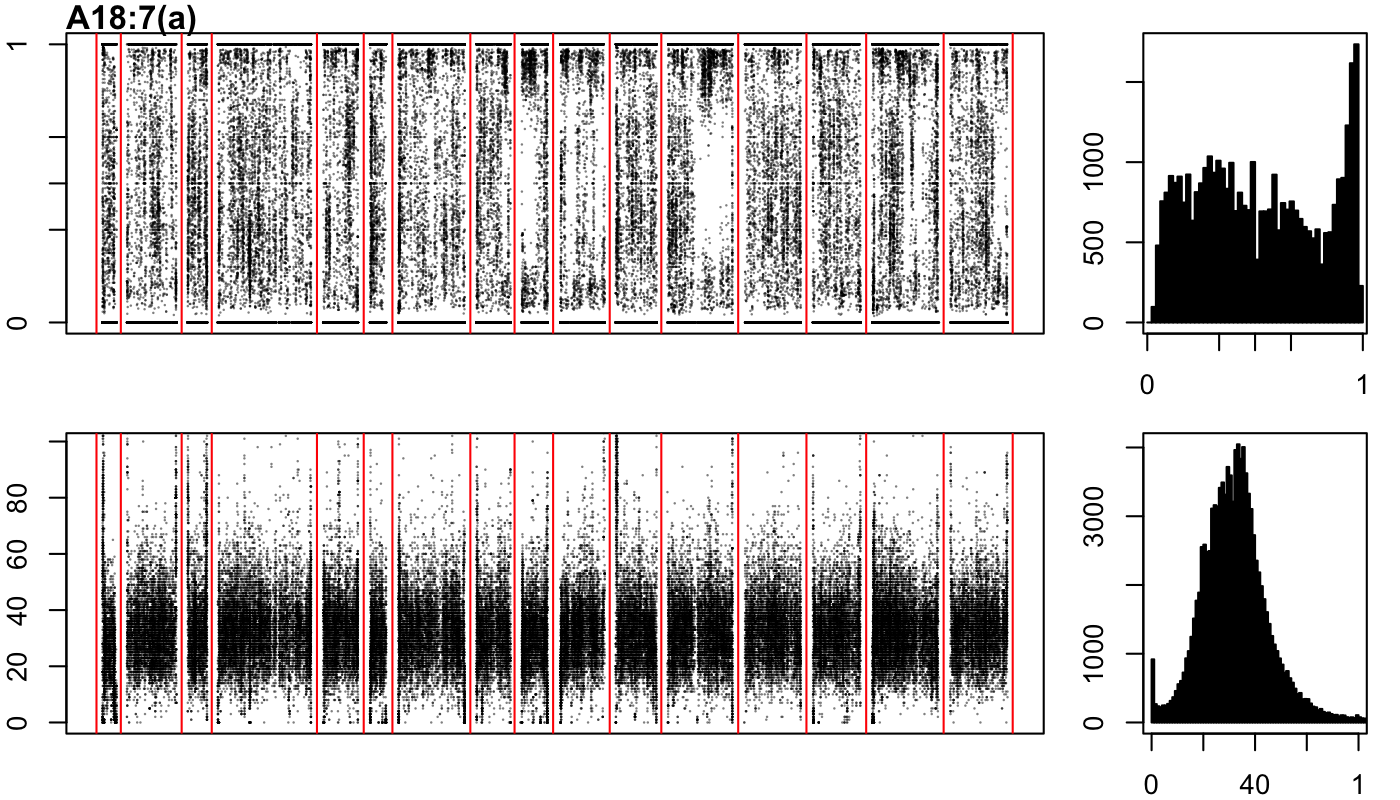

### A18-7-b.png

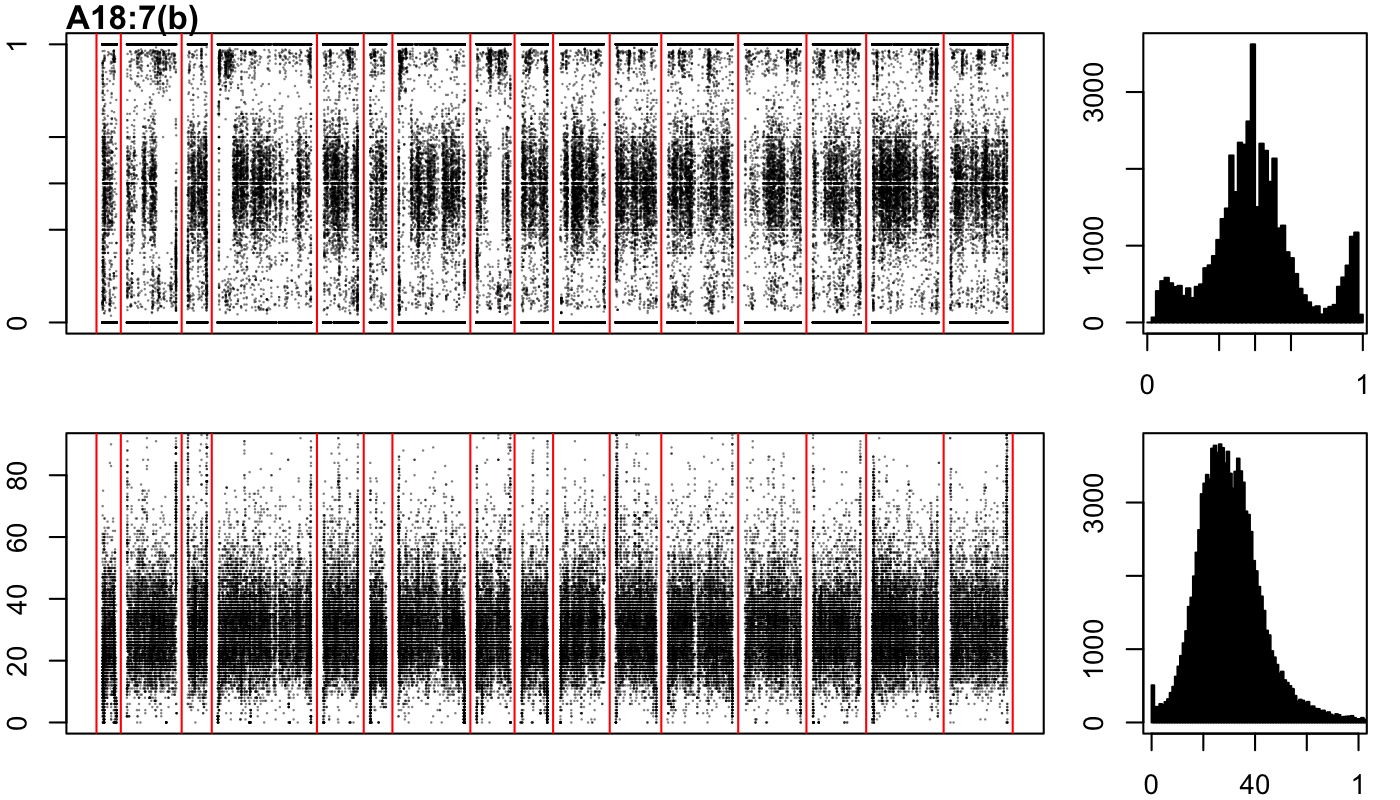

### A18-27-a.png

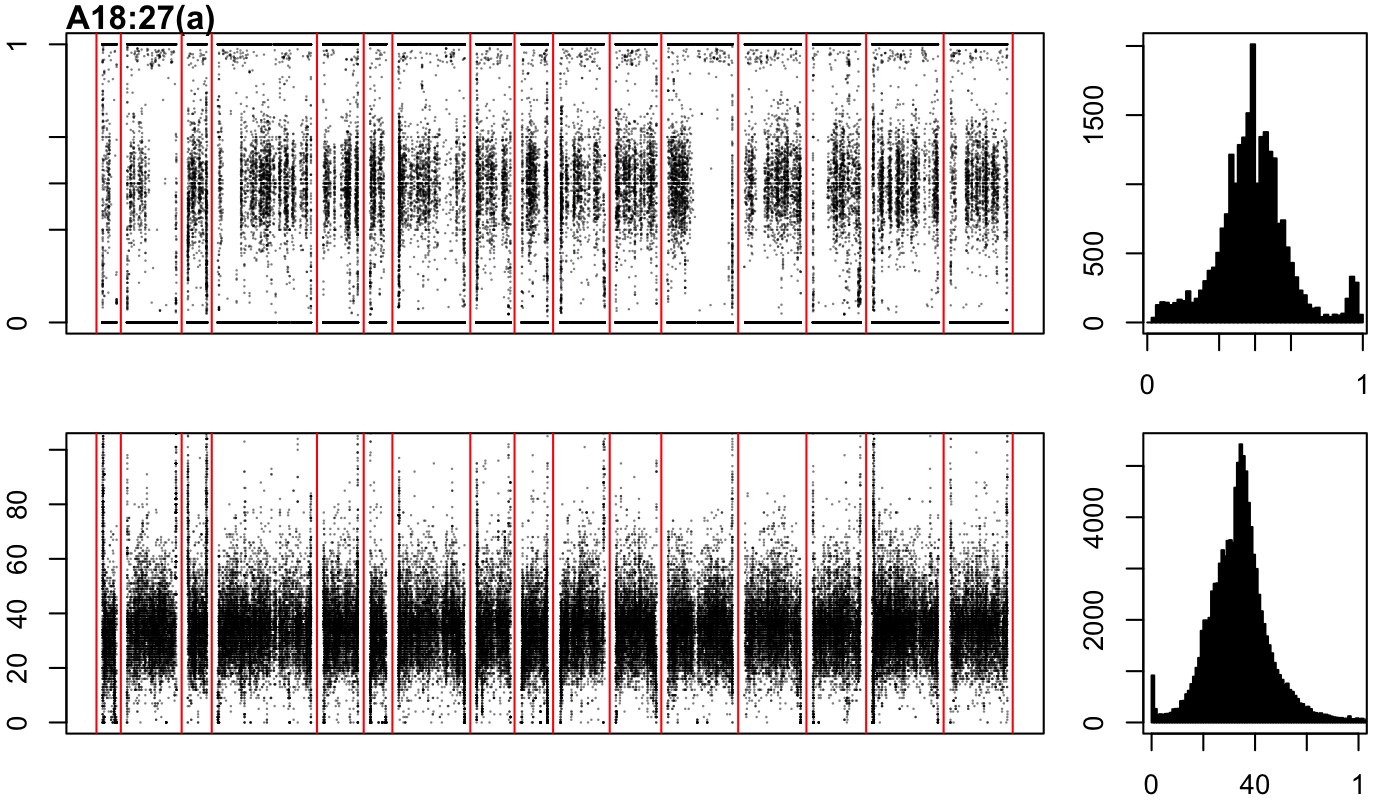

### A18-27-b.png

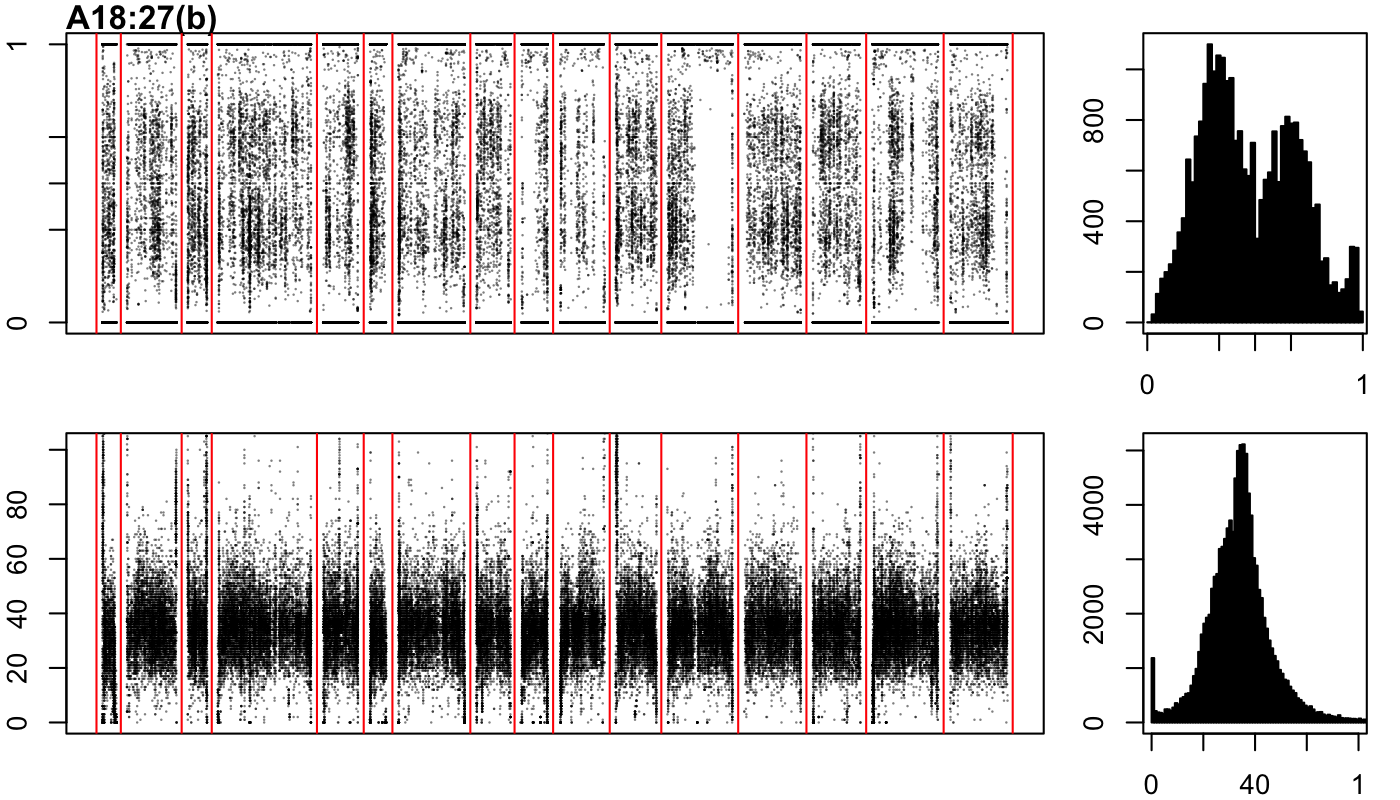

### A18-27-c.png

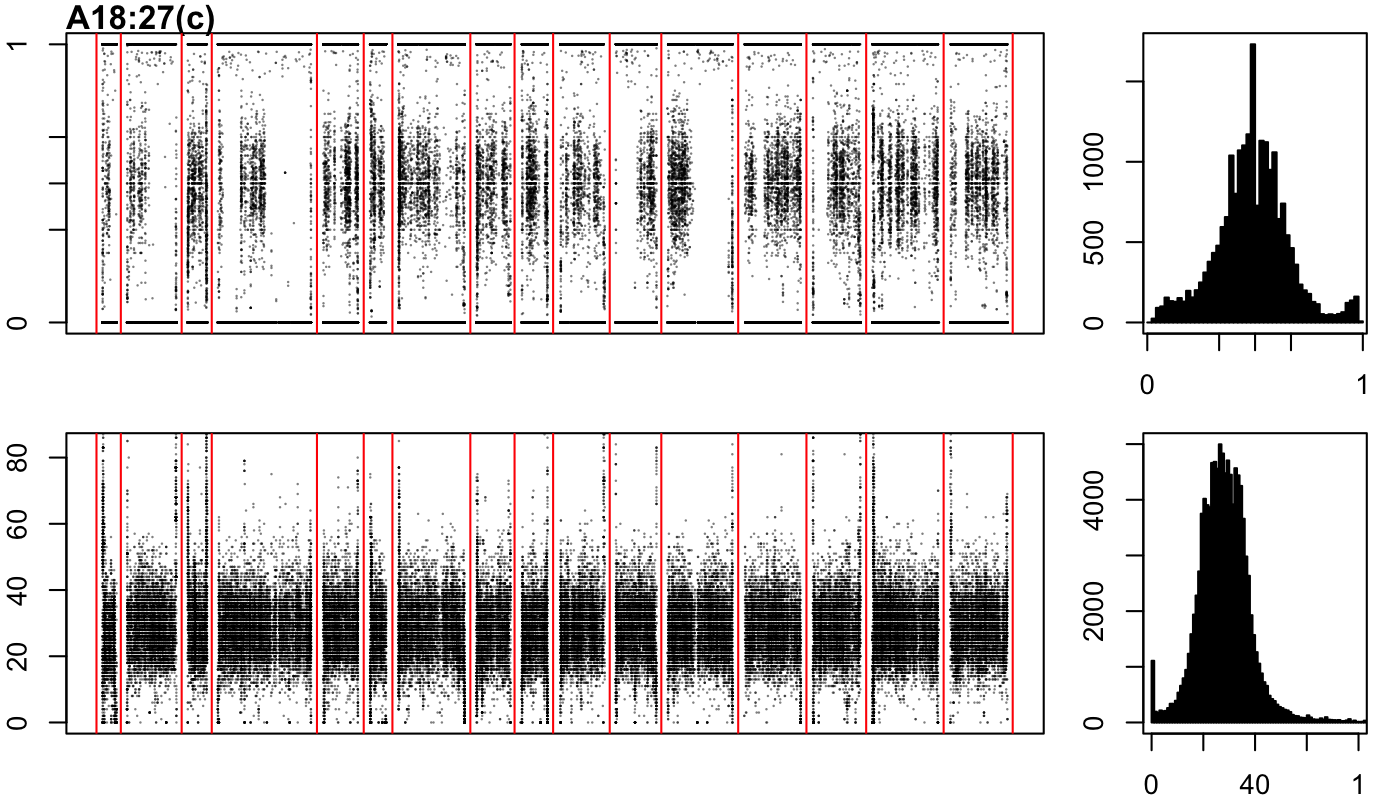

### A18-47-a.png

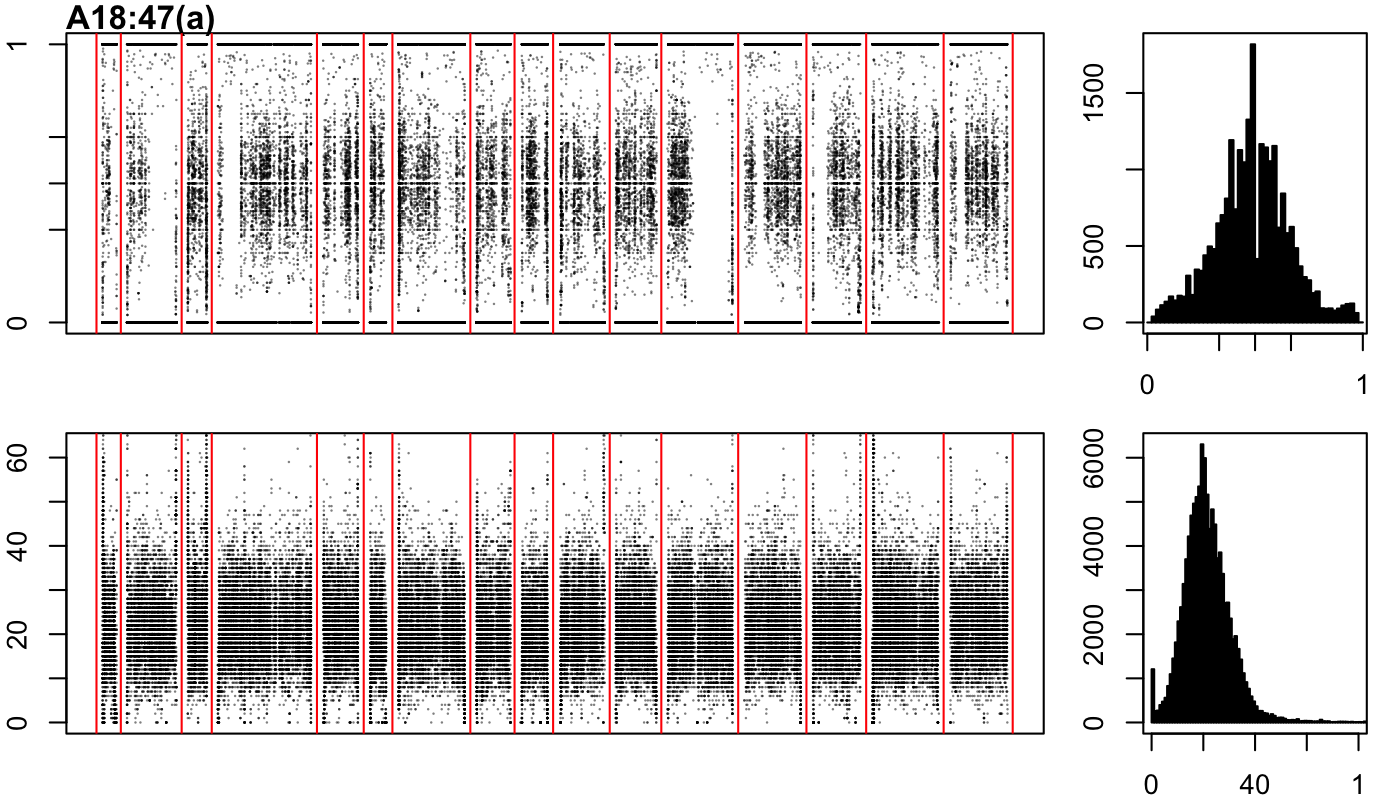

### A18-47-b.png

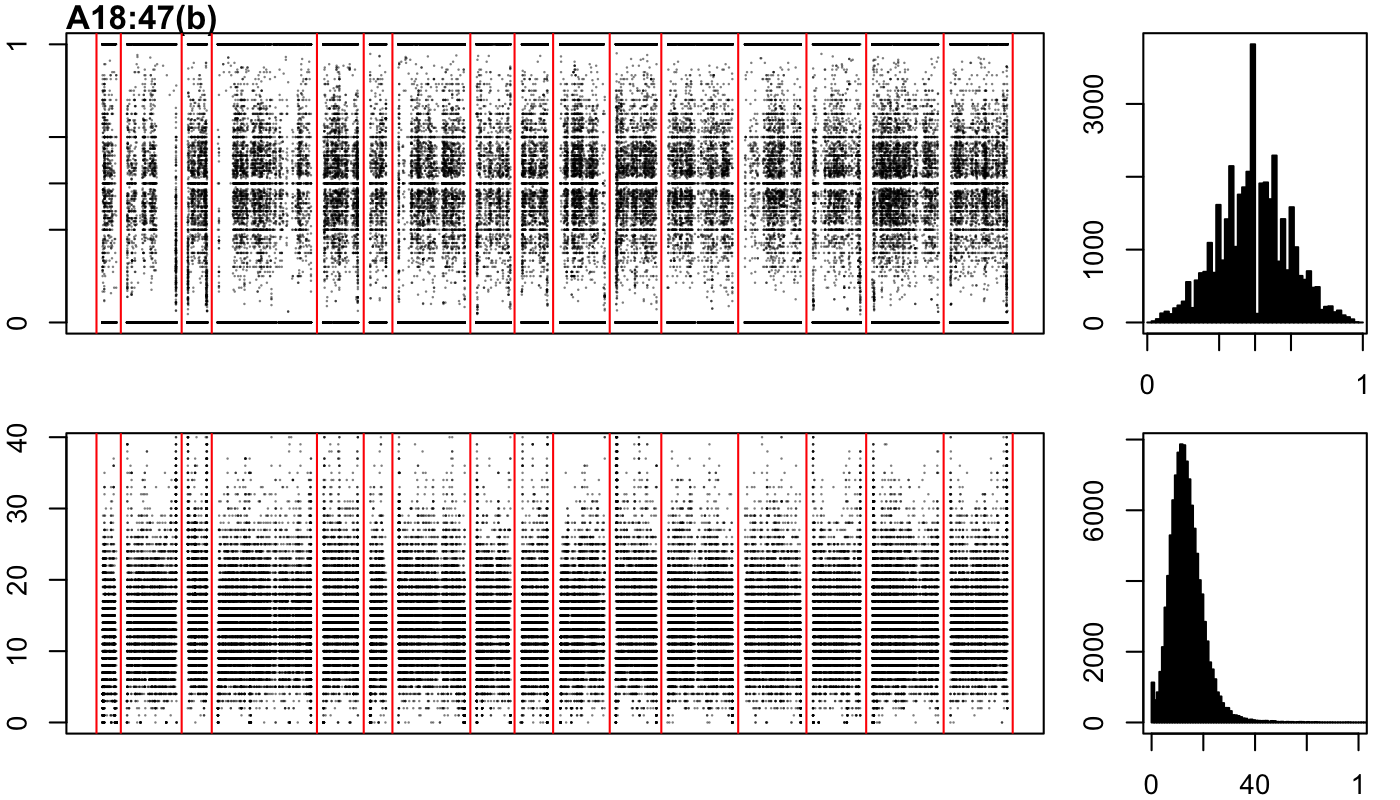

### A18-47-c.png

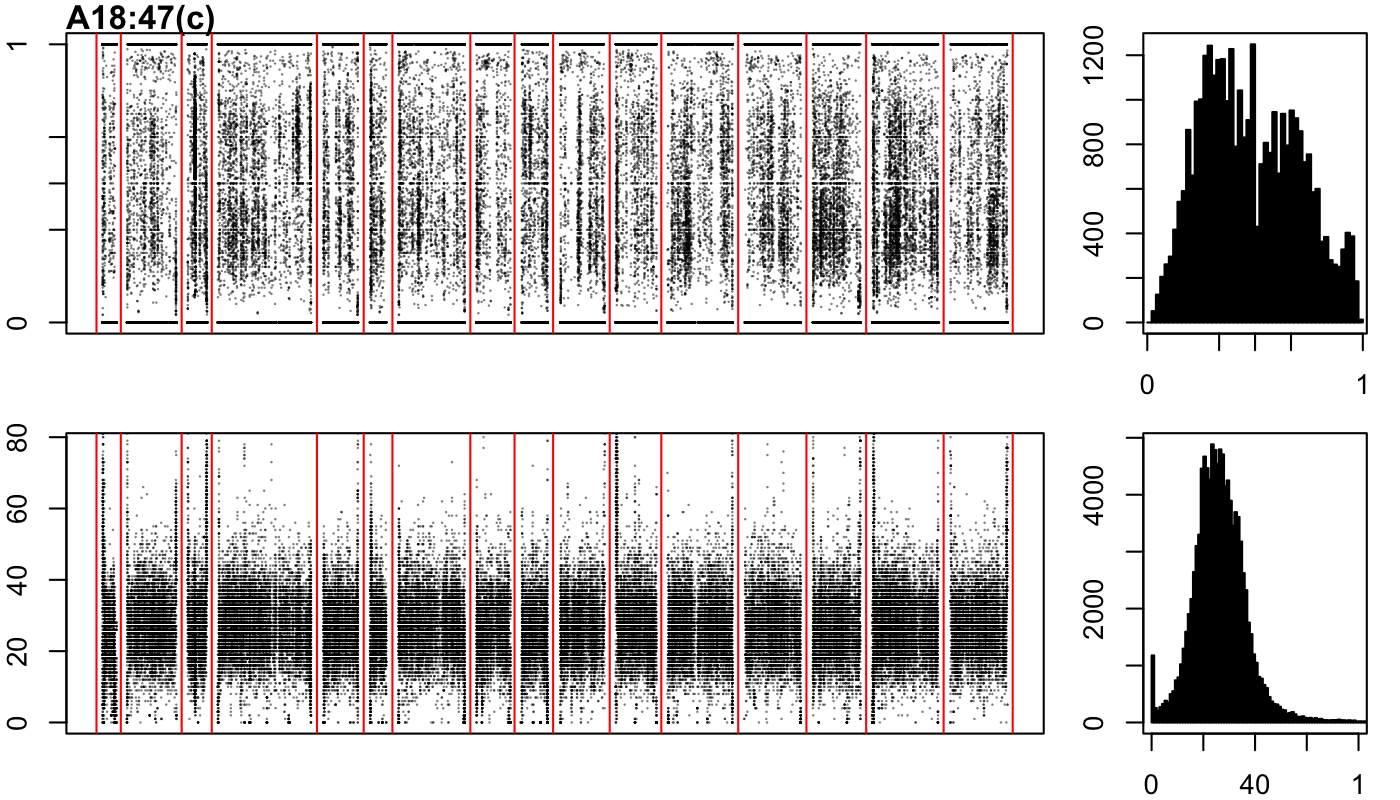

### A18-57-a.png

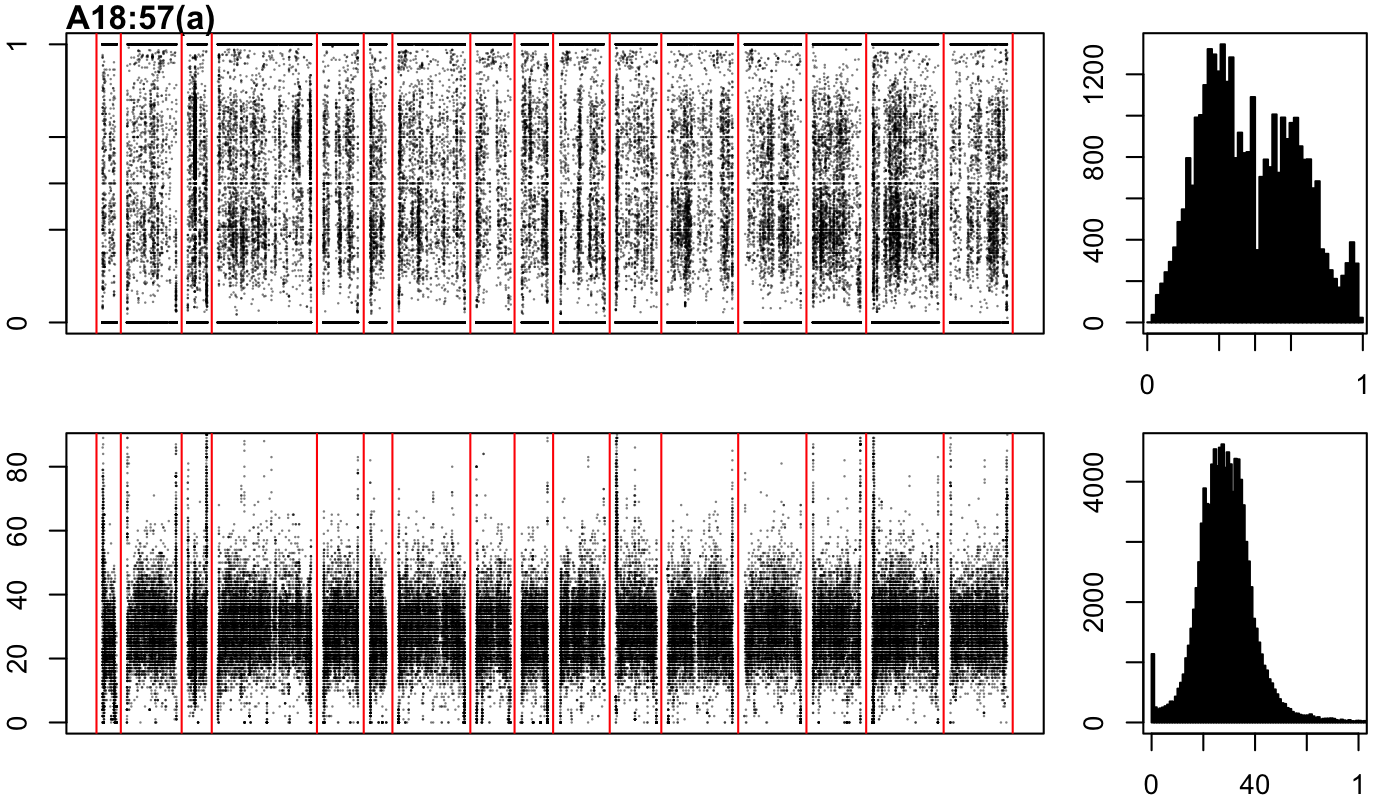

### A18-57-b.png

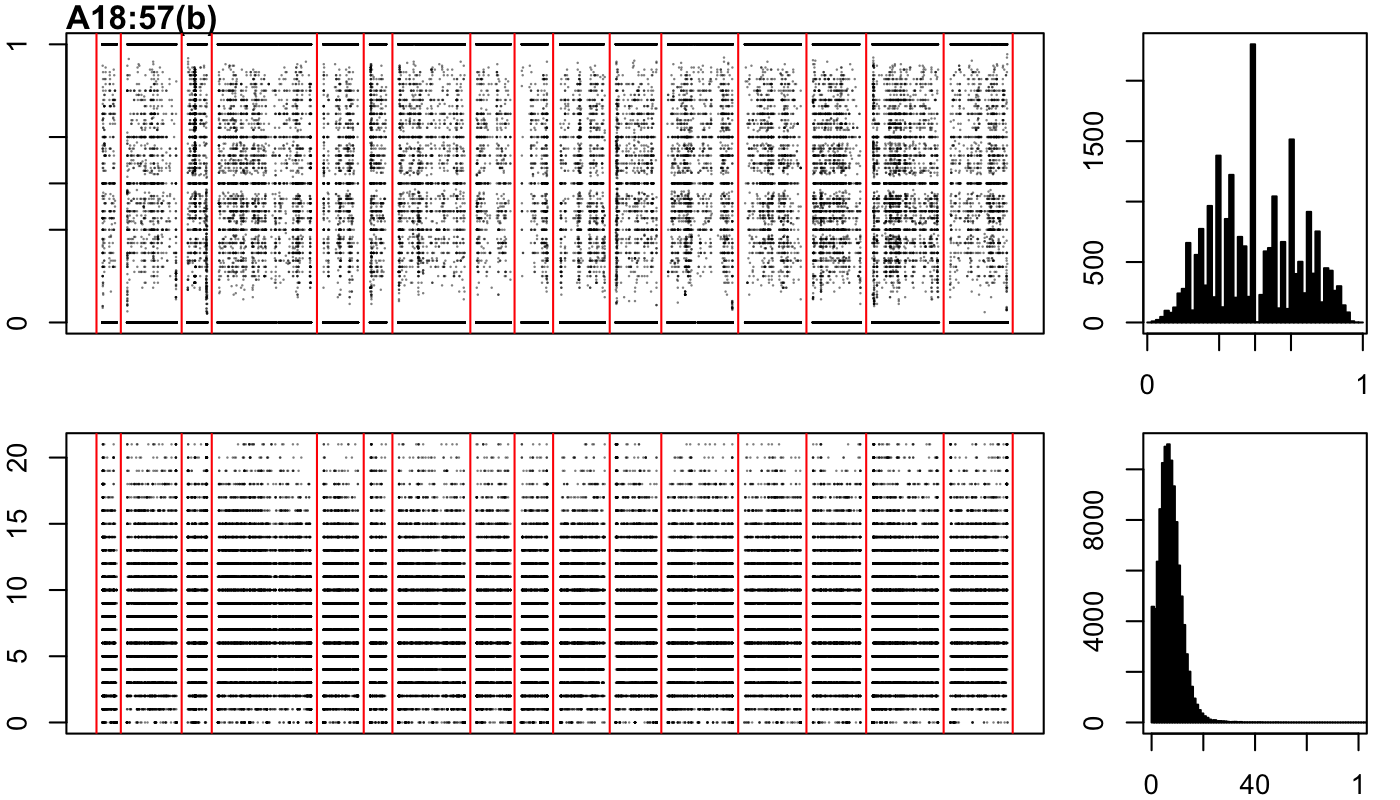

### A18-57-c.png

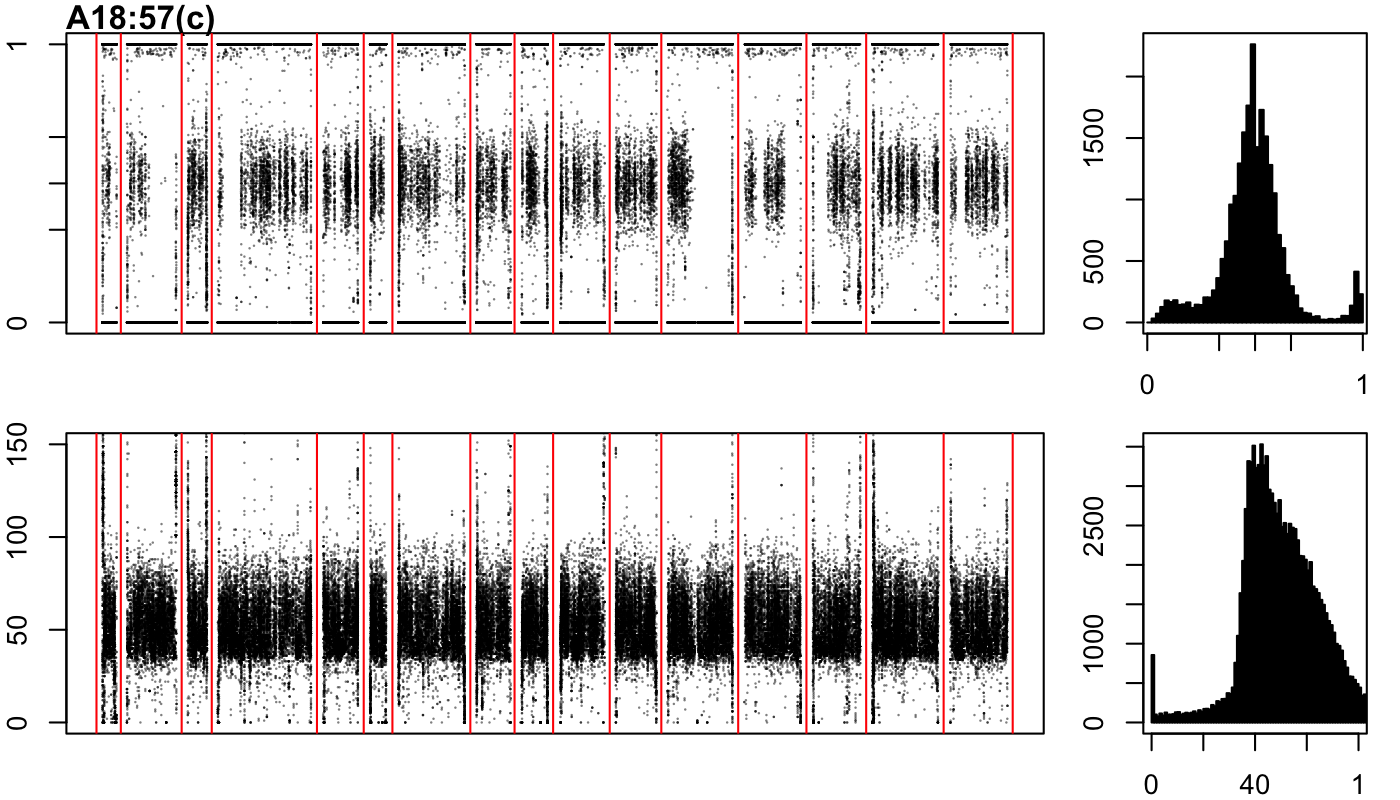

### A18-109-a.png

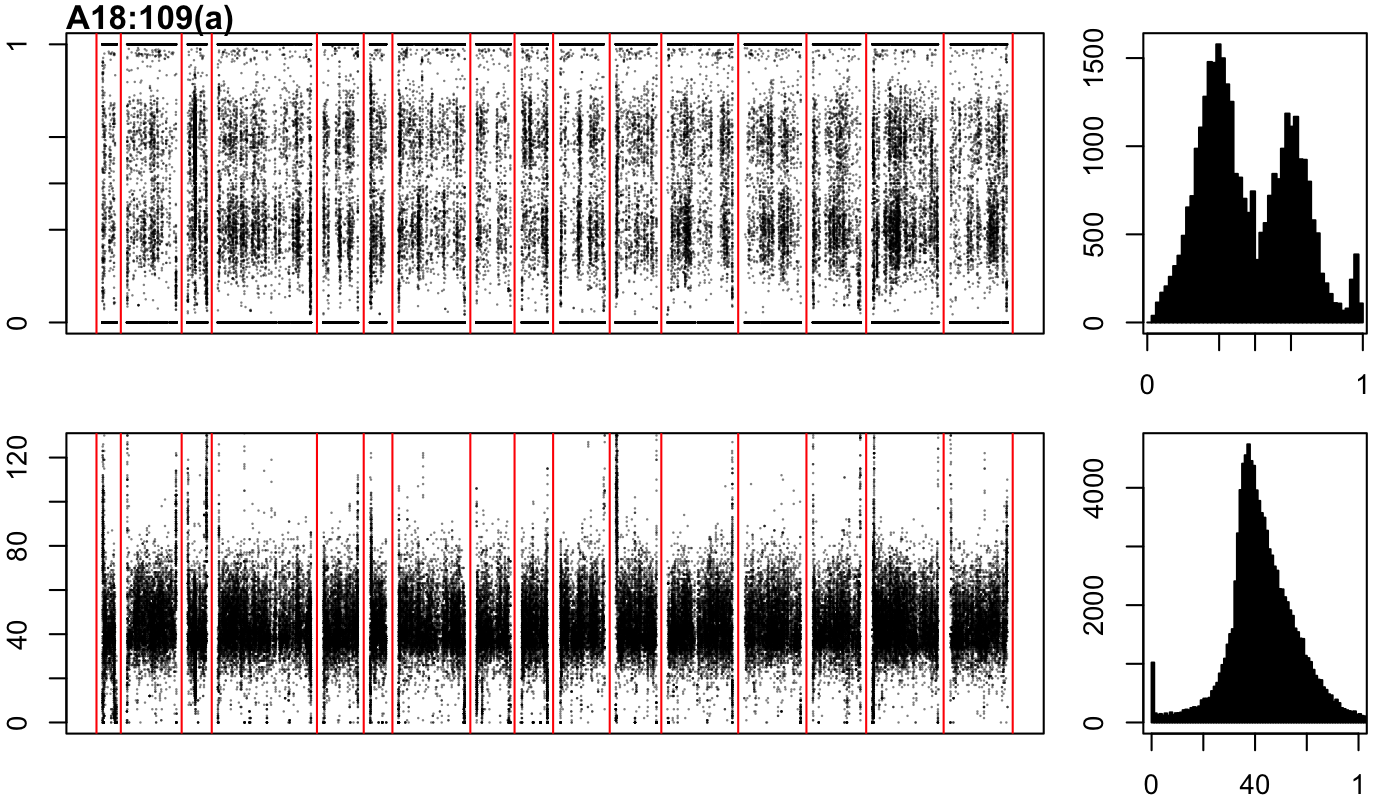

### A18-167-a.png

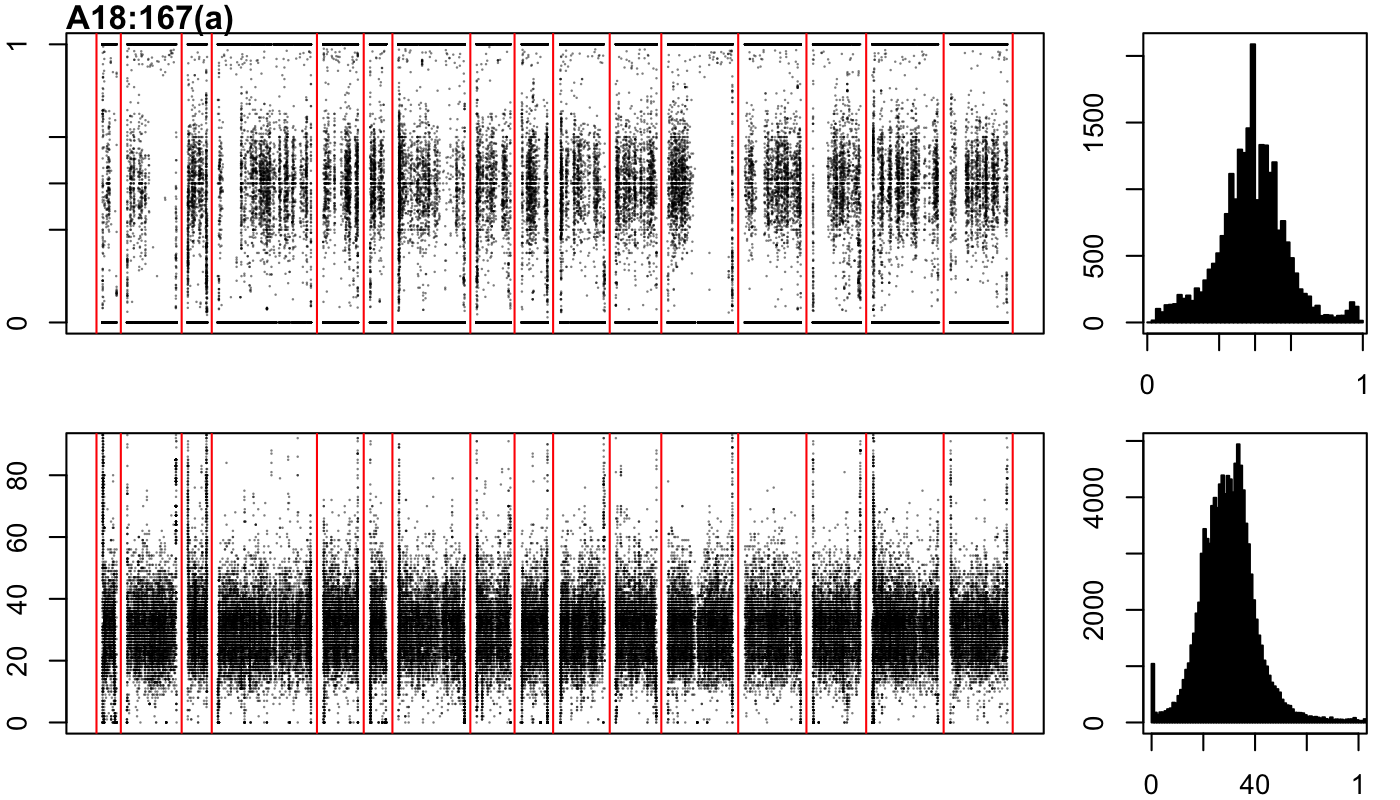

### A18-167-b.png

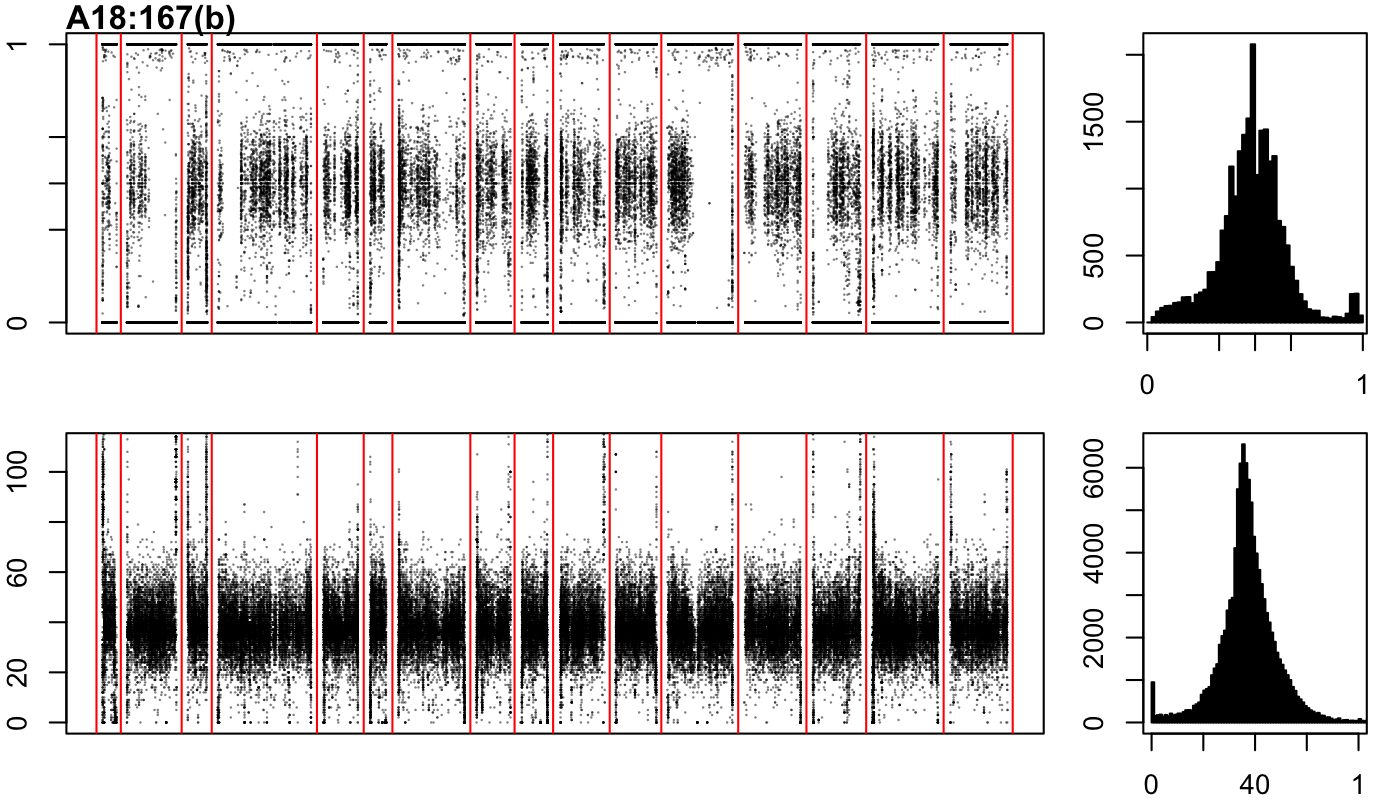

### A18-167-c.png

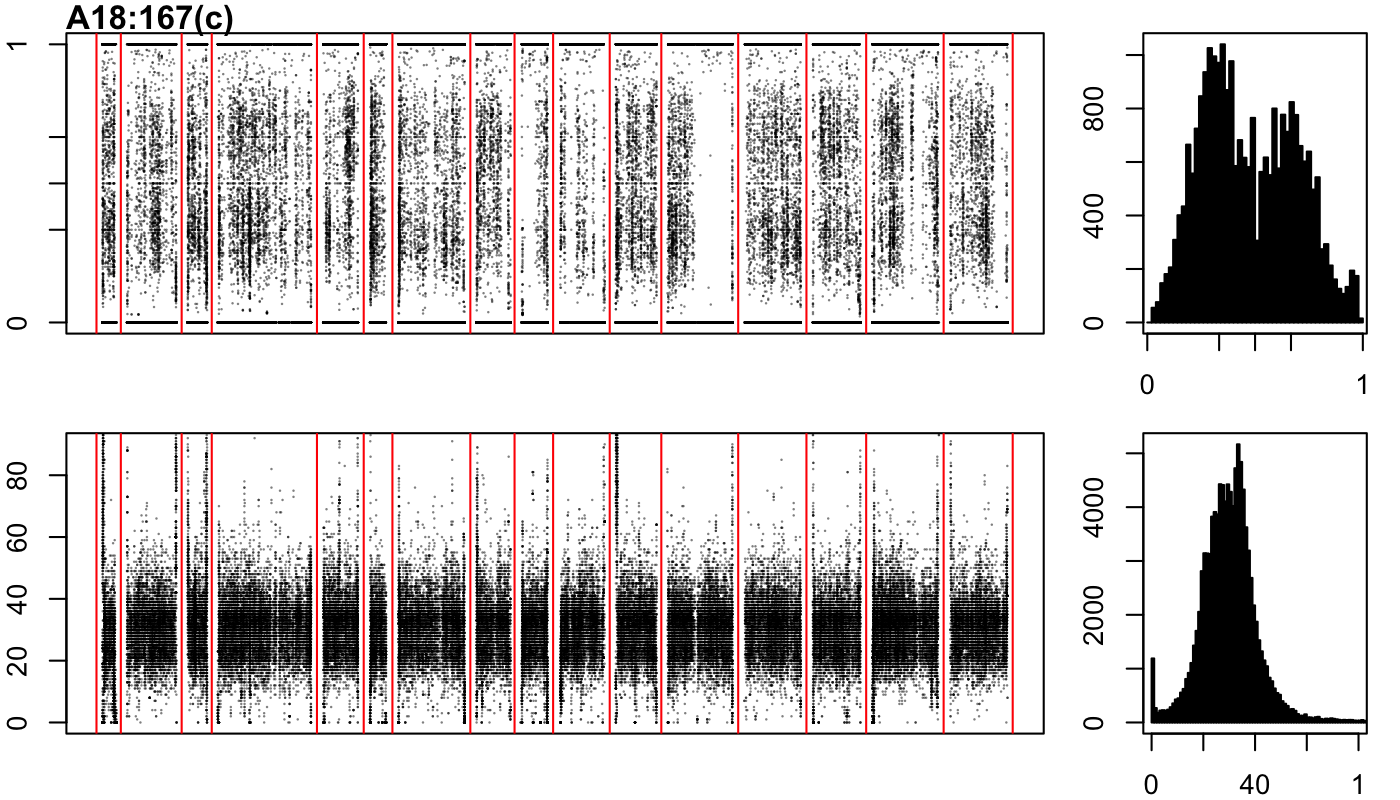

### A18-217-c.png

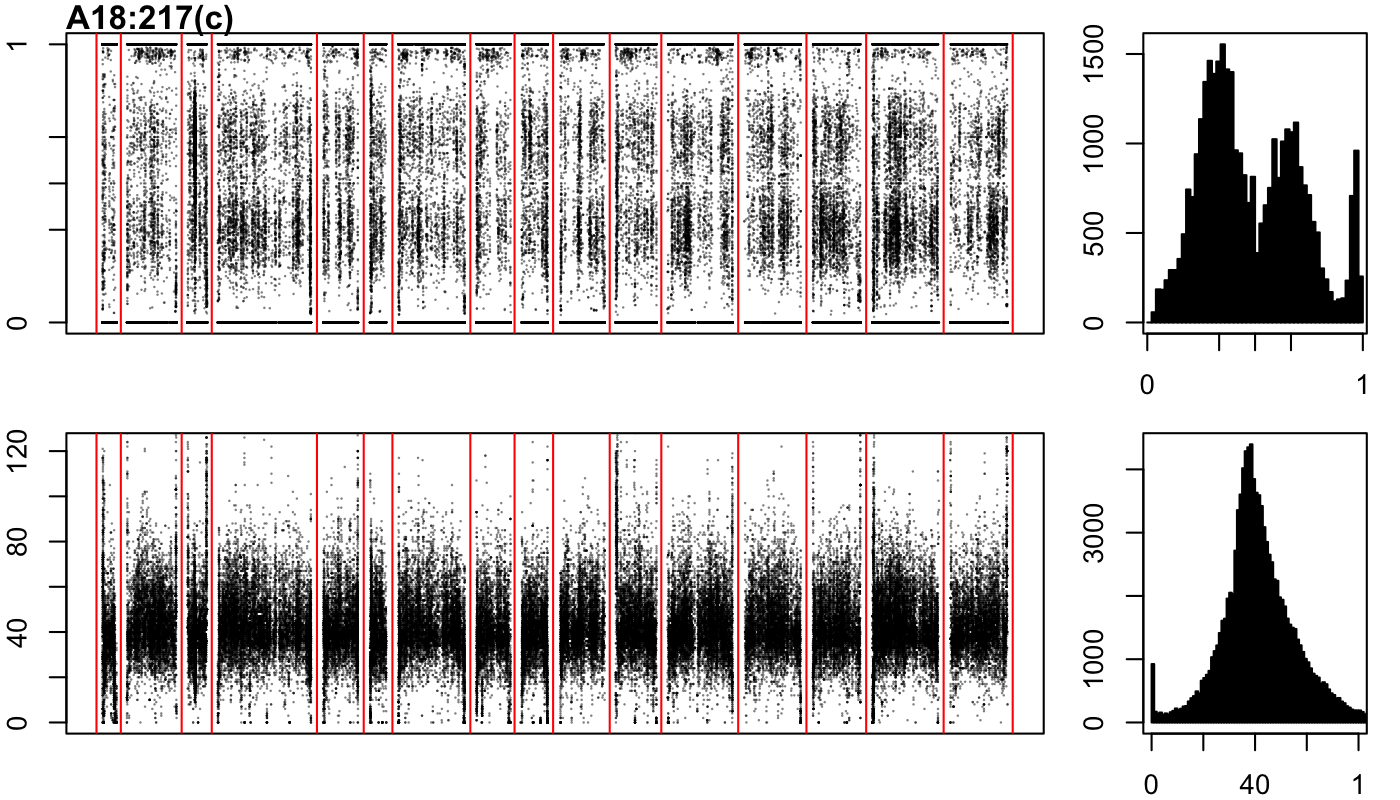

### A18-217-d.png

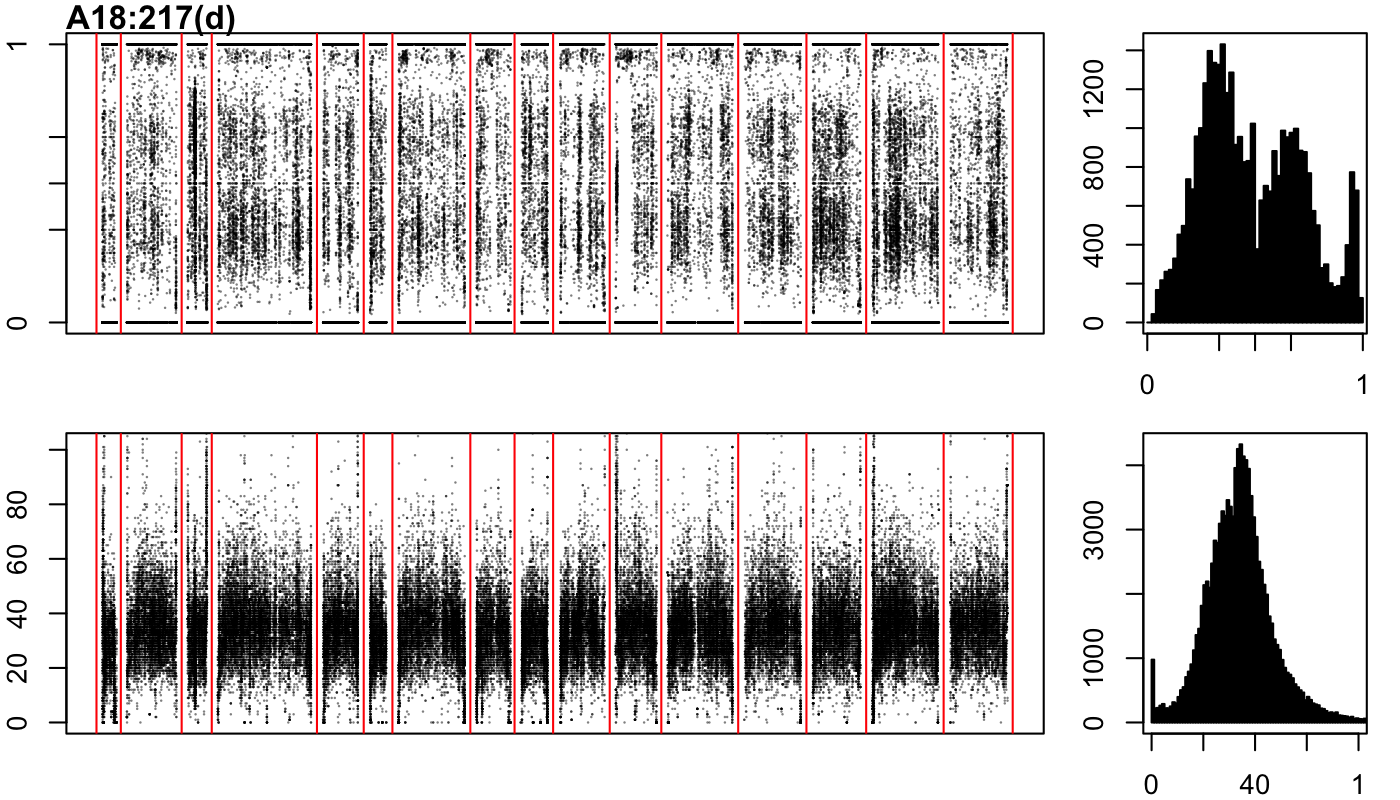

### A18-227-a.png

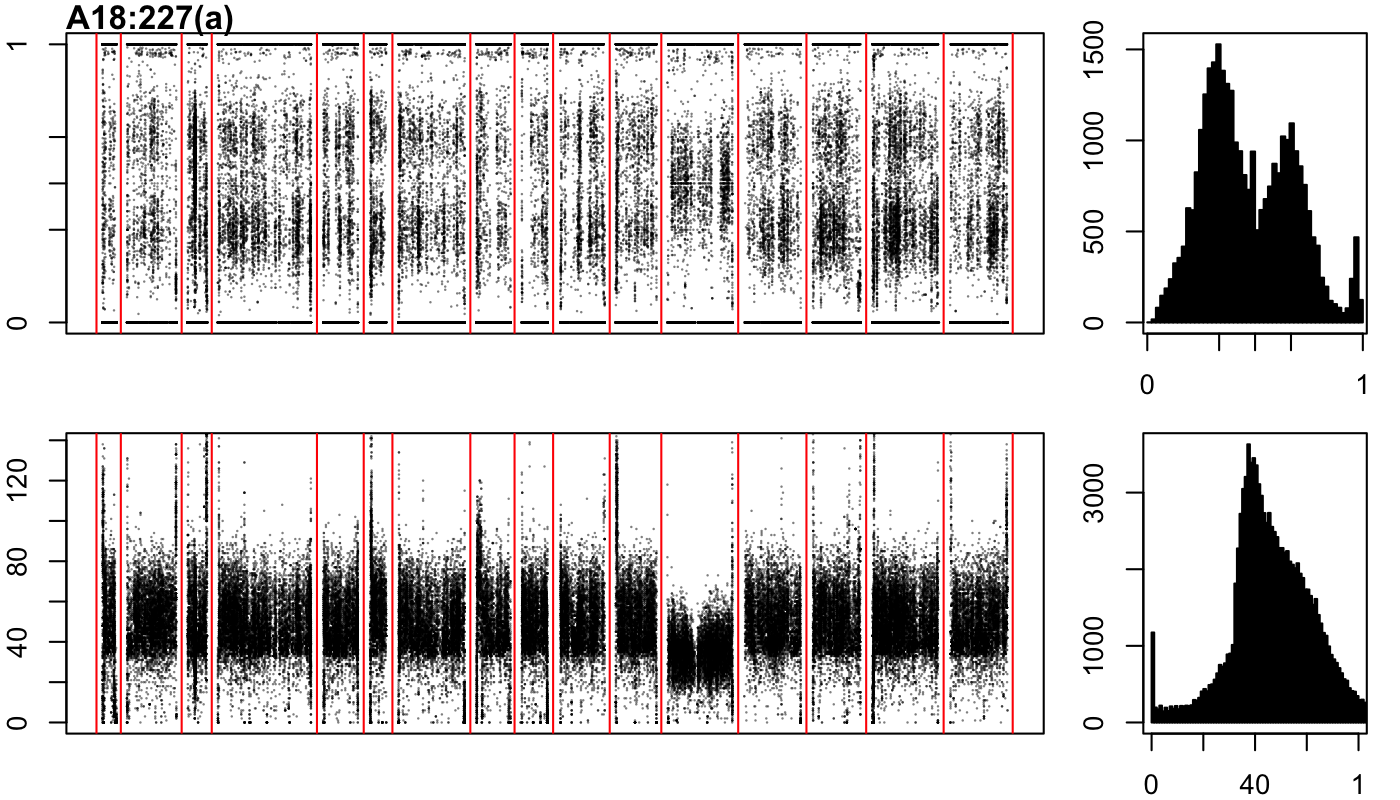

### A18-227-b.png

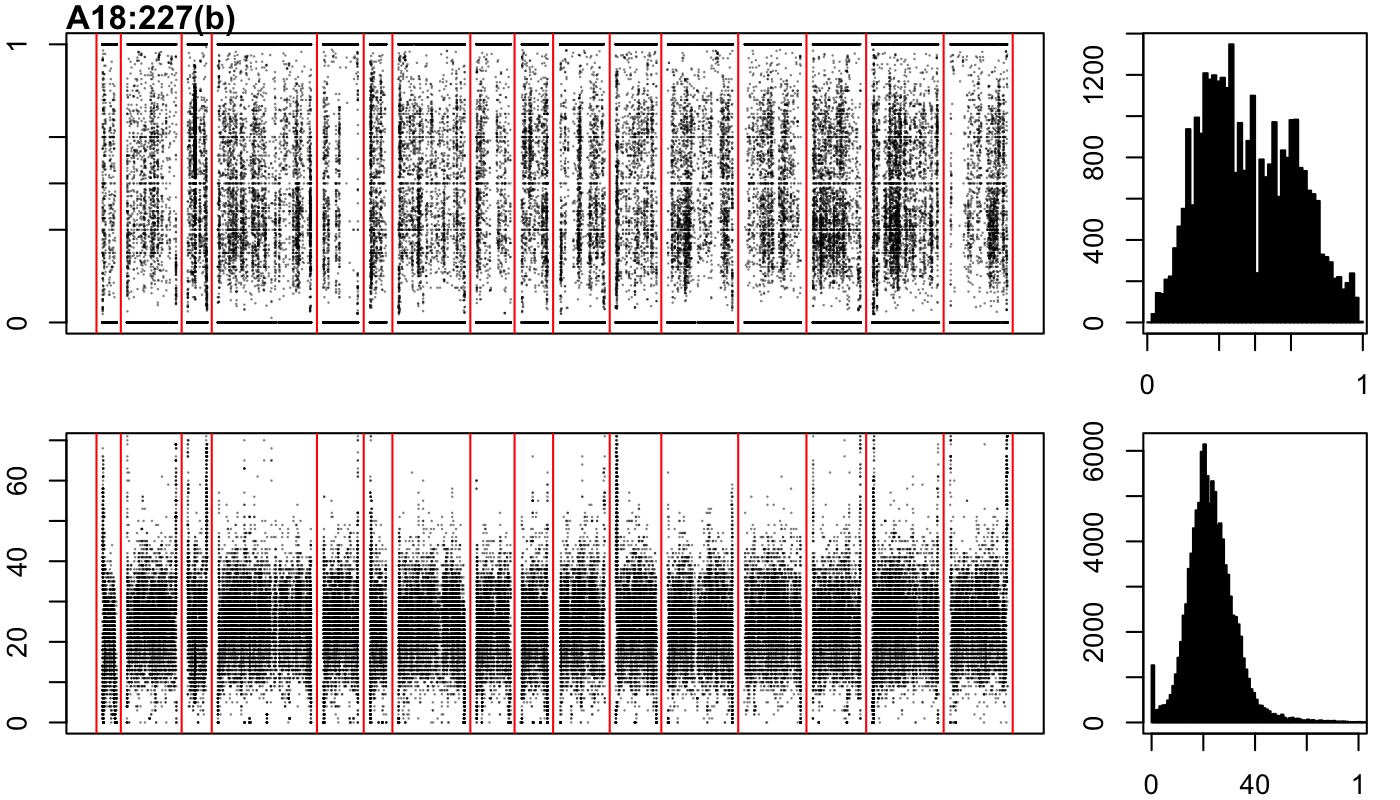

### A18-227-c.png

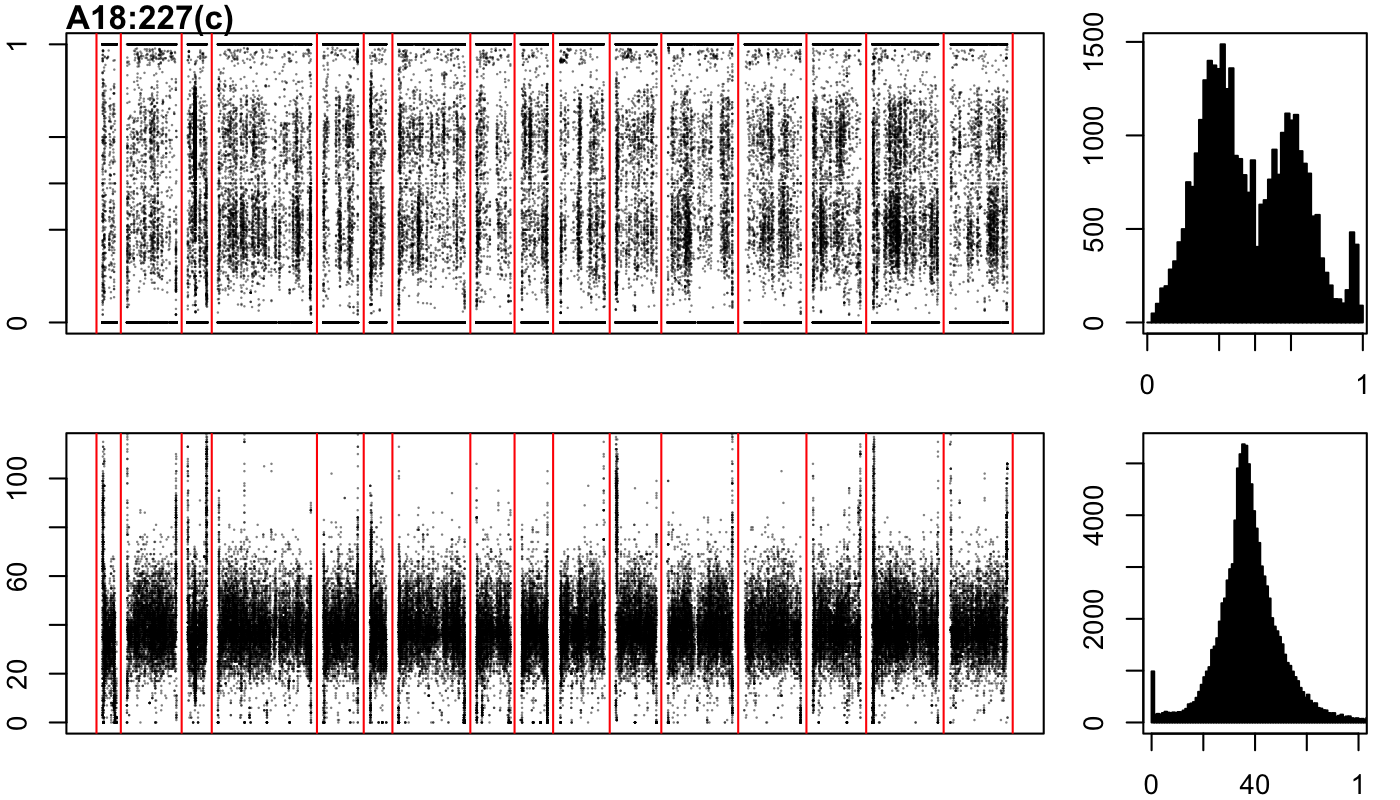

### A19-1-a.png

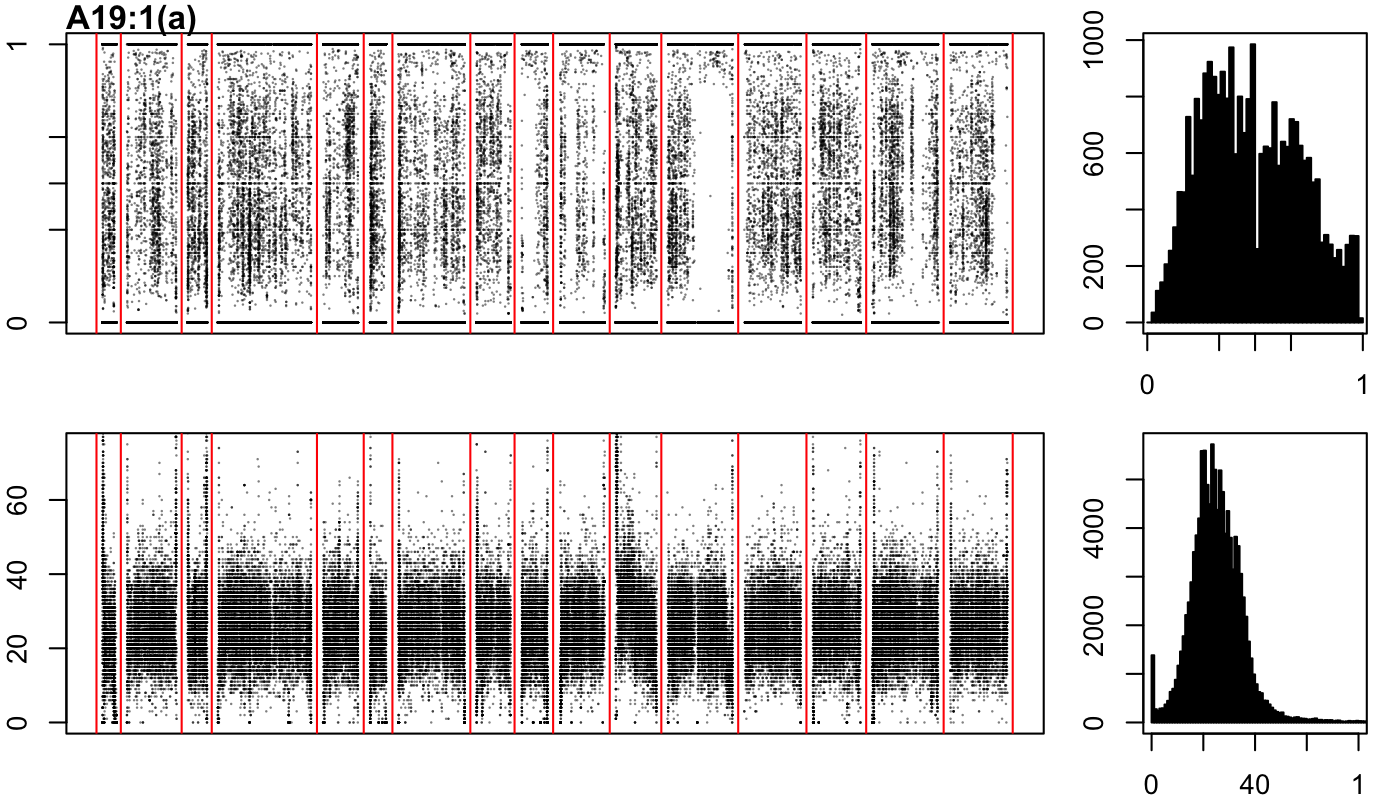

### A19-1-b.png

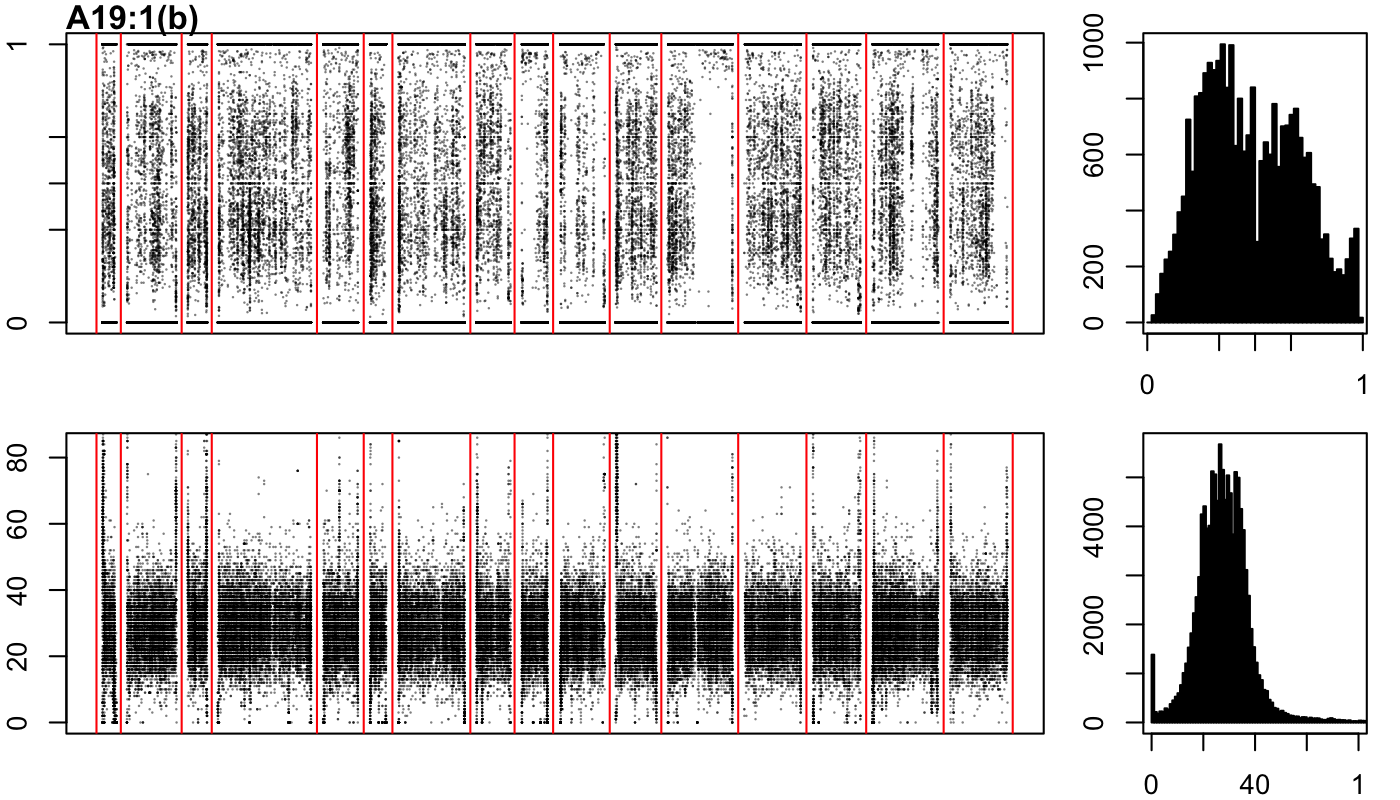

### A19-1-c.png

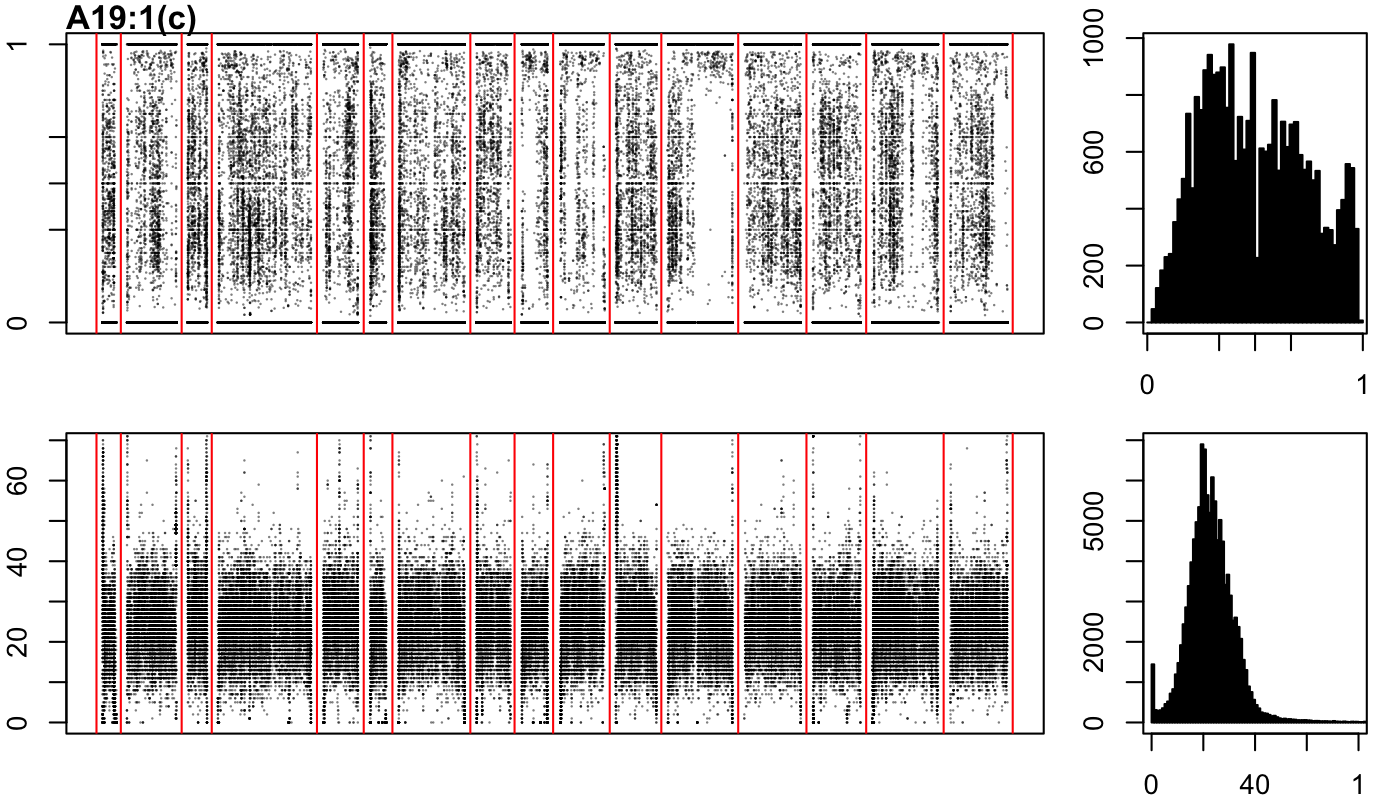

### A19-8-a.png

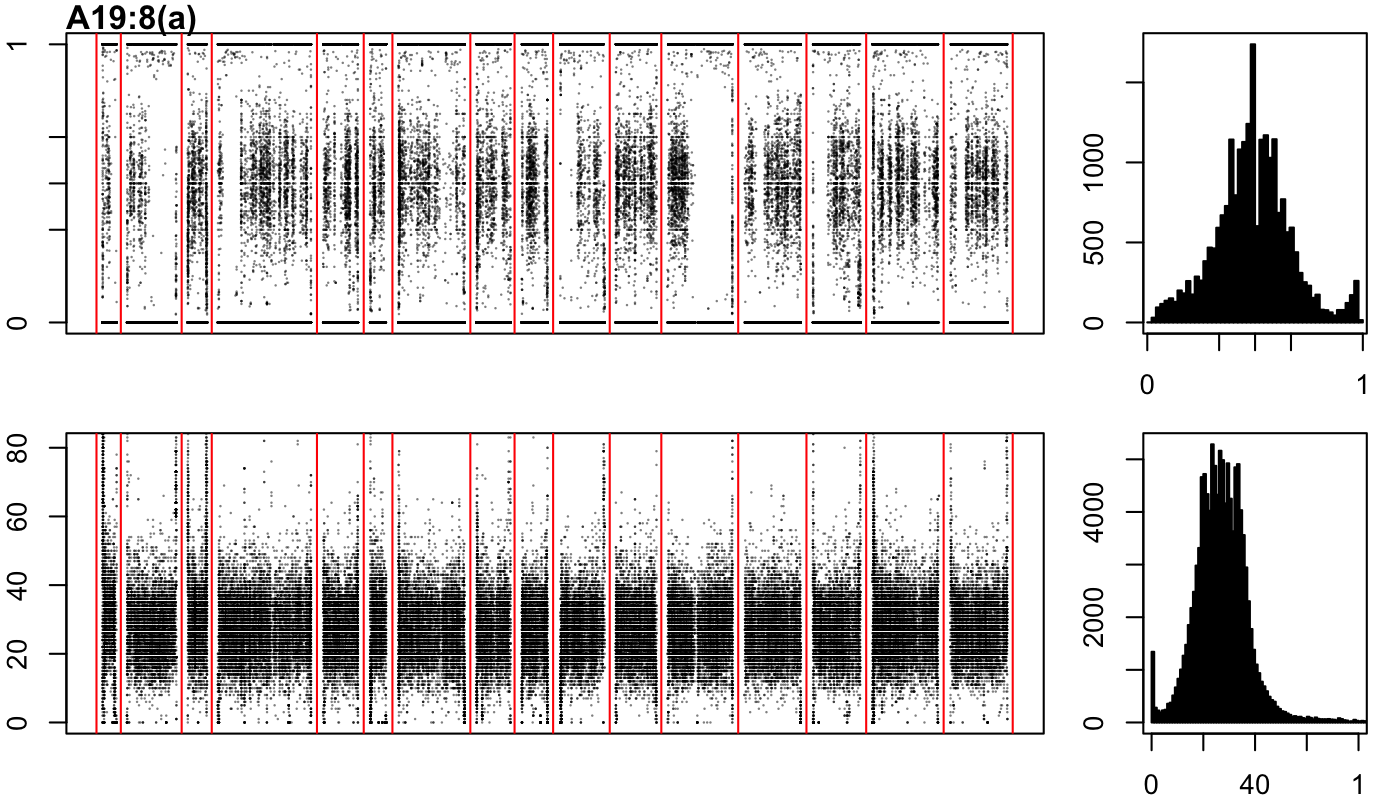

### A19-8-b.png

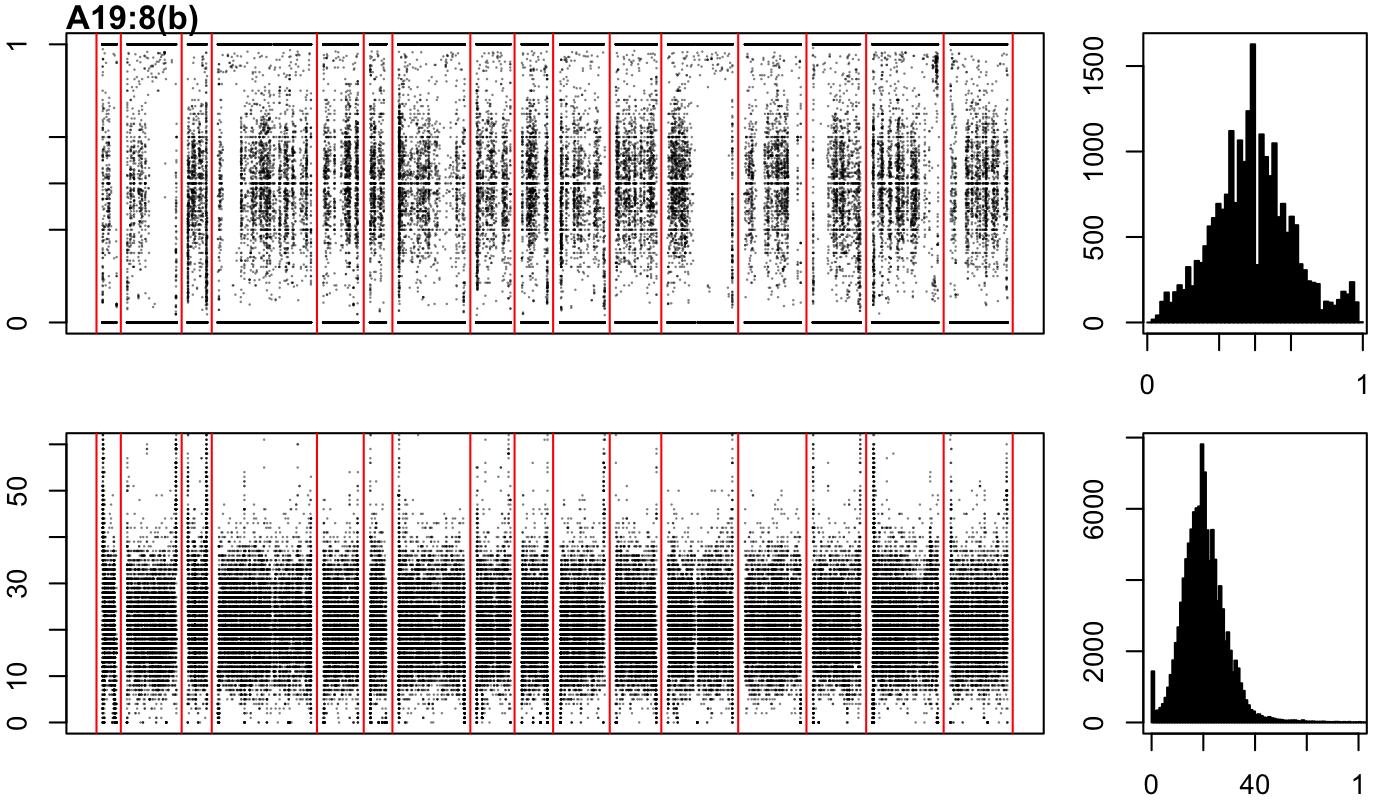

### A19-8-c.png

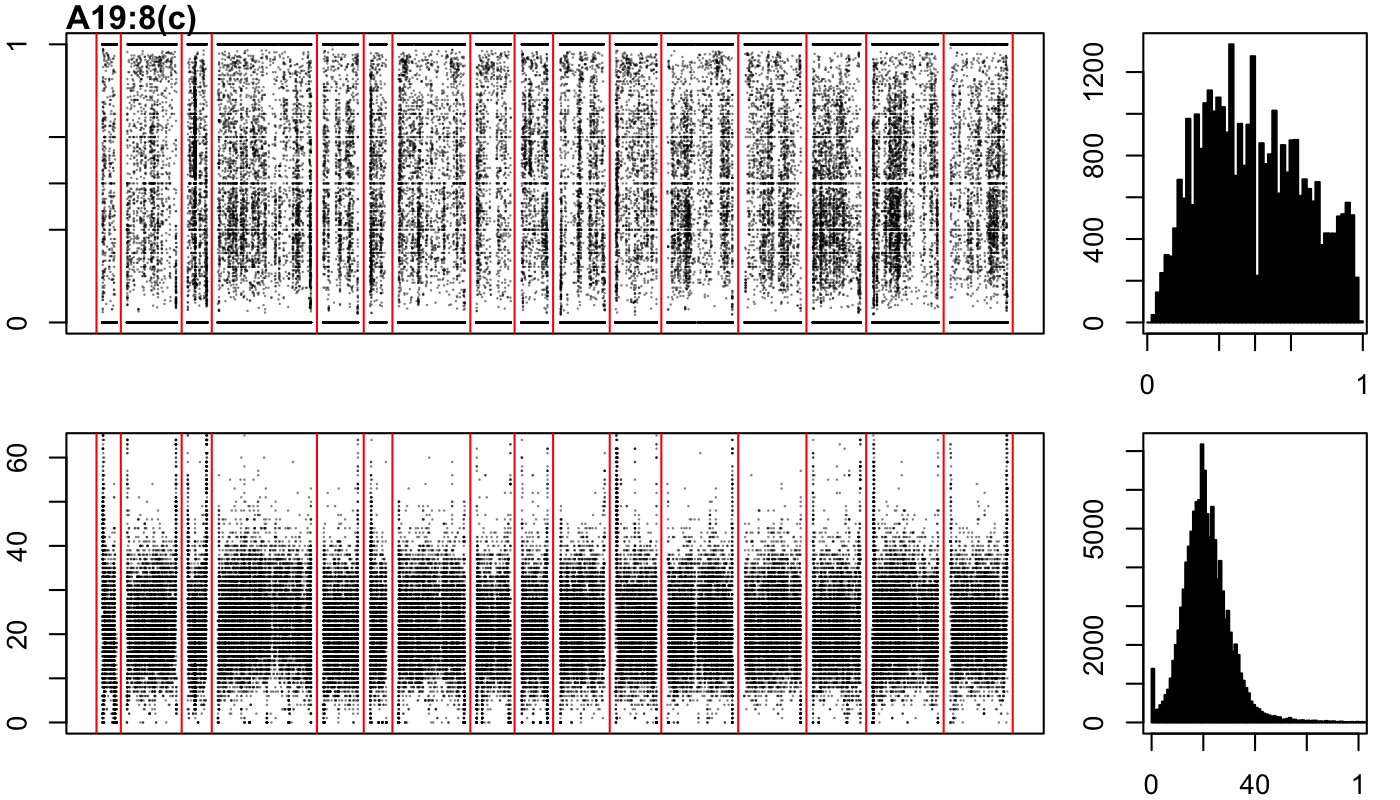

### A19-27-a.png

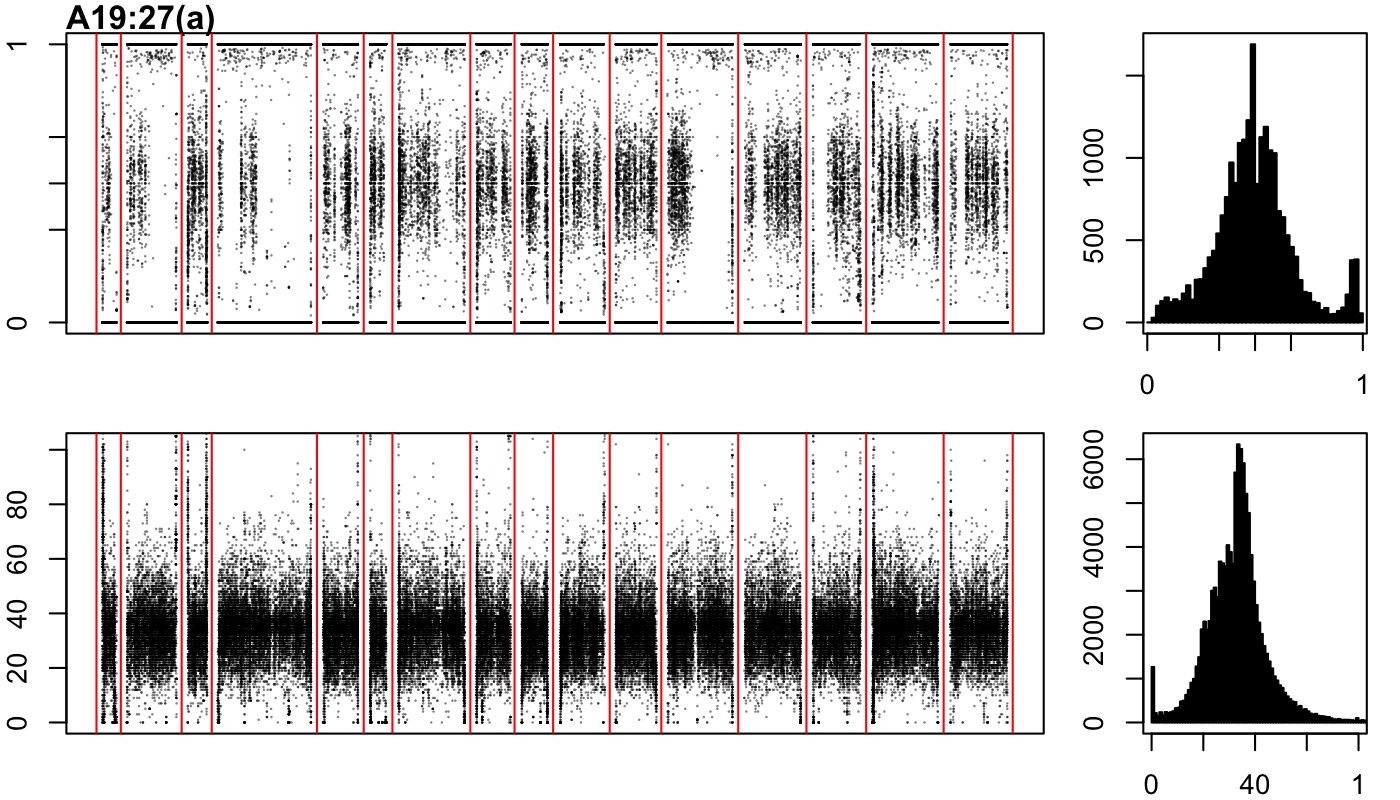

### A19-27-b.png

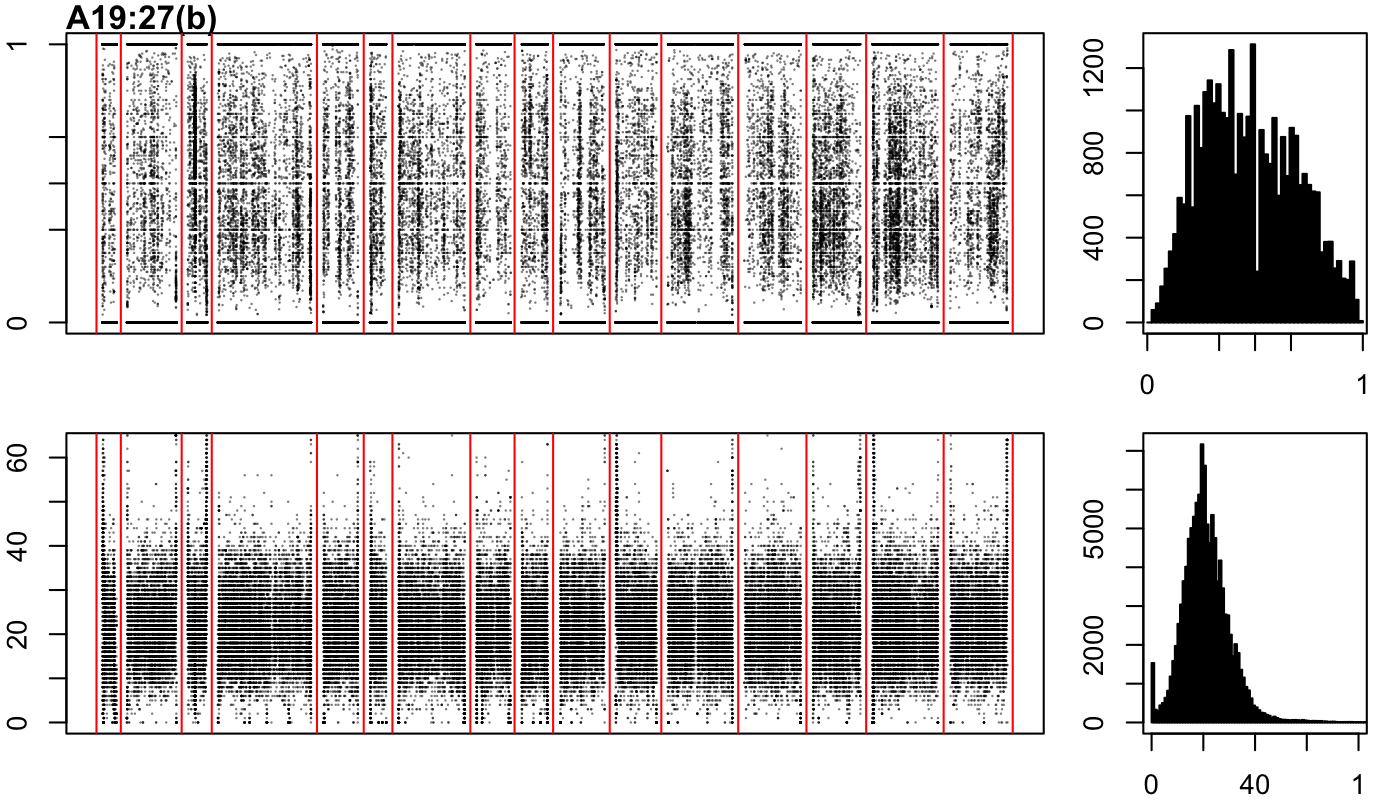

### A19-38-a.png

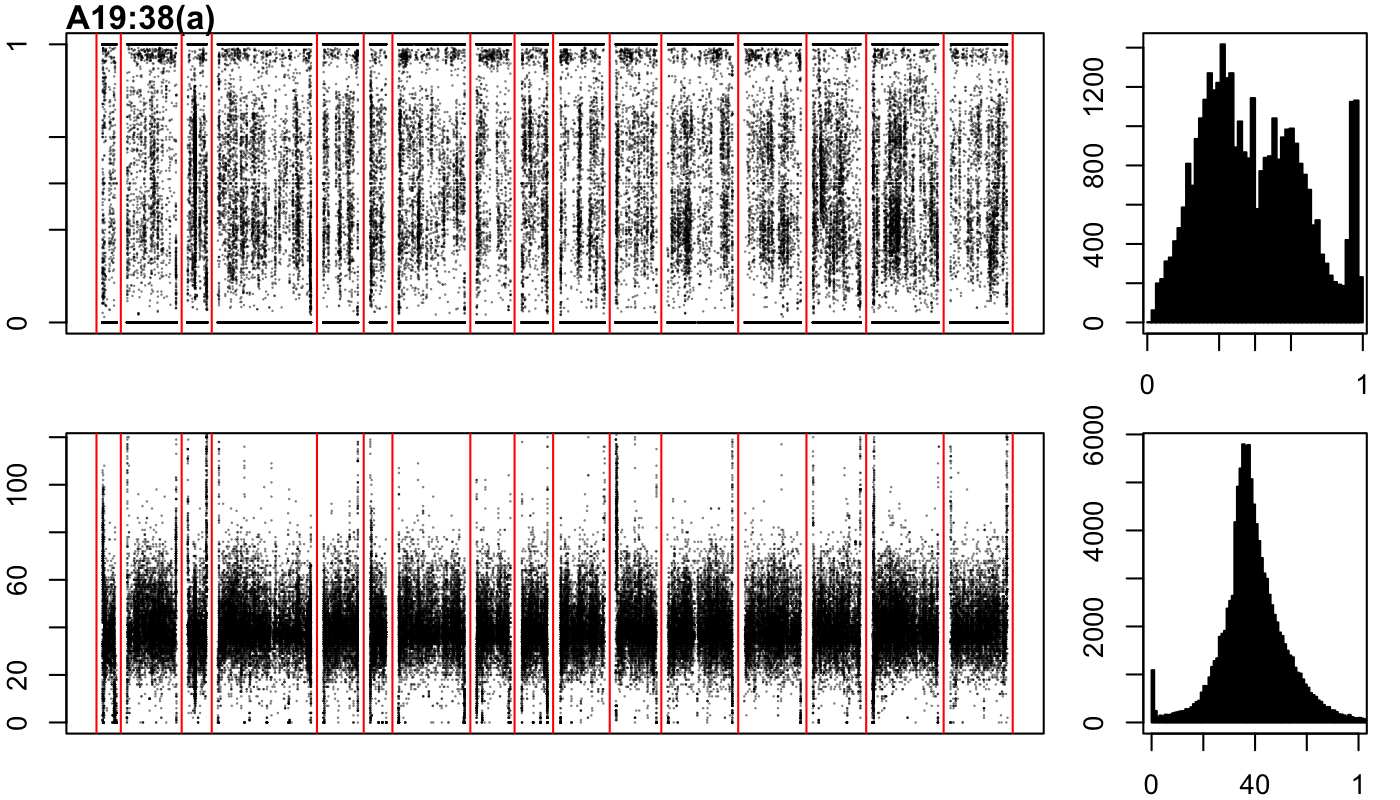

### A19-38-b.png

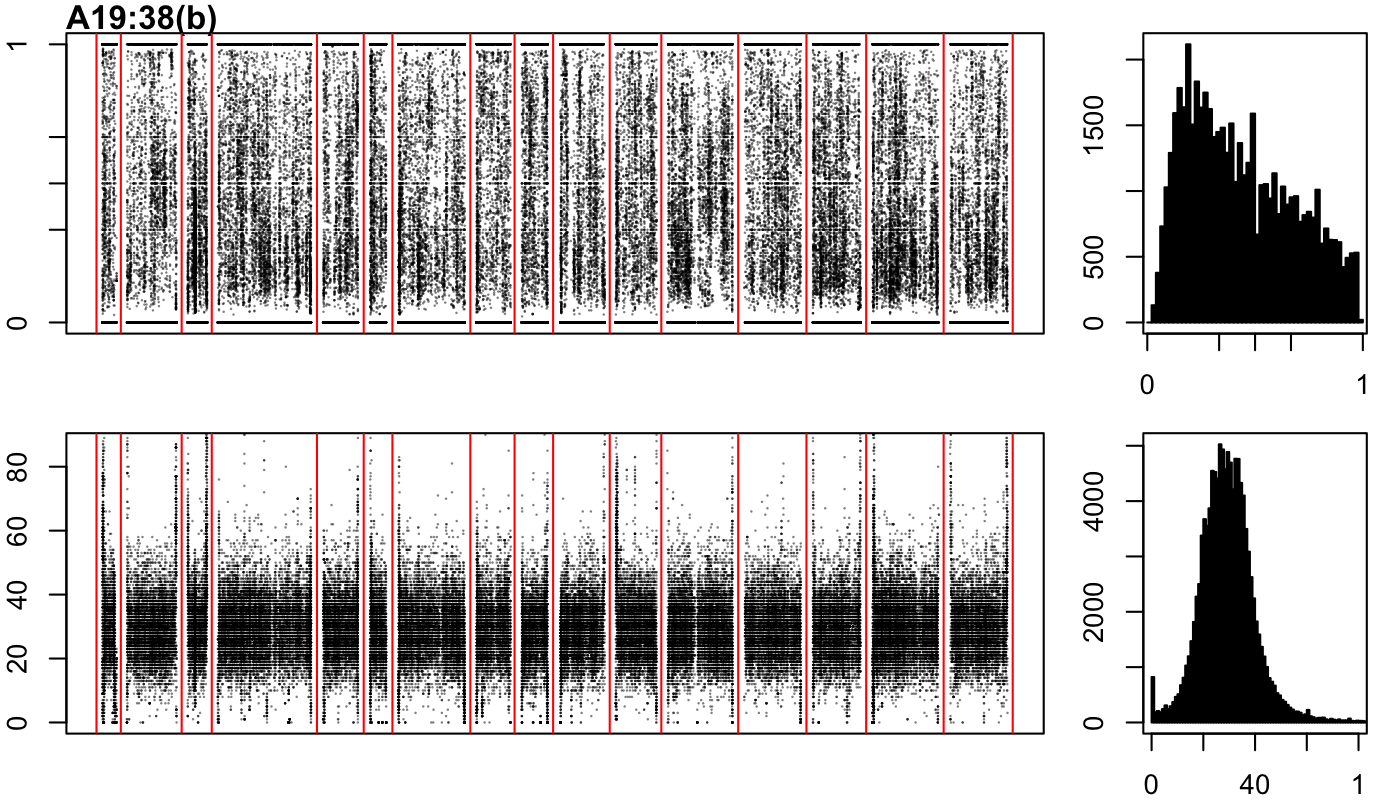

### A19-51-a.png

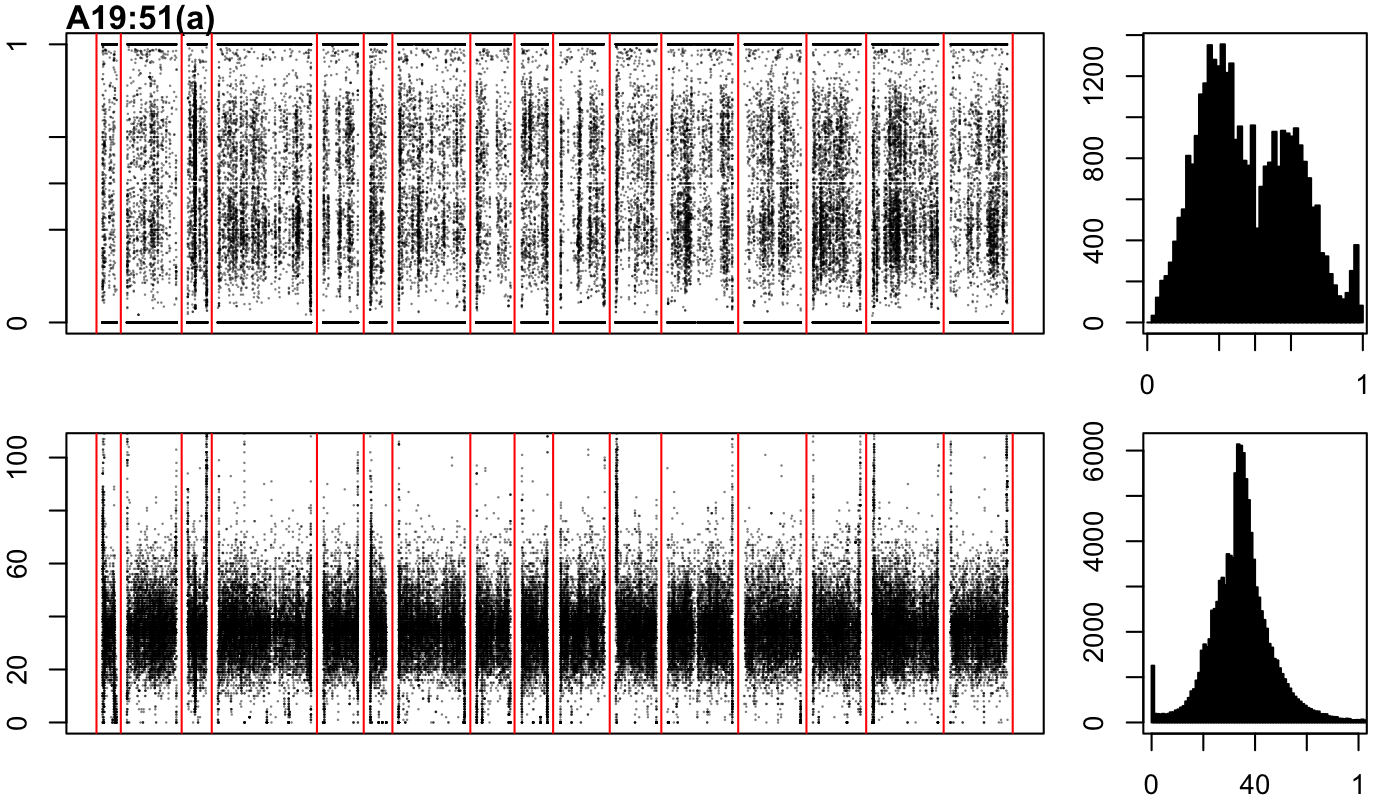

### A19-65-a.png

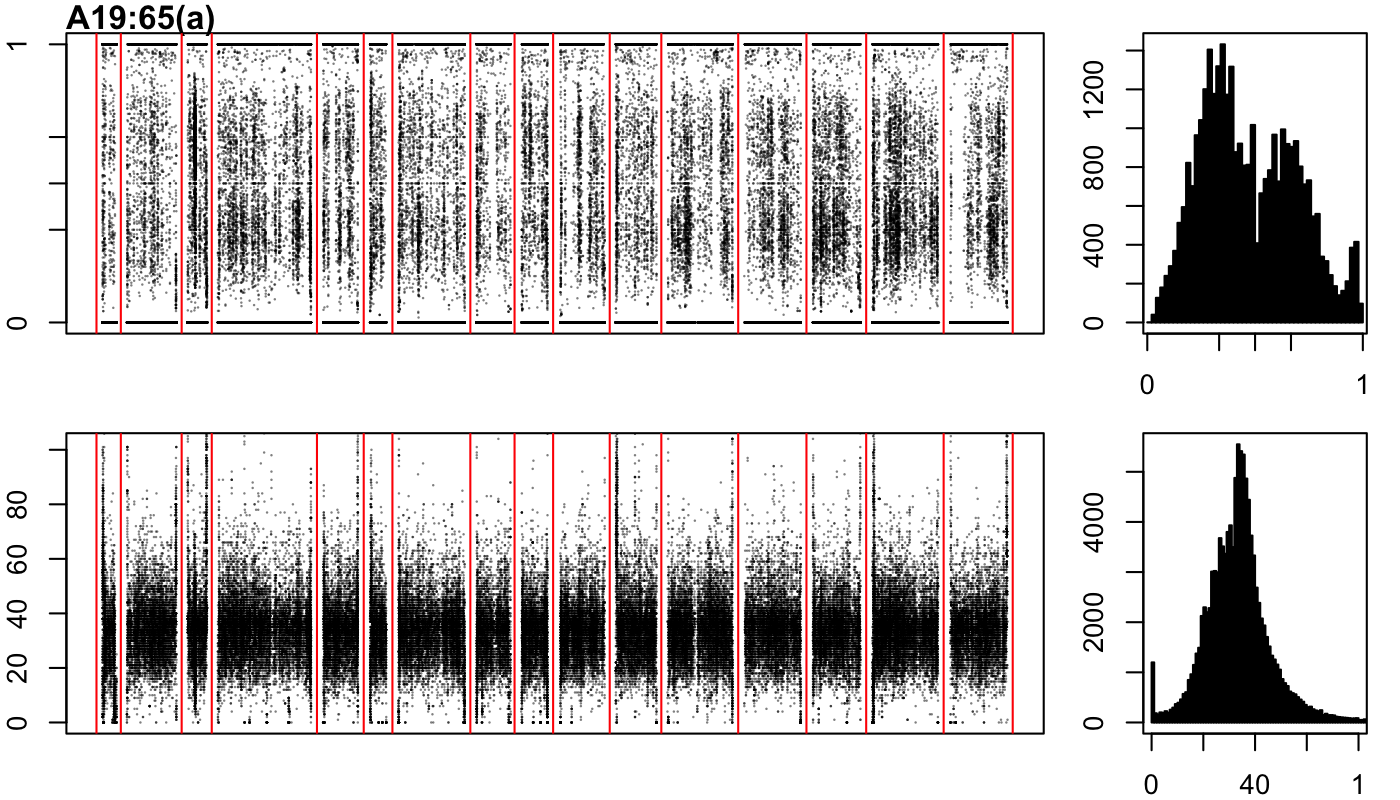

### A19-65-b.png

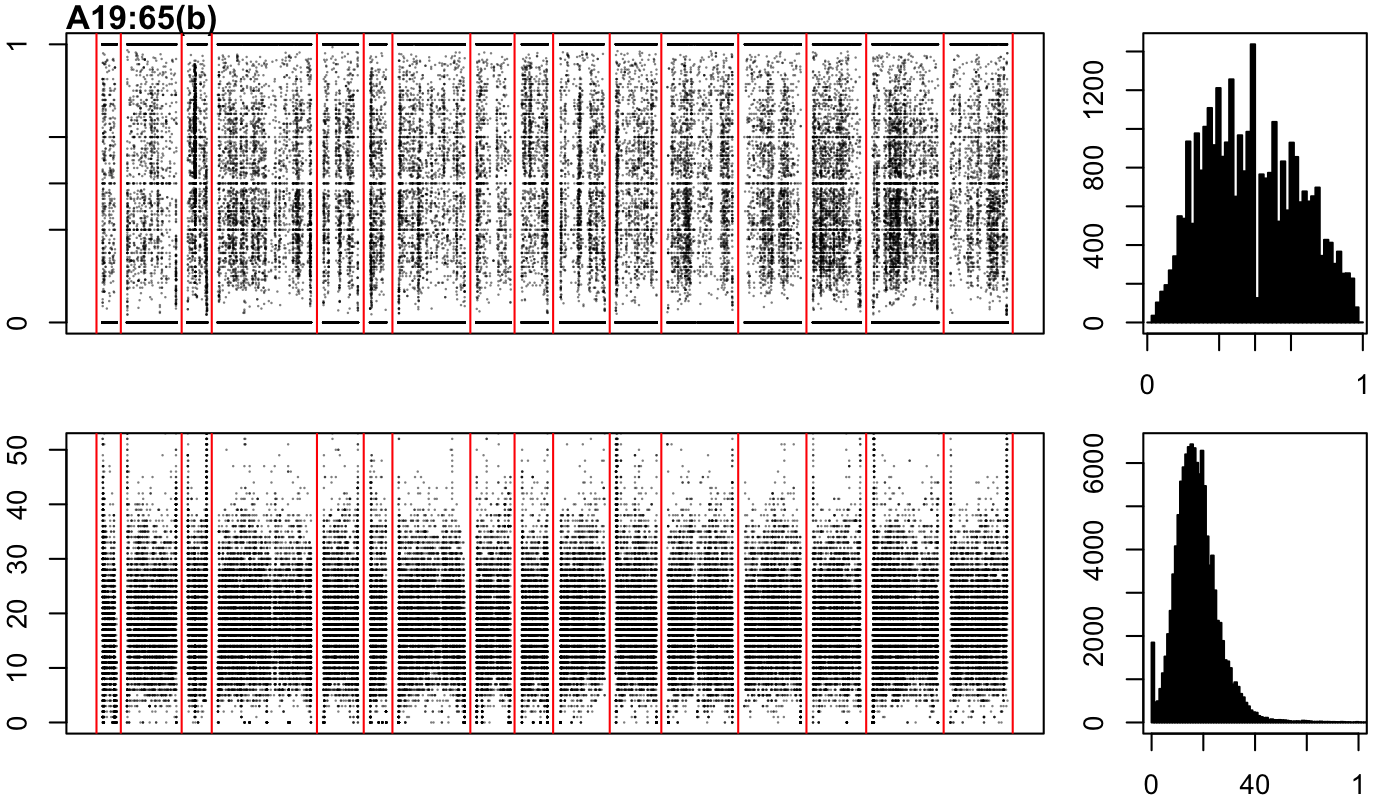

### A19-77-b.png

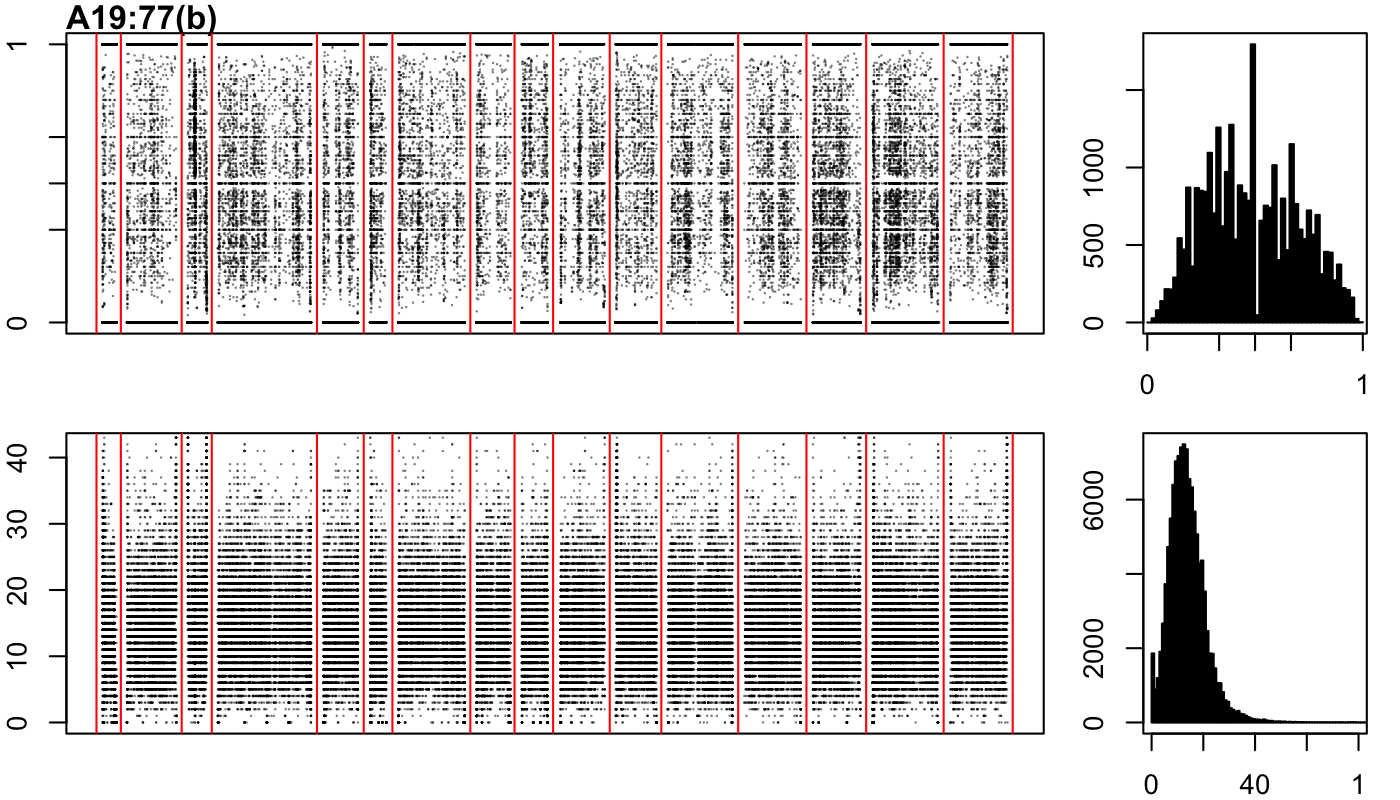

### A19-107-a.png

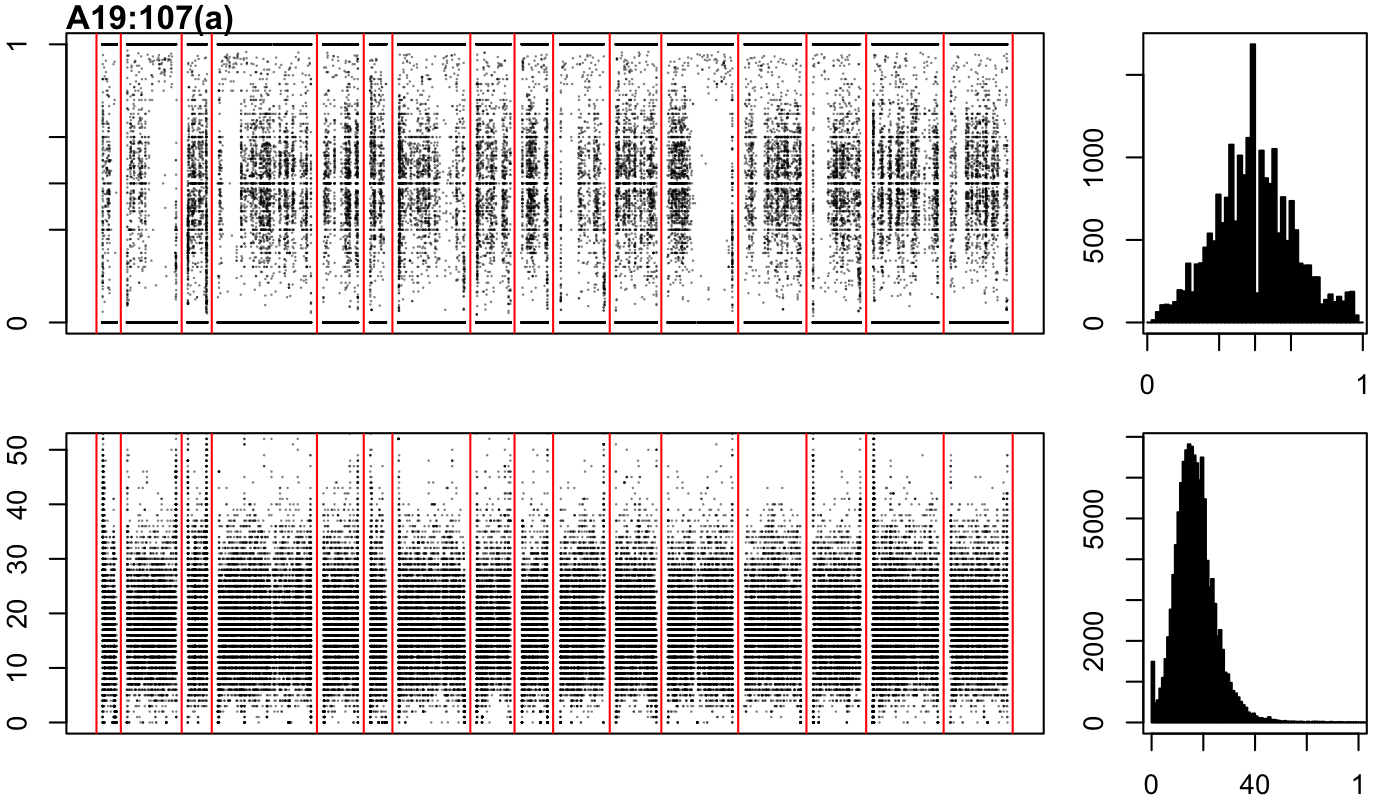

### A19-107-b.png

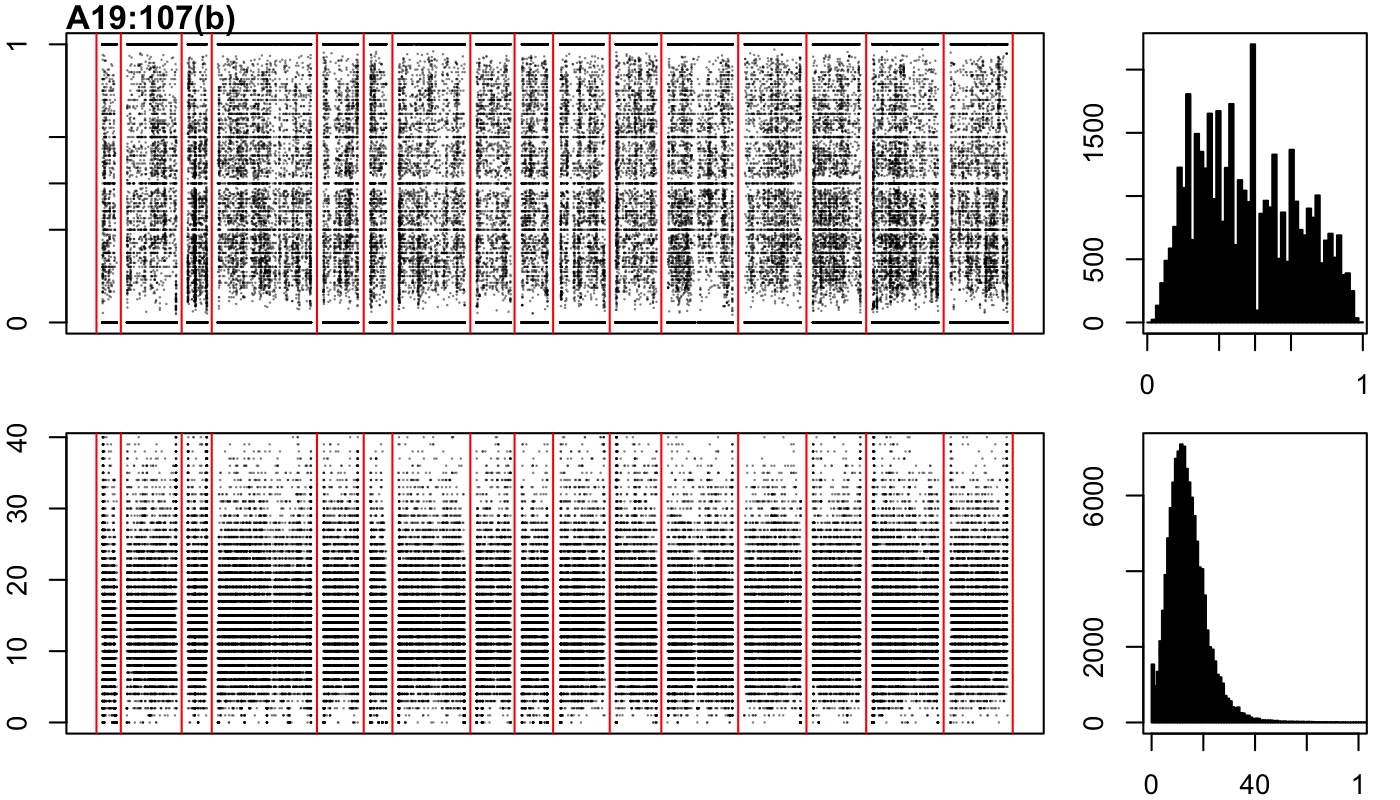

### A19-150-a.png

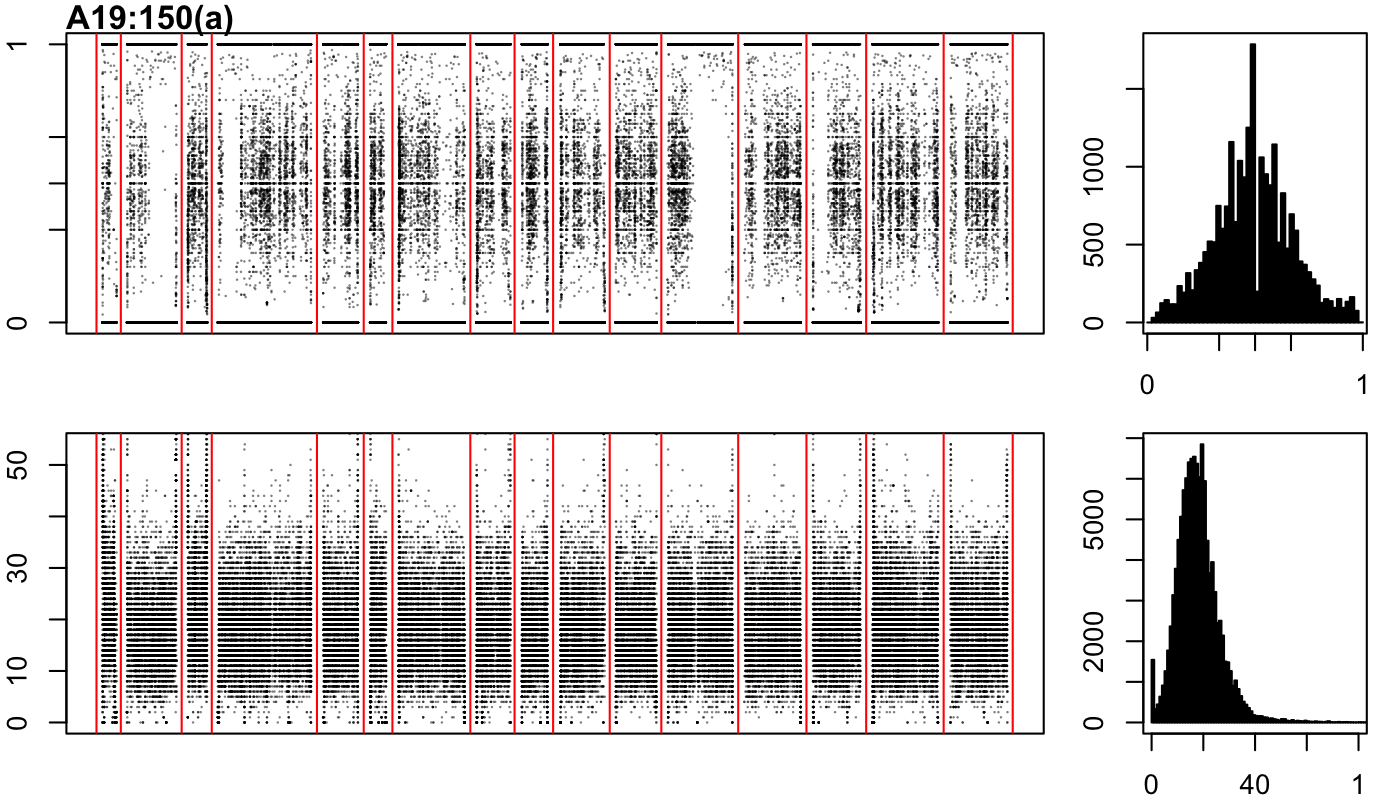

### A19-150-b.png

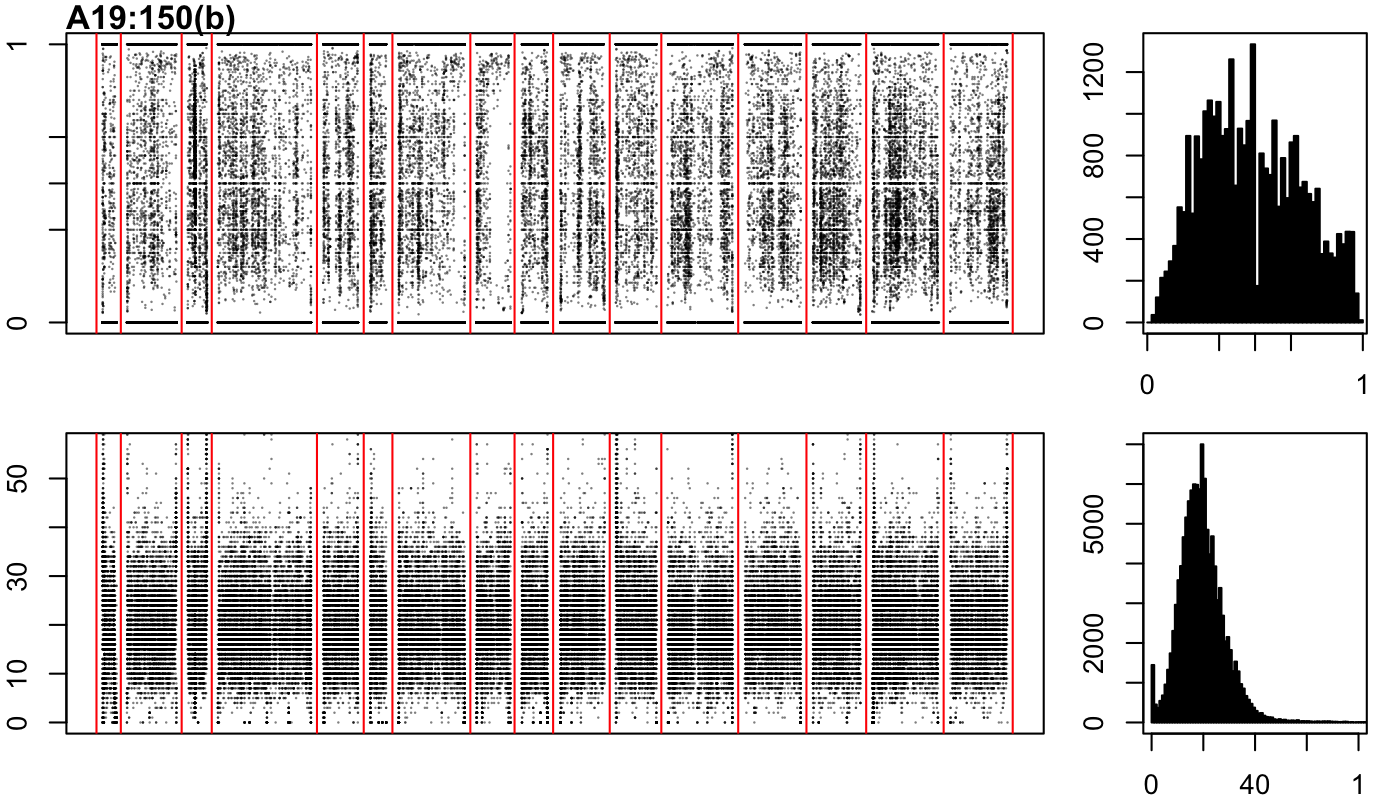

### A19-163-a.png

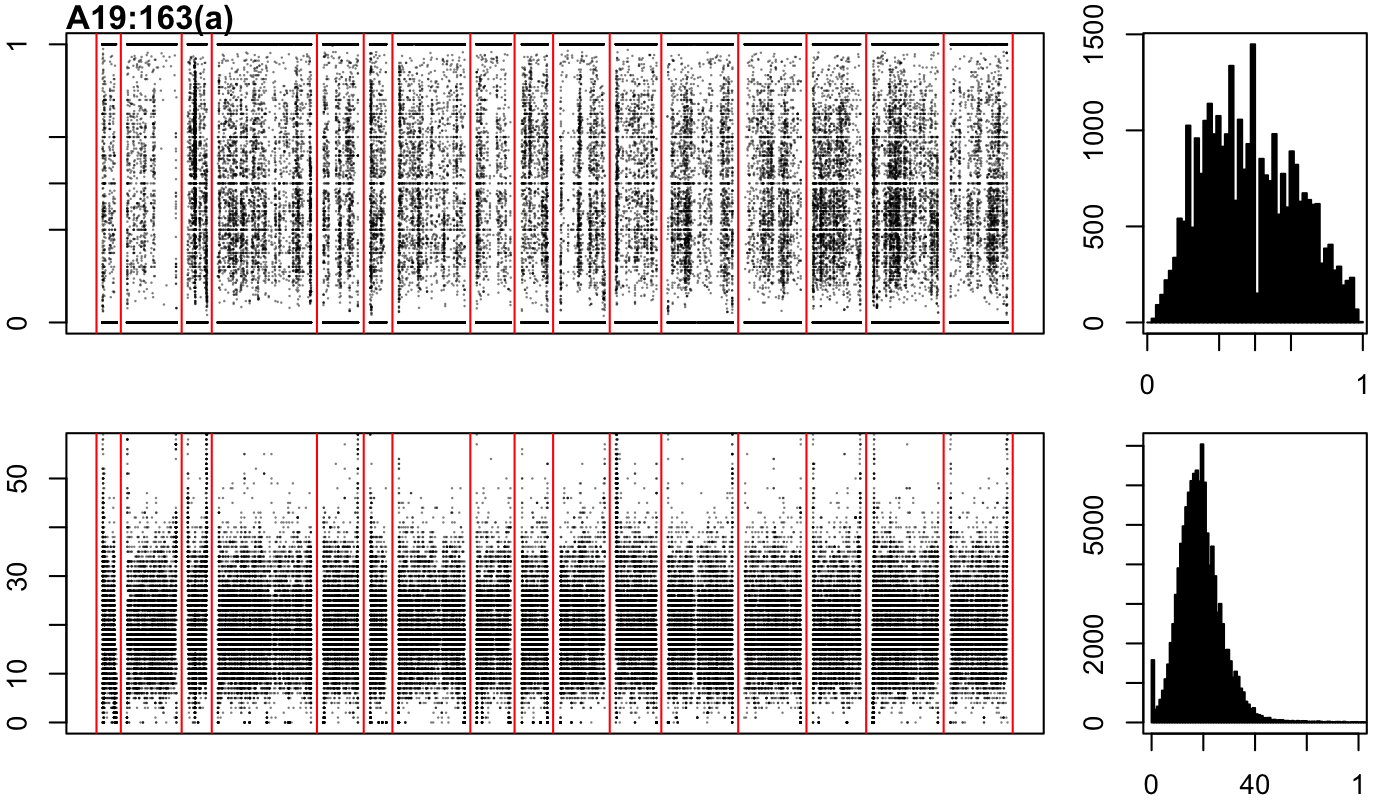

### A19-163-b.png

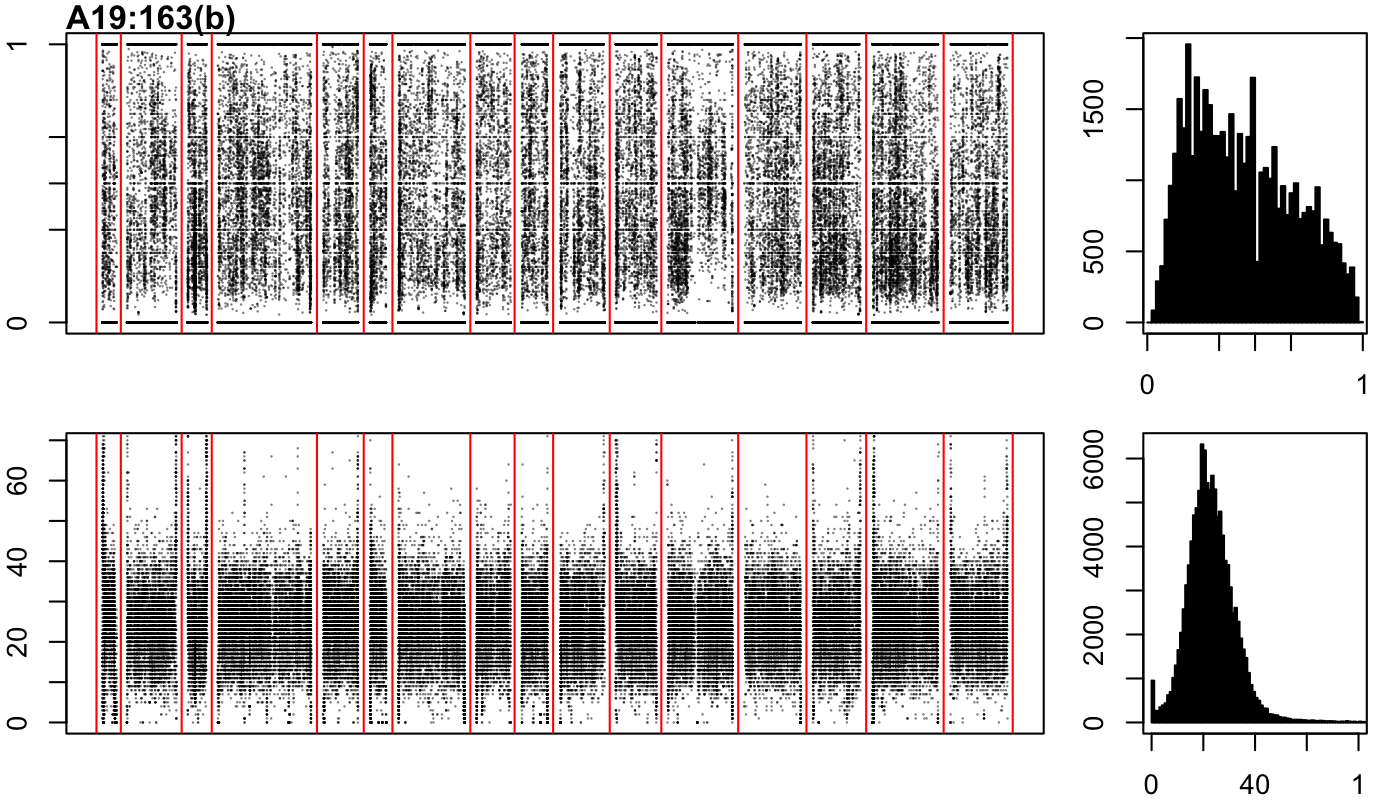

### A19-174-a.png

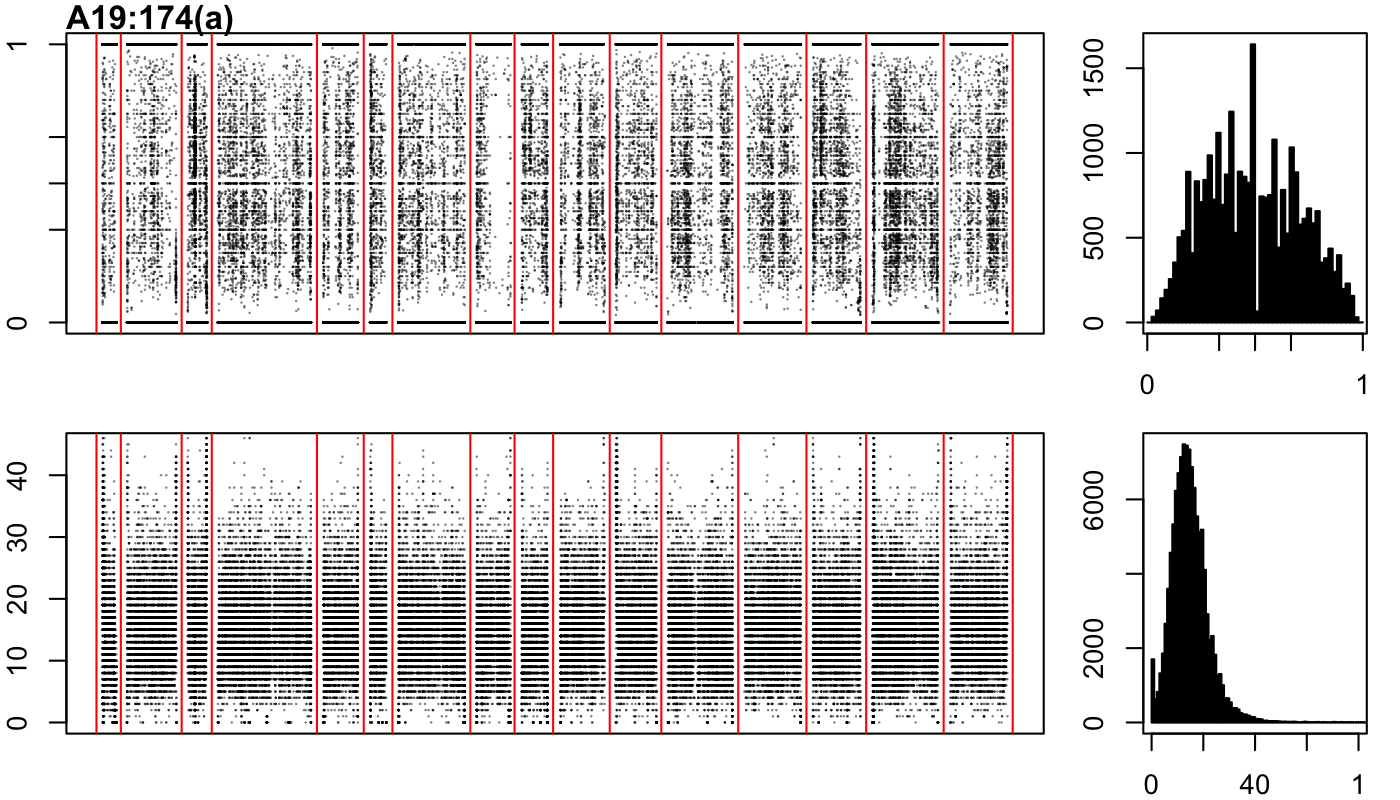

### A19-174-b.png

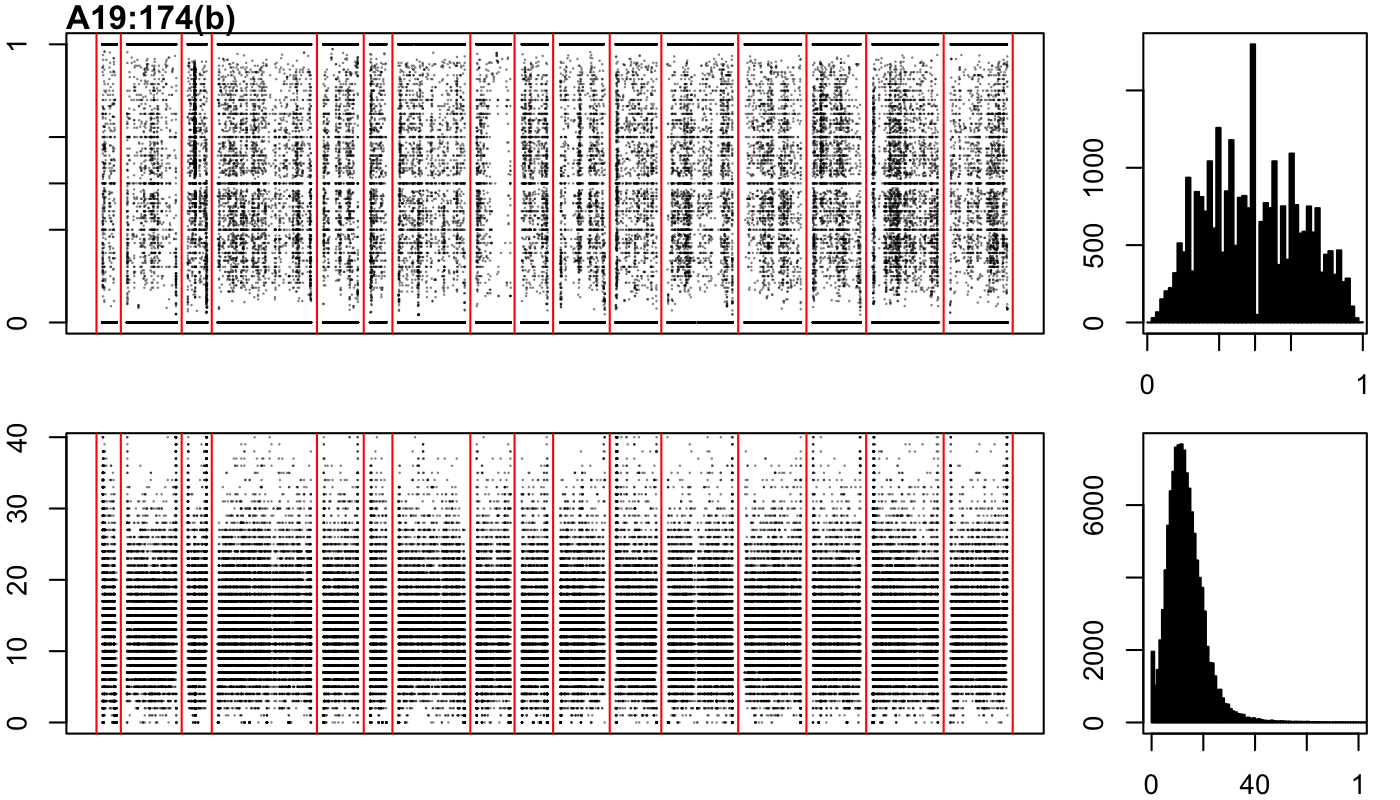

### A19-174-c.png

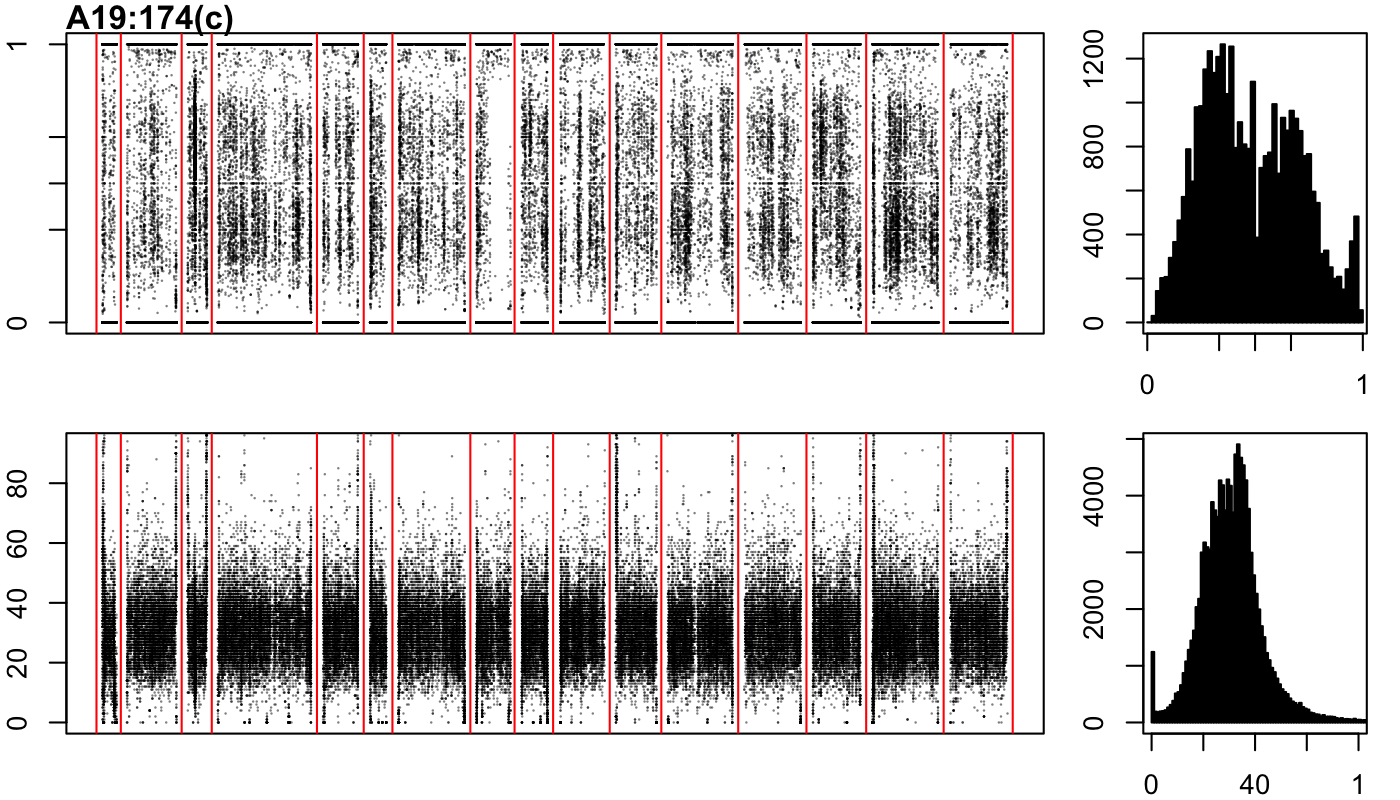

### B18-9-a.png

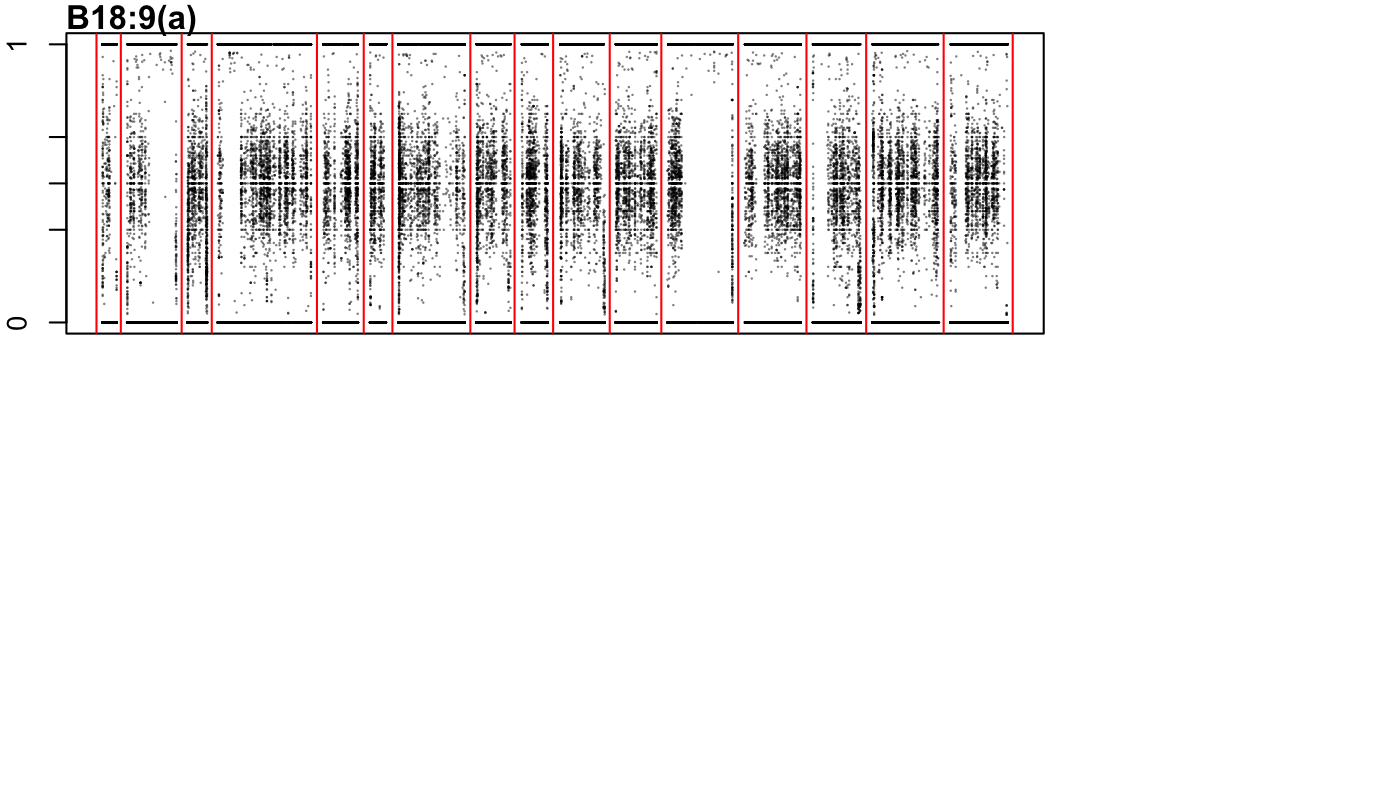

### B18-9-b.png

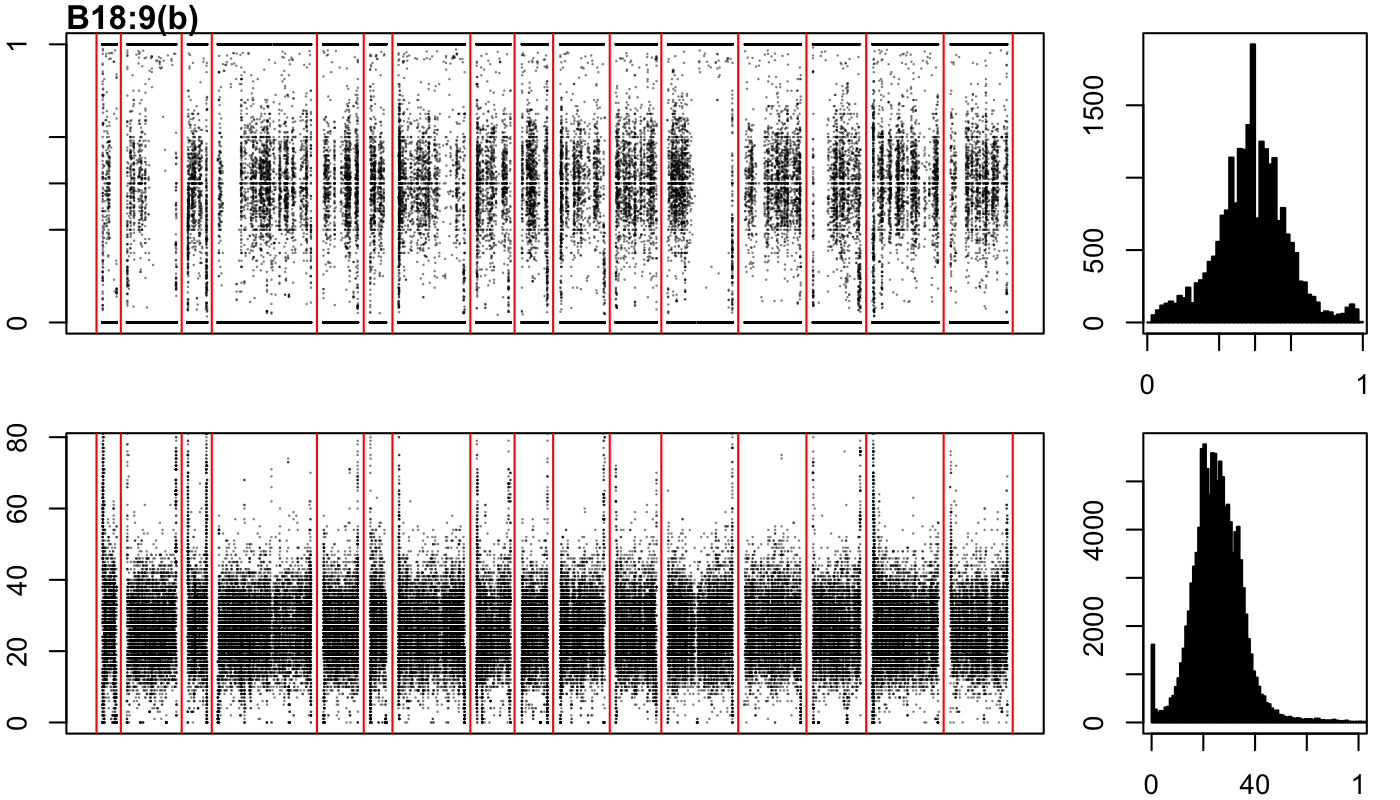

### B18-9-c.png

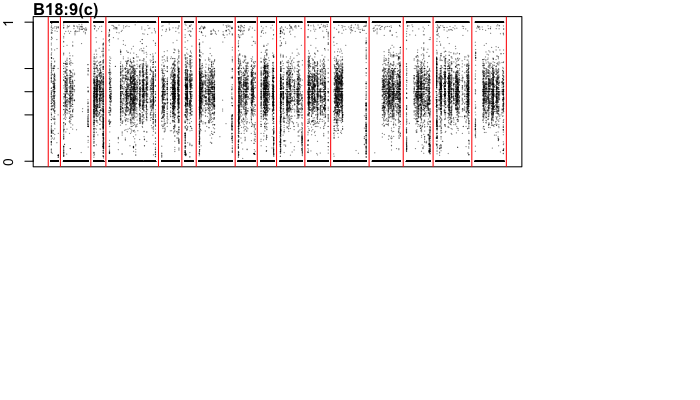

### B18-27-a.png

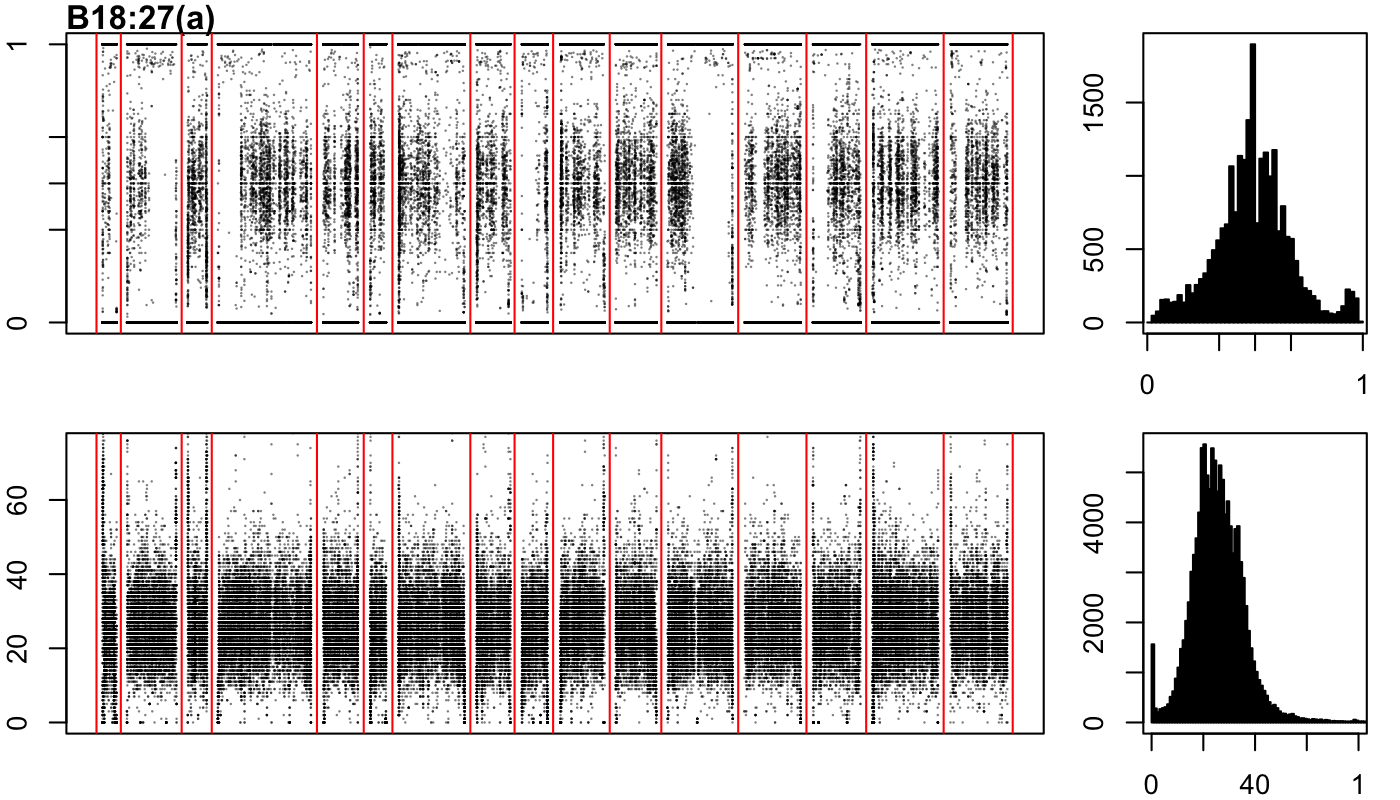

### B18-57-a.png

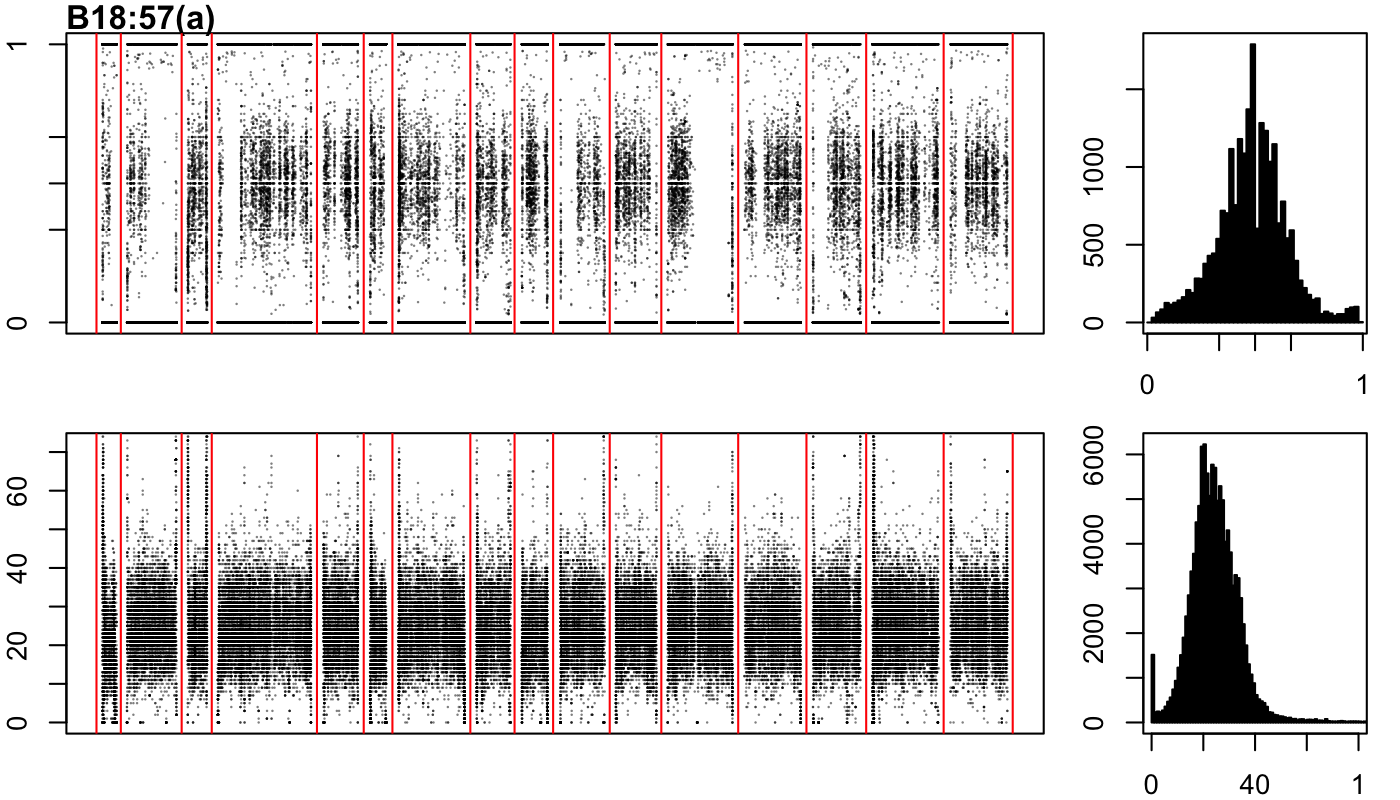

### B18-57-b.png

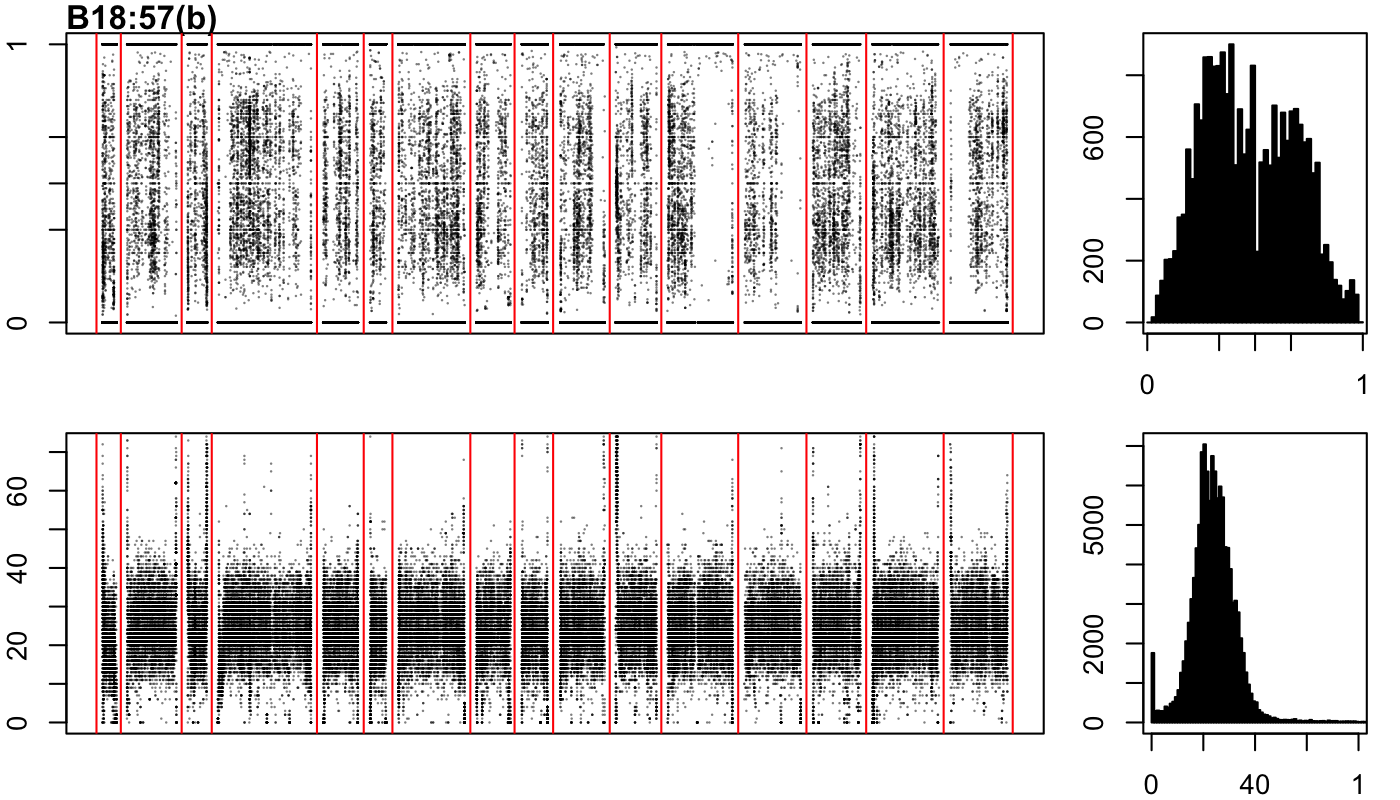

### B18-57-c.png

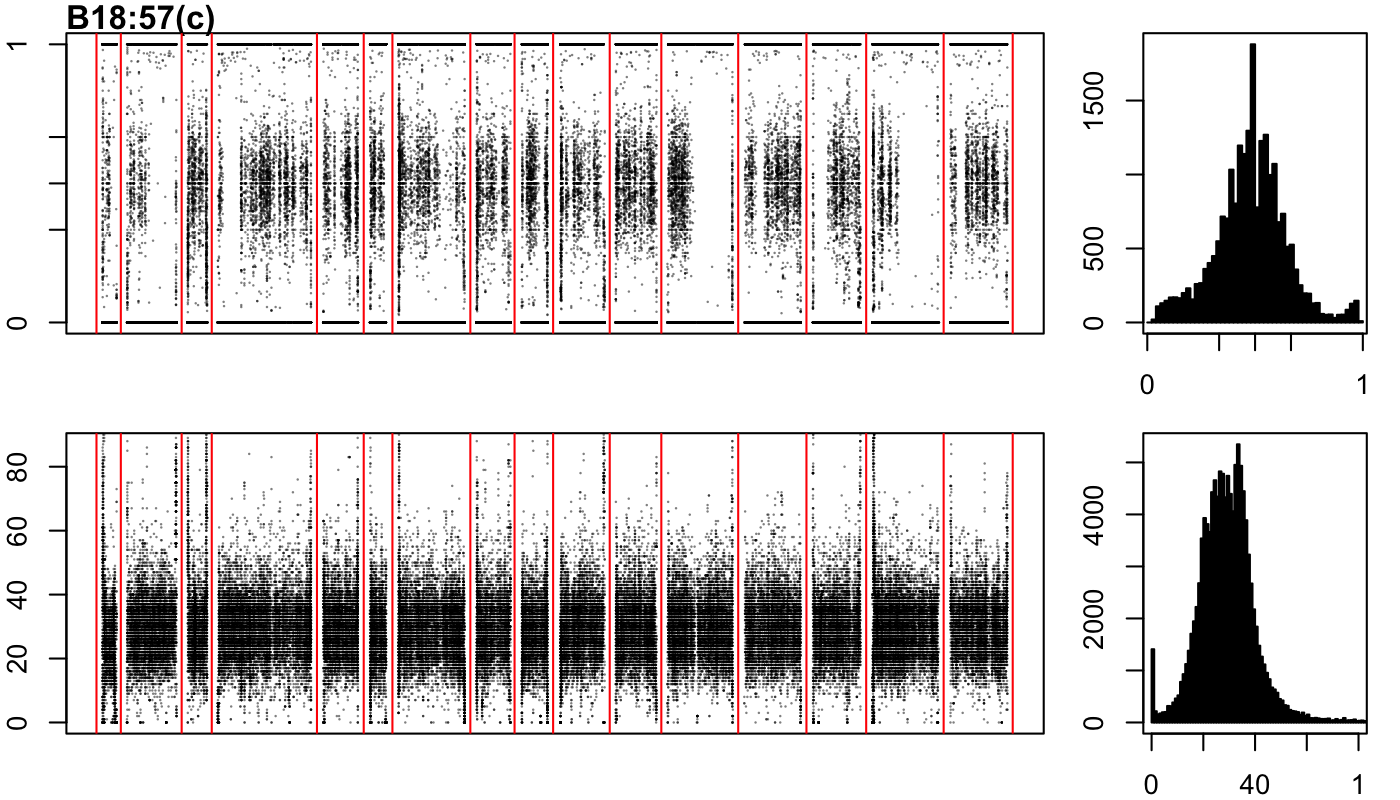

### B18-77-a.png

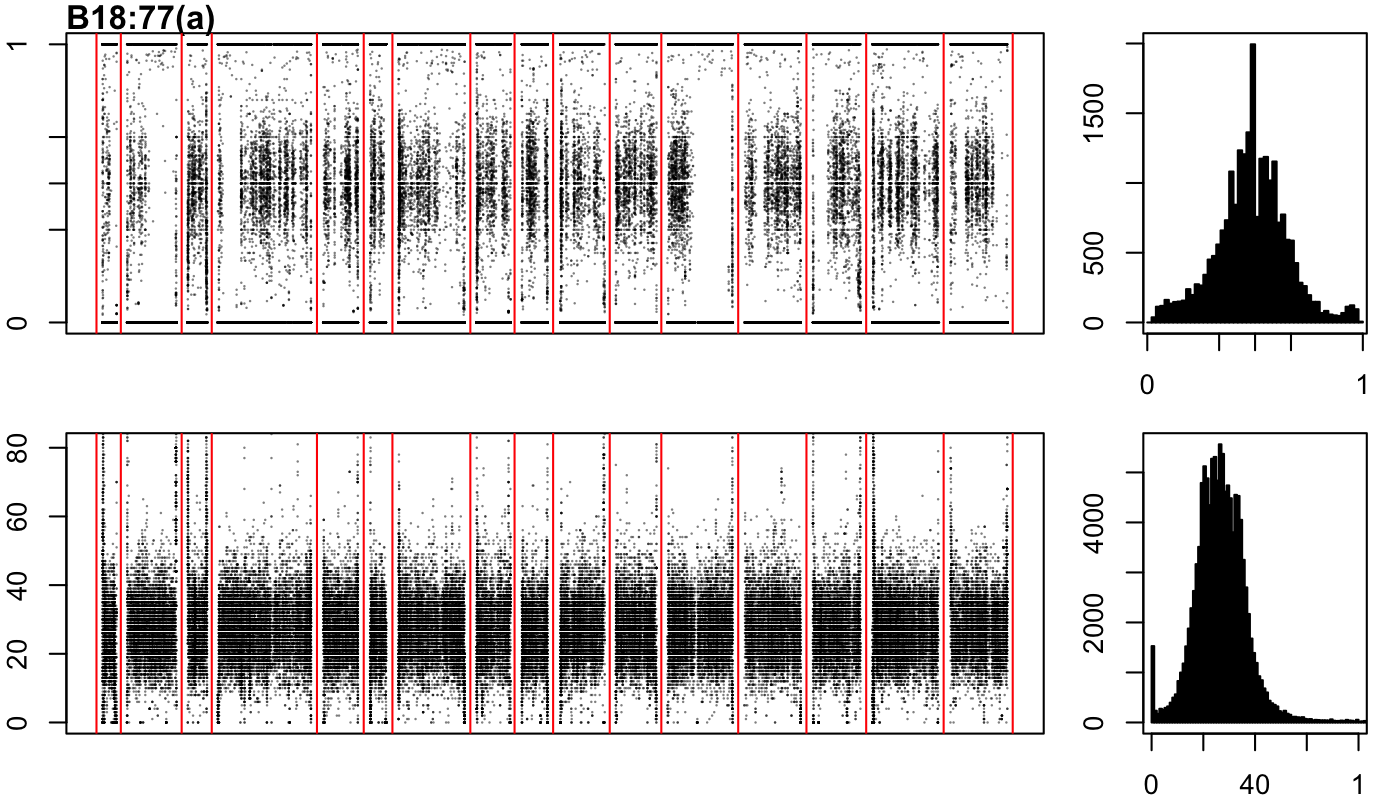

### B18-77-b.png

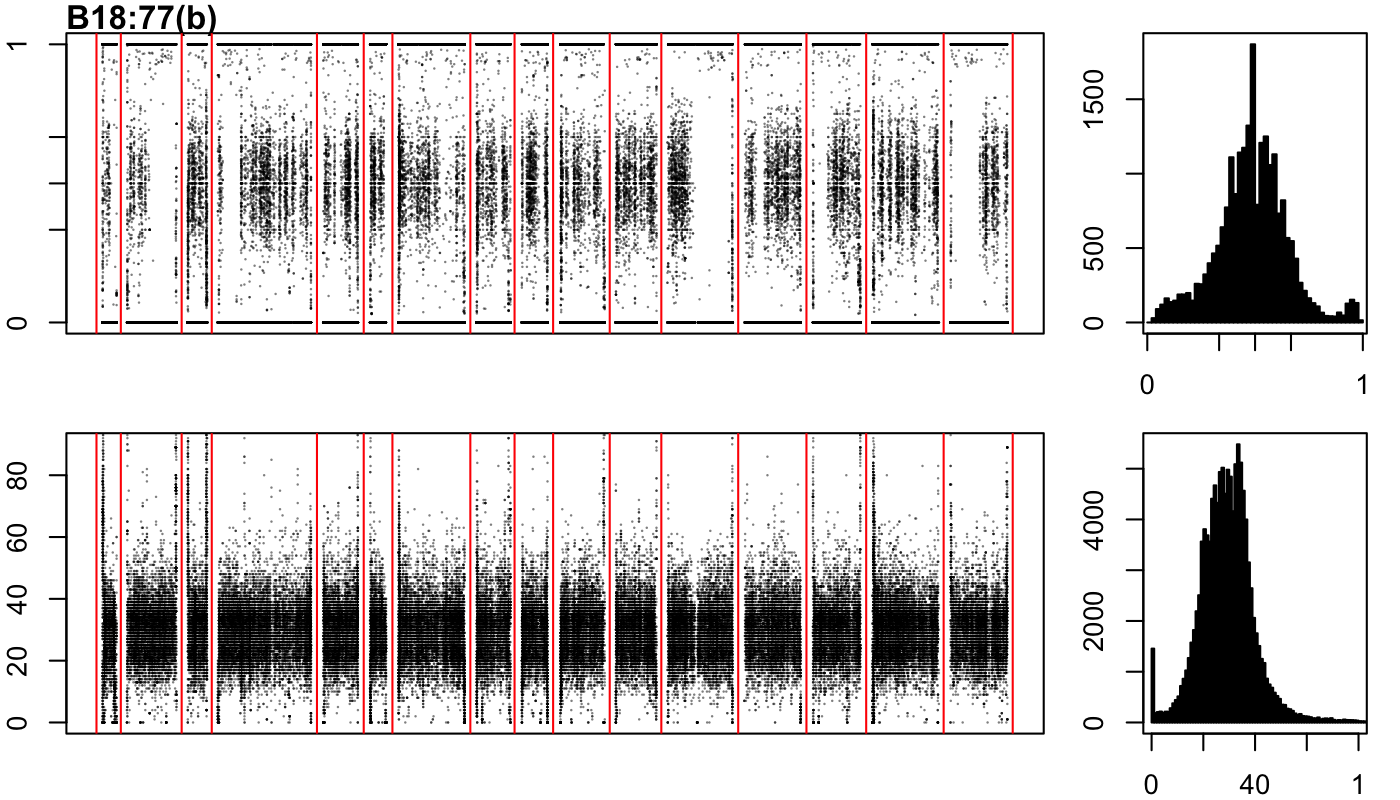

### B18-77-c.png

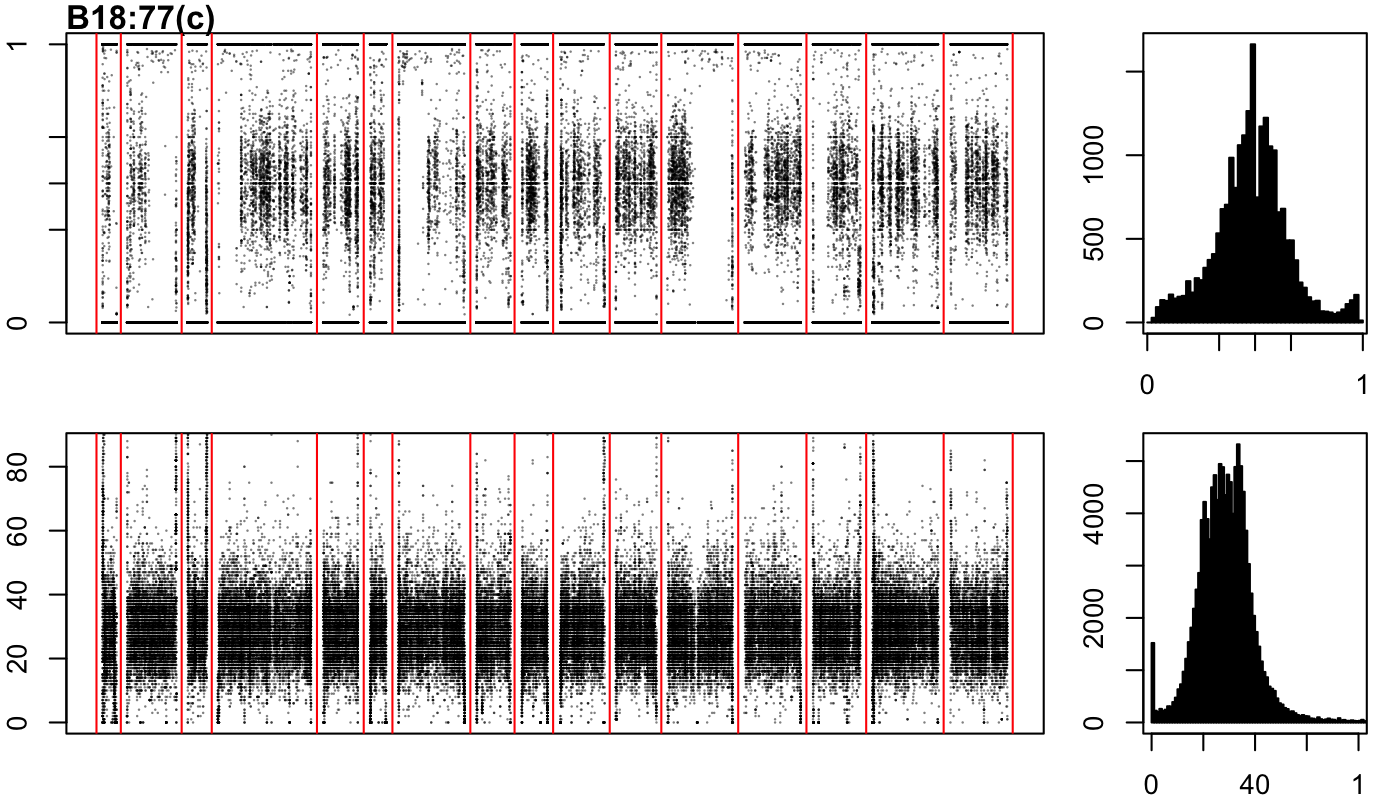

### B18-147-a.png

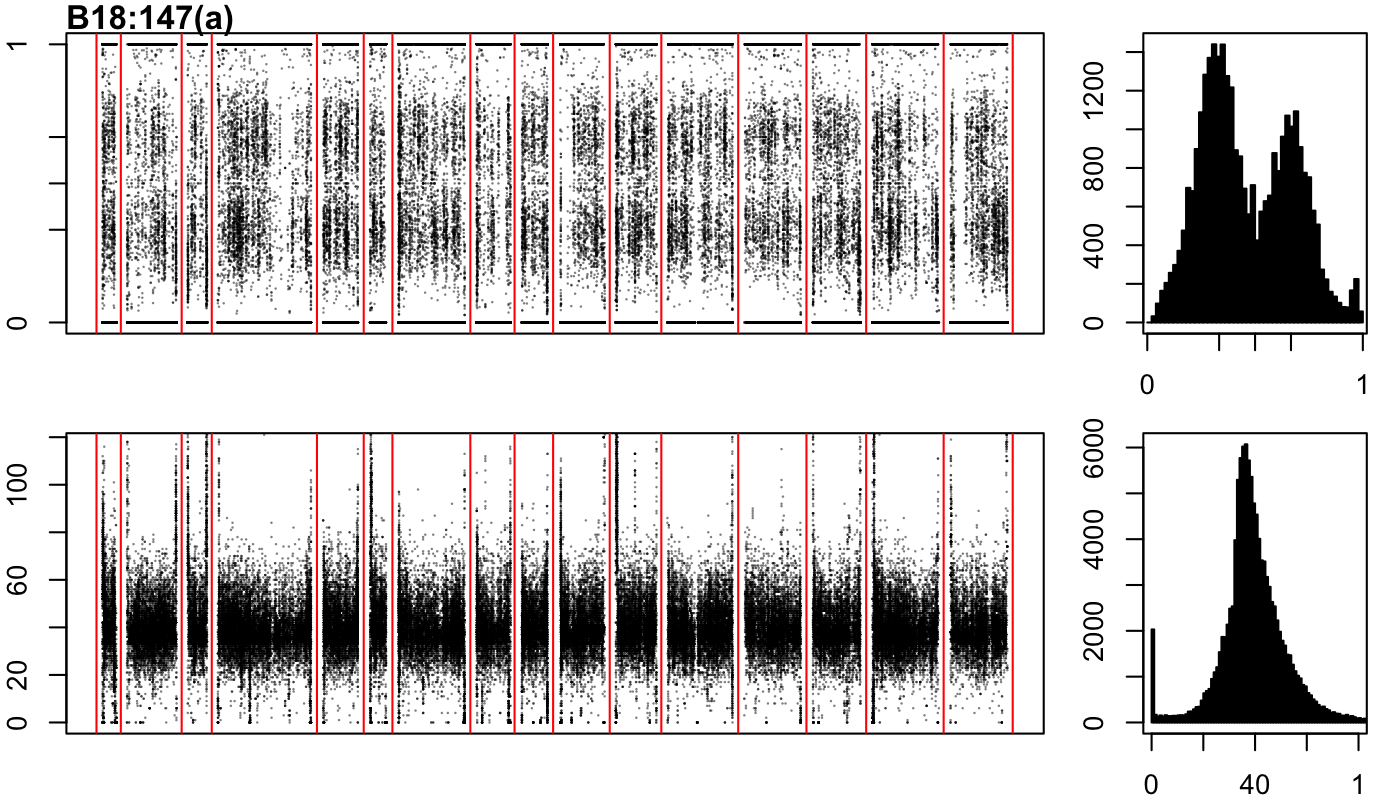

### B18-147-b.png

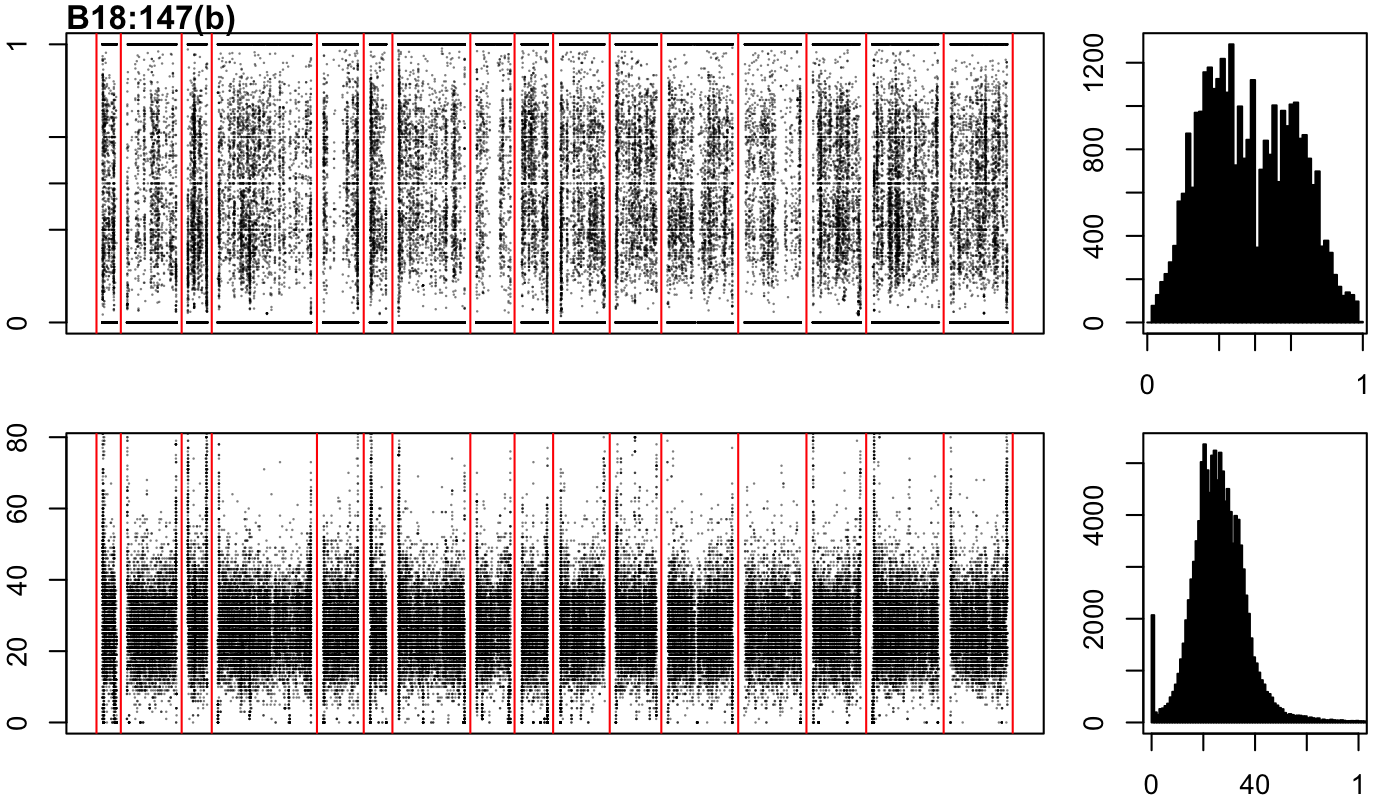

### B18-147-c.png

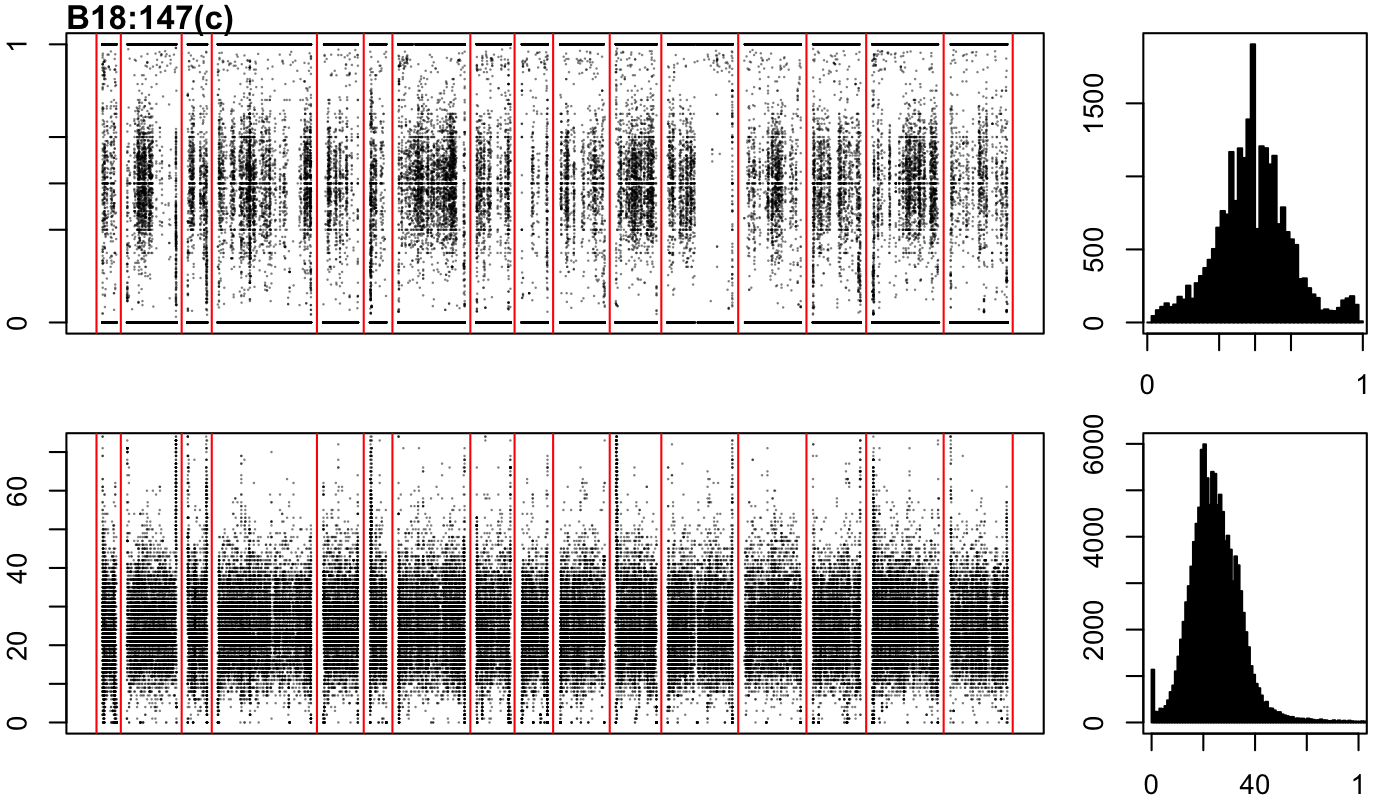

### B18-167-a.png

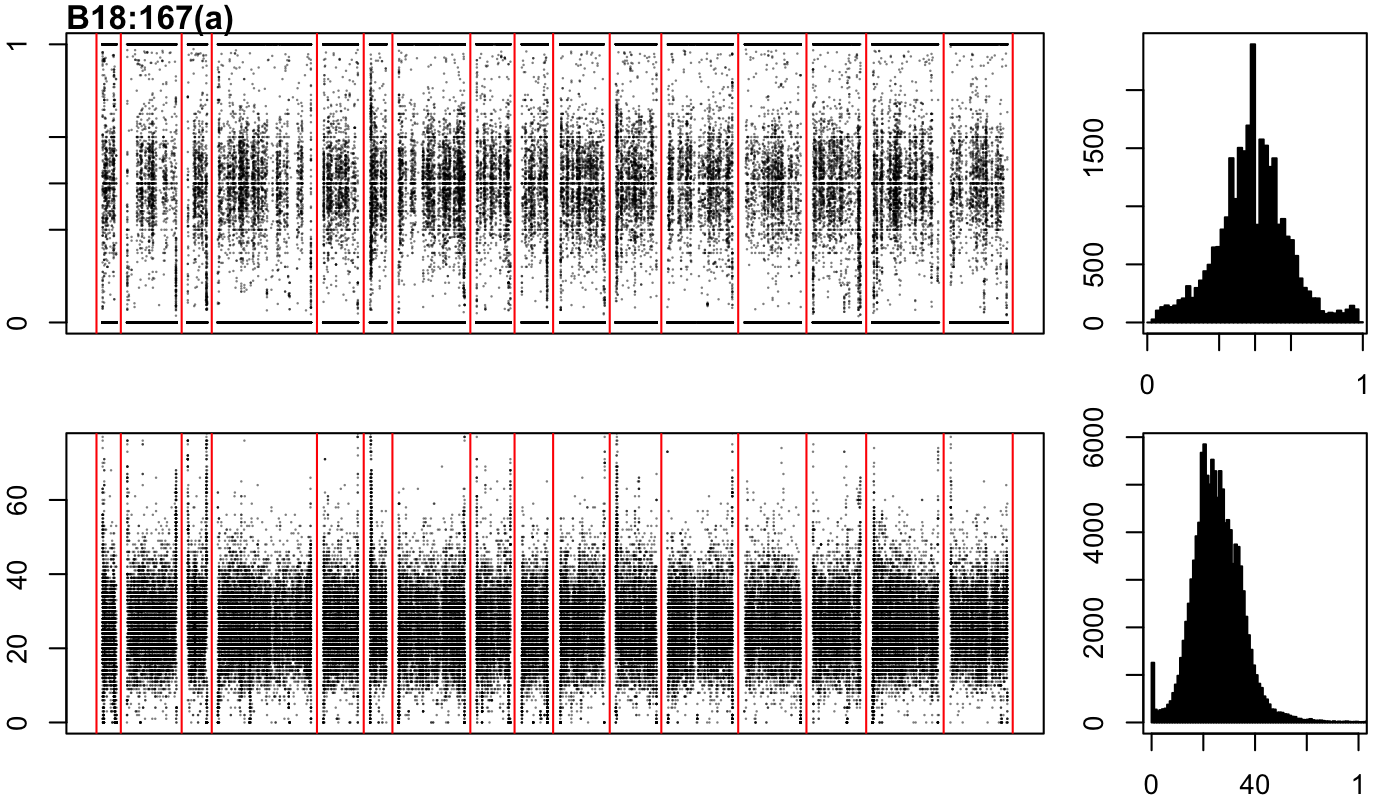

### B18-167-b.png

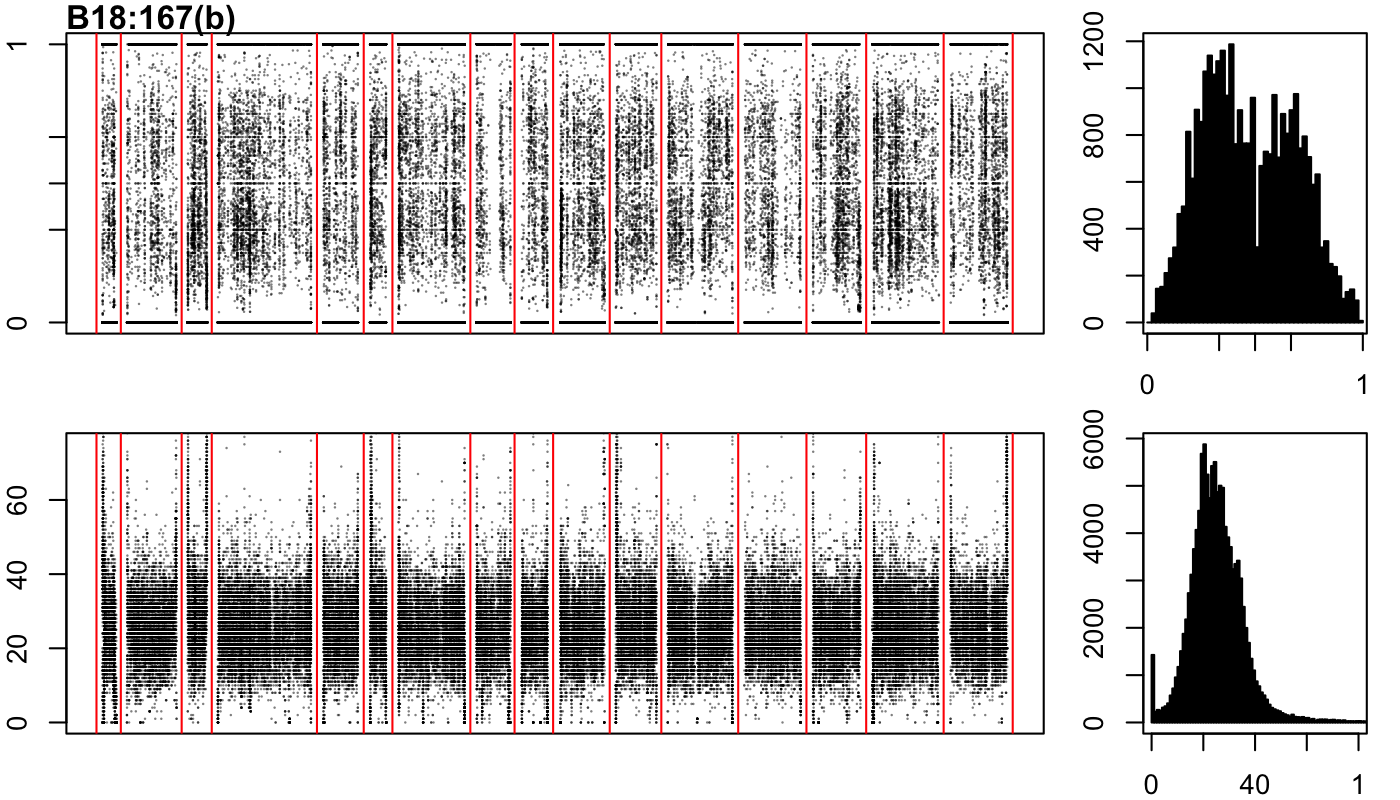

### B18-167-c.png

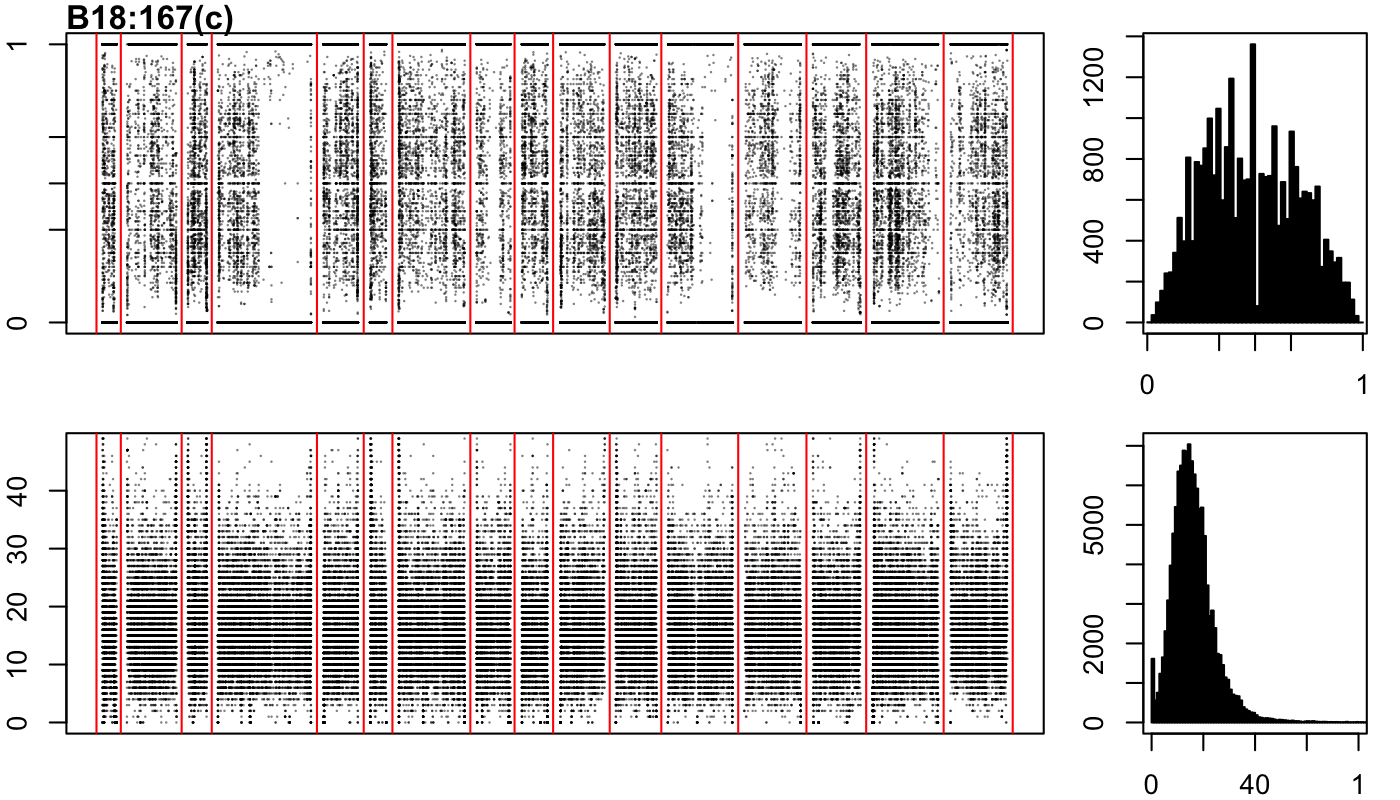

### B18-201-a.png

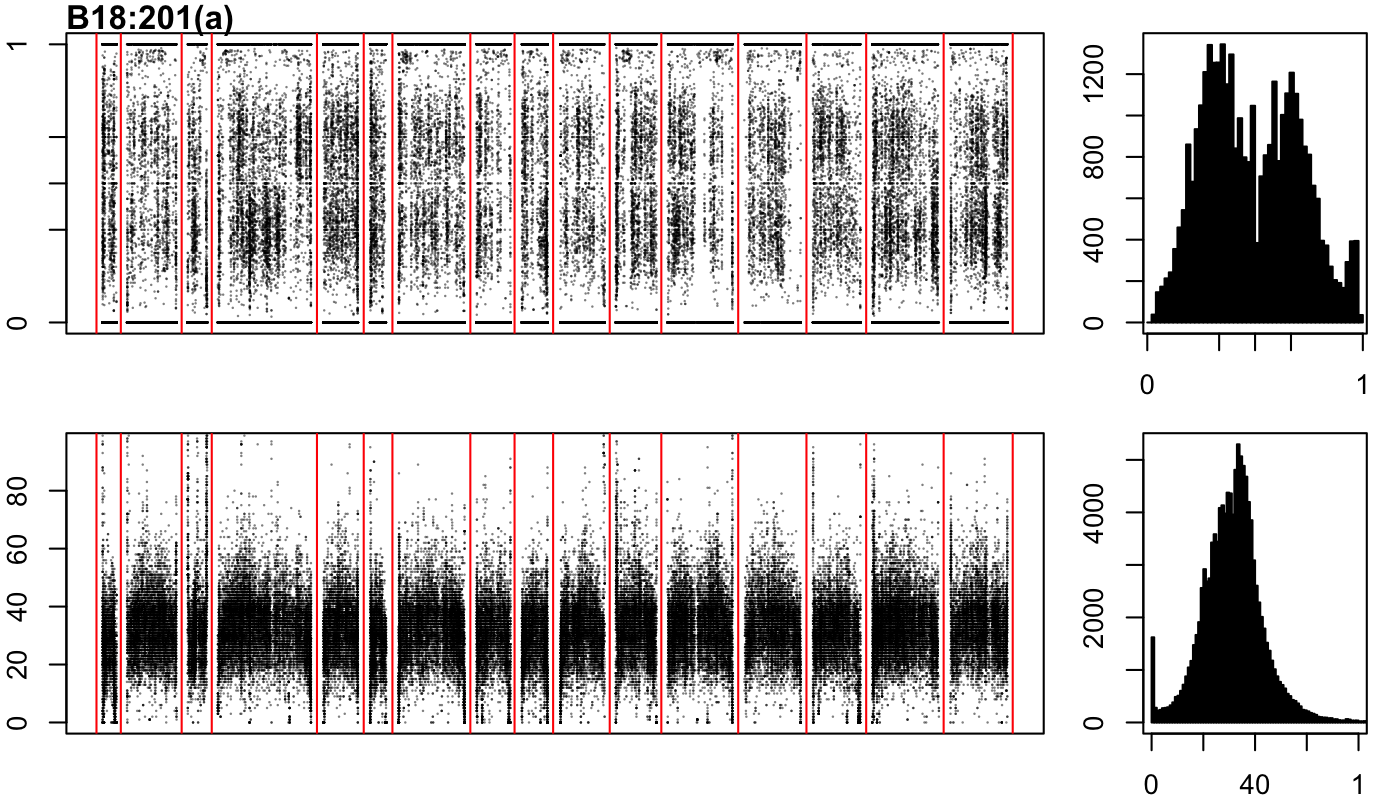

### B18-201-b.png

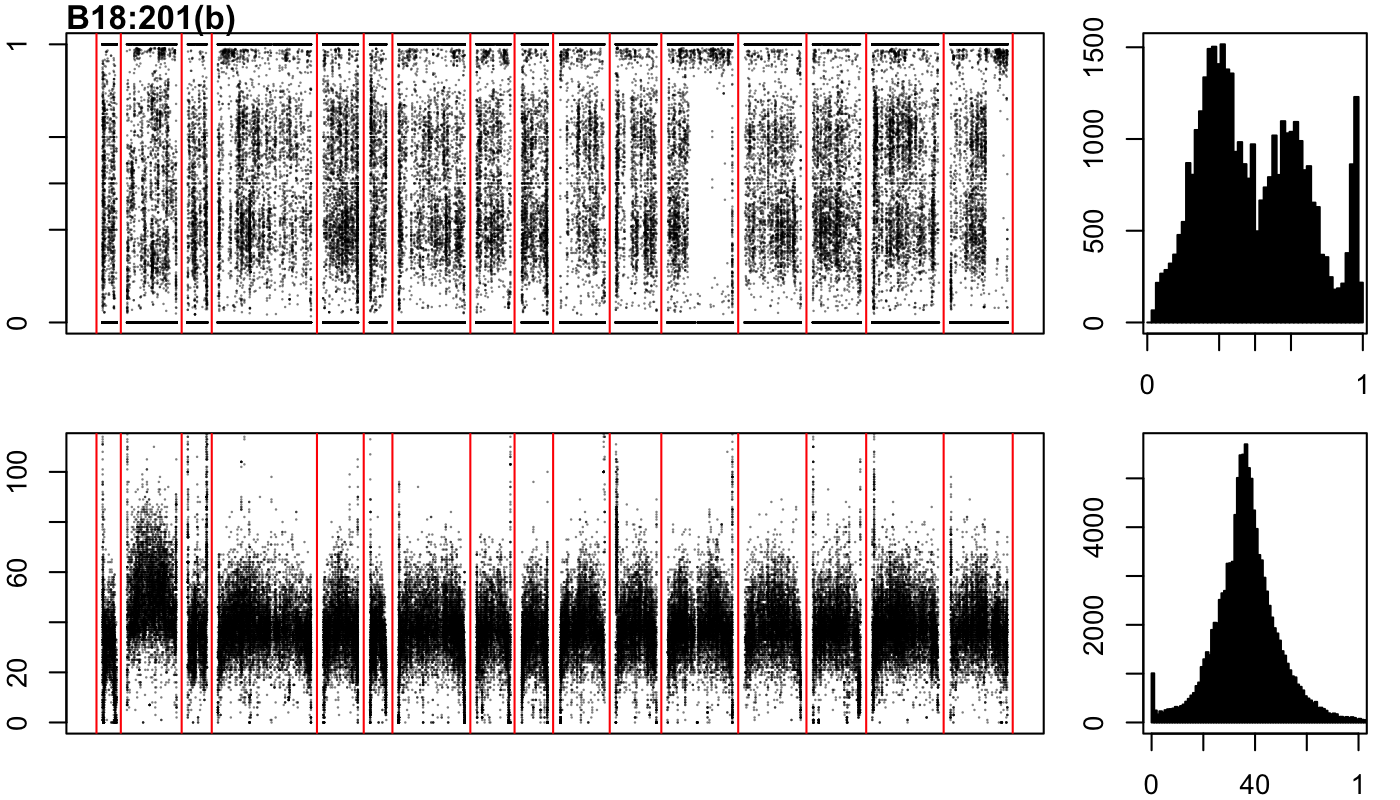

### B18-203-a.png

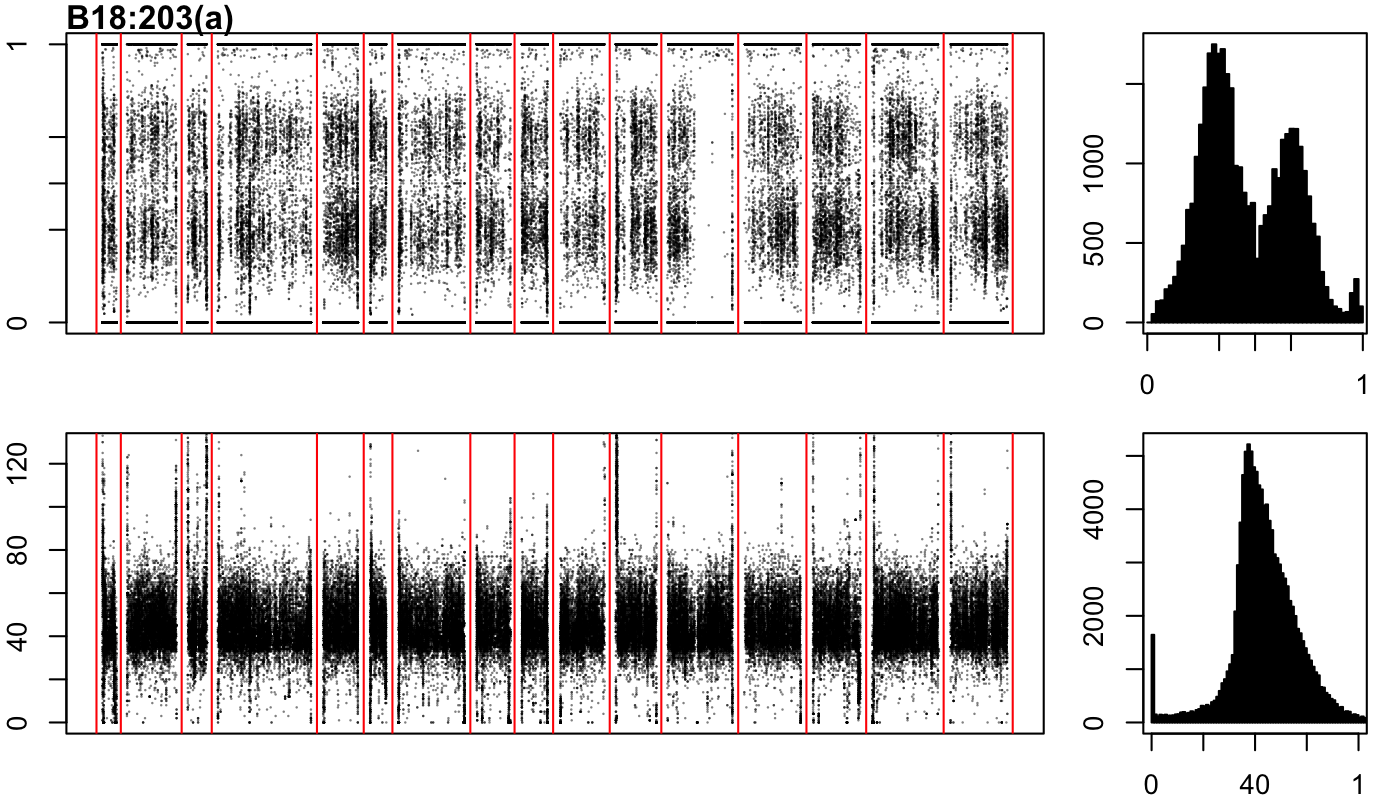

### B18-203-b.png

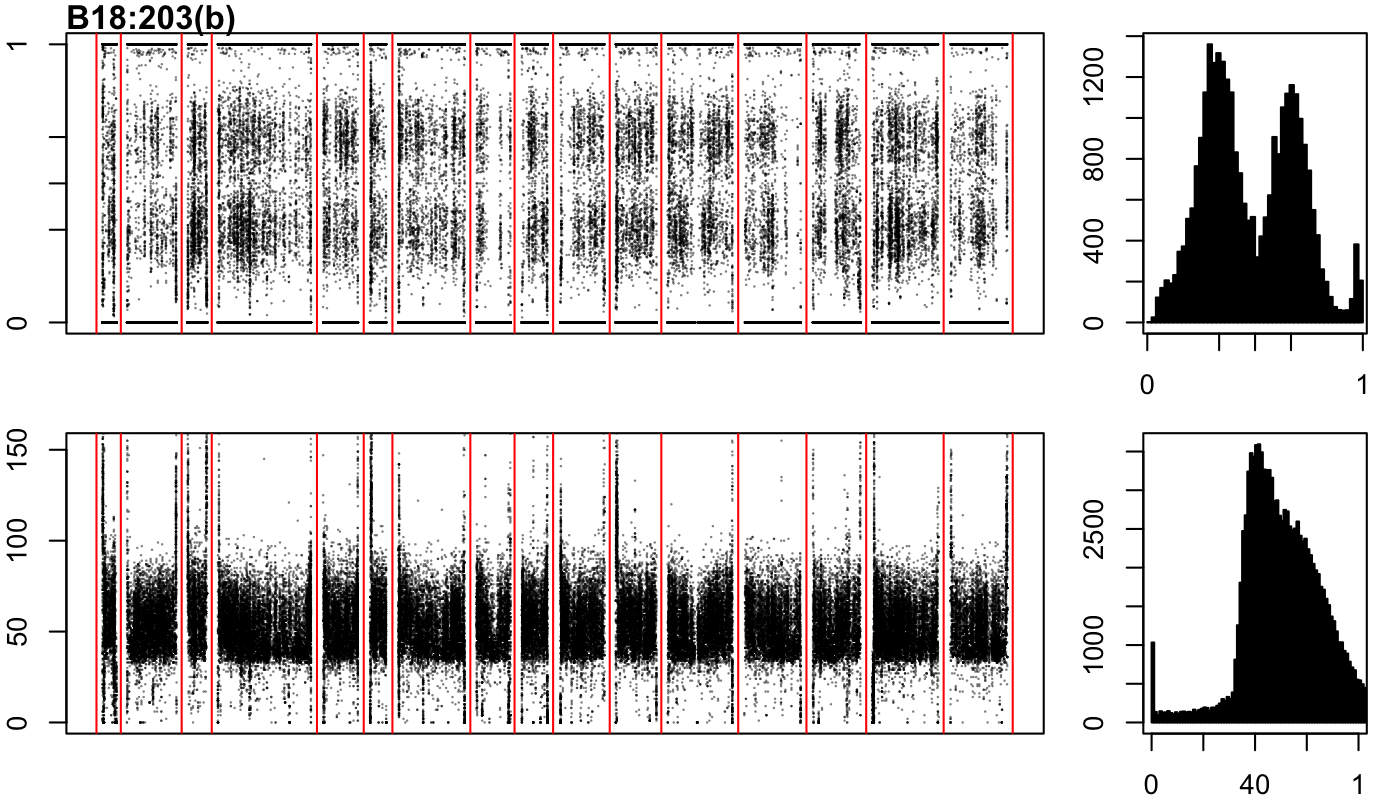

### B18-203-c.png

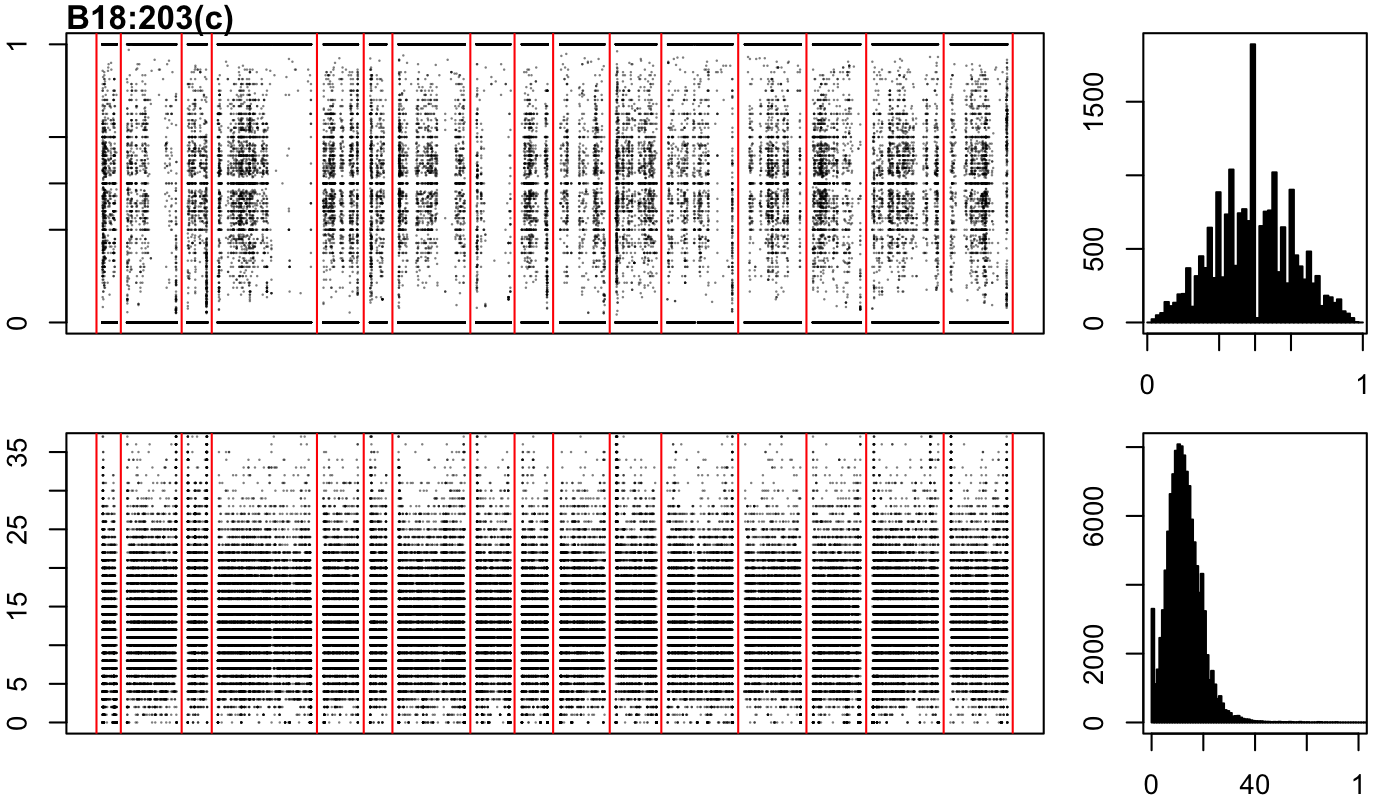

### B18:27(a).png

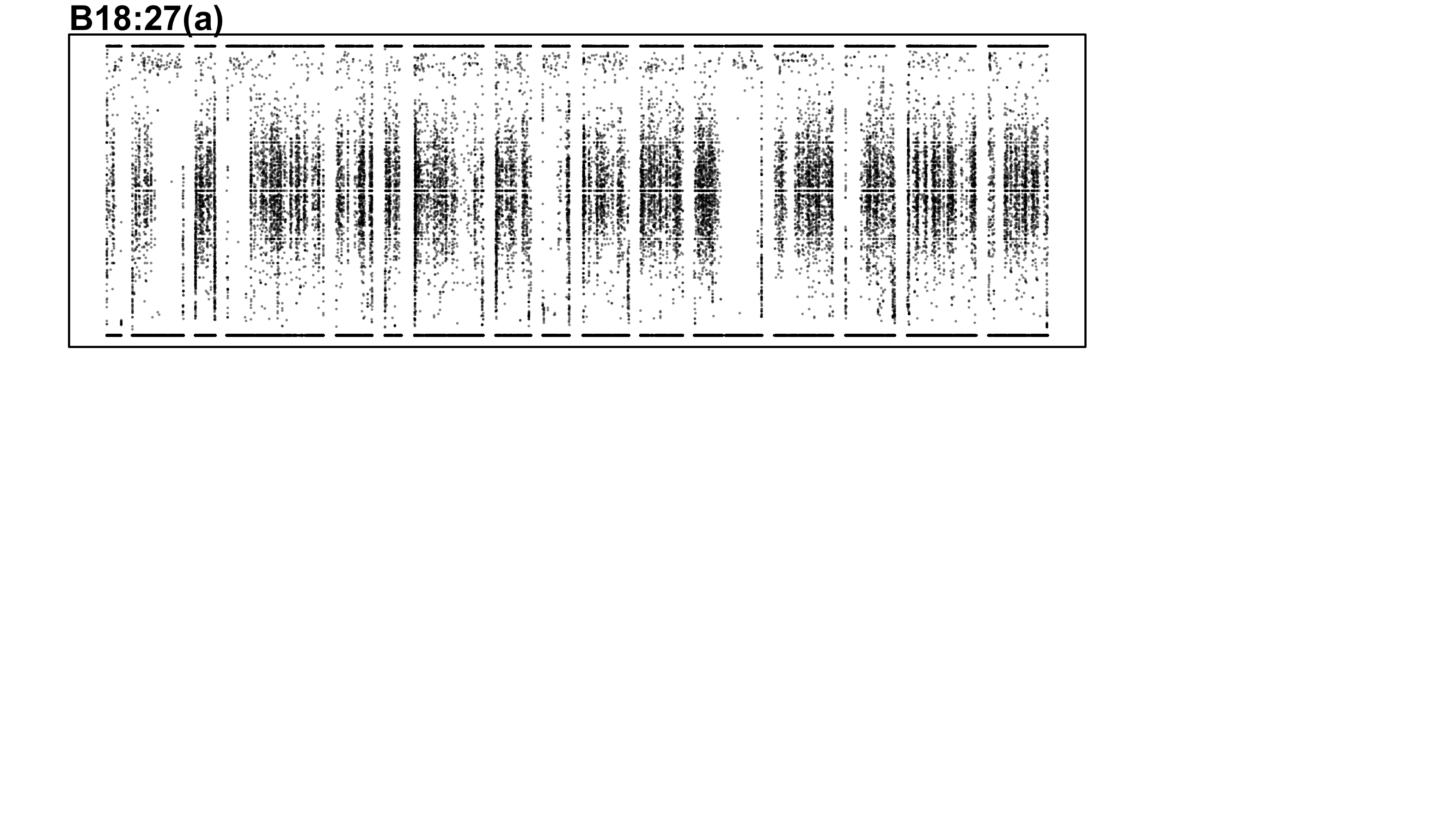

### B19-13-a.png

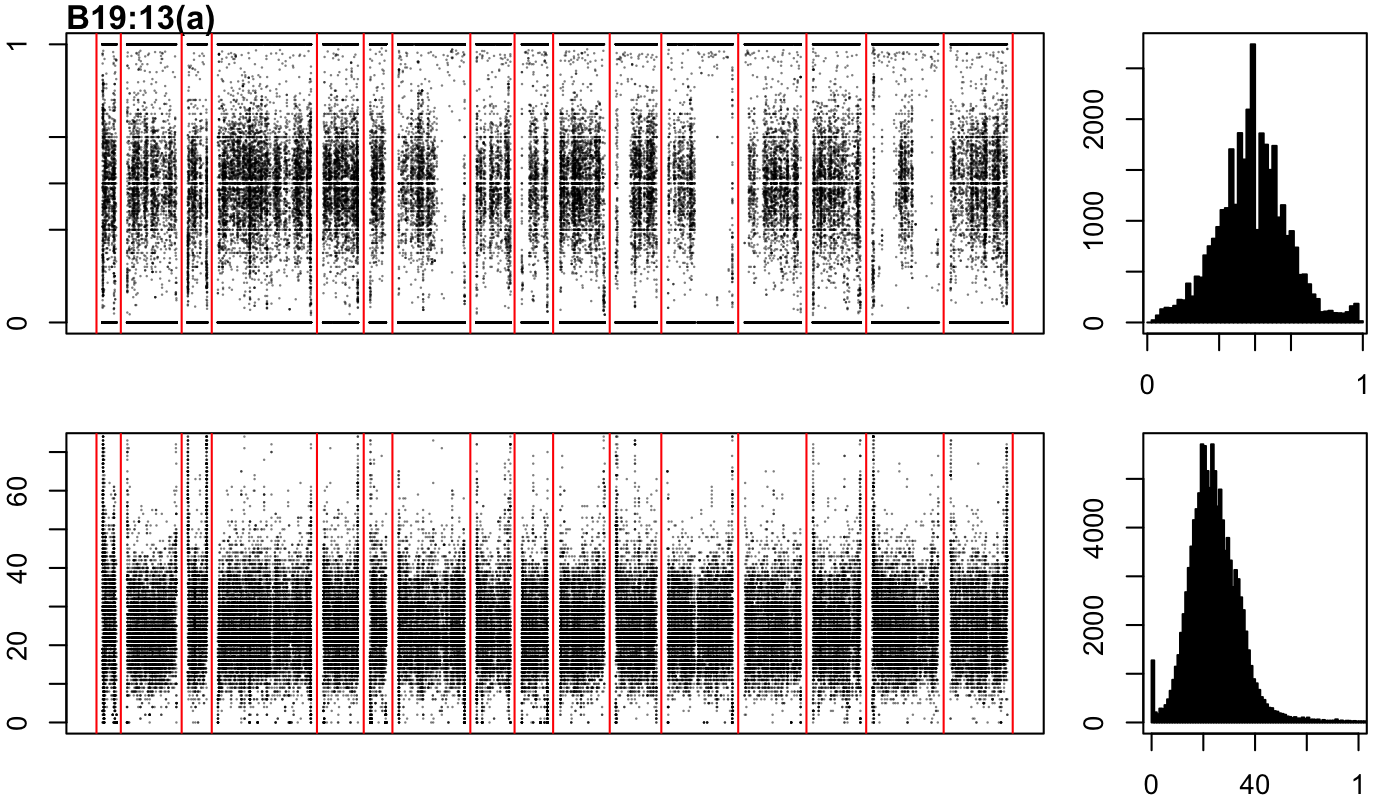

### B19-13-b.png

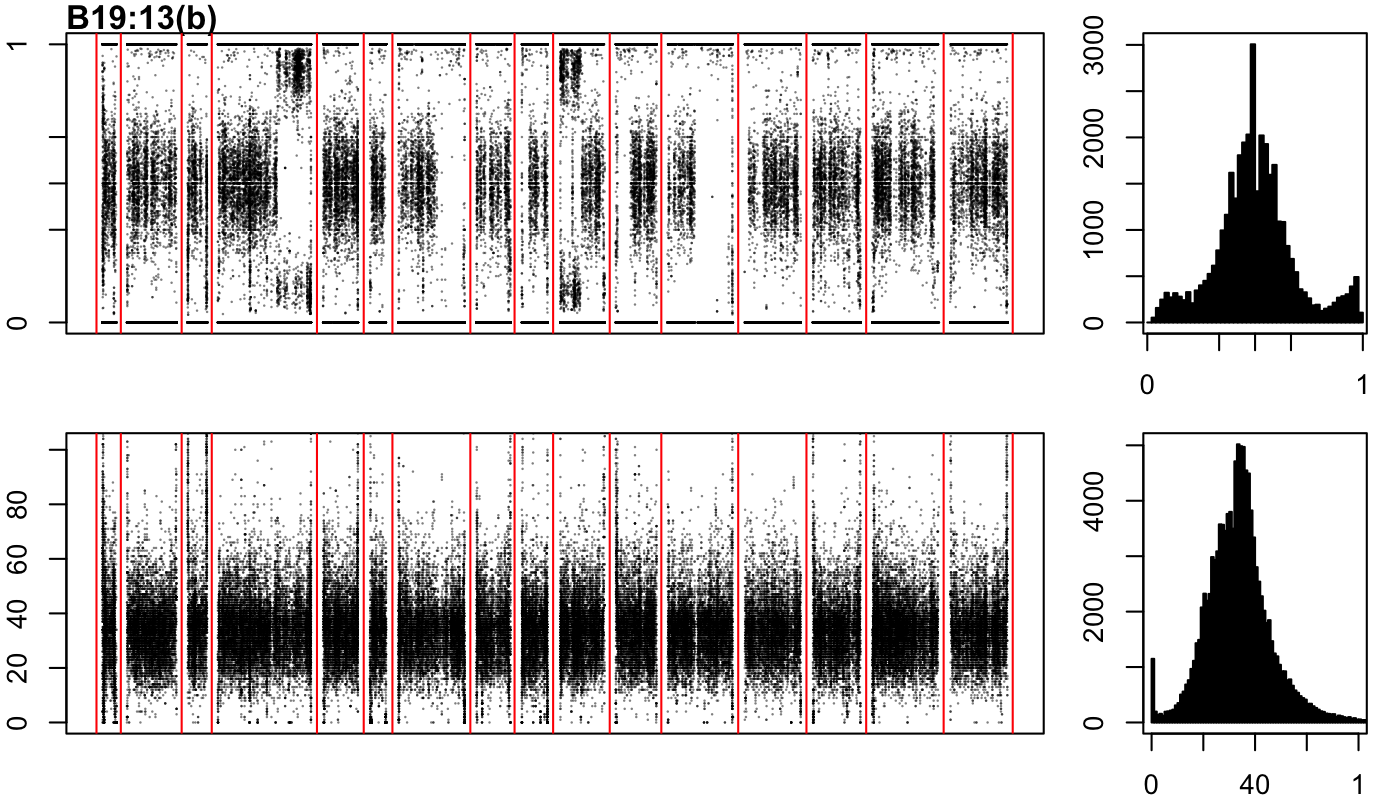

### B19-13-c.png

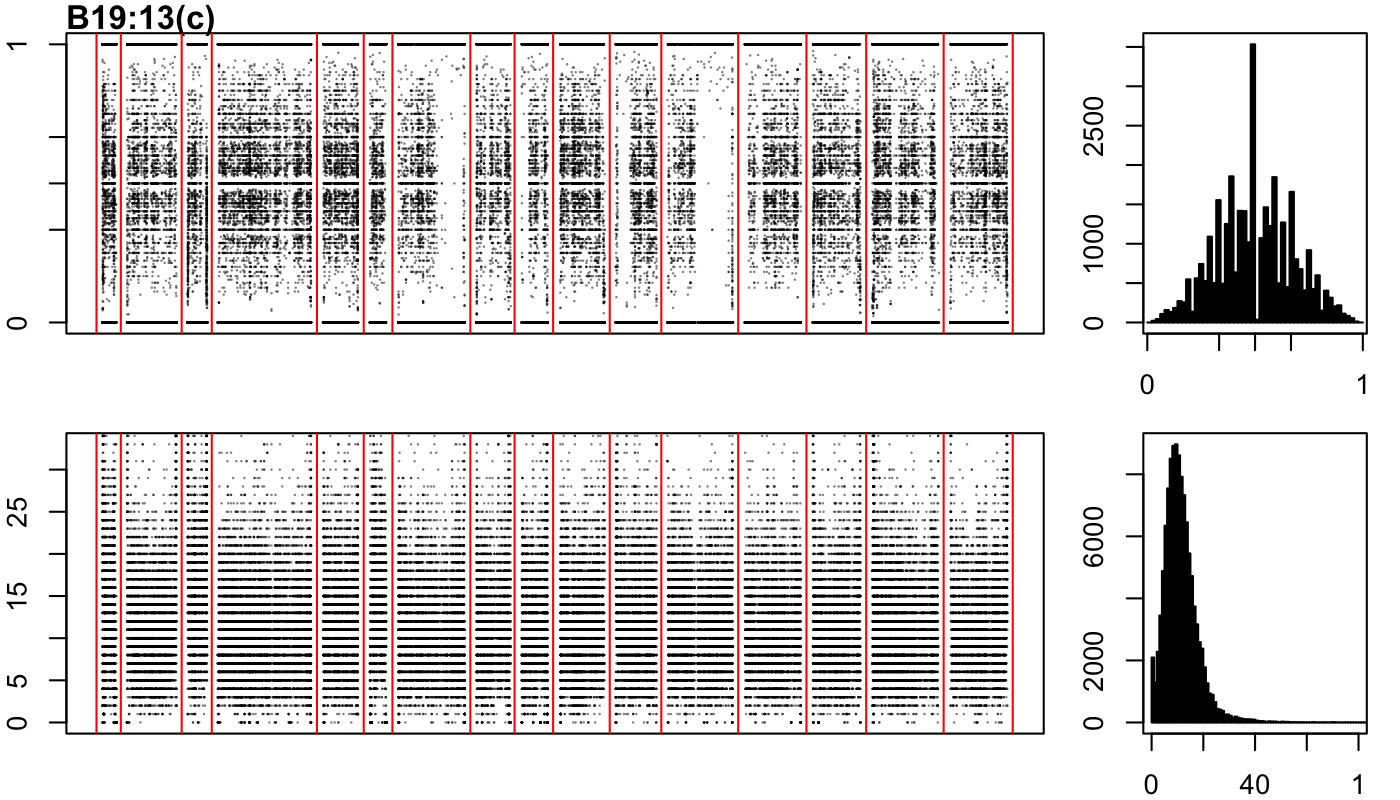

### B19-28-a.png

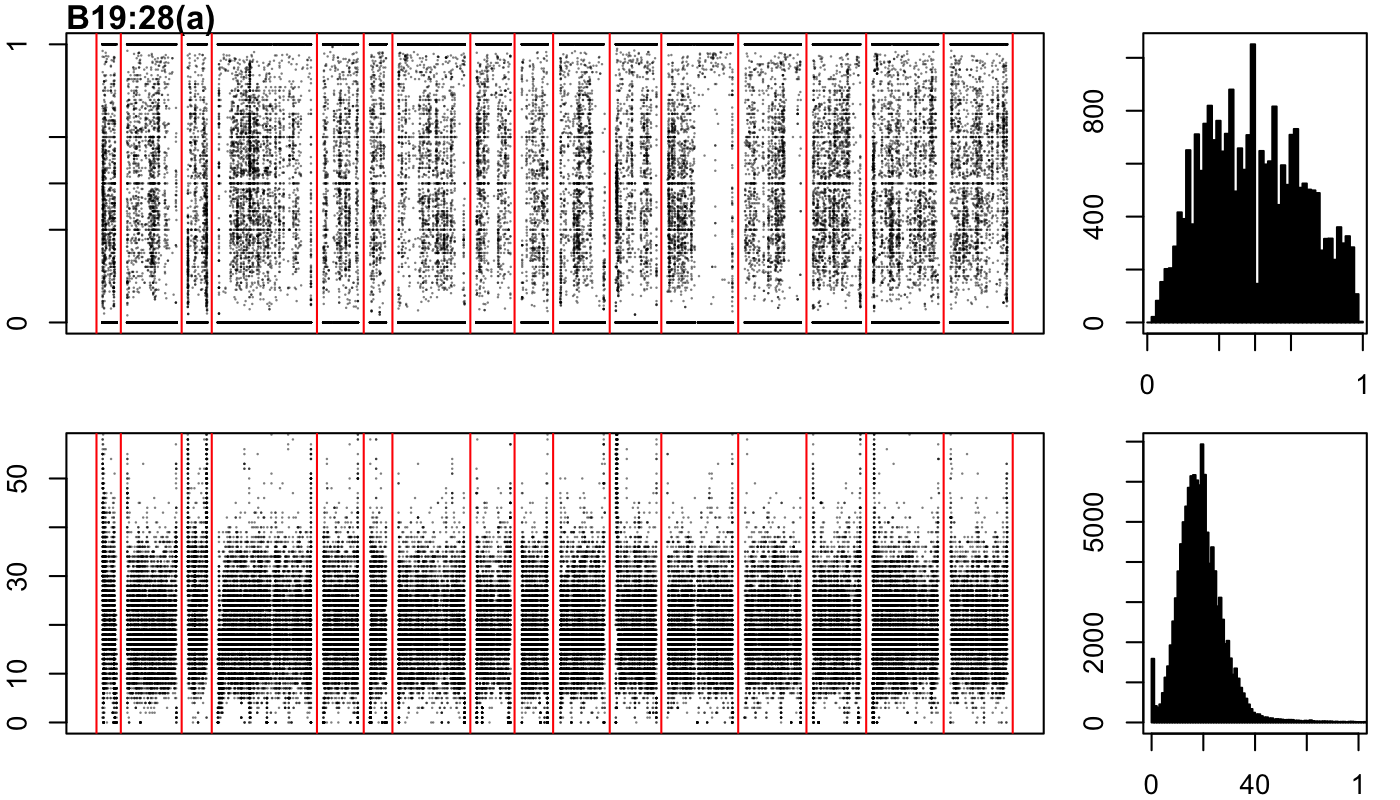

### B19-28-b.png

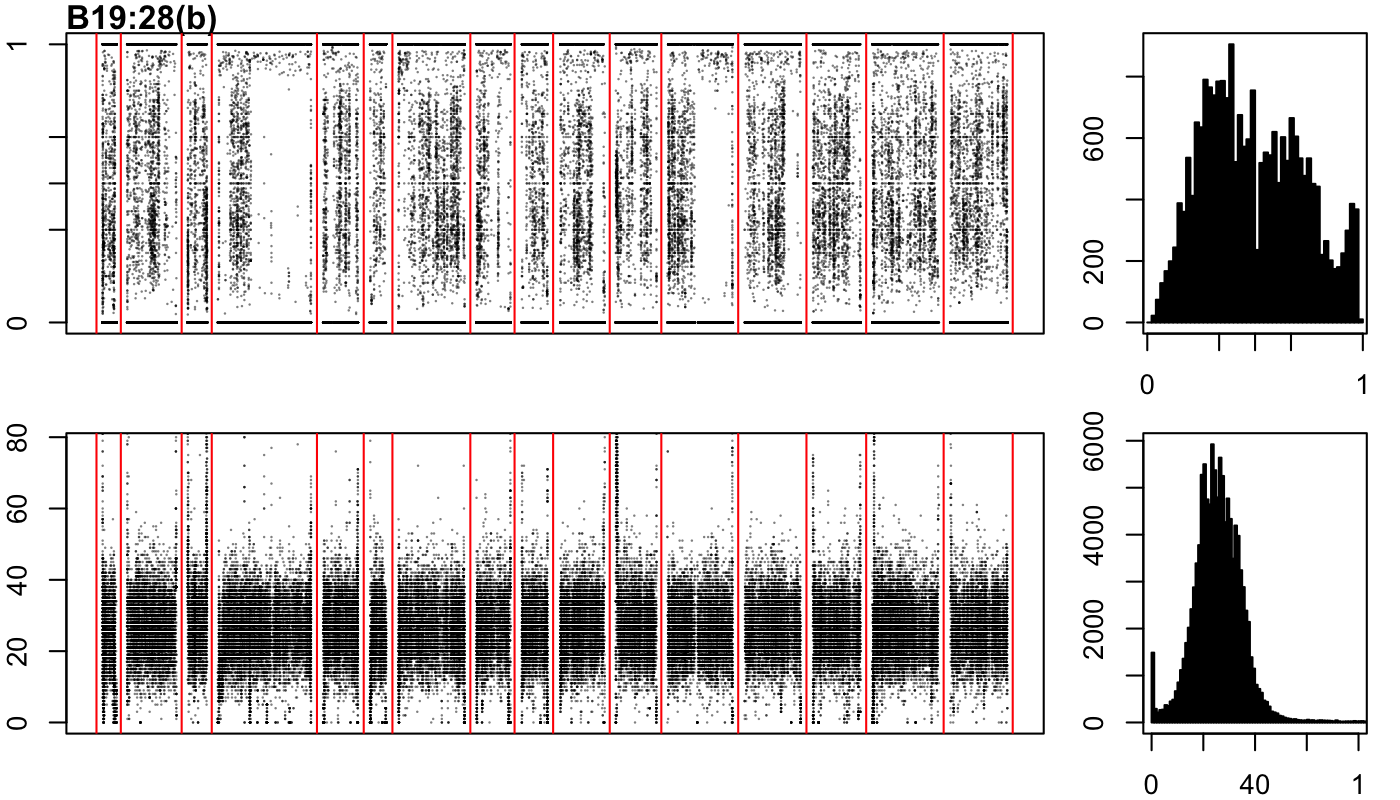

### B19-40-a.png

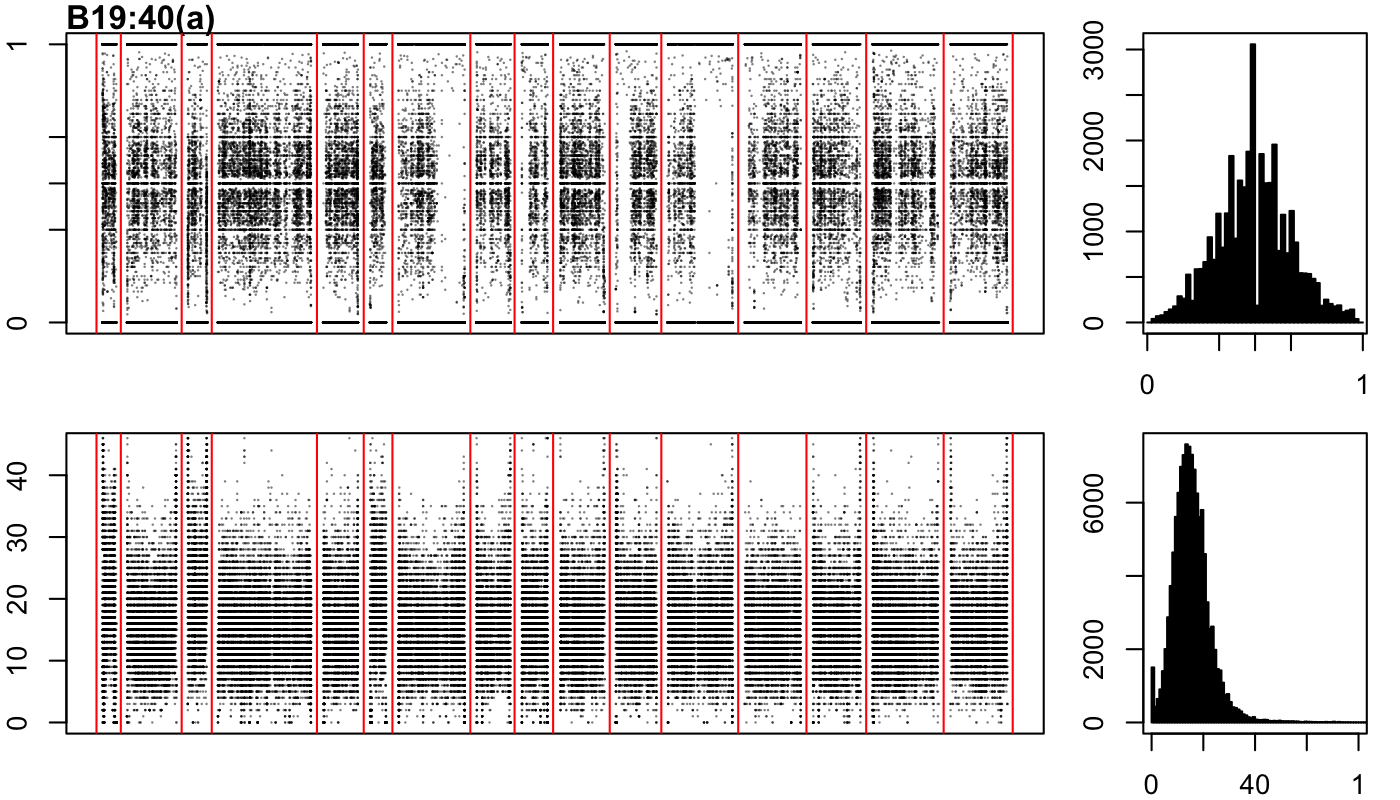

### B19-40-b.png

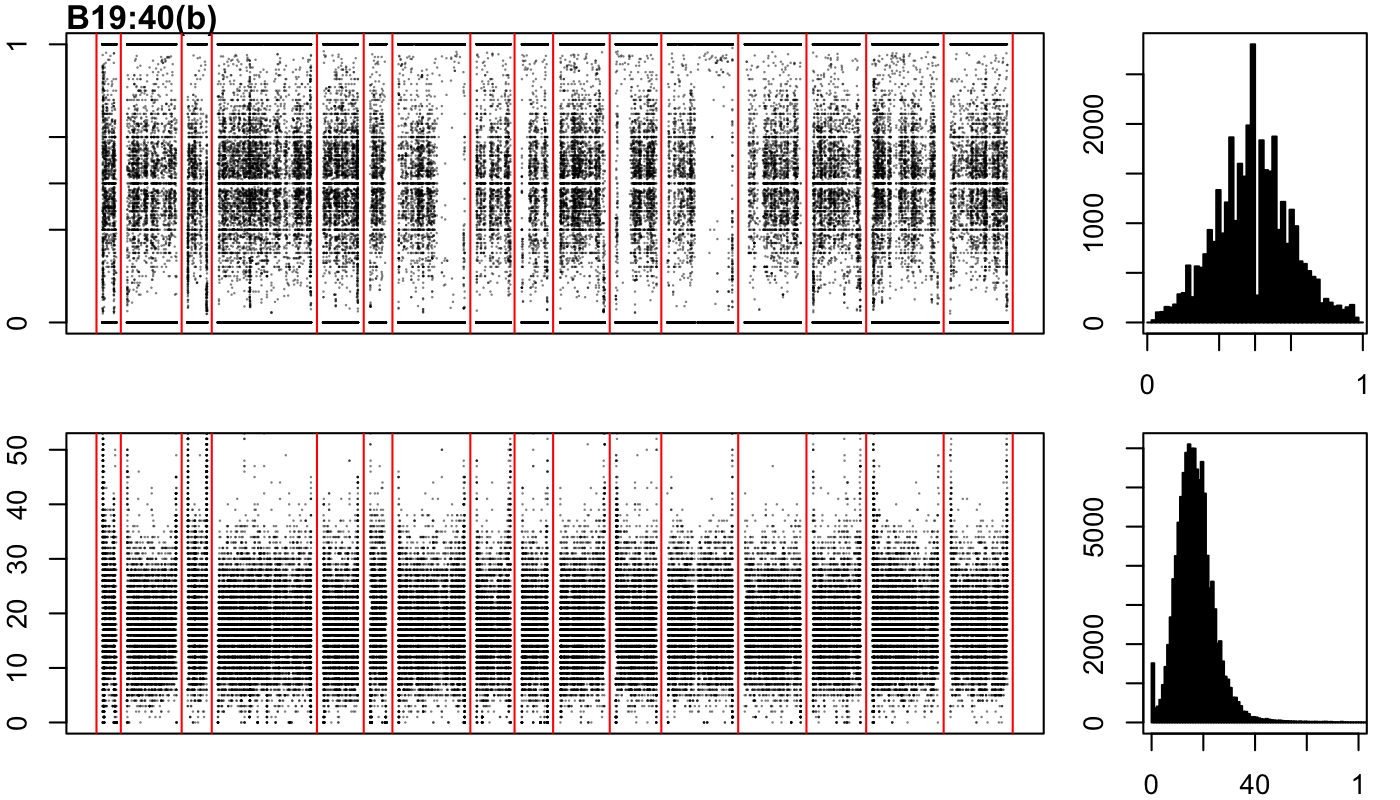

### B19-54-a.png

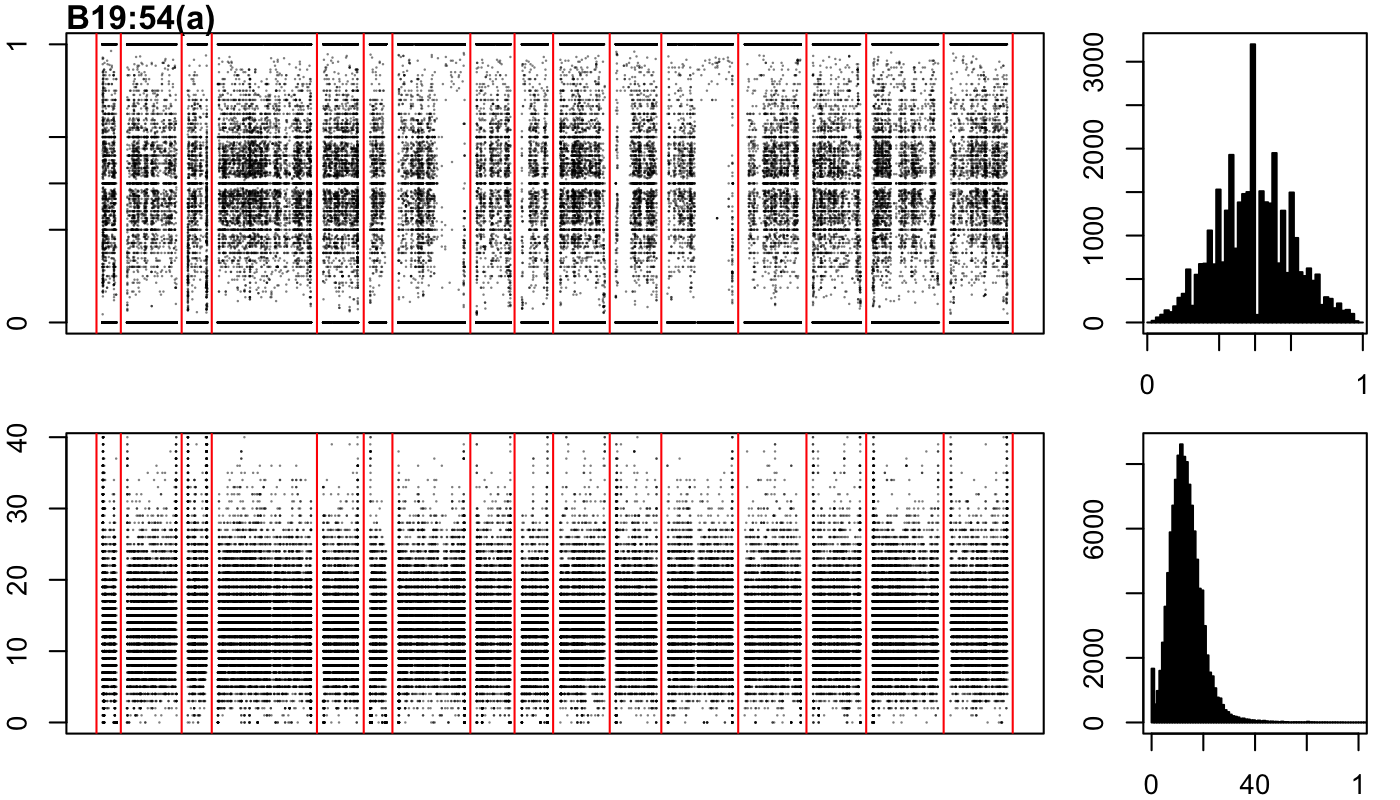

### B19-54-b.png

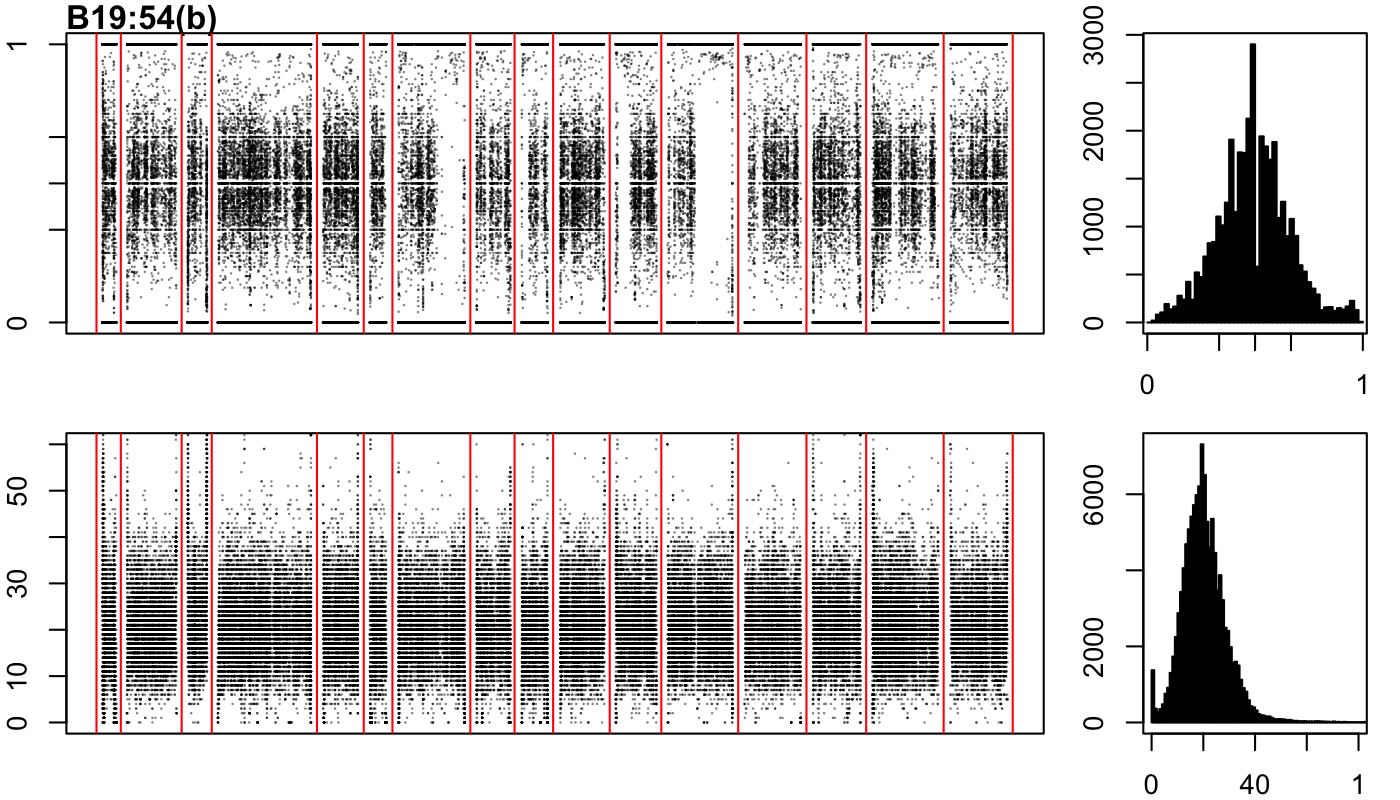

### B19-95-a.png

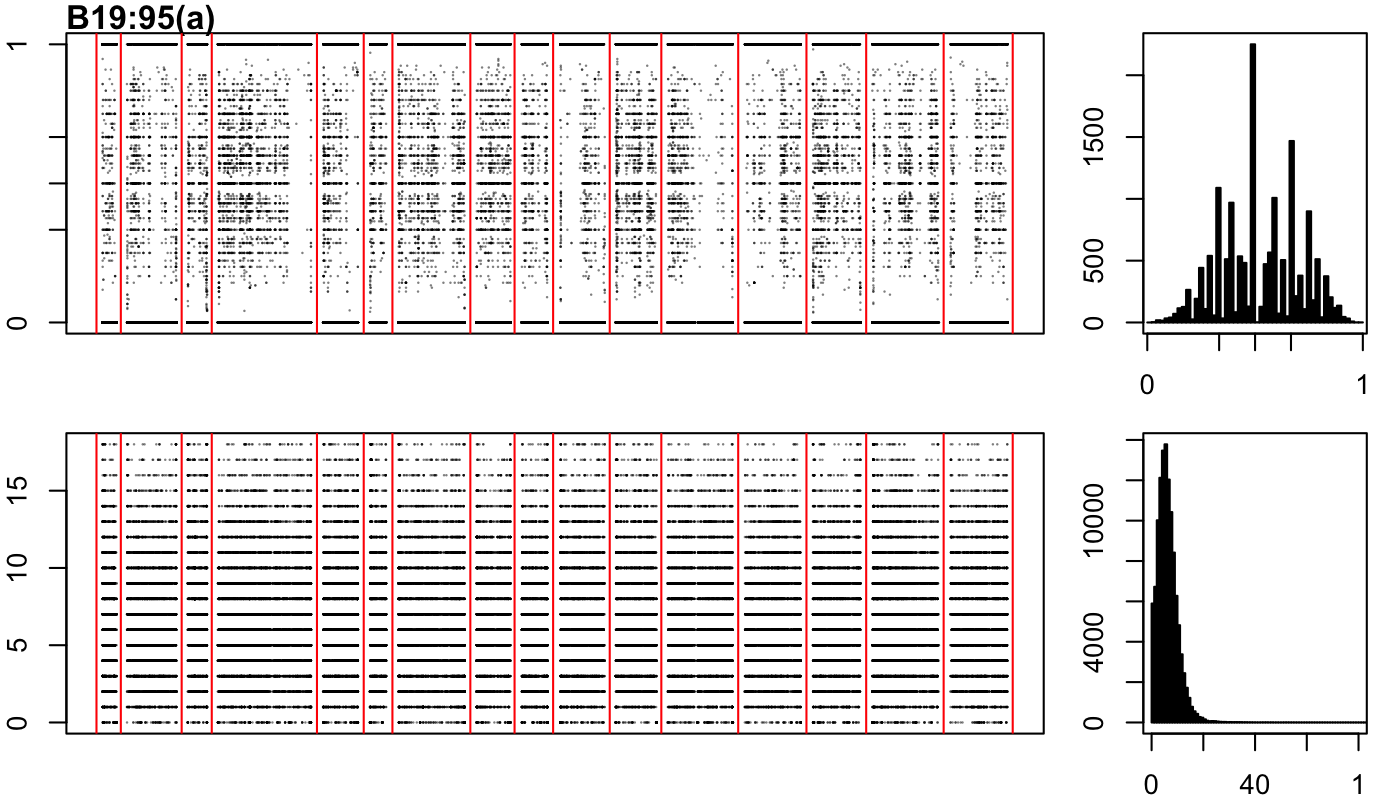

### B19-95-b.png

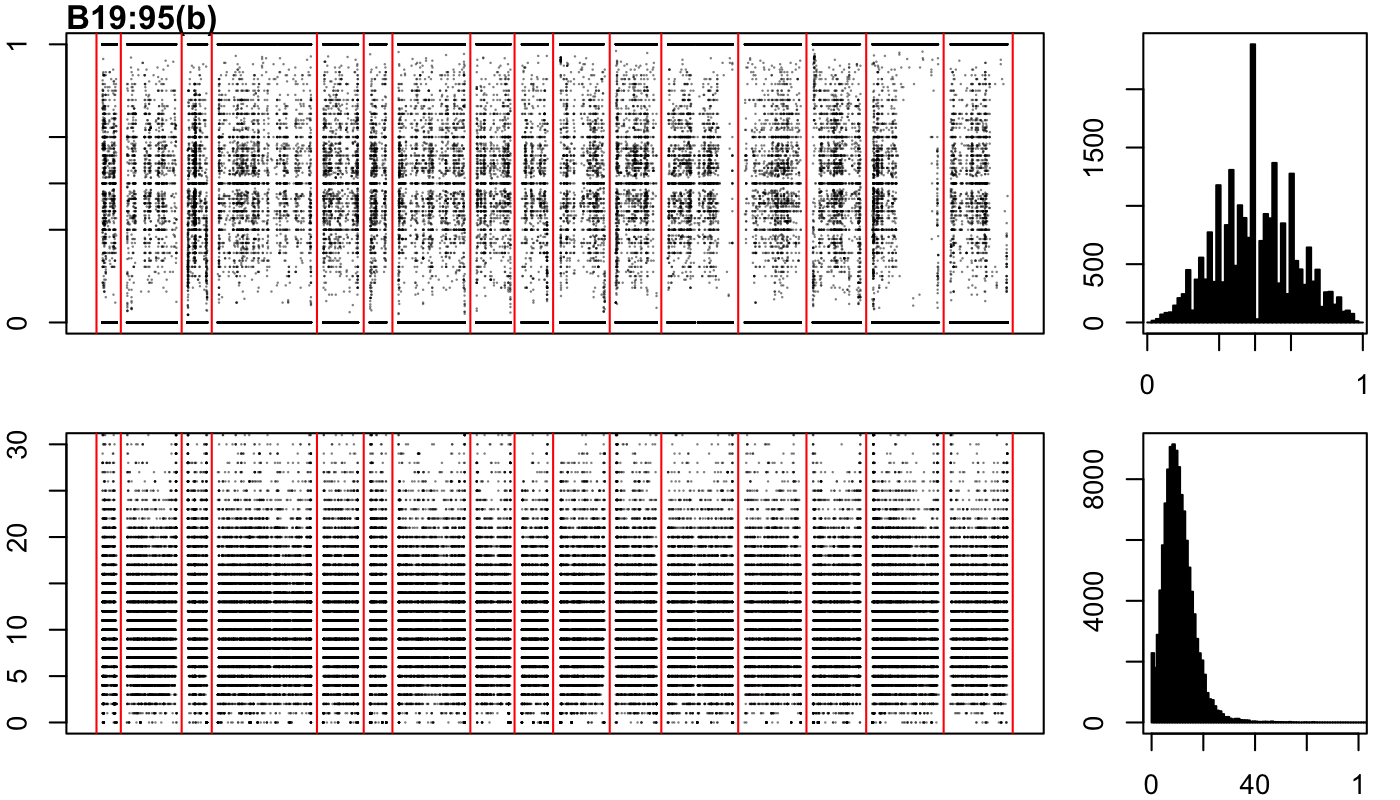

### B19-109-a.png

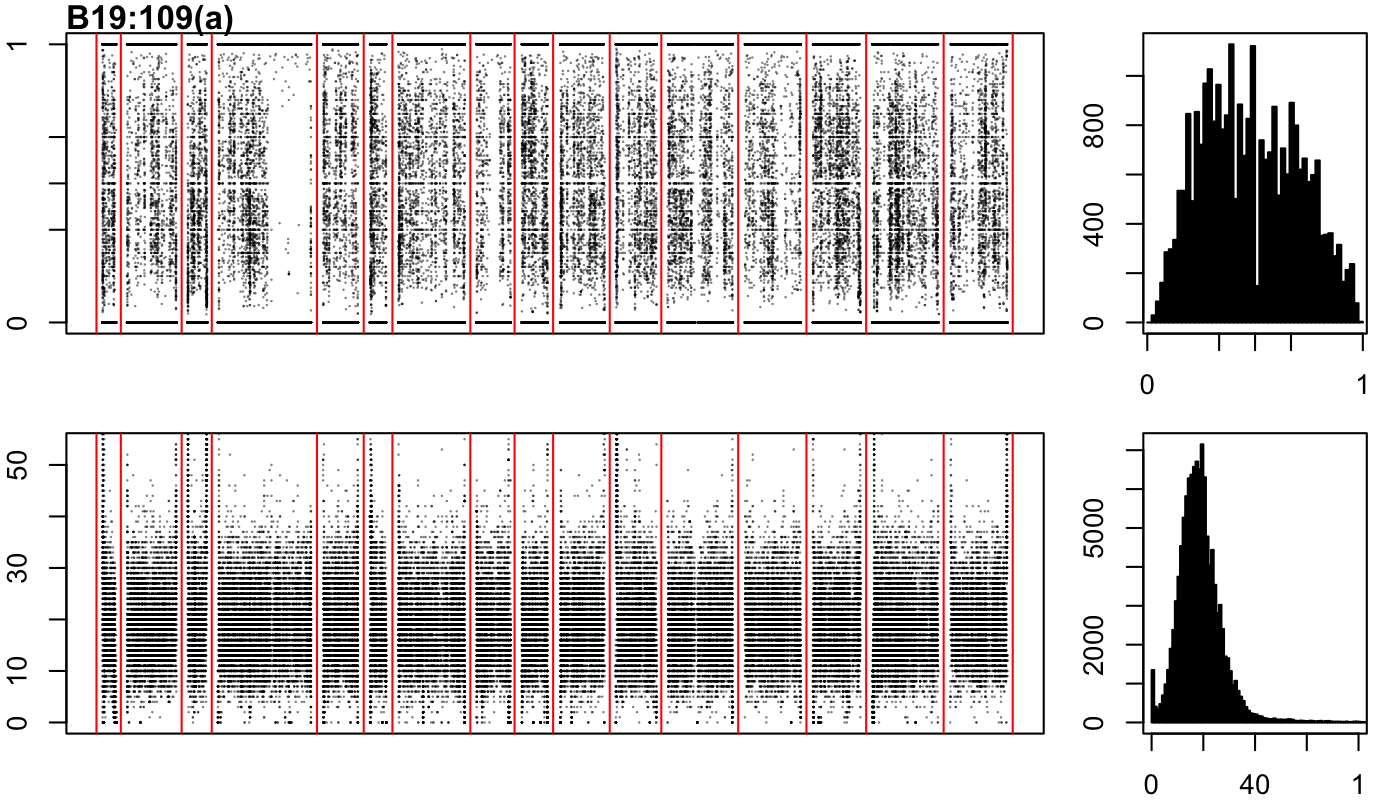

### B19-109-b.png

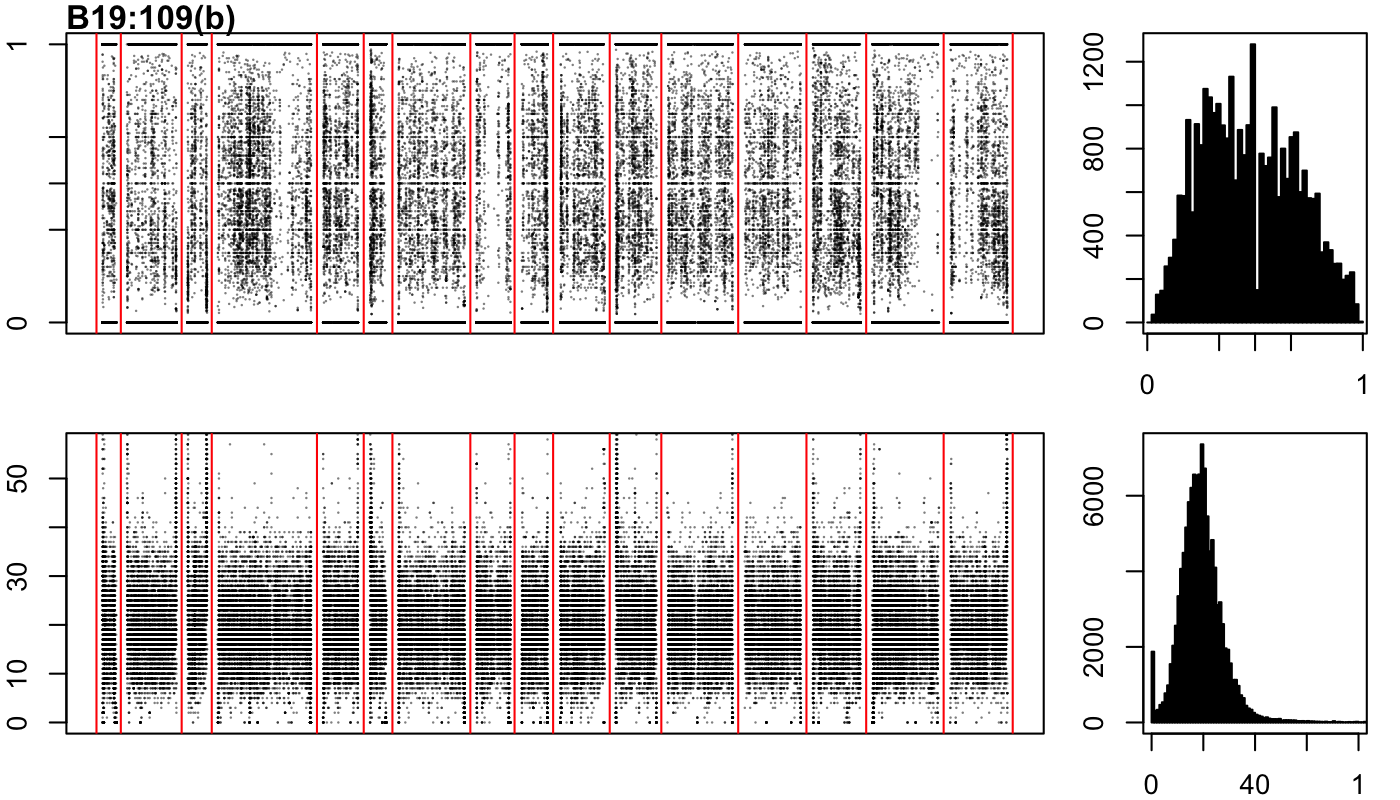

### B19-151-a.png

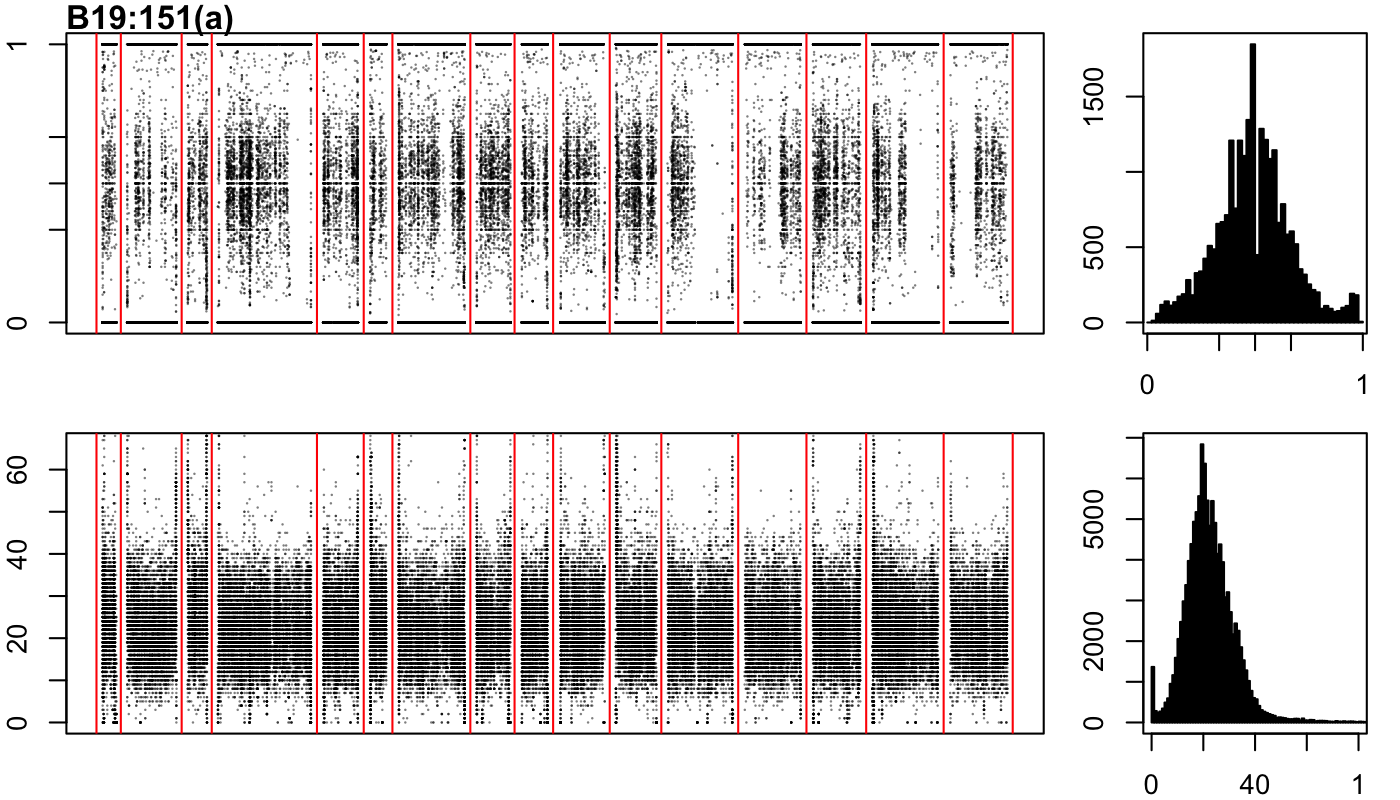

### B19-184-a.png

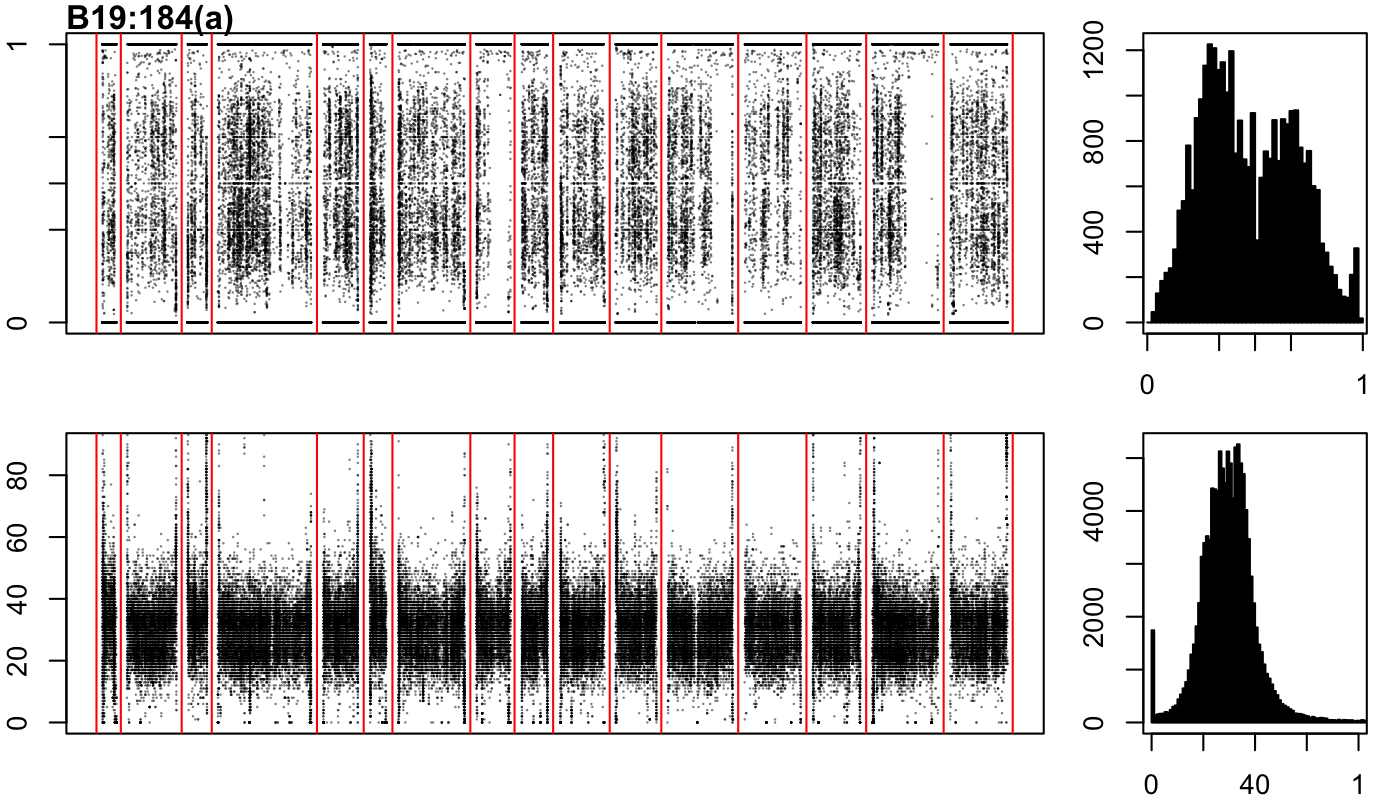

### B19-184-b.png

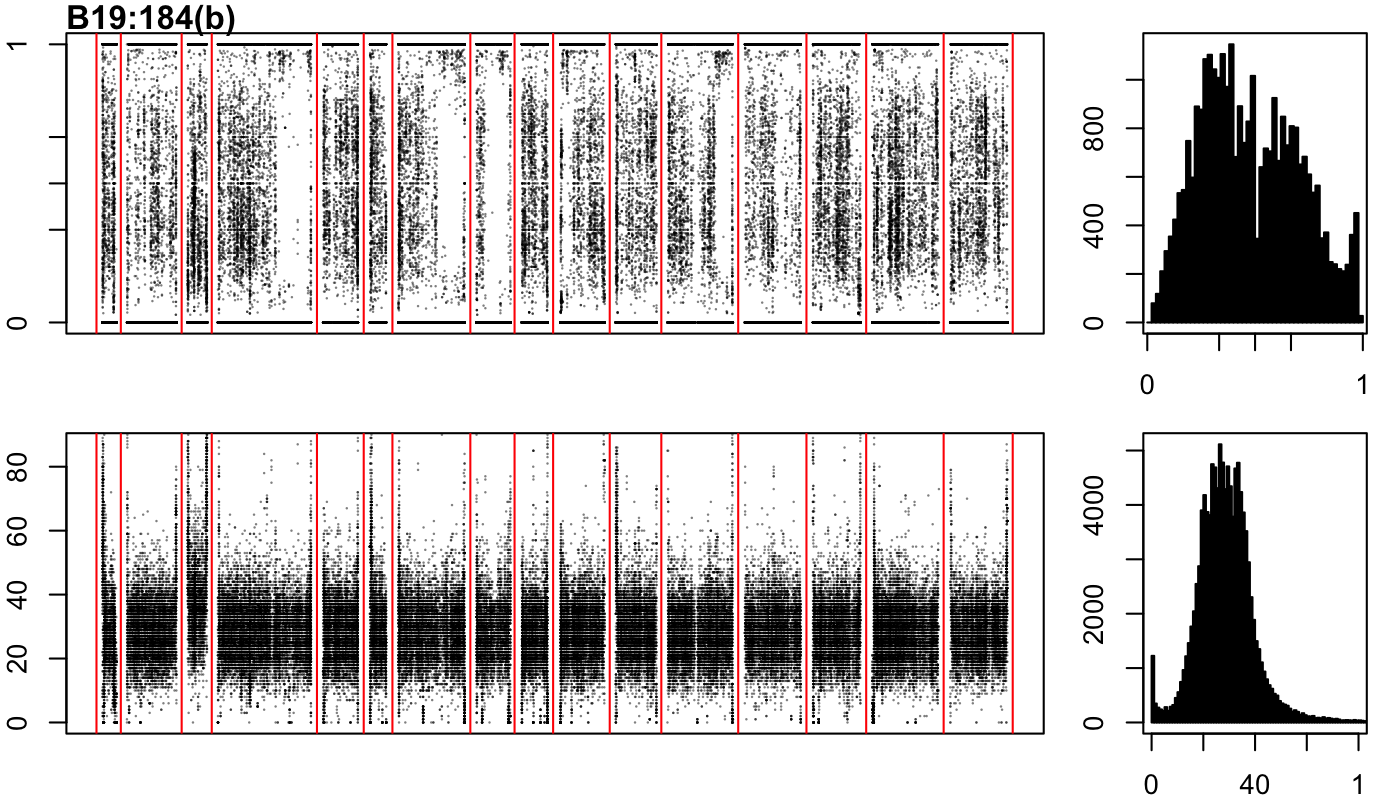

### B19-184-c.png

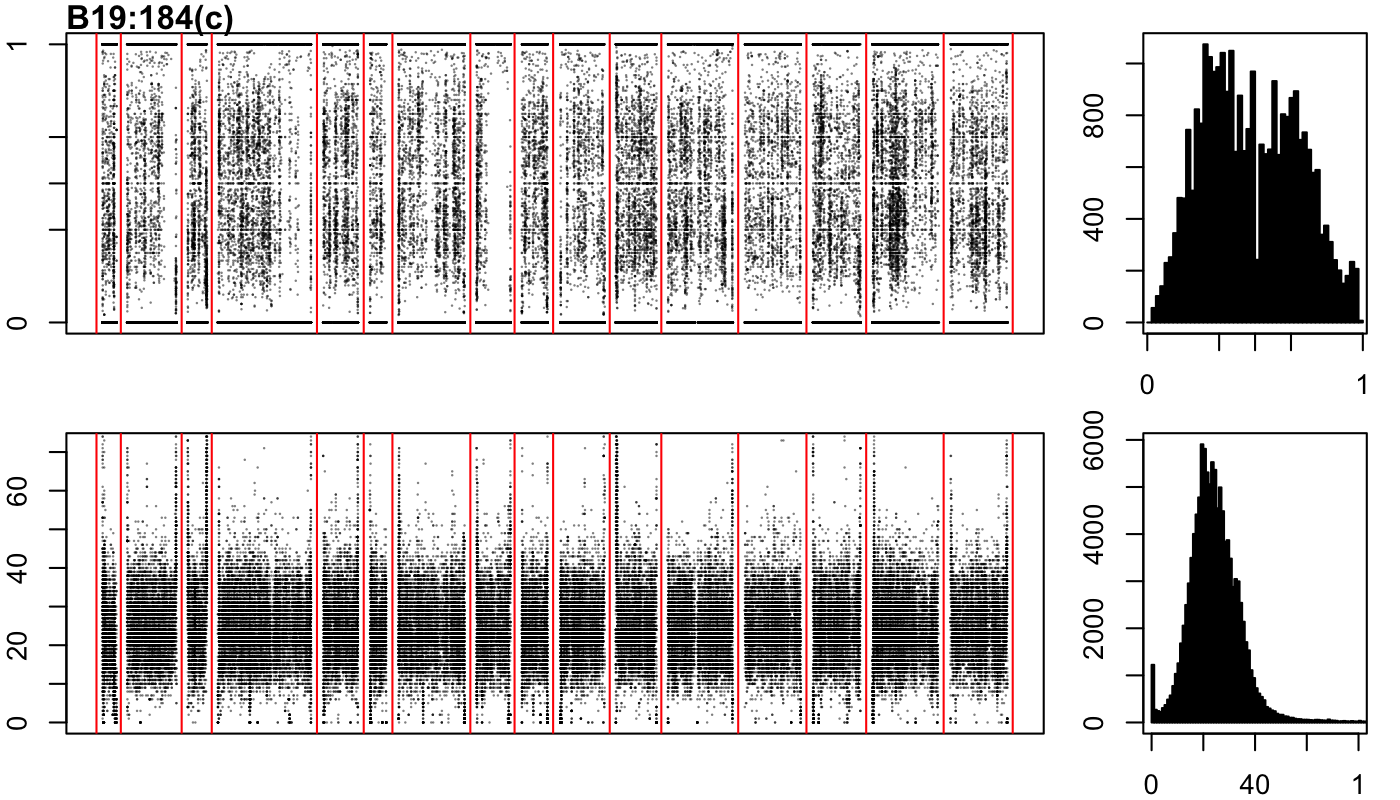

### FT-858-a.png

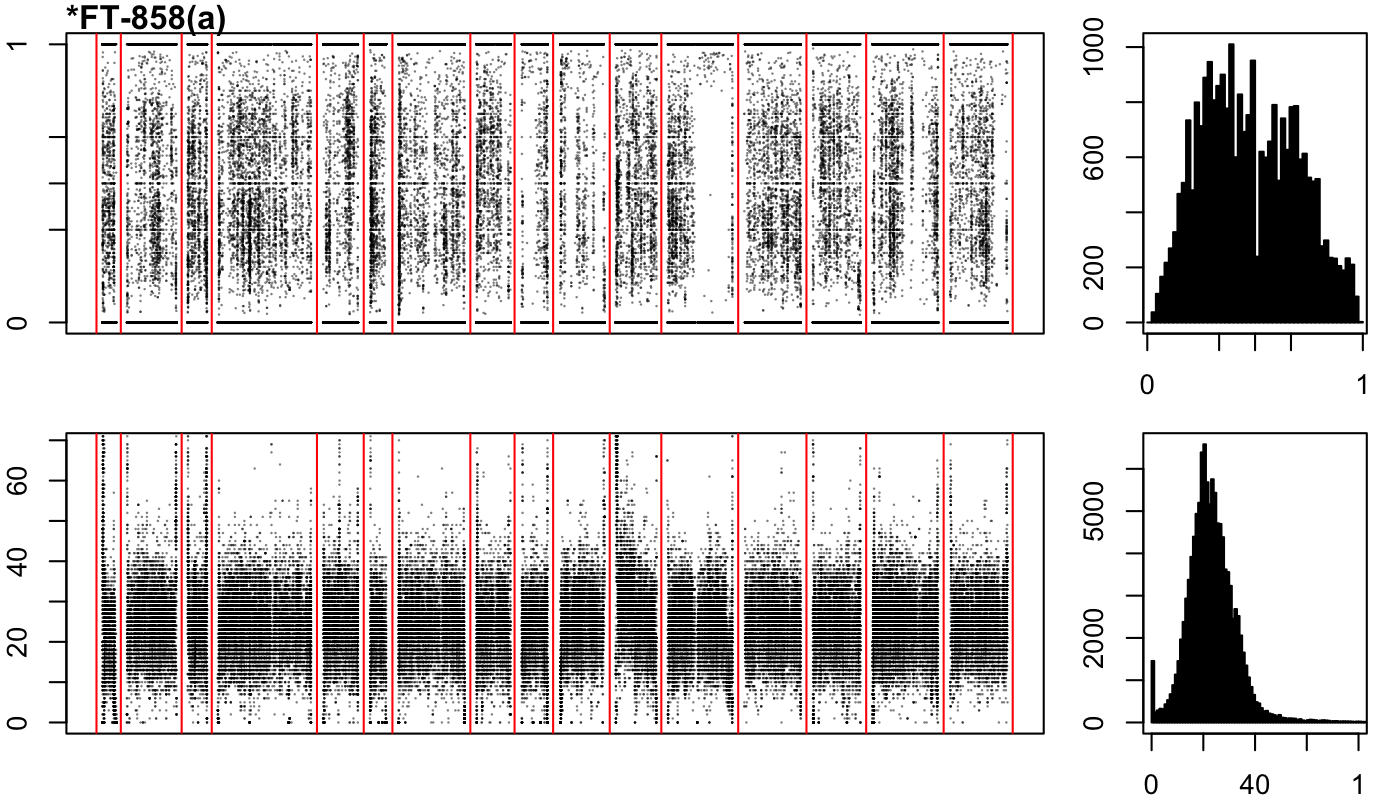

### FT-858-b.png

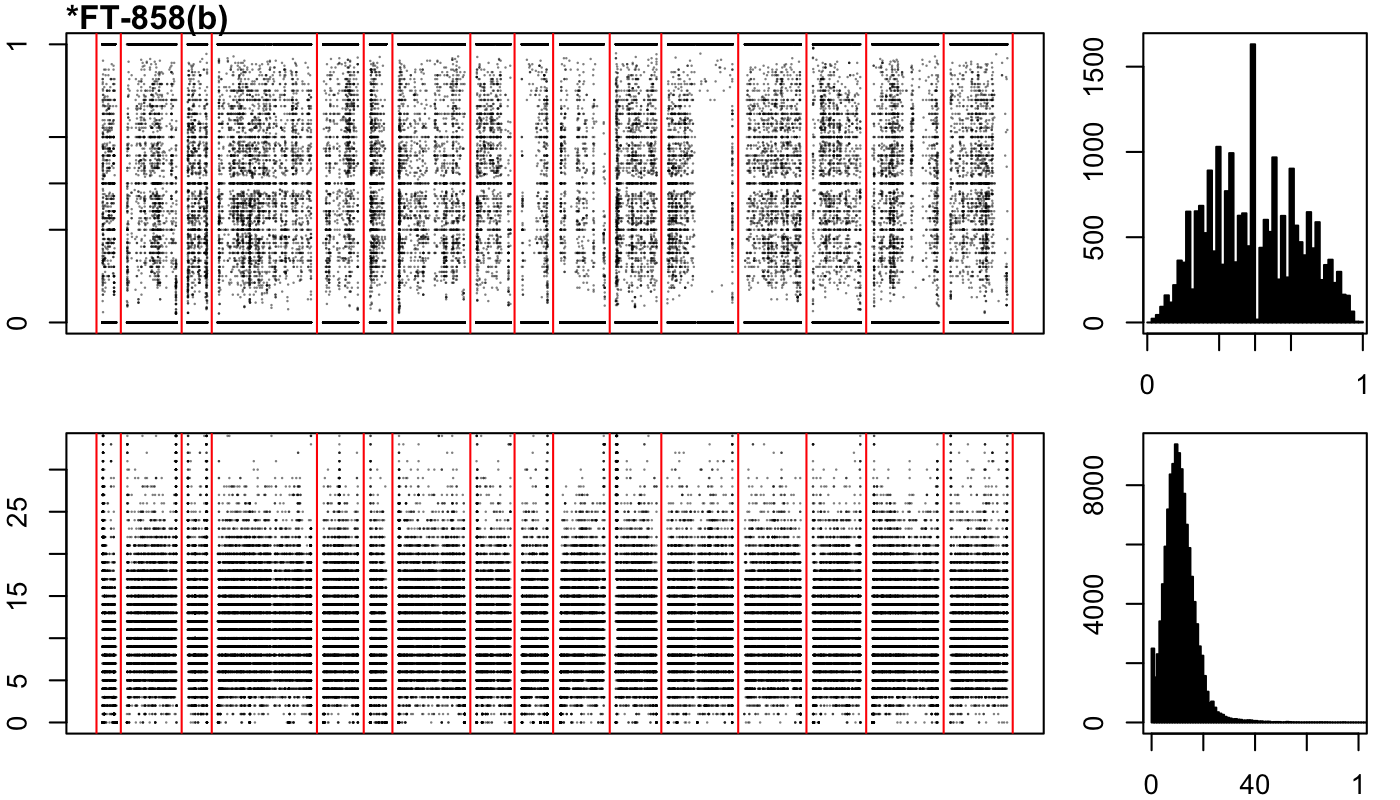

### FT-858-c.png

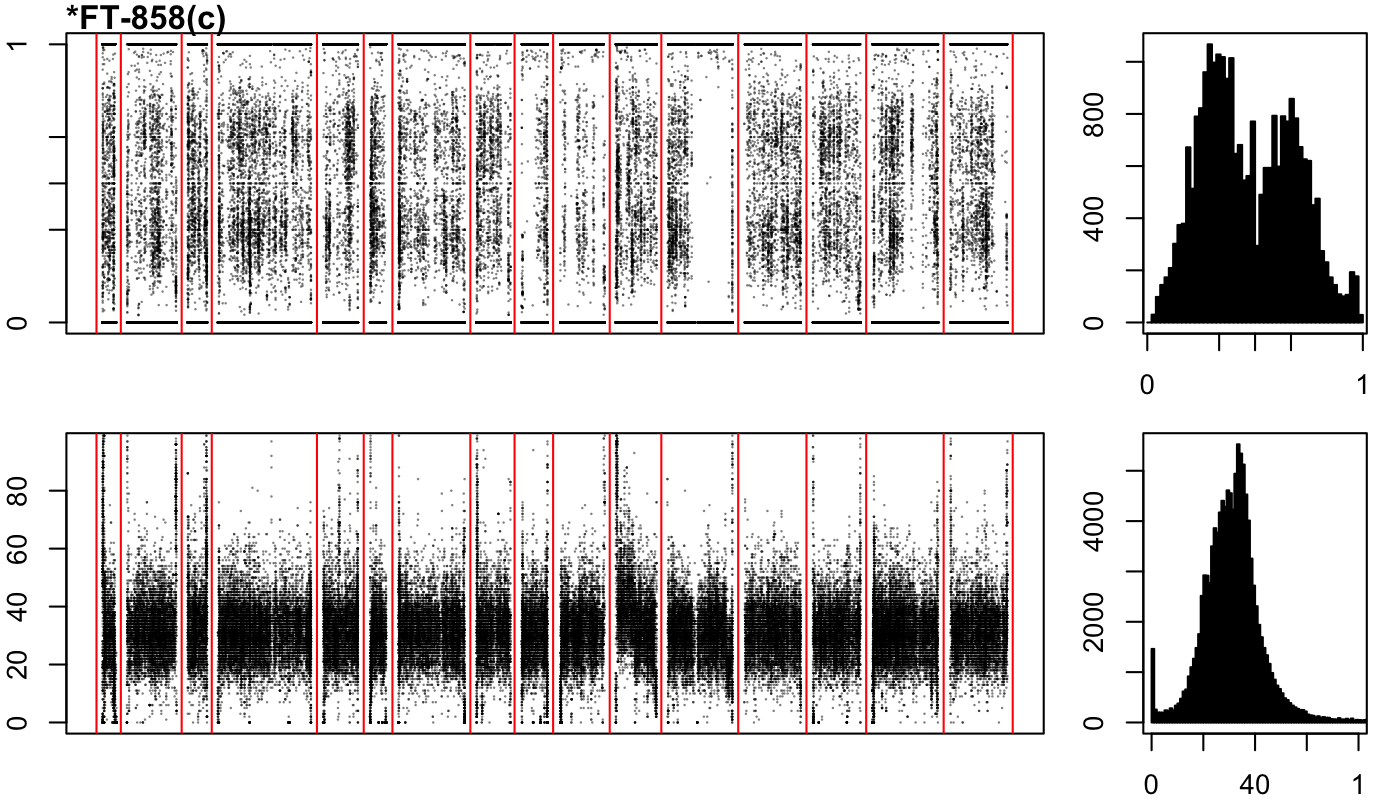

### IRA-D-a.png

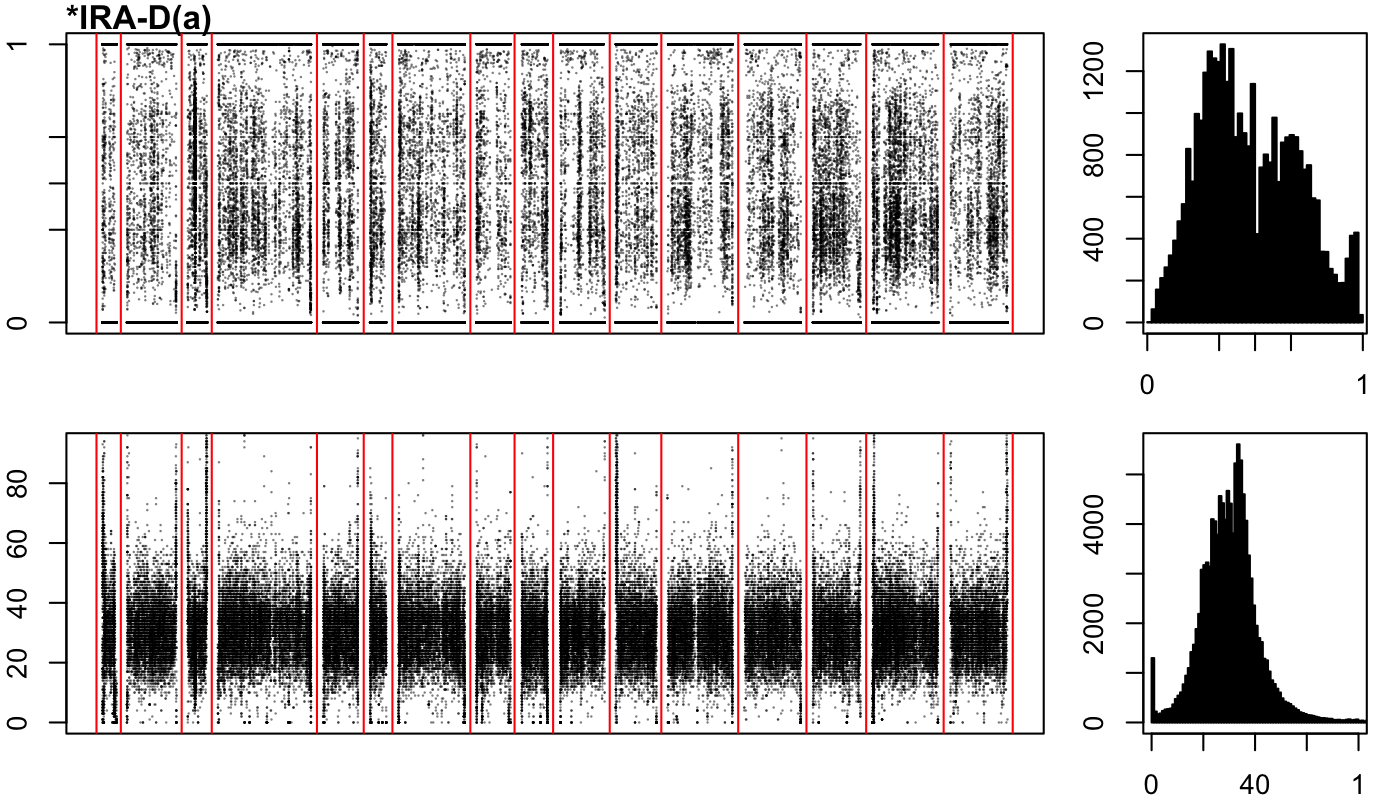

### IRA-D-b.png

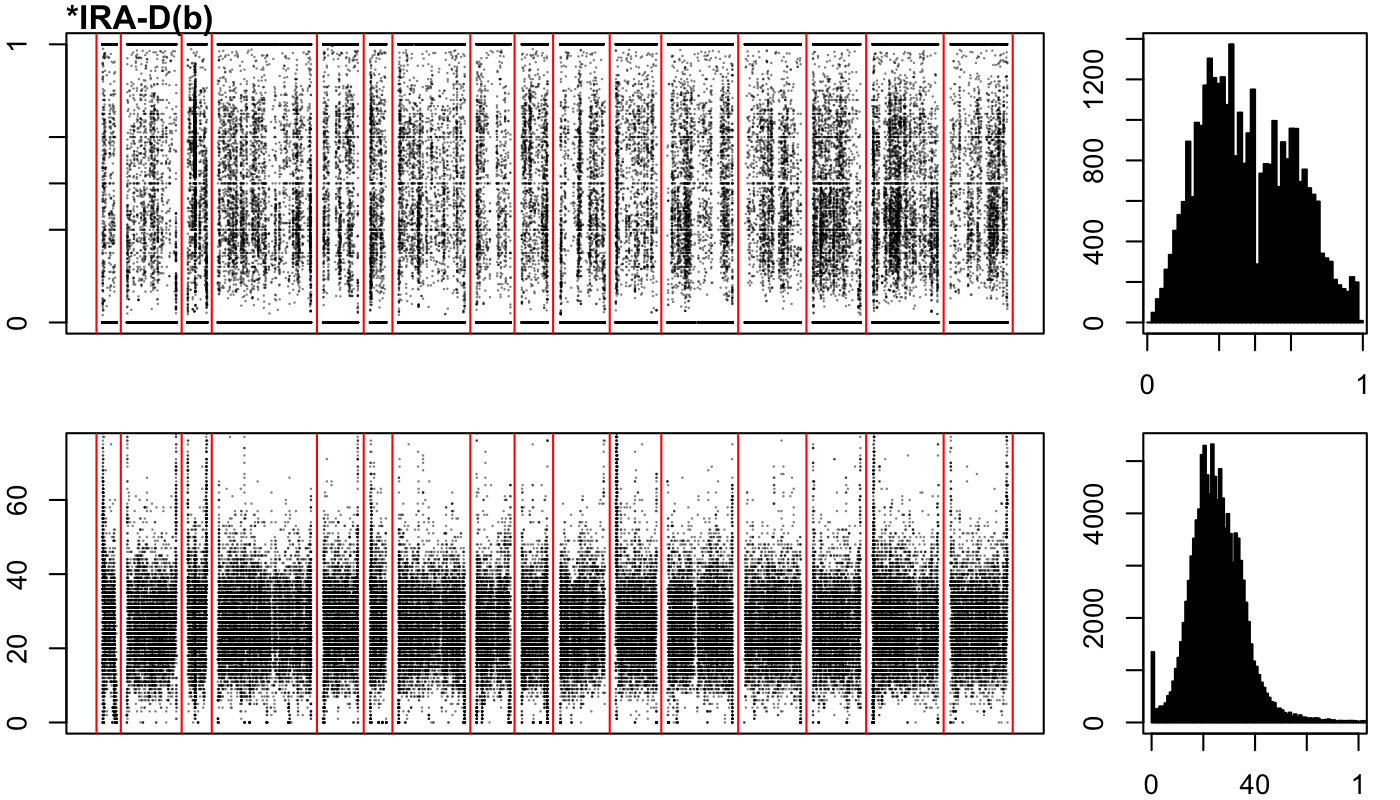

### IRA-D-c.png

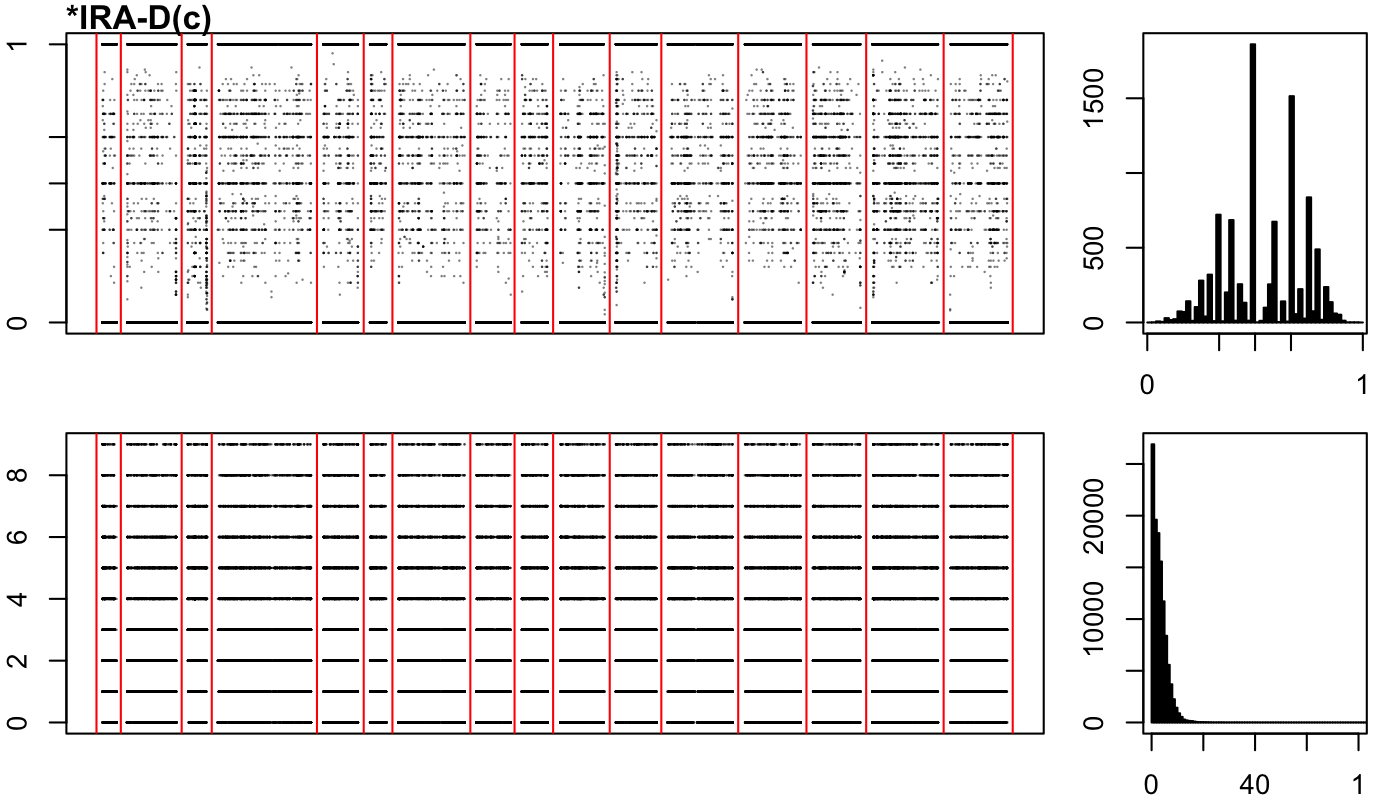

### PE-2-a.png

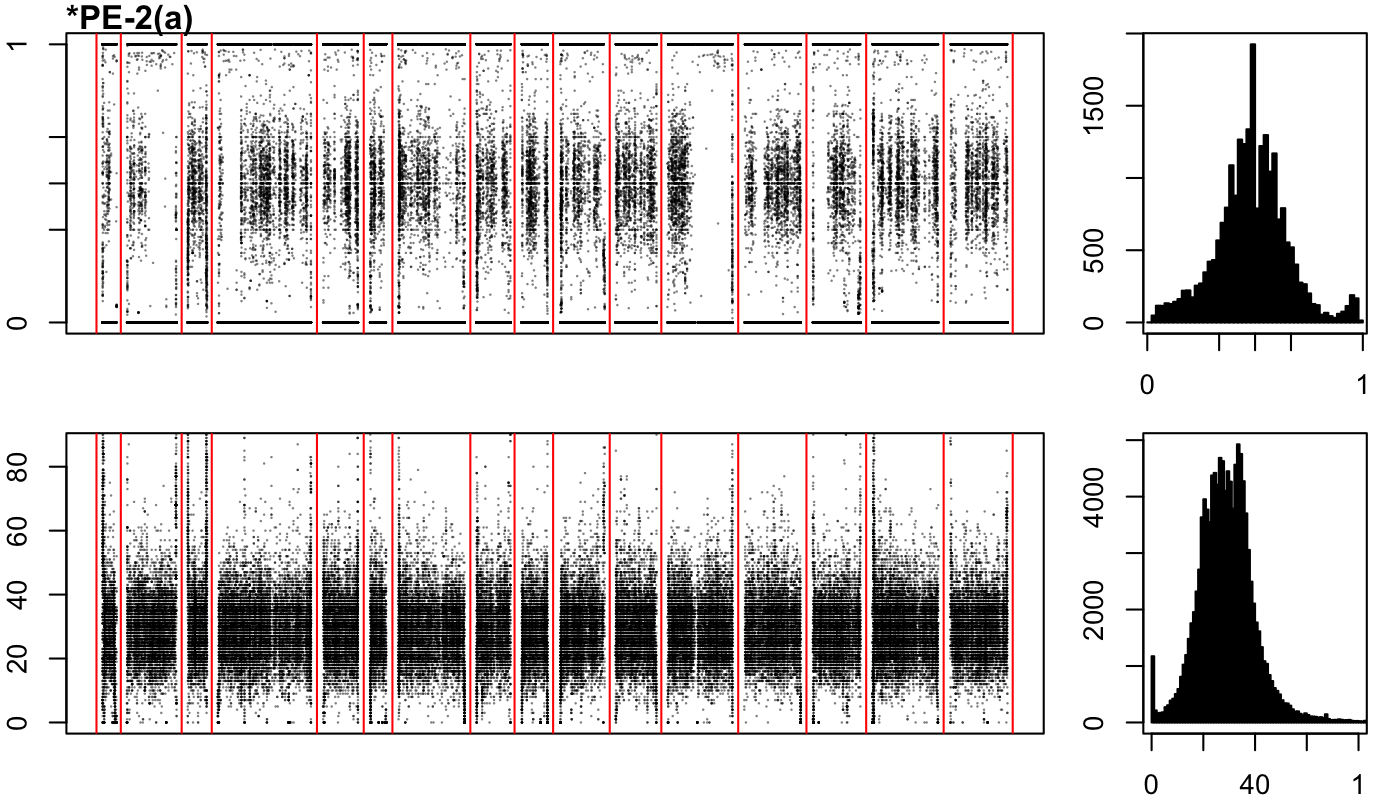

### PE-2-b.png

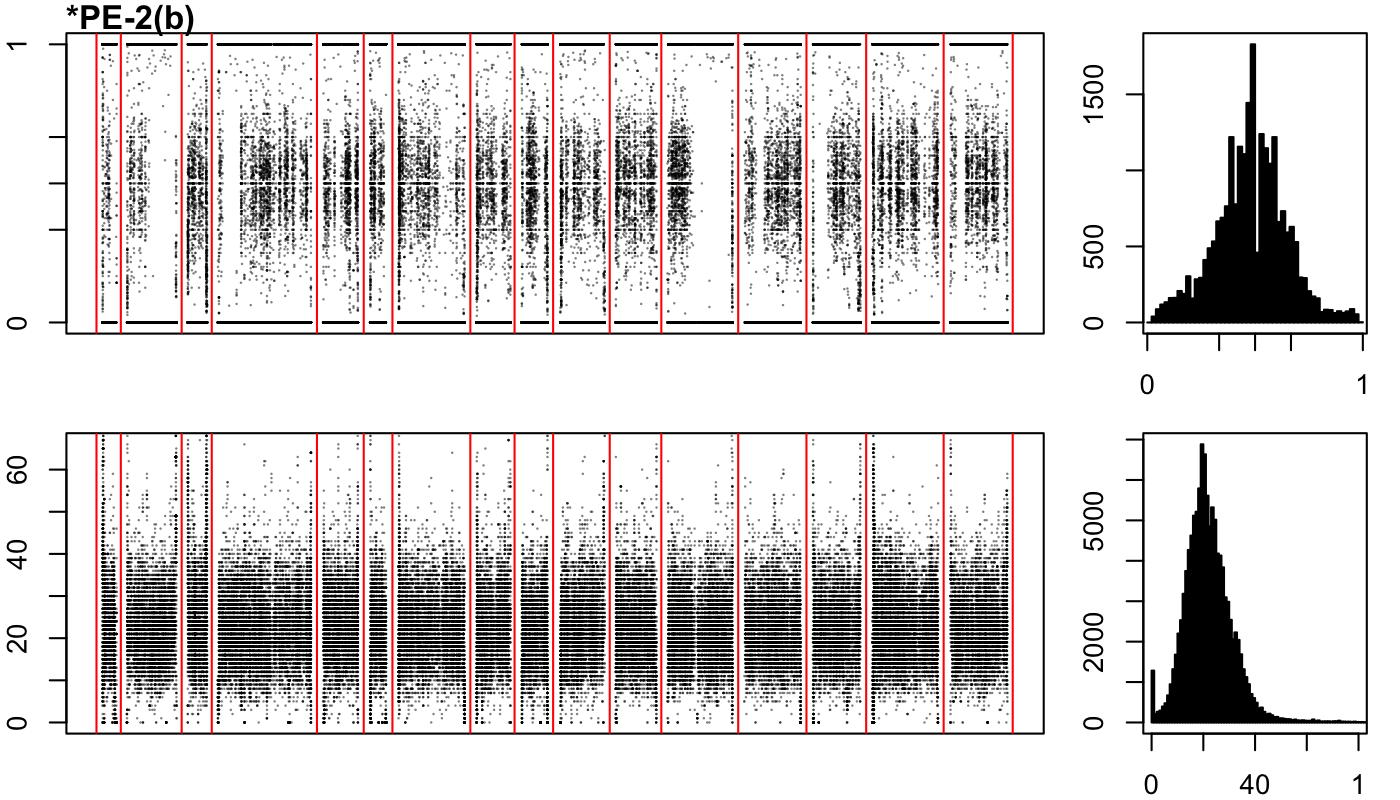

### PE-2-c.png

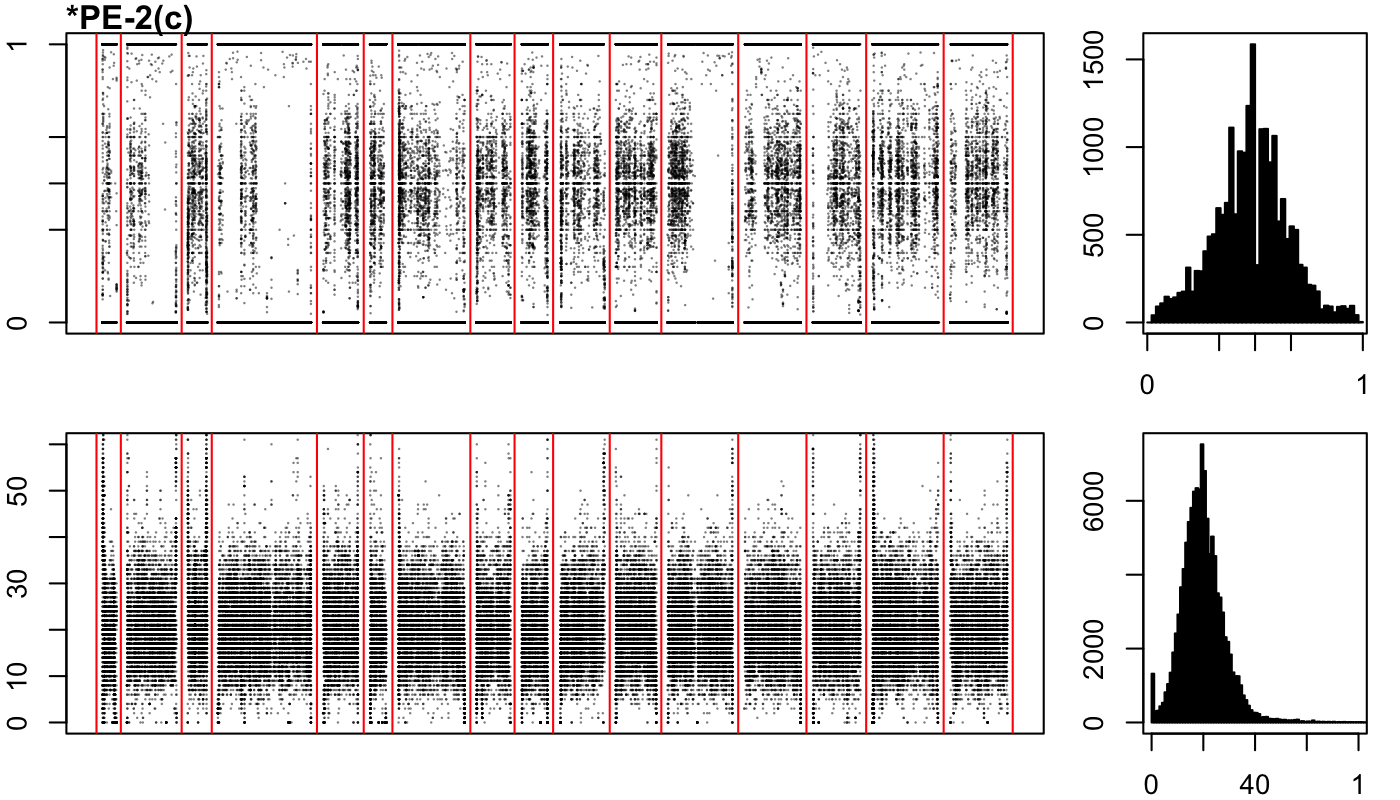

### SA-1-a.png

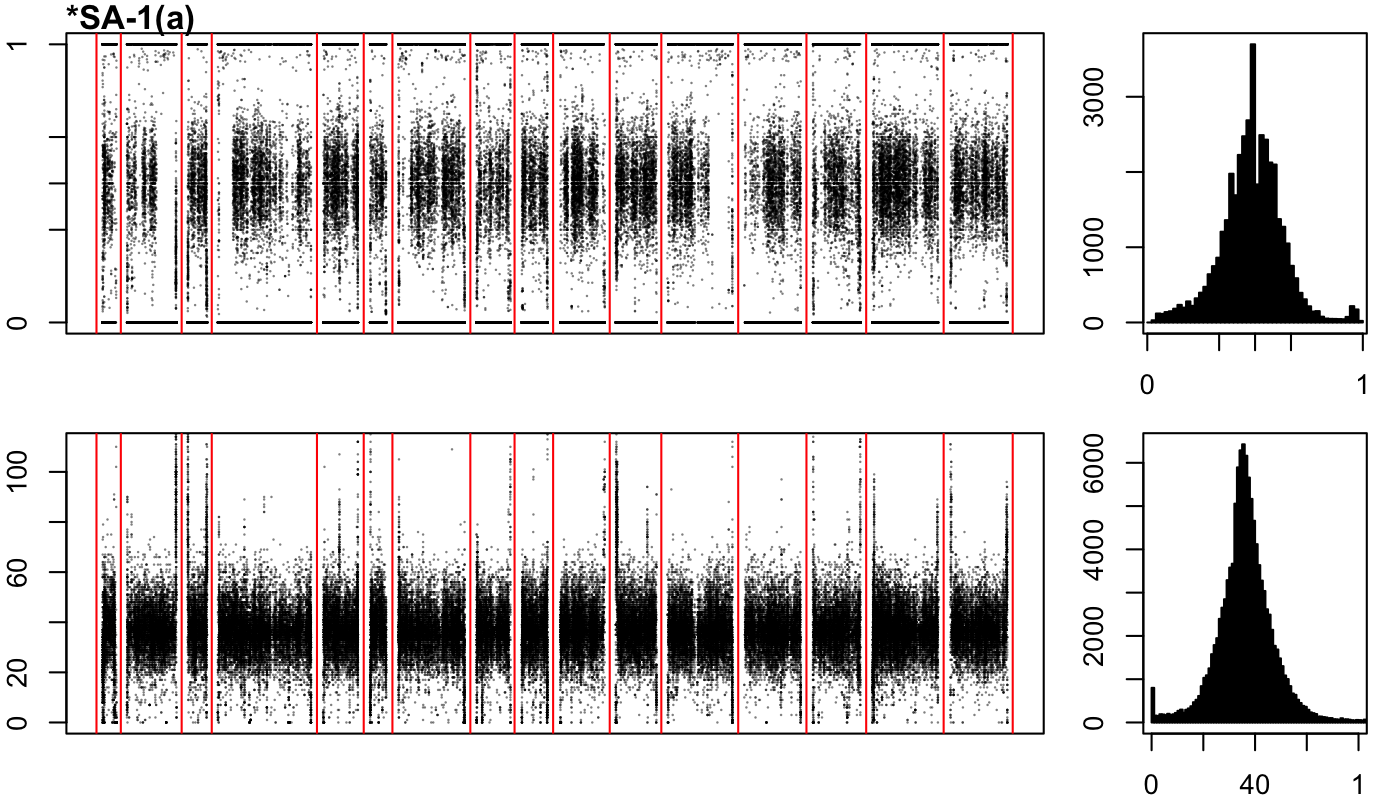

### SA-1-b.png

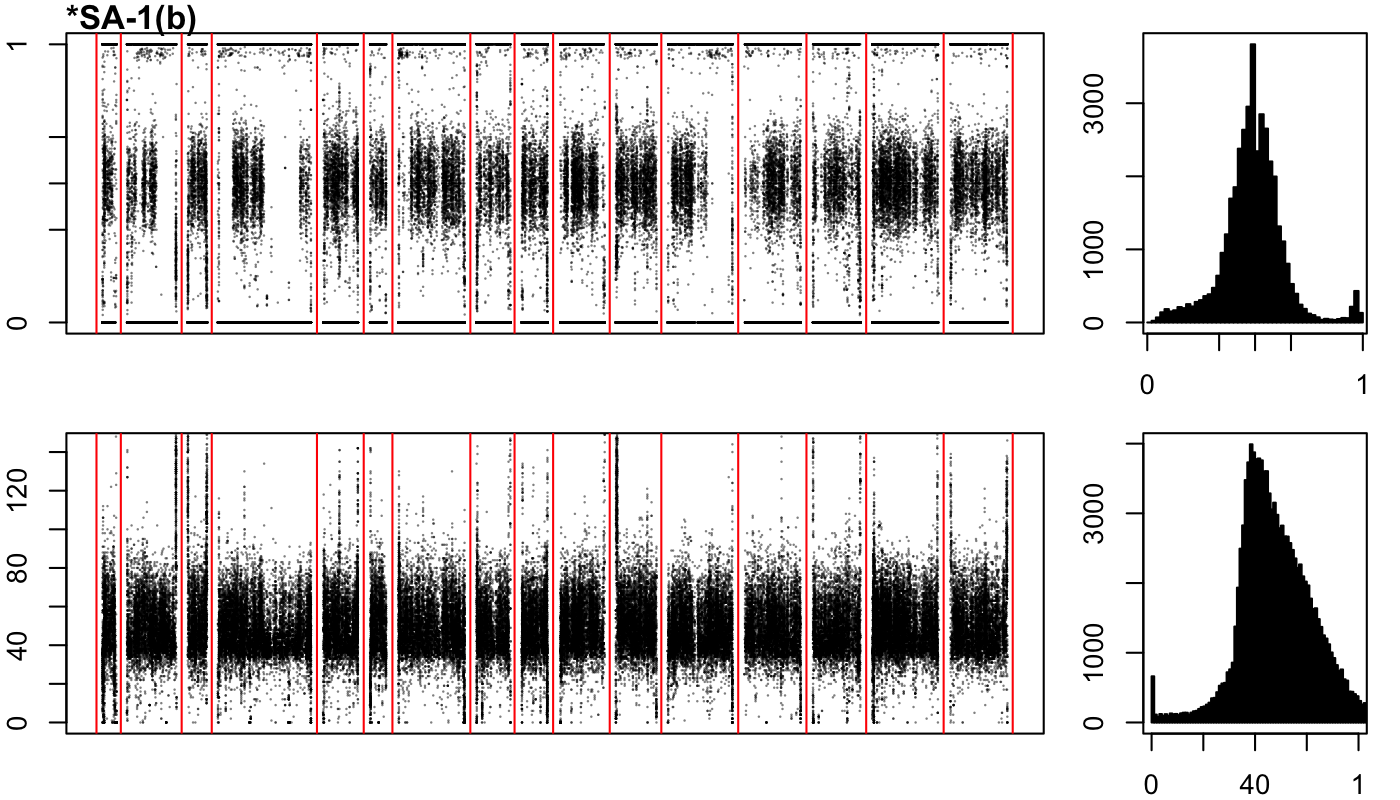

### SA-1-c.png

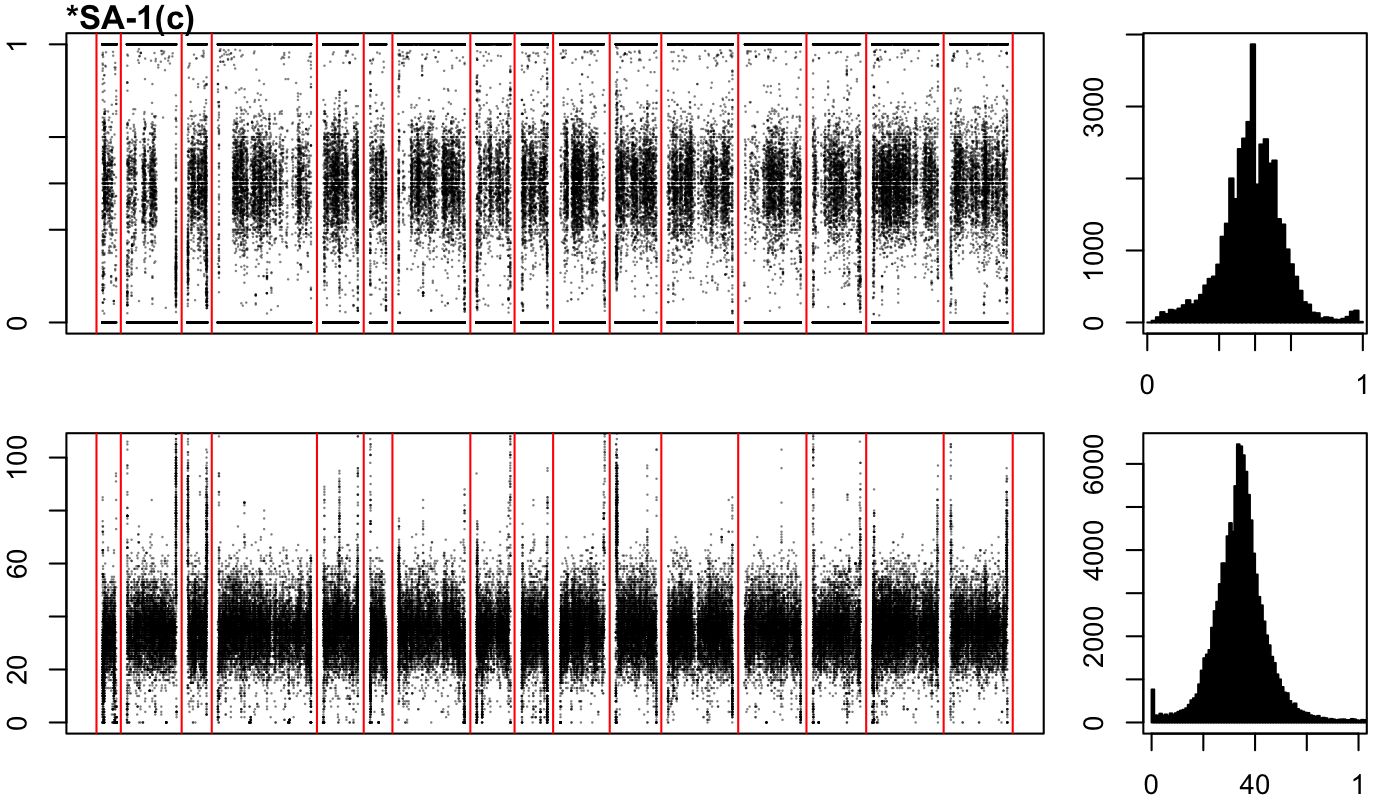
